# A mesoscale cortical lattice

**DOI:** 10.64898/2026.11.24.768659

**Authors:** Melanie Boly, Myeongjin Jeong, Lorenzo Murace, Kort Driessen, Graham Findlay, Larissa Albantakis, Rasmus M. Birn, Chiara Cirelli, William Marshall, Shuntaro Sasai, Olaf Sporns, Giulio Tononi

**Author notes:** Correspondence (M.B.), (G.T.).

## Abstract

Graph-theoretic analyses of functional connectivity have informed our understanding of cortical network architecture primarily at the macroscale of cortical parcels. To gain insights into whole-brain, near-mesoscale organization, we examined vertex-level resting-state fMRI correlation matrices, focusing on strong, short-range (3–20 mm) functional connections. In 151 Human Connectome Project 7T subjects, these connections reveal a partition between posterior-central and prefrontal cortex along a belt of regions anterior to the central sulcus. The posterior-central block is characterized by dense local connectivity and hierarchical organization (lattice cortex), whereas prefrontal connectivity is more distributed (paralattice cortex). A subset of vertices in higher-level areas of lateral and medial lattice cortex connects it to paralattice cortex. The partition largely corresponds to a basic anatomical subdivision of the underlying thalamus and associated basal ganglia and cerebellar circuits. The partition is preserved during movie watching, and replicates at 3T and in individuals studied with precision fMRI.

## INTRODUCTION

From its inception, the study of the human connectome has envisioned multiple scales of organization: a macroscale of cortical regions and areas, a mesoscale of cortical columns and local neuronal groups, and a microscale of individual neurons and synapses^1^. In the cerebral cortex, the macroscale captures the inter-areal, or parcel-level, wiring diagram and has been extensively mapped in humans and other mammals at both structural and functional levels. At the other extreme, the microscale is beginning to reveal the local circuitry of individual columns, largely through recent large-scale electron-microscopy reconstructions^2,3^. The intermediate mesoscale has been investigated primarily through tracing and electrophysiological studies in animal models^4–6^ and through some voxel-level functional magnetic resonance imaging (fMRI)^7–9^, and, in animals, through focal stimulation combined with high-field fMRI^10^. This scale approaches that of cortical columns and local groups of neurons, which have long been proposed as fundamental units of cortical organization^11–14^. Strong, short-range interactions among local neuronal groups, whose neurons tend to share inputs and outputs, may support a highly integrated, lattice-like architecture, distinct from the more distributed, flexible organization mediated by long-range interactions^15,16^. Here we take advantage of curated surface data sampled at individual vertices (point locations along the folded cortical sheet), primarily from the Human Connectome Project (HCP)^17^, to investigate whole-brain cortical functional connectivity at near-mesoscale resolution. We focus on strong, short-range functional connectivity that may reflect lateral gray-matter interactions within cortical areas and short U-fibers linking neighboring areas. This is the range in which anatomical connectivity is densest, given that cortical connection probability falls approximately exponentially with distance^6,18,19^. Throughout, analyses are performed at the level of individual cortical vertices rather than predefined parcels or areas, as mesoscale functional organization may not conform to conventional areal or cytoarchitectonic boundaries^20–22^.

Our primary analysis uses vertex-level resting-state fMRI correlation matrices from 151 HCP subjects scanned at 7 Tesla (7T; 1.6 mm isotropic resolution; 59,412 cortical vertices on the standard fs_LR surface after removal of the medial wall). After masking peri-allocortex, spectral decomposition of strong, short-range connectivity reveals a subdivision of the cortical mantle into two blocks. The larger block, which we term lattice cortex, encompasses posterior-central regions and is distinguished by dense local correlations and multimodal, hierarchical organization. Further analyses and replication across additional datasets establish the robustness of this cortical subdivision and its correspondence with subcortical organization.

## RESULTS

We employed resting-state fMRI data from 151 subjects in the HCP 7T cohort, projected onto the fs_LR 32k surface (59,412 cortical vertices; 29,696 in the left hemisphere (L) and 29,716 in the right (R)). Allocortex and peri-allocortex, hereafter referred to collectively as peri-allocortex, were excluded because of their well-established differences in cytoarchitecture and connectivity. These regions comprise agranular limbic, anterior insular and hippocampal cortex, in which layer 4 is essentially absent^23^. A total of 3,536 vertices, 6.0% of the cortical surface, were excluded (full list in Methods).

### Functional connectivity is reduced to vertex-level, strong, short-range edges

To investigate whole-brain cortical functional connectivity at near-mesoscale resolution, we first reduce the cortical connectivity graph to its strong, short-range edges.

#### Functional connectivity is reduced to strong edges

For each subject, we computed the Pearson correlation (r) between the BOLD time series of every pair of vertices. As detailed in the Methods (acquisition parameters in Table S1), we first removed vertex-to-vertex correlations unlikely to reflect direct neural interactions by estimating a subject-specific null floor τ_s_, which a correlation must exceed to count as supported in that subject. The null distribution is estimated from each subject’s correlation values between occipital cortex vertices and contralateral frontal pole and sensorimotor cortex vertices, pairs that have no plausible direct connection, and τ_s_ is defined, separately for each subject, as the smallest value at which the estimated false-discovery rate is 5%.

A pair of vertices is then included as an edge of the cohort graph (one graph per hemisphere) if its correlation exceeds τ_s_ in a majority of the 151 subjects (76 or more), and its weight w is the mean of r over the 151 subjects (positive for every edge; Methods). In an individual subject, on average 11.5% of vertex pairs within a hemisphere exceed τs (Figure 1A, left, and Figure 1B); after the majority rule the cohort graph retains 5.6% of all pairs (5.4% L and 5.7% R; Figure 1A, right). Thus, the edges of the cohort graph capture correlations that are strong across a majority of subjects.

**Figure 1.**
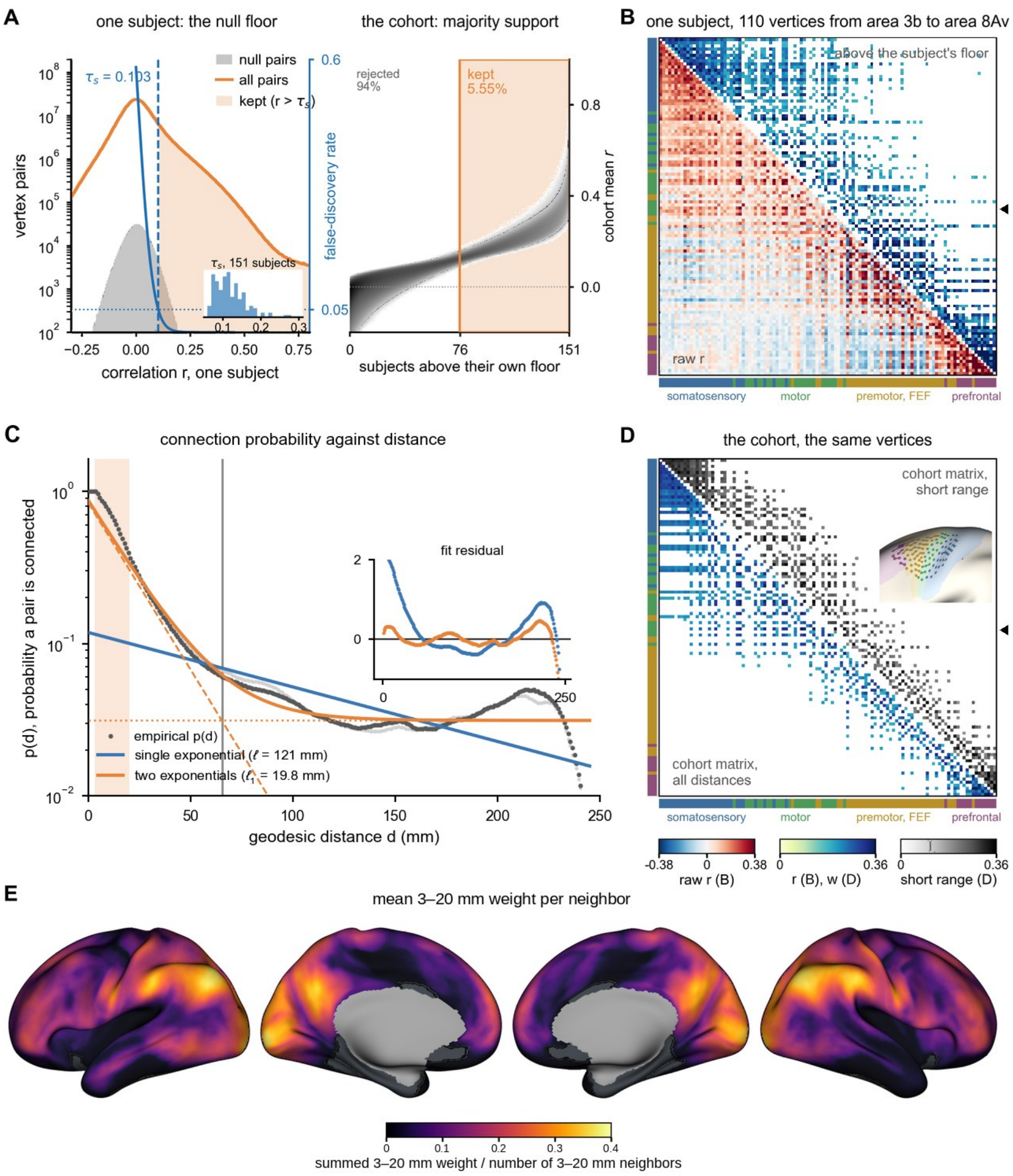
Construction of the cohort graph: strong edge selection and definition of the short-range window. **(A)** Strong edge selection. Left: for one representative subject (572045), the distribution of the correlation r over the subject’s within-hemisphere vertex pairs (orange) and over the null pairs (gray). The subject’s null floor τ_s_ (dashed line; 0.103) is set so that no more than 5% of the 3–20 mm pairs above it could have passed by chance (Methods); orange shading, the pairs kept. Inset: τ_s_ across the 151 subjects (median 0.113). Right: for every within-hemisphere vertex pair, the number of subjects in which r exceeds their own floor against the mean r over the 151 subjects (gray, log counts). Pairs above the floor in a majority of subjects (shaded) are edges of the cohort graph. **(B)** Correlation matrix of the same subject for 110 left-hemisphere vertices on a strip running from area 3b through motor and premotor cortex to area 8Av. Lower triangle, r; upper triangle, r of the pairs above the subject’s null floor. Colored bands, somatosensory, motor, premotor and prefrontal areas; arrowheads, the border between the two blocks of Figure 2B. **(C)** Short-range window. Edge probability p(d), the fraction of vertex pairs at geodesic distance d that are edges of the cohort graph (area-weighted; dark gray, left hemisphere; light gray, right), with single-exponential (blue) and two-exponential (orange) fits (left hemisphere). The two-exponential fit separates a short-range component (dashed) from a distance-independent long-range one (dotted). Shading, the 3–20 mm window. Inset: residuals of the two fits. **(D)** The same 110 vertices in the cohort graph. Lower triangle, all distances; upper triangle, the 3–20 mm window. Inset: the strip on the inflated left hemisphere. **(E)** Mean weight per 3–20 mm neighbor (summed edge weight divided by the number of neighbors within 3–20 mm), lateral and medial views of both hemispheres. The peri-allocortex is gray.

#### Functional connectivity contains a distinct short-range component

For each cortical vertex, the cohort graph retains a set of strongly correlated functional partners distributed across the cortex. We next asked whether these edges reflect a single distance-dependent process or a superposition of distinct short- and long-range components. Specifically, we sought to isolate short-range, near-mesoscale correlations that may reflect local interactions mediated by dense intra-gray-matter connections and short U-fibers linking neighboring cortical areas.

Within each hemisphere, the distance d between two vertices was defined as the shortest path along the folded mid-thickness surface (the exact geodesic), using the median distance across the 151 subjects’ individual surfaces. We quantified the edge probability p(d), defined as the fraction of vertex pairs within each 1-mm distance bin that are edges of the cohort graph (each pair weighted by the cortical area of its two vertices; Methods), separately for each hemisphere. We compared two models for p(d): a single exponential, p(d) = A·exp(−d/*l*), with the sum of two exponentials, p(d) = A_1_·exp(−d/*l*_1_) + A_2_·exp(−d/*l*_2_). Each *l* is a length constant, the distance over which a component falls by a factor of *e*, and each A is the component’s amplitude, its value extrapolated to d = 0. Both were fit by least squares to ln p(d), each 1-mm bin weighted by its area-weighted number of pairs (Methods). The two-component model provided a markedly better fit in both hemispheres (Figure 1C; ΔBIC = 355 L, 298 R). The two-component model identified a short-range component with a length constant of *l*_1_ = 19.8 mm L and 22.1 mm R, superimposed on a long-range component that is effectively distance-independent in both hemispheres (A_2_ = 0.031 L and 0.030 R; *l* _2_ → ∞). The two components crossed at d = 65 mm L and 71 mm R. Thus, beyond the initial short-range decay, the probability of an edge does not continue to fall with distance. The increase of p(d) beyond 200 mm, which neither model captures (Figure 1C, inset), comes primarily from correlations linking the angular gyrus and posterior superior temporal sulcus (PGi, PGs, STSvp, TE1p) with dorsomedial and frontopolar prefrontal cortex (9m, 10d, 9p, 9a, 8BL, 8Ad). These exceptionally long-range pairs, the longest in the hemisphere, account for 51% L and 40% R of the edges beyond 200 mm; without them, p(d) stays at the level of the long-range component up to 210 mm (0.028 L and 0.031 R at 200–210 mm) and falls below it at longer distances. On the right, connections of inferior parietal and intraparietal cortex (PGs, PGi, PFm, IP1) with frontopolar areas p10p and a47r also contribute.

Resting-state correlations were also substantially stronger at short than at long distances: the mean correlation between vertices within a hemisphere decreased from 0.41 at 3–4 mm to 0.11 at 19– 20 mm, compared with 0.02 for randomly selected vertex pairs.

#### Short-range connectivity is densest in posterior-central cortex

The following analyses focus on short-range functional connectivity, defined as the geodesic-distance window 3 ≤ d < 20 mm. The inner bound (3 mm) was chosen to exclude the range in which the hemodynamic point-spread function and the projection from volume to surface can produce spurious correlations (see Supplementary Results S1). The outer bound (20 mm) lies at approximately one short-range length constant (1.0 *l*_1_ L, 0.9 *l*_1_ R). At that distance, the short-range component still exceeds the long-range component by about tenfold in both hemispheres (9.8 L and 10.2 R), well short of the 65 mm L and 71 mm R crossovers, and 64% L and 60% R of the fitted short-range component falls within 20 mm. The same range also corresponds to the scale of the short U-fibers linking nearby neuronal populations within and across neighboring cortical areas.

Within the 3–20 mm window, the cohort graph keeps 3.48 million L and 3.52 million R edges, 56% L and 57% R of the pairs in the window (Figure 1D). Altogether, the strong, short-range pairs are slightly less than 1% of the total vertex pairs. Out to 4 mm, essentially every pair is connected (p(d) = 0.998 at 3 mm and 0.96 at 4 mm); thereafter, the probability falls steadily, to 0.72 at 10 mm, 0.53 at 15 mm and 0.39 at 19 mm, with values differing by no more than 0.02 between the two hemispheres.

For each vertex, we then summed the weights of its 3–20 mm edges and divided the sum by the number of its neighbors in the window (Methods). This short-range density was highest in the inferior parietal lobule and in early visual and dorsal extrastriate visual cortex (median 0.25–0.31 per neighbor), high in the intraparietal sulcus, the precuneus and posterior cingulate (0.20–0.23), intermediate over postcentral, lateral prefrontal and motor cortex (0.10– 0.18), and lowest, below 0.02, over the temporal pole, the cingulate motor areas, orbitofrontal cortex and the insula (Figure 1E). Counting edges without their weights gives the same map (Spearman ρ = 0.98 in both hemispheres).

### A vertex-level partition of strong, short-range cortical functional connectivity identifies two blocks

The observed variations in the density of strong, short-range connectivity across the cortical surface raise the question of whether the cortical mantle is organized into distinct territories. If it is, the graph should contain a fault line crossed by fewer strong connections relative to those on either side. To find out, we applied spectral decomposition to the 3 ≤ d < 20 mm graph using its symmetric normalized Laplacian, L = I − D^(−1/2) W D^(−1/2), where W contains the edge weights (each weighted by the cortical area of its two vertices; Methods) and D the per-vertex totals. Decomposition of L yields a series of eigenmodes. The first non-trivial one, the Fiedler eigenmode (v_2_, λ_2_), changes sign across the weakest cut in the graph. An abrupt increase in eigenvalues, quantified by the ratio of successive eigenvalues λ_k+1_/λ_k_, indicates that subsequent modes no longer capture well-defined boundaries.

The eigenspectrum thus provides a data-driven estimate of the number of cortical blocks, without imposing predefined partitions (Methods). The normalized cut places boundaries where connectivity between blocks is weakest relative to connectivity within them, effectively assigning each vertex to the block with which it is most densely interconnected^24,25^. The normalized cut between two blocks A and B is defined as cut(A,B)/vol(A) + cut(A,B)/vol(B), where cut(A,B) is the summed weight of the edges between the blocks and vol is the summed weight of all edges of a block’s vertices. In practice, the vertices are ordered along the Fiedler eigenvector, so that vertices strongly connected to each other tend to lie close together, the graph is divided at the position along this order with the lowest normalized cut, and the border is then refined by moving single vertices between the blocks wherever this lowers the normalized cut (Methods).

Partitions are compared with the adjusted Rand index (ARI), the fraction of vertex pairs placed together or apart by both partitions, corrected for chance (1 for identical partitions, 0 for chance agreement); we use it throughout to compare the partitions obtained in the two hemispheres, with different parameters, and in other datasets.

#### Spectral decomposition of cortical connectivity at 3–20 mm yields two blocks

Spectral decomposition of the cohort 3 ≤ d < 20 mm cortical graph gave a single large eigenvalue ratio in both hemispheres, λ_3_/λ_2_ (5.06 L and 4.54 R), after which the ratios fall toward one (Figure 2A). The single gap identifies K = 2 as the data-driven number of blocks. Dividing the graph along the Fiedler ordering at the lowest normalized cut and refining the cut separates each hemisphere into a posterior-central block and an anterior block (Supplementary Results S2). The results agree between the two hemispheres: 99.1% of vertices at homotopic locations receive the same assignment (ARI 0.97). Figure 2B maps the two blocks onto the inflated cortical surface. The posterior-central block (orange; 18,952 vertices, 63.8% L; 18,966, 63.8% R) spans the posterior-central cortical mantle. It encompasses all primary sensory cortex (V1 and V2, 100%; primary somatosensory 3b, 100%; primary auditory cortex, the belt and the parabelt, 100% each), M1 (96% L and 91% R), the posterior parietal lobule, much of lateral temporal cortex, the fusiform complex (100%), the parietal operculum and the posterior insula (OP1, OP2-3, OP4, Ig, 98–100%), and retrosplenial and posterior cingulate cortex (100%). The anterior block (teal; 9,003, 30.3% L; 8,955, 30.1% R) encompasses prefrontal cortex (areas 46, 9a and 10d, 100%), anterior cingulate, premotor (except caudal dorsal premotor area 6d) and supplementary motor cortex and the anterior insula. The peri-allocortex (gray; 1,741 L and 1,795 R vertices, 5.9% L and 6.0% R) is not assigned. A complete parcel-level breakdown is given in Table S2.

**Figure 2.**
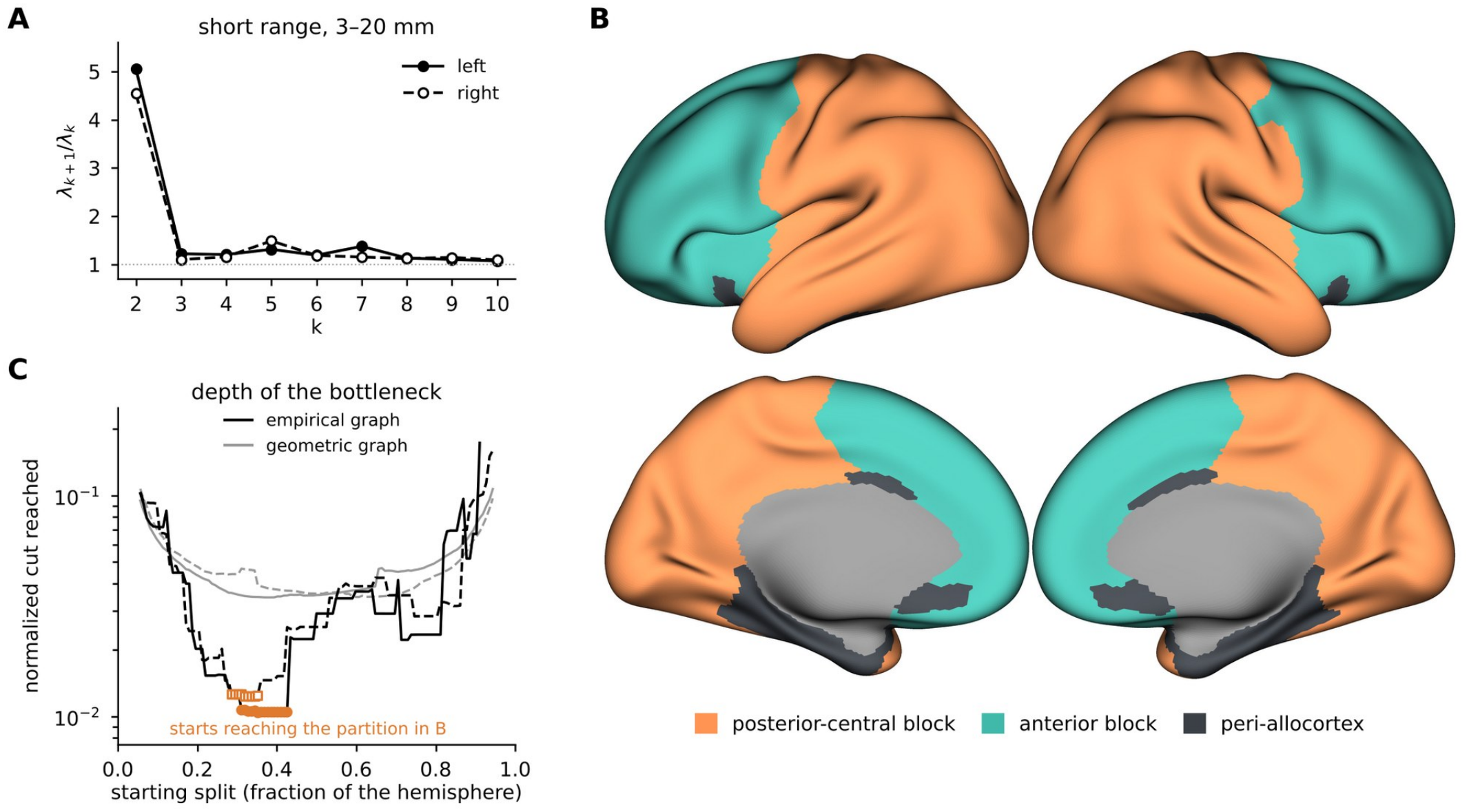
Spectral decomposition of the short-range graph and the resulting two-block partition (HCP 7T, N = 151). **(A)** Eigengap ratios λ_k+1_/λ_k_ of the symmetric normalized Laplacian of the 3–20 mm graph, k = 2–10, L (filled) and R (open) hemisphere. A single gap follows the second eigenvalue (λ_3_/λ_2_ = 5.06 L, 4.54 R); the subsequent ratios approach unity. **(B)** The two blocks on the inflated cortical surface, after refinement of the border by single-vertex descent: posterior-central block (orange; 18,952 vertices L, 18,966 R) and anterior block (teal; 9,003 and 8,955). The peri-allocortex, set aside before the analysis, is dark gray; the medial wall is light gray. Lateral views, top; medial views, bottom. **(C)** The bottleneck between the two blocks is deep. The partition is the two-way split of the Fiedler ordering with the lowest normalized cut, refined by single-vertex descent (Methods). Black: the descent is started from 121 splits evenly spaced along the Fiedler ordering (abscissa, as a fraction of the hemisphere). Only starts near the border reach the lowest cut (orange, within 5% of it). Gray: the same procedure on the geometric graph, in which pairs are weighted by distance alone (Methods). Solid lines and filled markers, left hemisphere; dashed and open, right.

#### The partition identifies a connectivity bottleneck between a cortical expanse with higher density of strong connections and one with lower density

The partition identifies a connectivity bottleneck between the two blocks: a border across which strong short-range connections are far sparser than within either block. The border lies close to a shallow fault line of the geometric graph, which reflects the shape of the cortical sheet^26,27^, but it is about three times deeper and distinct from other candidate partitions (Figure 2C; Supplementary Results S3).

The two blocks differ in density (Figure 1E): the median posterior-central vertex is connected to about three quarters of its neighbors within the 3–20 mm window, whereas the median anterior vertex is connected to about half. The difference increases at the outer bound of the window (at 19–20 mm, connection probability is 0.44 vs. 0.21 L and 0.45 vs. 0.21 R). The difference in density is sufficient to determine the approximate location of the border regardless of the local connectivity pattern within each block. However, at the border the local connectivity deepens the bottleneck. Within the window, pairs of vertices that straddle the border are connected four to five times less often than pairs at the same distance within a block (Supplementary Results S3). For example, the average correlation for pairs of vertices that cross the border is about half of what would be expected based on their geodesic distance, even without imposing any threshold (Supplementary Results S3).

#### The partition is tied to short-range, strong connections

The partition depends on selecting short-range connections. The partition disappears when considering the cohort graph over the full distance range (ARI 0.03 L and R) or over edges of 20 mm or longer alone (ARI −0.06 L and −0.02 R; Supplementary Results S3). Long-range connectivity, by contrast, recovers the established unimodal–transmodal gradient and canonical functional networks (Supplementary Results S3). It also disappears if the same number of strongest connections is chosen regardless of distance (ARI −0.06 L and −0.04 R).

The depth of the partition further depends on the strength of connections because strength determines differences in connection density across vertices. If the same number of edges is drawn at random from all pairs in the 3–20 mm window, so that each vertex keeps on average about 56% of its neighbors, the graph still divides near the same border (ARI 0.83–0.86 L, 0.88–0.89 R), as the geometric graph does, but the cut is four to five times shallower (normalized cut 0.051 L and R; Supplementary Results S3).

Finally, the partition is robust with respect to several parameters: it does not depend on the precise choice of short-range window, on the number of subjects required to support a connection, the way connection weight is calculated, or particular preprocessing steps (Supplementary Results S2).

The two-block organization shown in Figure 2, hereafter termed the main partition, is therefore a specific property of short-range functional cortical connectivity restricted to strong connections, whereas longer-range connectivity recovers the established large-scale organization (Supplementary Results S3).

### The posterior-central and anterior blocks show distinct internal connectivity: lattice-like vs. discontinuous

The partition at 3 ≤ d < 20 mm assigns every cortical vertex outside the peri-allocortex to one of two blocks. We next asked whether and how these blocks differ in their internal organization. Because the posterior-central and anterior blocks differ in connection density, comparisons use the 450 strongest partners of each vertex, about the number of vertices in its short-range neighborhood (the results below are stable from 100 to 450 partners). The partners are taken from the full-range cohort graph, built by the same rule as the short-range graph but over all distances (Methods).

#### The blocks differ in the organization of their connectivity

Figure 3A shows, for one representative subject (550439), the 450 strongest partners of six example vertices (seeds), three from each block, on the left hemisphere. Posterior-central block seeds are V1 (primary visual), V2 (secondary visual) and area 3b (primary somatosensory). Each of the three seeds resolves into a single compact territory containing 99.8–100% of its 450 partners, with almost none beyond 100 mm. Anterior block seeds are three pre-frontal regions: area 46 (dorsolateral), area 9a (dorsomedial) and area 10d (frontopolar). Here, partners are sparse around the seed and form many small territories at long distances (16–17 patches, defined as contiguous clusters of partners within 10 mm of one another).

**Figure 3.**
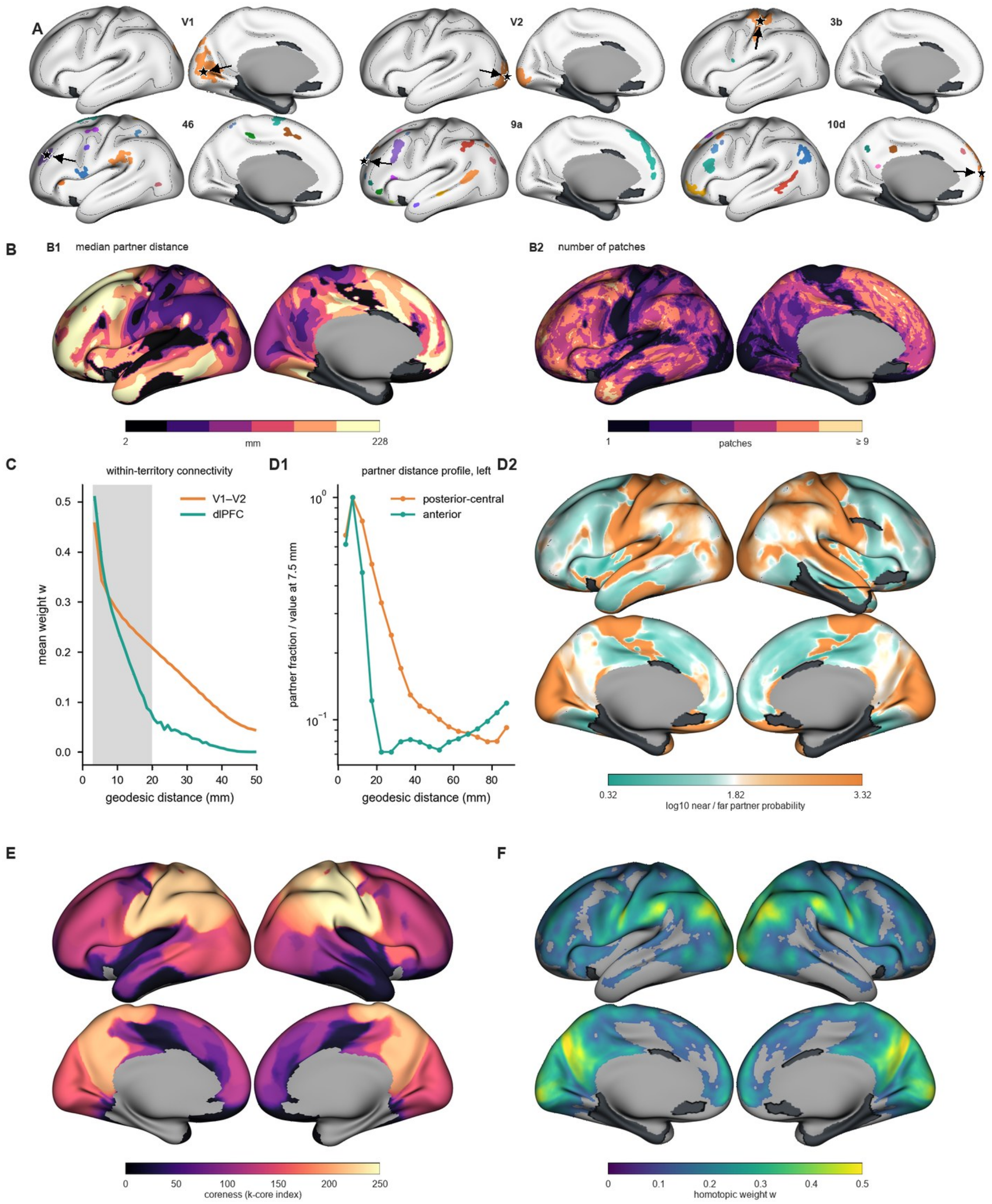
The posterior-central and anterior blocks differ in the organization of their connectivity (HCP 7T, N = 151). Partners are each vertex’s 450 strongest same-hemisphere partners; patches group partners lying within 10 mm of one another. Unless stated otherwise, panels are computed on the full-range cohort graph, using the cohort-median geodesic distance, each hemisphere alone. **(A)** Representative seed vertices on the left hemisphere of one subject (550439), lateral and medial views; seed in black and marked by an arrow, that subject’s 450 strongest partners (pairs with r > τ_s_, ranked on r) colored by patch. Top, posterior-central block seeds (V1, V2, area 3b), whose partners form a single compact territory. Bottom, anterior block seeds (areas 46, 9a, 10d), whose partners form many small territories at long distances. **(B)** Left-hemisphere maps of two measures of partner organization: median partner distance **(B1)** and number of patches **(B2)**. B1 is colored by percentile rank, with the ends of the ramp labeled by the actual minimum and maximum (distances along the folded mid-thickness surface; the largest, up to 228 mm, are those of anterior prefrontal vertices whose strongest partners lie far back in the hemisphere); B2 is on an absolute six-step scale (1, 2, 3, 4–5, 6–8 and ≥ 9 patches). Distance and patch count are highest over the anterior block. Values in the text are medians of the block-wise distributions; Cohen’s d is computed on log_10_ values for B1 and on raw values for B2. **(C)** Mean weight w against within-territory geodesic distance for early visual cortex (V1–V2, the posterior-central block) and dorsolateral prefrontal cortex (areas 46, 9a and 10d, the anterior block), in 1-mm bins from 3 to 50 mm, left hemisphere; every within-territory pair contributes, pairs that are not edges of the cohort graph are set to zero. Shading marks the 3–20 mm short-range window. Posterior-central block connectivity declines gradually and remains graded; anterior block connectivity falls steeply at short distances and then flattens. **(D1)** Fraction of each vertex’s strong partners at geodesic distance d, for the posterior-central block (orange) and the anterior block (teal), in a 3–5 mm band and 5-mm bands beyond it, normalized to the value of the 5–10 mm band, logarithmic scale, left hemisphere. The posterior-central block profile declines steadily with distance; the anterior block profile reaches a minimum at 20–30 mm and rises to the 90 mm limit of the profile. **(D2)** Per-vertex ratio of connection probability near to far, p(3–20 mm) / p(≥ 60 mm), on lateral and medial surfaces of both hemispheres, on a logarithmic scale centered at the Youden cut between the blocks (the threshold on the log ratio that maximizes the sum of the fraction of posterior-central block vertices above it and anterior block vertices below it). Orange, connectivity concentrated near; teal, connectivity comparable or greater far away; white, intermediate. **(E)** Per-vertex coreness from k-core decomposition of the 3–20 mm graph of each hemisphere (0–250). Coreness is highest over the posterior-central block (early visual, sensorimotor, posterior medial-wall, lateral parietal and temporal cortex) and lowest over prefrontal, orbitofrontal and anterior cingulate cortex. **(F)** Per-vertex weight of the edge to the homotopic vertex for both hemispheres. Homotopic weights are strongest in primary sensorimotor, auditory and medial parieto-occipital cortex and weaker in lateral prefrontal and lateral parietal association cortex; gray marks vertices whose homotopic pair is supported in fewer than 76 subjects, which includes most of the peri-allocortex.

Figure 3B presents two measures characterizing the functional connectivity of vertices in the two blocks, computed over the full-range cohort graph.

The first measure (Figure 3B1) is the median distance between each seed and its partners. It is short over occipital, central and superior temporal cortex and long over prefrontal cortex. The block medians are 18.1 mm in the posterior-central block and 80.9 mm in the anterior block L, 18.3 and 91.1 mm R (Cohen’s d 0.99 L and 1.08 R; the test is taken on log distances since they are right-skewed; spin test p = 0.03 L and 0.04 R, 1,000 rotations^28^; the spin test rotates the block map on the sphere, which keeps its size and spatial smoothness, and asks how often a contrast this large arises from a division of the cortex with those properties; Methods).

The second measure (Figure 3B2) asks whether a vertex’s partners form a single territory or several spatially separated patches, where a patch is a contiguous cluster of partners within 10 mm of one another. The median number of patches is 2 in the posterior-central block vs. 5 in the anterior block in both hemispheres (means 2.9 vs. 4.9 L and 2.8 vs. 4.7 R; d = 1.04 L and 1.02 R, spin test p = 0.003 and 0.001; see Supplementary Results S4 for additional details).

#### The blocks differ in their within-area connectivity-by-distance profiles

Figure 3C shows that the distinction between the posterior-central and anterior blocks is also evident in the distance dependence of local connectivity within individual cortical territories. We compared two large, contiguous territories whose intrinsic organization is well characterized in cats and primates: early visual cortex in the posterior-central block (V1–V2), where horizontal connections are dense and local^29–31^, and dorsolateral prefrontal cortex in the anterior block (areas 46–9a–10d), where intrinsic connections are less locally organized^32^. For each territory, we plotted the weight w against the within-territory geodesic distance d between vertices, using the full-range cohort graph, excluding pairs <3 mm apart.

To characterize the decline of w with distance, we used a curvature index κ, the second derivative of a parabola fitted to w against d over 3–50 mm, with both scaled to the unit interval (κ = 0 for a straight line, κ > 0 for a convex curve; Methods). κ was low in the posterior-central block territories (V1–V2, 1.05 L and 1.05 R) and about 2.8 to 2.9 times higher in the anterior block territories (areas 46–9a–10d, 3.03 L and 2.93 R). Thus, posterior-central block connectivity declines gradually and remains graded across the neighborhood, whereas anterior block connectivity falls steeply at short distances before flattening.

#### The blocks differ in overall connectivity-by-distance profiles

We next examined how each vertex’s connections are distributed with distance across its hemisphere (Figure 3D). For each vertex, we again took its 450 strongest partners, binned them by geodesic distance (again excluding pairs closer than 3 mm), and calculated the fraction of its partners in each band. As shown in Figure 3D1, in the posterior-central block (orange) the fraction of strong partners falls steeply with distance and then stays low. In the anterior block (teal) it falls to a minimum at 20–30 mm and then rises again: an anterior block vertex has on average 1.6 times as many of its strong partners in the 85–90 mm band as in the 25–30 mm band (1.7 R). Figure 3D2 maps, for each vertex, how much of its connectivity is near rather than far. It plots the ratio between the fraction of available vertices at 3–20 mm that are strong partners and the same fraction at 60 mm and beyond (60 mm is about where the distance-independent component of the edge probability overtakes the short-range one; Figure 1C). The posterior-central block is rich in vertices whose connectivity is concentrated nearby (high ratios), whereas the anterior block is rich in vertices that connect as readily far as near (low ratios); the block contrast is d = 1.16 L and 1.20 R on log values, p = 0.002 in both hemispheres by the spin test.

#### The blocks differ in internal interconnectedness at short range

Another aspect of internal organization is the degree of interconnectedness, or coreness, obtained by iteratively removing the least-connected vertices through k-core decomposition^33^. A k-core is the maximal subgraph in which every vertex has at least k partners within that subgraph. High coreness therefore identifies vertices embedded within a large, mutually interconnected core, rather than vertices that simply have many connections; maximum coreness defines the innermost shell whose vertices retain at least that many mutual connections. Figure 3E maps coreness onto the cortex for connections within the 3–20 mm window. The innermost cores lie within the posterior-central block territory, with coreness falling to its lowest values across prefrontal, orbitofrontal and anterior cingulate cortex and over the temporal pole (Supplementary Results S4).

#### Interhemispheric coupling is strongest where short-range connectivity is densest

It is known from both neuroanatomical and neurophysiological studies of early sensory areas that the dense horizontal circuitry of posterior-central cortex continues through the corpus callosum to homologous contralateral territory^34,35^. For each vertex, we took the weight of the edge to its homotopic vertex (the same fs_LR vertex in the other hemisphere). As shown in Figure 3F, homotopic weight is greatest in primary sensorimotor, auditory and medial parieto-occipital cortex. The distribution of strong cross-hemispheric homotopic weights resembles that of intra-hemispheric short-range weight (Figure 1E), as confirmed by their vertex by vertex correlation (Spearman ρ = 0.85 and 0.87, L and R). The median homotopic weight is about 1.3 times higher in the posterior-central block than in the anterior one (0.209 vs. 0.163 L, 0.210 vs. 0.164 R). Because homotopic pairs lie in different hemispheres, these correlations are free of the hemodynamic point-spread function and of the projection from volume to surface.

In summary, posterior-central block vertices connect primarily locally, cover their neighborhoods densely, concentrate their connection weight within one or two contiguous territories, and more than a third (35% L and 35% R) have no strong partner beyond 60 mm. Anterior block vertices connect farther, distribute their weight across several smaller territories, and have more strong partners at longer distances. The innermost cores of the cortical graph lie entirely within the posterior-central block, where homotopic interhemispheric connectivity is also strongest.

To capture these differences in internal organization, we term the posterior-central block lattice cortex and the anterior block paralattice cortex. The lattice–paralattice partition recovered from strong, local connectivity at 3 ≤ d < 20 mm thus reflects distinct geometries of cortical connectivity at the near-mesoscale.

### Lattice cortex is organized internally as an integrated multi-stream hierarchy and externally by connectors to paralattice and peri-allocortex

We next asked whether, at the near-mesoscale, additional structure can be identified within lattice cortex. Several features emerge.

#### Lattice cortex can be subdivided into four sectors that are not sharply demarcated

We first applied the same spectral decomposition procedure used to find the main partition to the lattice cortex of each hemisphere (18,952 L and 18,966 R vertices), using the same 3 ≤ d < 20 mm window (2.66 L and 2.68 R million within-lattice edges). Unlike the cortex-wide spectrum, whose single large eigenvalue ratio fixes the number of blocks at two (Figure 2A), the within-lattice spectrum has no dominant ratio (largest 1.88 L and 1.92 R, compared with 5.06 and 4.54 for the whole hemisphere). Thus, lattice cortex is not separable into sharply demarcated blocks. Nevertheless, a natural subdivision into four sectors can be identified by asking at which K a partition reproduces independently in the two hemispheres and across two independent halves of the cohort (Supplementary Results S4).

The resulting four sectors are shown in Figure 4. The first sector (Figure 4A; 4,086 L and 4,265 R vertices) covers V1 to V4, V3A and V6; the second sector (Figure 4B; 6,466 L and 6,404 R vertices) covers the lateral pre- and postcentral strip together with the inferior parietal and intraparietal areas (PFm, PF, PGi, AIP, PGs); the third sector (Figure 4C; 4,143 L and 3,824 R vertices) covers A1, the belt, the superior temporal sulcus and lateral temporal cortex; and the fourth sector (Figure 4D; 4,257 L and 4,473 R vertices) covers medial parietal cortex (POS1, POS2, RSC, 23c, PCV, 7m) together with the medial and caudal motor strip. Thus, the first three sectors include primary areas in the visual, somatomotor, and auditory modalities, respectively, while the fourth sector does not.

**Figure 4.**
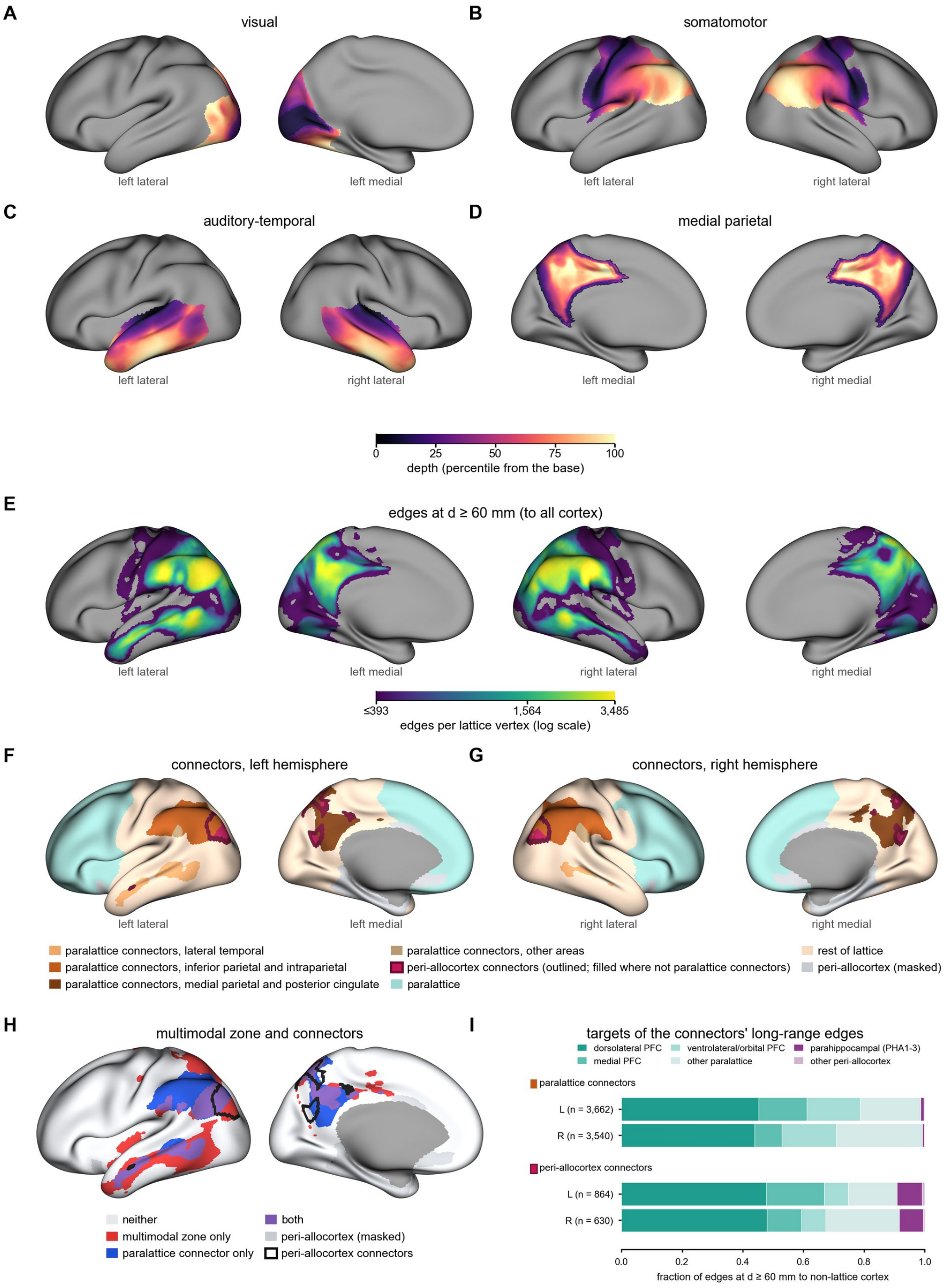
Internal organization of lattice cortex, and its connections with paralattice cortex and the peri-allocortex. Each hemisphere was analyzed separately. **(A–D)** Within-sector depth for the four sectors obtained by spectral decomposition of the lattice of each hemisphere at 3 ≤ d < 20 mm (K = 4, refined by single-vertex descent; see text). Depth is the length of the shortest path from the base through the sector’s full-range graph (all pairs at least 3 mm apart), each edge counting as the inverse of its weight, expressed as a percentile within the sector (0, dark, at the base; 100, light, farthest from it). The base is V1 for the visual sector (**A**; left lateral and medial views), area 3b for the somatomotor sector (**B**; left and right lateral views), A1 for the auditory-temporal sector (**C**; left and right lateral views) and, for the medial parietal sector, which has no primary area, its border with other cortex and the medial wall (**D**; left and right medial views; depth computed within medial parietal cortex proper). In C, the anterior branch reaches the temporal pole, and the posterior branch, toward STSvp, ends near the middle of the depth range. **(E)** Number of edges at d ≥ 60 mm (to all cortex) per lattice vertex, both hemispheres, logarithmic scale. **(F, G)** Paralattice and peri-allocortex connectors, left **(F)** and right **(G)** hemisphere, lateral and medial views. For each target (paralattice cortex or the peri-allocortex), the connectors are the lattice vertices with the most edges at d ≥ 60 mm into it that together send three quarters of all such edges from lattice cortex. Paralattice connectors are shown by region: lateral temporal (light orange), inferior parietal and intraparietal (dark orange), medial parietal and posterior cingulate (brown) and other areas (tan). Peri-allocortex connectors are outlined in dark crimson and filled in crimson where they are not also paralattice connectors. The rest of the lattice is shown in pale orange, paralattice cortex in pale teal and the peri-allocortex in gray. **(H)** Overlap of the paralattice connectors (blue) with the multimodal zone (red; purple where they overlap); peri-allocortex connectors outlined in black. **(I)** Targets of the edges at d ≥ 60 mm into non-lattice cortex of the paralattice and the peri-allocortex connectors, by hemisphere (color keys as in F and G; the two populations are not exclusive: 506 L and 346 R vertices belong to both); n, number of vertices.

#### Each sector has its own within-stream hierarchy

Within each sensory sector, depth from the primary area (V1, area 3b or A1) along strong connections of any length within the sector orders the cortical areas much as the hierarchies established by anatomy and physiology in primates: in the visual sector from V1 through V2–V4 then branching to dorsal and ventral extrastriate areas, in the somatomotor sector from 3b through areas 1 and 2 and the anterior parietal areas to PFm and the angular gyrus, and in the auditory-temporal sector from A1 through the belt and the parabelt into anterior and posterior temporal branches. Distance along the surface from the same base already gives much of this order; the strong connections add to it in the visual, somatomotor and anterior auditory streams. The medial parietal sector, which has no primary area, deepens from its border, the only one of five bases tried from which depth behaves as in the sensory sectors, toward the posterior cingulate cortex and the central precuneus (Supplementary Results S4).

#### The hierarchies converge on shared posterior zones

The apices of the three sensory hierarchies and the base of the medial parietal hierarchy adjoin in posterior association cortex: the visual stream ends at the temporo-parieto-occipital junction, the somatomotor stream in PFm and the angular gyrus, and the posterior auditory branch in STSvp, near MIP and 7PC at the base of the medial parietal sector. Many vertices there connect across sectors. For each lattice vertex, we calculated the fraction of its lattice partners in the full-range graph that lie in sectors other than its own. The top quintile (3,791 L and 3,794 R vertices), which we call the “multimodal zone”, lies in the inferior parietal lobule and intraparietal sulcus (PGi, PGs, MIP, LIPv, VIP), in lateral temporal cortex and the temporo-parieto-occipital junction (TE1p, STSvp, PHT, PH, PGp), less extensively in the precuneus and posterior cingulate cortex (7Pm, 7Am and, on the left, 7m and 31pd) and, on the right, along the border between the somatomotor and medial parietal sectors (7PC, 2, 23c). This shared zone helps explain why the streams do not separate as blocks. The auditory-temporal sector also connects with the top of the medial parietal hierarchy (Supplementary Results S4).

#### Lattice “connectors” to paralattice cortex and the periallocortex lie in parietal, medial parietal and lateral temporal cortex

Some clusters of vertices of posterior-central cortex, which we call “lattice connectors”, combine the dense short-range connectivity of lattice cortex with many long-range edges (d ≥ 60 mm; Figure 4E). Lattice connectors can be further distinguished by their targets outside lattice cortex, which gives two populations that are not mutually exclusive. “Paralattice connectors” occupy the inferior parietal lobule and intraparietal sulcus, the precuneus and smaller parts of lateral temporal cortex and, on the left, anterior temporal cortex (Figure 4F,G). Their long-range edges go mainly to prefrontal cortex, most densely to its caudal and mid-lateral part (Figure 4I), and can be subdivided into separate channels (Supplementary Results S4). A second, smaller population sends edges to the peri-allocortex, nearly all of them to parahippocampal cortex (PHA1–3). These “peri-allocortex connectors” lie in the angular gyrus, the precuneus and the parieto-occipital sulcus, and more than half of them are also paralattice connectors (506 of 864 L and 346 of 630 R vertices belong to both; Figure 4F,G,I). About two fifths of the paralattice connectors on the left and a third on the right, and more than half of the peri-allocortex connectors, lie in the multimodal zone (Figure 4H). More details are provided in Supplementary Results S4.

In summary, lattice cortex comprises four reproducible but weakly separated sectors, each with its own hierarchy, which converge in posterior association cortex. There, connectors combine dense local connectivity with long-range connections to prefrontal paralattice cortex, and a subset of them, in the angular gyrus and the precuneus, also connects with parahippocampal cortex.

### The main cortical partition largely corresponds to thalamic basal ganglia / cerebellar organization

Next, we asked whether the main cortical partition defined by short-range functional connectivity corresponds to a basic anatomical subdivision of the underlying thalamus and associated basal ganglia and cerebellar circuits.

We focus on “specific” thalamic nuclei that provide point-to-point, driving inputs to their cortical targets, terminating primarily in granular layers^36,37^. As shown in Table S3, “specific” nuclei can be divided according to whether they receive basal ganglia and/or cerebellar inputs. First-order sensory relay nuclei (lateral geniculate, medial geniculate, ventral posterior), as well as higher-order specific nuclei (pulvinar, lateral posterior), receive little or no such input^36–40^. These specific nuclei connect primarily to posterior-central cortex. Conversely, specific thalamic nuclei receiving basal ganglia and/or cerebellar inputs include the ventral anterior, ventral lateral, and mediodorsal nuclei and connect primarily to anterior cortex and peri-allocortex. In primates, cortical territories in anterior regions (motor, premotor, prefrontal) participate in numerous partially segregated, primarily unidirectional cortico-basal-ganglia-thalamo-cortical loops^41–43^. They also participate in cortico-ponto-cerebellum-thalamo-cortical loops^42^. The cortical somatomotor strip’s motor side is a major recipient of thalamic inputs participating in basal ganglia and cerebellar loops, whereas the sensory side is not^44^.

#### Thalamic nuclei receiving anatomical basal ganglia / cerebellar input show preferential functional connectivity to paralattice cortex, those that do not to lattice cortex

We asked whether the main partition obtained from the spectral decomposition of short-range functional connectivity separates specific nuclei that are involved in basal ganglia and/or cerebellar loops from those that are not. The mean time course of each nucleus was partially correlated with every cortical vertex of the same hemisphere, controlling for the mean signal of the rest of the thalamus (Methods).

The results for the 55 nucleus estimates (Figure 5A–C, Supplementary Results S5 and Table S4) show that the basal ganglia-recipient motor nuclei (ventral anterior, parvocellular and magnocellular, and ventral lateral anterior) couple more strongly with paralattice than with lattice cortex in every estimate and so does the cerebellar-recipient ventral lateral posterior nucleus. This holds even though the ventral lateral nucleus provides the main thalamic input to primary motor cortex, which lies in lattice cortex: in the present analysis, the ventral lateral anterior nucleus couples less with primary motor cortex than with the rest of lattice cortex, and less still than with paralattice cortex (Supplementary Results S5).

**Figure 5.**
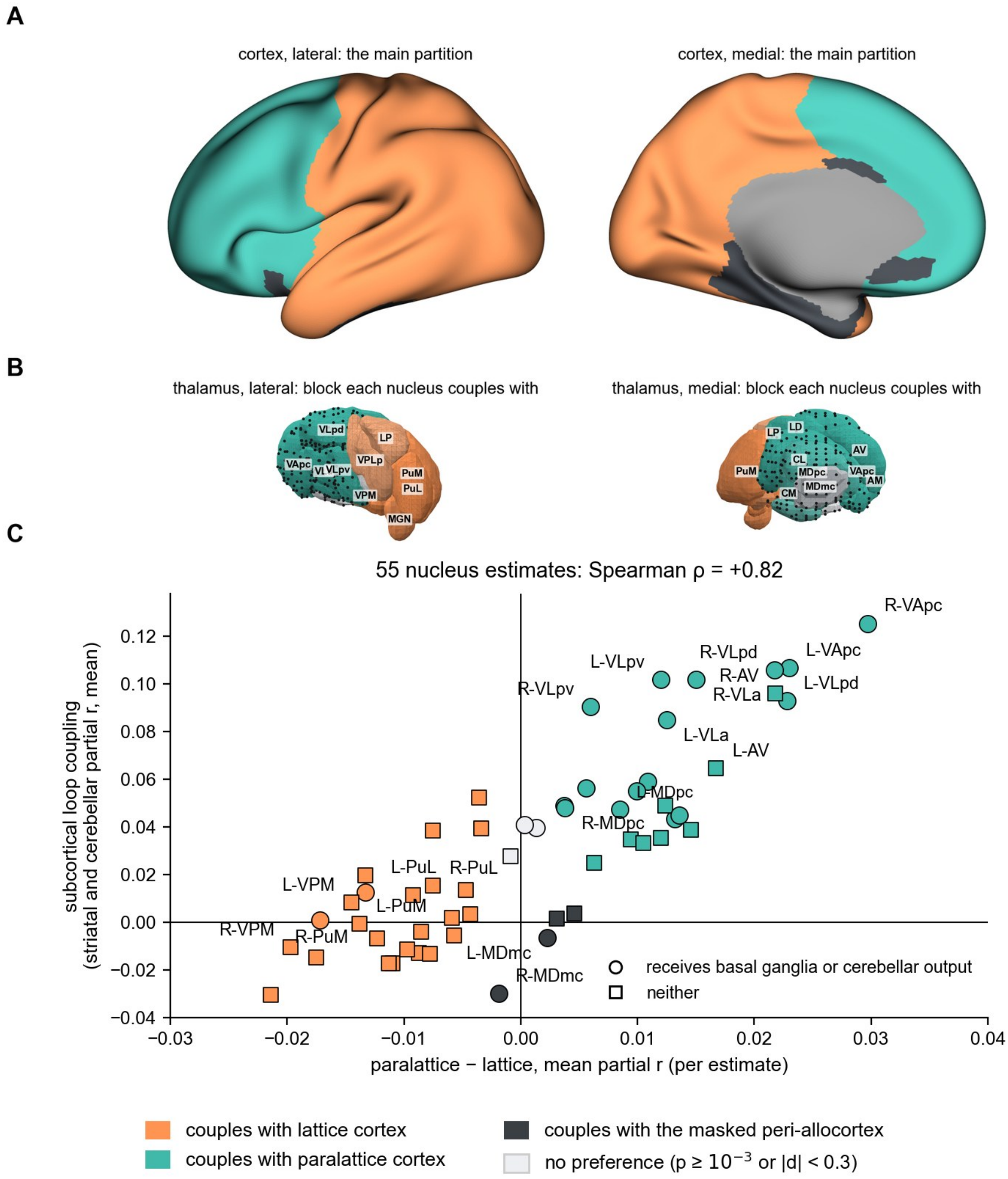
Thalamic nuclei that receive basal ganglia or cerebellar output couple with paralattice cortex and with the subcortical loops; relays and pulvinar couple with lattice cortex. **(A)** The main partition on the left hemisphere (lateral and medial views; lattice orange, paralattice teal; gray, the peri-allocortex, as in Figure 2B). **(B)** The left thalamus (Morel atlas, lateral and medial views), each nucleus colored by the cortical territory it couples with, in the colors of A (orange lattice, teal paralattice, gray the peri-allocortex), lighter where |d| is smaller, near-white where no territory is preferred (p ≥ 10^−3^ or |d| < 0.3); dots mark nuclei that receive basal ganglia or cerebellar output. In the medial view the thin midline nuclei are omitted and the magnocellular mediodorsal division is brought forward so that it is visible. **(C)** The 55 nucleus estimates: preference for paralattice over lattice cortex (mean difference of partial correlation across 151 subjects) against subcortical loop coupling (mean of the striatal and cerebellar partial correlations); circles, nuclei that receive basal ganglia or cerebellar output; squares, neither; colors represent the result of the paired test (Table S4).

The specific nuclei that do not receive basal ganglia and/or cerebellar inputs are instead coupled with lattice cortex. The first-order relays (ventral posterior medial, posterior part of the ventral posterior lateral, ventral posterior inferior, medial geniculate), and the higher-order nuclei (pulvinar, lateral posterior, posterior, limitans and suprageniculate nuclei) all couple more strongly with lattice cortex.

The anterior nuclei (anteroventral, anteromedial, lateral dorsal), which receive neither basal ganglia nor cerebellar input and project to cingulate and retrosplenial cortex, couple more with paralattice cortex and with the peri-allocortex than with lattice cortex. Among the intralaminar nuclei, the central lateral and central medial nuclei prefer paralattice cortex while the centromedian nucleus prefers lattice cortex.

Finally, across nuclei (the two hemisphere estimates of each nucleus averaged), the coupling of a nucleus with subcortical loops predicts its cortical preference: the more strongly a nucleus couples with paralattice relative to lattice cortex, the more strongly it couples with the striatum and the cerebellum (Supplementary Results S5).

In summary, specific thalamic nuclei that relay basal ganglia or cerebellar output couple with paralattice cortex, and first-order sensory relays and higher-order nuclei such as the pulvinar couple with lattice cortex.

### The main partition generalizes across stimulus-driven activity and other datasets

The next section aims at establishing the generality of the main lattice–paralattice partition. We asked whether the partition persists during movie watching, and whether it replicates in a larger 3T cohort and, subject by subject, in a precision-fMRI cohort. In every case the partition is re-derived at the 3–20 mm range with the same criteria as for the 7T resting-state cohort: the same peri-allocortex, a null floor τ_s_ solved for each subject on its own 3–20 mm population, an edge present in a majority of the cohort’s own subjects with weight the cohort mean of r, and the same spectral decomposition and descent, separately for each hemisphere (Methods).

#### The main partition is preserved during stimulus-driven activity

We first asked whether the main partition observed in the awake resting state (Figure 6A) persists during continuous audiovisual stimulation in the HCP 7T movie-watching cohort (N = 160). In the raw movie dataset we identified the same two blocks (Figure 6B). The disagreement with the resting state is small and mostly at the border, in the premotor and cingulate motor belt and the posterior insula and parietal operculum (dashed line in the figure). Removing the stimulus-locked component makes almost no difference (Figure S1), showing that the main partition does not depend on the stimulus being present or absent from the signal. Details are provided in Supplementary Results S6 and a full per-parcel listing in Table S2.

**Figure 6.**
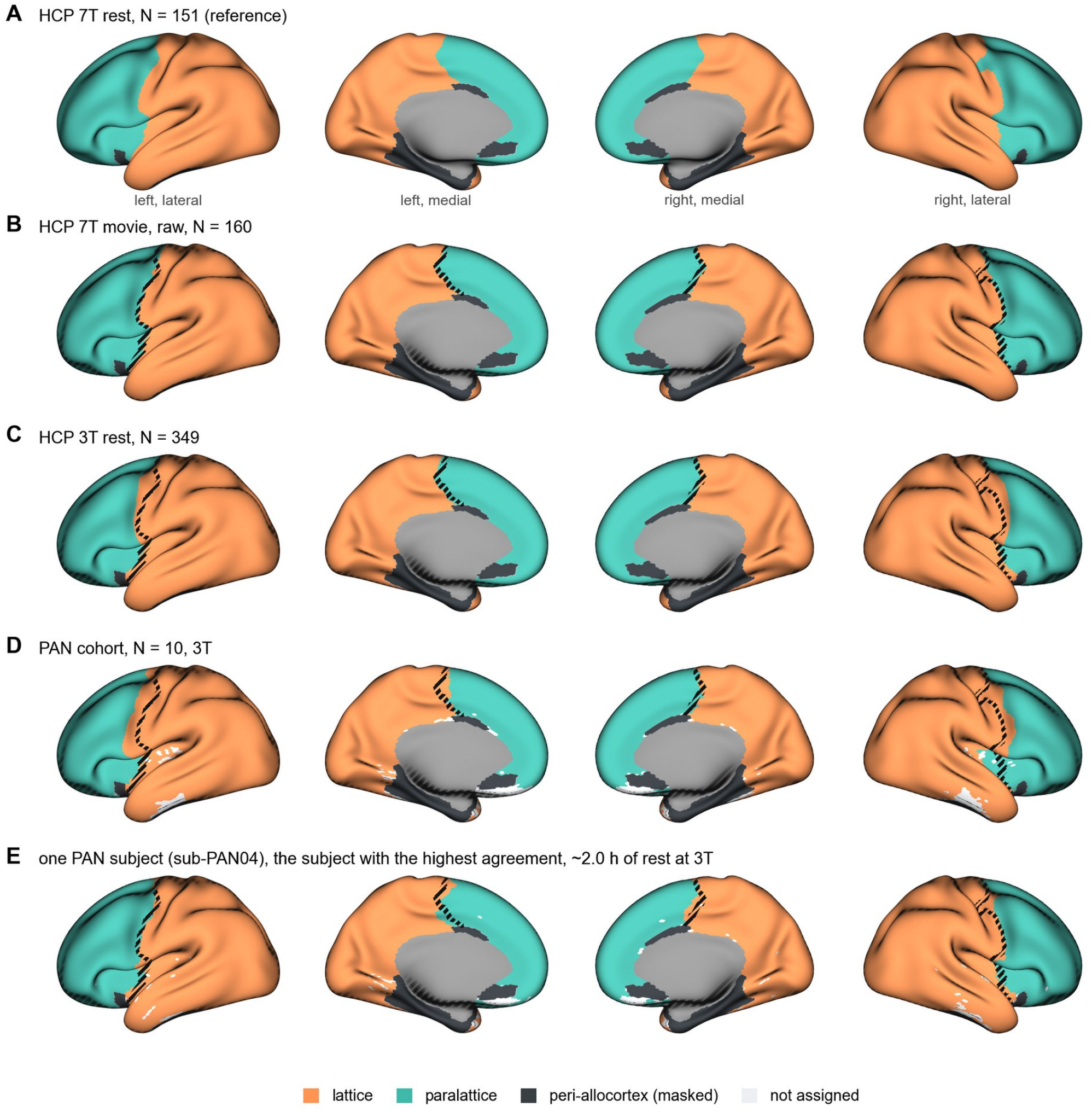
The main partition replicates across conditions, cohorts and field strengths (both hemispheres, lateral and medial views). **(A)** The HCP 7T resting-state reference partition (N = 151; each hemisphere partitioned separately). **(B)** HCP 7T movie watching (N = 160; ARI 0.98 L and 0.97 R against rest); the movie data after removal of the stimulus-locked component are shown in Figure S1. **(C)** HCP 3T (N = 349; ARI 0.91 and 0.89 against the 7T reference). **(D)** The ten-subject PAN cohort (3T; ARI 0.82 and 0.86). **(E)** One PAN subject, from that subject’s data alone (∼2.0 h of rest at 3T): sub-PAN04, the subject with the highest agreement with the reference (ARI 0.91 and 0.90). Every replication was re-derived by the same construction with every parameter re-solved on its own cohort or subject; for PAN the sweep margin is proportional (Methods). Orange, lattice; teal, paralattice; gray, the peri-allocortex; near-white, vertices with no edge in the graph, not assigned. In B–E the dashed line marks the border of the 7T resting-state partition shown in A, so that where a replication departs from it can be read directly.

#### The main partition replicates at 3T in a larger cohort

We next asked whether the main partition identified at 7T generalizes to standard-resolution 3T acquisitions (N = 349), with the same preprocessing chain (Methods).

As shown in Figure 6C, the 3T graph subdivides into two blocks, closely corresponding to 7T (ARI 0.91 L and 0.89 R). In both hemispheres the border lies slightly more anteriorly at 3T. The two 3T hemispheres, each partitioned alone, agree with each other (ARI 0.95), also similar to the 7T hemispheres (ARI 0.97). Details are provided in Supplementary Results S6 and a full per-parcel listing in Table S2.

#### The main partition replicates at the single-subject level with precision fMRI

We further tested the primary HCP 7T findings using the Precision Targeting of Association Networks dataset (PAN^21^; OpenNeuro ds006598, 3T, N = 10). Despite its lower field strength, PAN combines multi-echo acquisition and deep within-subject sampling, making it well suited to single-brain replication and complementary to the population-level power of the primary cohort. The multi-echo acquisition was used to remove non-BOLD signal components from each run. Each PAN subject was treated as an independent dataset (Methods).

Again, the ten-subject cohort graph matched the 7T reference (ARI 0.82 L and 0.86 R) with the two hemispheres agreeing at 97.8% of homotopic positions (ARI 0.91) (Figure 6D). As in the HCP 3T cohort, the border lies more anteriorly than at 7T, more so in the left hemisphere. The partition is also recovered from every single subject, in both hemispheres (Figure 6E). Details are provided in Supplementary Results S6 and a full per-parcel listing in Table S2.

Altogether, the above replications confirm the location of the main partition described in the 7T cohort. Across the replications the partition changes slightly, mostly at the border of the lattice: at the premotor and cingulate motor strip (6mp, 6d, 24dd) and at its Sylvian end, in the posterior insula and operculum (PoI2, 43, FOP2); at 3T, the frontal eye field, 55b and ventral premotor cortex also move to lattice cortex.

#### The main partition aligns with cortical dynamics more strongly than cytoarchitecture

Finally, we asked how the main partition recovered from BOLD coupling at 3 ≤ d < 20 mm relates to other measures of cortical organization. We compared the block-wise distributions of 27 published cortical maps from the neuromaps annotation library^45^ (Methods). For each map, we calculated the block-wise mean and the Spearman correlation ρ(block) between its per-vertex values and block ordinal index (lattice = 1, paralattice = 2, peri-allocortex = 3, Table S5). Positive ρ indicates increasing values from lattice through paralattice to peri-allocortex, whereas negative ρ indicates the reverse.

The strongest associations were with cortical dynamics measured by magnetoencephalography (MEG): delta power (ρ = +0.70: lattice low, paralattice high), alpha power (ρ = −0.71: lattice high, paralattice low), gamma power (ρ = +0.63 at 30–60 Hz and +0.57 at 60–150 Hz: lattice low, paralattice high), and intrinsic timescale, the decay time constant of the temporal autocorrelation of the regional MEG signal (ρ = +0.66: lattice short, paralattice long^46^). None of these associations survives the spin test after correction across the 27 maps (Table S5), so they are reported descriptively. Details are provided in Supplementary Results S6.

On the other hand, the lattice–paralattice subdivision does not map directly onto cytoarchitectonic divisions of the cerebral cortex. Assigning each vertex a laminar-differentiation type from BigBrain histology^47^, graded by granular-layer (layer 4) development, reveals a graded relationship with block membership. Details are provided in Supplementary Results S6.

In summary, the main partition follows cortical dynamics most closely, and laminar differentiation less closely.

## DISCUSSION

We analyzed resting-state fMRI functional connectivity at vertex resolution in a primary 7T cohort of 151 subjects and several replication datasets. To target near-mesoscale interactions, we restricted the correlation matrix to strong, short-range edges (3 ≤ d < 20 mm). Spectral decomposition of this short-range graph partitions the cortical mantle into two blocks, robust across parameter choices, experimental conditions and participant cohorts. The larger is a posterior-central expanse, about twice the size of the anterior block. These two blocks are separated by a border evident in short-range, near-mesoscale connectivity and differ in both their internal and their external connectivity. As discussed below, the internal organization of the posterior-central block supports its designation as lattice cortex, and we refer to the anterior block as paralattice cortex.

### The cortical main partition and strong, short-range functional connectivity

The main partition emerges only when functional connectivity is restricted to near-mesoscale, strong, short-range correlations. The 3–20 mm window was chosen to reflect interactions potentially mediated by intra-areal gray matter fibers and short U-fibers connecting neighboring areas: horizontal axons of layer 2 and 3 pyramidal neurons extend 2–8 mm within gray matter^4,29^, and U-fibers linking adjacent gyri are usually defined as association fibers no longer than 30–40 mm along their course, with a median length of about 20 mm in tractography^48,49^. Such connections are thought to mediate strong, direct interactions among neuronal populations whose neurons share much of their input and output. Longer connections, mediated by anatomically distinct white matter tracts, are organized differently and are mostly excluded by the cutoff. The choice of window size is supported by the observed decay of edge probability with distance, which separates a short-range component from a component that is independent of distance. Human intracranial recordings are consistent with this distinction: after a small focal lesion, sleep-like slow waves prevail in perilesional cortex and spread farther only along specific long-range connections^50^. The inner bound of 3 mm was chosen to exclude distances where substantial spurious correlations are most likely to occur, from hemodynamic and vascular spread^51,52^ or from volume-to-surface mapping^53,54^ (Supplementary Results S1).

A single, reproducible border divides the short-range graph into two contiguous blocks. Spectral decomposition shows that the second (Fiedler) eigenmode is associated with a large eigengap, after which eigenvalues flatten. This sets the number of well-defined blocks to two. The Fiedler eigenmode places posterior-central cortex and anterior cortex at opposite ends, with the border identified through a sweep minimizing the normalized cut followed by single-vertex descent. The posterior-central block comprises all primary sensory cortex, the posterior parietal lobule, much of lateral temporal cortex, the fusiform complex, the parietal operculum and the posterior insula, and retrosplenial and posterior cingulate cortex. The anterior block comprises prefrontal cortex, anterior cingulate, premotor and supplementary motor cortex, and the anterior insula (Figure 2).

The border between the two blocks marks a bottleneck of short-range connectivity. The normalized cut at the border is about two and a half to three times deeper than other divisions of the same graph and about three times as deep as the best cut of the geometric graph, a nearby, shallow fault line which reflects the shape of the cortical sheet^26,27^. Moreover, pairs of vertices that straddle the border are connected four to five times less often than predicted by their distance. The border appears to mark where the intrinsic cortical wiring changes from the locally connected, topographic regime of sensory and motor cortex to the discontinuous, striped regime of prefrontal cortex, with premotor, cingulate motor and opercular-insular cortex as the belt between them (see Supplementary Results S3). The two blocks are, however, heavily interconnected by long-range edges (see lattice connectors below).

The main partition does not depend on the precise bounds of the window, as it remains essentially unchanged when the inner bound is raised or the outer bound lowered by several millimeters. It is also robust to methodological choices and potential confounds. The same two blocks persist across changes in windows, thresholds, number of subjects above threshold, and cohorts (Table S6), with or without global-signal regression, and in two halves of the cohort processed separately. Most importantly, the main partition persists during movie watching, replicates in a larger 3T cohort, and is recovered in individual subjects of a precision-fMRI dataset (Figure 6).

Conversely, the main partition vanishes when considering the full-range cortical functional connectivity, confirming its dependence on short-range edges: without the distance restriction the graph yields a single dominant cluster. Moreover, long-range edges alone show hardly any trace of the main partition (Supplementary Results S3). Instead, when spectral decomposition is restricted to longer-range connectivity, enforcing two clusters yields a partition with mainly unimodal areas on one side and mainly transmodal areas on the other, along a vector similar to the principal gradient^55^; enforcing seven clusters yields six of the seven Yeo networks^56^, save for the limbic network (which lies in the orbitofrontal and temporopolar signal-dropout zone) (Supplementary Results S3). Recovering these known large-scale divisions supports the spectral approach itself; what sets the main partition apart is that its border is a deep, localized bottleneck, and it appears only in the short-range graph. The main partition also depends on the short-range correlations being strong: for the same edge density, choosing the edges at random, regardless of strength, yields a division that is four to five times shallower (Supplementary Results S3).

In the fMRI literature, correlations below roughly 20 mm are often discarded or averaged over, usually without testing whether the cutoff removes mainly artifactual correlations or also valid ones. However, the distance range up to 20 mm is the one in which cortical synaptic connectivity is densest and thus likely to generate strong functional dependencies in neural signals. Anatomically, connection probability falls approximately exponentially with distance, such that short connections outnumber long ones by orders of magnitude^6,19^; intrinsic horizontal collaterals of supragranular pyramidal neurons extend for millimeters and form clustered terminations at that scale^4,29^; and short U-fibers, which run beneath the cortex to an adjacent gyrus, are up to 30–40 mm long, with a median of about 20 mm in tractography^48,49^. As detailed in Supplementary Results S1, concerns regarding hemodynamic spread, vascular blurring and other noise sources^51,52,57^ can be addressed: a spatially uniform non-neural signal component falls off within a few millimeters, well inside the window; correlations between facing banks of a sulcus follow distance along the cortical surface rather than through the volume; and the vascular topography does not follow the partition. Such artifacts should also affect short-range correlations across the whole cortical surface, whereas the partition rests on differences in short-range connectivity between cortical territories.

### The internal organization of lattice cortex

The main partition reflects differences in the internal organization of the two blocks, most clearly revealed by the distinctive functional connectivity of lattice cortex (Figure 3). These differences should not be considered as absolute, since neither block is uniformly organized inside, and there can be substantial local variations. Thus, posterior-central cortex, here the focus of further analysis, shows varying degrees of local connectivity, can be further subdivided into modality streams that are hierarchically organized, and comprises multimodal regions as well as regions with dense local and long-distance connectivity.

Even so, several features justify designating the posterior-central block as “lattice cortex” (without implying a lattice in a strict graph-theoretical sense). Lattice cortex has denser short-range connectivity: the median lattice vertex is connected to about three quarters of its neighbors within 3–20 mm, the median paralattice vertex to about half, and the probability of a connection at the outer bound of the window is more than twice as high. Its connections are local and spatially compact, forming in most cases one or two patches. Within lattice areas, connectivity declines gradually with distance and remains graded across each vertex’s neighborhood. Across the hemisphere, more than a third of lattice vertices do not have any of their strongest partners at long range. This dense, local, compact organization motivates the term lattice cortex. Lattice cortex is also more integrated than paralattice cortex at short range, as indicated by its coreness: the innermost cores of the cortical graph lie entirely within lattice cortex. By contrast, short-range connectivity in paralattice cortex is more distributed. Its vertices connect to much more distant and dispersed partners, distributing them over more patches, about five against two. Within paralattice areas, correlations fall steeply before flattening with distance. Across the hemisphere, the fraction of strongest partners reaches a minimum just beyond the window and then rises again, with more than one and a half times as many strong partners at long range.

Further spectral decomposition of lattice cortex separates it into four sectors (Figure 4): visual, somatomotor, auditory-temporal, and medial parietal. This provides an internal validation of the spectral approach by recovering established anatomical and functional subdivisions^58,59^. Unlike the cortex-wide partition, these subdivisions do not represent genuine bottlenecks (the eigenmode profile shows no large eigengaps) and the number of sectors is chosen based on reproducibility across hemispheres and independent halves of the cohort. Even so, within each sector, the analysis of near-mesoscale connectivity recovers a hierarchical organization from primary toward higher-order areas, consistent with the progressive spread of intra-areal connectivity from early to higher areas^4,60,61^. The medial parietal sector has no primary area of its own. Its hierarchy runs inward from its border, where it meets the visual and somatomotor sectors, retrosplenial cortex and the medial wall, to the dorsal posterior cingulate cortex and the central precuneus.

Vertex-level functional connectivity also shows that the four streams are not isolated but interact, particularly at higher levels (Figure 4), consistent with substantial evidence for multimodal responses in posterior cortex^62^. The lattice vertices with the largest share of their partners in other sectors form a “multimodal zone”, concentrated in the inferior parietal lobule and intraparietal sulcus, in lateral temporal cortex and the temporo-parieto-occipital junction, and extending into the precuneus. Spectral decomposition of short-range connectivity complements gradient-based approaches such as diffusion embedding across all connectivity distances^55^, which identifies two orthogonal gradients: one extending from unimodal sensory and motor cortex to transmodal default-mode cortex, and another placing visual cortex at one pole and somatomotor and auditory cortex at the other.

Another subset of lattice vertices, the “lattice connectors”, combines the dense local connectivity of lattice cortex with the largest number of long-range connections into paralattice cortex and peri-allocortex (Figure 4E–I). “Paralattice connectors” occupy the inferior parietal lobule and intraparietal sulcus, the precuneus (continuous with the parietal territory on the right, separate on the left), and lateral temporal cortex along the posterior superior temporal sulcus. Their edges to non-lattice cortex are long and target mainly prefrontal cortex, nearly half of them dorsolateral prefrontal cortex; the rest reach ventrolateral and medial prefrontal cortex and other paralattice regions, and few reach orbitofrontal cortex, with fewer reaching medial prefrontal cortex on the right, especially from the lateral temporal territory. A small fraction of the edges leaving the lattice reaches the peri-allocortex (“peri-allocortex connectors”), almost entirely parahippocampal cortex, from the angular gyrus and medial parietal cortex, in line with the parieto-medial temporal pathway; these edges are concentrated in few vertices (about one twentieth of lattice cortex sends three quarters of them), and about a third of them come from vertices that are both paralattice connectors and multimodal.

Lattice connectors overlap the multimodal zone in the inferior parietal lobule and lateral temporal cortex, the peri-allocortex connectors more extensively than the paralattice connectors. Whether their short- and long-range connections originate from distinct cortical layers or cell populations remains an open question.

Lattice connectors also partly overlap with the structural “rich club” identified by diffusion tractography, the set of nodes more densely interconnected than expected from their degree. The posterior cortical contingent of the rich club, in precuneus and superior parietal cortex, lies within lattice cortex, whereas its superior frontal contingent lies within paralattice cortex (the rich club also includes subcortical nodes^63,64^). Similarly, the “diverse club” identified from resting-state fMRI, defined by high-participation hubs whose connections span multiple communities^65,66^ and concentrated in the fronto-parietal control network, has its parietal nodes within lattice cortex and its prefrontal nodes within paralattice cortex. The lattice connectors do not correspond to any one canonical resting-state network. Paralattice connectors are spread over four of the seven networks of Yeo et al.^56^ (default, frontoparietal, dorsal attention and ventral attention), while peri-allocortex connectors are most enriched in the medial temporal subsystem of the default network.

### Relation to previous characterizations of cortical connectivity

The two blocks identified here can be compared with established descriptions of cortical functional organization. Canonical functional networks like Yeo et al.^56^ and Cole–Anticevic^67^ are defined from parcel-level, full-range connectivity, and therefore overlap only partially with the present partition (Table S7). The visual network and most of the somatomotor network fall within lattice cortex, as does most of the dorsal attention network. By contrast, the fronto-parietal control and default-mode networks span across the lattice/paralattice boundary, with their posterior parietal and precuneus components in lattice cortex and their prefrontal components in paralattice cortex.

Cortical organization has also been characterized using eigenvectors of graph Laplacians and related operators. These embeddings, however, have generally been analyzed as continuous axes over full-range connectivity, as in the principal gradient^55^. Connectome harmonics^68^ combine short surface connections and long-range tracts into standing-wave patterns that resemble resting-state networks. Geometric eigenmodes^26^ reconstruct cortical activity from shape-based modes; task-evoked maps are captured mainly by modes with wavelengths above ∼60 mm, which average over the short-range scale. A related model derives the structural connectome itself from geometric eigenmodes weighted toward long wavelengths, fitted to a human diffusion-MRI connectome at 4,386 vertices of one hemisphere^27^. Because such a model couples every point of the surface to its surroundings in the same way everywhere, it corresponds, at a coarser scale, to the geometric graph used here as the comparison for the empirical spectral decomposition. Tracer-eigenmode decomposition of marmoset cortex^69^ separates fMRI activity into wiring-coupled and decoupled areas along a smooth dorsal-posterior to ventralanterior axis.

Two previous studies similarly found short-range connectivity concentrated in sensory cortex and long-range connectivity in transmodal cortex. One^7^ classified voxels by short-vs. long-range (14 mm) connectivity into predominantly local (primary sensory and motor cortex), predominantly distant (lateral parieto-temporal and frontal cortex), and mixed (posterior cingulate and medial prefrontal cortex) classes. Another^70^ analyzed 200 regions and found greater long-distance connectivity in transmodal and paralimbic cortex and lower values in primary areas, closely aligned with the principal gradient. The present results extend these observations by showing that connectivity in lattice cortex remains a decreasing function of distance across the hemisphere, whereas in paralattice cortex it reaches a minimum just beyond the window and then rises again (Figure 3D1).

A recent precision-fMRI study shows that small, non-adjacent cortical patches can form time-ordered “chains” sharing a striatal target^71^; chains are densest in frontal cortex and sparse in primary sensory and motor cortex and posterior default-mode nodes, so that chain-rich regions may broadly correspond to paralattice cortex, which is coupled to thalamic nuclei receiving basal ganglia or cerebellar output (Tables S3 and S4), and chain-poor regions to lattice cortex. Together, these observations suggest a possible distinction between anterior cortical territories organized into multiple functionally segregated chains extending through cortico-basal ganglia-thalamo-cortical loops, and posterior-central territories organized as a highly integrated lattice^16^.

Beyond functional connectivity, several other dimensions of cortical organization show partial correspondence with the main partition (Table S5). Most of lattice cortex shows lower inter-subject variability, higher myelin content (T1w/T2w), and lower μ-opioid and 5-HT1A receptor densities. Paralattice cortex is more variable, has lower myelin content, and has higher μ-opioid and 5-HT1A densities, whereas GABA-A, 5-HT4, D2/D3 and serotonin-transporter densities differ little between the two blocks. Cytoarchitectonically, lattice cortex broadly overlaps Mesulam’s idiotypic and unimodal tiers and Barbas’ granular eulaminate cortex^72,73^. However, paralattice cortex corresponds less closely to heteromodal and dysgranular tiers, and the main partition can cross cytoarchitectonic boundaries. None of these correspondences is closer than what a rotation of the block map yields (Table S5), so they should be considered as purely descriptive.

Among the variables most strongly associated with the two blocks are cortical oscillatory dynamics. MEG delta power is higher in paralattice cortex, whereas alpha power is higher in lattice cortex; correspondingly, the decay time constant of regional MEG autocorrelation is longer in paralattice and shorter in lattice cortex^46^. To what extent these dynamical differences arise from the underlying organization of short-range connectivity remains an open question.

### Relation to short-range anatomical connectivity

The main partition rests on strong, short-range functional connectivity, potentially reflecting neural interactions mediated by dense intracortical fibers running within gray matter and short U-fibers arching beneath sulci to connect neighboring areas. These anatomical connections greatly outnumber long-range ones.

Interareal weights decline approximately exponentially with distance, making short connections dominant in the structural connectome^6,19^. Within an area, connection density also falls steeply with distance: in the macaque, about 80% of the neurons projecting to a cortical site lie in the same area, 80% of them within 1.2 mm and 95% within 1.9 mm^18^. Between pairs of pyramidal cells, connection probability in human cortex falls off with a lateral spread of about 130 μm^74^. Because the number of potential partners grows with distance, most excitatory synapses on a neuron come from within a few millimeters but from outside its immediate column: in cat V1, about three quarters of those near the axis of a 0.5–1 mm column arise from neurons outside it^75,76^. The bulk of the input to any cortical location thus arises within about 2 mm, mostly below the 3 mm lower limit of our window, which we set because of hemodynamic and vascular spread and of the spatial blurring introduced by acquisition and preprocessing (Supplementary Results S1); the 3–20 mm window captures the outer part of this distribution, the patchy horizontal connections linking columns several millimeters apart within an area^4,29,60^, and the short U-fibers to neighboring areas^48,49^.

Short-range connections are also distinctive in their organization and termination, mediating dense, direct interactions among neighboring neuronal groups. Neurons connected within fine-scale subnetworks share common inputs, whereas unconnected neighbors share little, and local wiring is highly nonrandom: a small number of strong, often reciprocal connections forms a skeleton within a much larger population of weak connections^77–79^. Such convergence can make a coherent local population more effective than an equal number of dispersed afferents, because co-active inputs on the same dendritic branch summate supralin-early^80,81^, while a few strong, like-tuned inputs can disproportionately shape stimulus-specific responses^77^. Long-range feedback fibers, by contrast, terminate over broader, more divergent fields than the focal terminations of short pathways^82^. An important exception is the corpus callosum: in visual cortex, callosal axons terminate in register with the visuotopic map, linking corresponding points of the two hemispheres^34,35^. As shown here (Figure 3F), homotopic callosal connectivity is highly correlated with strong, local connectivity.

Anatomical evidence also supports differences in short-range organization between lattice and paralattice cortex. Dense, local interactions are well established in the posterior-central territories that comprise most of lattice cortex. In primary visual and somatosensory areas, intrinsic horizontal connections link patches of neurons with similar response properties across neighboring neuronal groups in an orderly, topographic, distance-limited manner^29,30,60^, with callosal connections linking corresponding points of the visual map across hemispheres^34^. The resulting architecture resembles a hierarchical multi-grid: overlapping modality-specific sheets in which each location connects predominantly to its neighbors, extending from lower-to higher-order areas.

Paralattice cortex appears to follow a different organization, with disparate functional domains connected non-topographically over larger distances. In marmoset prefrontal cortex, labeled projections form many discrete, submillimeter columnar patches, about 1 mm apart, in nearby and distant regions^32^. In macaque prefrontal areas 9 and 46, intrinsic projections of supragranular pyramidal cells do not spread uniformly around the injection site but form narrow terminal bands separated by regions without terminations, spaced 500–600 μm apart and extending 7–8 mm across cortex. This contrasts with the more local, compact patches of V1, V2, V4, and motor and somatosensory areas^60,83^. The same injections also label associational clusters reached through white matter and arranged in stripes of similar dimensions^84^, while dual retrograde labeling indicates that intrinsic and associational axons arise from the same supragranular pyramidal neurons^85^. This organization may provide a cellular counterpart to the centimeter-scale pattern in Figure 3D1, where a paralattice vertex has more strong partners at long range than just beyond the window. The exponential distance rule, whereby cortico-cortical projection strength declines exponentially with distance^19^, may therefore apply most strongly within lattice cortex and less so within paralattice cortex, where beyond the short range the number of strong partners no longer declines with distance. Tractography provides convergent evidence: fiber-length distributions vary across human cortex, with very short fibers enriched in highly myelinated, thinner cortical regions, largely corresponding to lattice cortex, and longer fibers showing the opposite pattern^86^. Rodent data point in the same direction, although with important species differences^87^. However, inter-areal connectivity is substantially denser in rodents^88^, suggesting that the strongly local, neighbor-dominated organization of lattice cortex may be more pronounced in primates.

In primates, the lattice/paralattice contrast may also extend to single-neuron morphology. Prefrontal pyramidal neurons have larger, more branched, and more spinous dendritic trees than their sensory counterparts, allowing them to sample a broader and more heterogeneous set of inputs^89^. This organization supports mixed selectivity, whereby individual neurons respond to nonlinear combinations of multiple variables, enabling populations to represent arbitrary conjunctions in a high-dimensional format suited to flexible behavior^90,91^. In humans, layer-2/3 pyramidal neurons have especially elaborate dendritic arbors and form networks of higher topological dimension than in mouse, a structural property proposed to increase computational capacity and flexibility^92^. Lattice cortex, by contrast, is dominated by neighborhood relations, an organization well matched to the statistics of sensory inputs: signals vary smoothly across space and time, making a lattice of short connections among neighboring units an economical architecture for capturing this structure.

The organization of paralattice cortex is also consistent with the functional architecture of prefrontal cortex into partially segregated, closed loops through the basal ganglia and thalamus, which preserve distinct functional channels rather than blending neighboring representations^41^. Prefrontal representations can be sustained and flexibly routed through thalamocortical interactions in a context-dependent manner^93,94^. This organization is also consistent with the recent characterization of much of paralattice cortex as a mosaic of small, non-adjacent patches linked into functional chains^71^, and with the anatomical correspondence identified here between paralattice cortex and specific thalamic nuclei receiving pallido-nigral or cerebellar input, in contrast to lattice cortex and specific thalamic nuclei that lack such input (Table S3). Our functional connectivity results follow the same axis: thalamic nuclei receiving basal-ganglia or cerebellar output (ventral anterior, ventral lateral) couple preferentially with paralattice cortex and the striatum, whereas first-order sensory relays and the pulvinar couple with lattice cortex but not the striatum. Therefore, paralattice cortex participates preferentially in closed cortico-basal-ganglia-thalamo-cortical loops, whereas lattice cortex, with a few exceptions, notably primary motor cortex (Table S3), does not.

Finally, the peri-allocortex, which was excluded from the analysis, has an anatomical organization distinct from both lattice and paralattice cortex: reduced laminar differentiation, including regions in which layer 4 is sparse or absent, and transitional architectures between neocortex and hippocampal or limbic structures^72,95^. When peri-allocortex is not excluded, however, the two-block partition is nearly the same (Table S6): the eigengap still falls at K = 2, and the peri-allocortex joins the two blocks, four fifths in lattice cortex. The next eigenmode separates medial temporal, temporopolar and inferior temporal cortex into a third cluster, without any marked eigengap.

### The bottleneck as a border between a perceptual and an action hierarchy

Lattice cortex extends forward to include primary motor cortex and most of caudal dorsal premotor cortex, and the border runs between these and rostral premotor and prefrontal cortex. Macaque tracing studies indicate the premotor strip as a connectional transition zone: F2 and F3, the caudal premotor and supplementary motor areas, are linked with F1, with the cingulate motor area and with somatosensory and superior parietal cortex, and receive little or no prefrontal input, whereas the rostral fields F7 and F6 are linked with area 46 and have meager parietal connections^96–98^. Their thalamic afferents shift in the same direction: F3 and F2 receive mainly from the ventral lateral and ventral posterior lateral nuclei that also supply F1, whereas F6 and F7 receive mainly from the parvocellular ventral anterior nucleus and area X; all receive a strong mediodorsal projection, but only for F6 and F7 does it arise from the mediodorsal territory that also projects to prefrontal cortex^99^.

Classical accounts place primary sensory and primary motor cortex at the bottom of two hierarchies, one posterior ascending through unimodal to heteromodal association cortex and one anterior descending from prefrontal to premotor to motor cortex^72,100^. Importantly, however, the frontal hierarchy is not the mirror image of the posterior one, since its intrinsic connections are striped and discontinuous rather than lattice-like.

To effectively interact with a complex environment, high-level action plans should be guided by high-level perceptual categorization^72,100^. Accordingly, the upper levels of the two hierarchies should interact directly rather than through chains of neighbor-to-neighbor connections at the lowest levels. This implies that, if the two hierarchies expand and diverge during evolution, the connections between the highest levels should stretch correspondingly. In fact, successive posterior stages project to progressively more rostral frontal targets^101^. Conversely, short-range connections between the lower levels of the two hierarchies would become progressively less relevant, which could contribute to the origin of the bottleneck.

From this perspective, the sites of termination of the lattice connectors may reflect the location of the upper levels of the frontal hierarchy. As shown above, their input is densest in caudal and mid-lateral prefrontal cortex, including area 46 and the 9-46 complex, and sparse in orbitofrontal cortex and in the cingulate motor areas at the foot of the slope. This agrees with functional evidence that places the apex of the frontal hierarchy in mid-lateral prefrontal cortex rather than in the frontal pole^102,103^ (rostral area 10 is a partial exception, as 10d, and on the right p10p, receive substantial input). The projections run in parallel: the anterior inferior parietal lobule, the intraparietal sulcus and the anterior precuneus (PF, PFt, AIP, 7Am) reach mainly 6r, 6a, 46 and IFSa, whereas the angular gyrus and posterior medial parietal cortex (PGi, PGs, 7m, 31pv) reach mainly 8Ad, 8Av and 9m, recalling the segregated parieto-frontal channels traced in the macaque^104,105^.

### Limitations of the study

Using fMRI connectivity to probe local neuronal interactions has clear limitations, foremost spatial and temporal resolution. Vertex spacing at fs_LR 32k (∼2 mm) exceeds the scale of a macrocolumn (300–500 μm) and far exceeds that of a minicolumn (30–80 μm); hemodynamics further blur the signal, and the relevant neuronal timescale (tens to hundreds of milliseconds) lies beyond fMRI resolution. (Sub-millimeter fMRI can resolve resting-state connectivity between ocular dominance columns in human primary visual cortex^106^, but only over a small part of the brain.) Nevertheless, 7T fMRI resolves lattice and paralattice cortex reproducibly and identifies modality streams and their hierarchy within lattice cortex.

The fMRI signal is also weak in orbitofrontal cortex, the temporal pole and anterior ventral temporal cortex, where the nearby air-filled sinuses and ear canals cause signal dropout. These regions retain very few strong short-range edges, so their assignment to one block or the other rests on scarce evidence: orbitofrontal cortex falls in paralattice cortex, and the temporal pole and anterior ventral temporal cortex fall in lattice cortex, at the end of the auditory-temporal stream. The main partition is robust to the removal of these regions, but their own assignment to one block or the other should be considered provisional.

Most analyses were performed on cohort-level 7T data, and individual-subject estimates are necessarily noisier. However, the PAN replication shows that long, high-quality acquisitions can recover the partition in individual subjects. This raises the possibility of detecting altered blocks or boundaries at the individual level, potentially including changes associated with specific pathological conditions such as dissociative disorders.

The present data also cannot resolve potential laminar differences in functional connectivity. Whole-cortex laminar fMRI may eventually determine whether the lattice/paralattice partition primarily reflects the organization of dense, short-range horizontal supragranular connectivity rather than sparser, longer-range infragranular intra-telencephalic connections. Another question is whether the partition observed in human short-range functional connectivity extends to other species, particularly non-human primates, and perhaps less strongly to rodents given their more compact connectivity^88^. Preliminary analysis of high-quality 9.4T resting-state data from marmoset monkeys suggests that an analogous main partition can also be recovered, although with a smaller proportion of paralattice cortex (unpublished results).

An obvious limitation is that functional connectivity can only be taken as an indirect, imperfect, and partial reflection of neural interactions along the anatomical connectivity. A subject-specific null floor (as used here) or global-signal regression (as a further control with its own limitations^107–109^, Table S6) are intended to reduce but cannot completely eliminate contributions from non-neural sources and diffuse common input. On the other hand, strong, short-range, vertex-level functional connectivity may at least partially reflect the synergistic effects of neurons sharing inputs and outputs across short recurrent loops.

Even so, pairwise correlations cannot be interpreted as direct causal interactions, provide no information about the direction of the interactions, and cannot establish whether interactions among multiple vertices are specific (informative) or redundant. Nevertheless, a hierarchical, multi-grid lattice supported by dense local interactions among neuronal groups and demarcated from cortical and subcortical territories with more modular organization provides a potential anatomical substrate for high information integration^15,16^.

## Supporting information

Supplemental data: subject lists

## DATA AND CODE AVAILABILITY

- The HCP Young Adult data are available from the Human Connectome Project (https://www.humanconnectome.org) under the HCP data use terms. The PAN data are available from OpenNeuro (accession ds006598, version 2.0.1, raw data; https://doi.org/10.18112/openneuro.ds006598.v2.0.1).
- The subject lists of every cohort, in the order used by the analysis code and with the excluded subjects and the reason for each exclusion, are provided as Data S1. The public inputs, the definitions and the check values of every step are listed in Table S8.
- Derived group-level data (cohort graphs, cohort-median distance matrices and block labels) and the analysis code, with the pinned software environment, will be deposited at time of publication. Per-subject HCP-derived data are not redistributed, in accordance with the HCP data use terms.
- Any additional information required to reanalyze the data reported in this paper is available from the corresponding authors upon request.

## ACKNOWLEDGMENTS

This work was made possible through support from the Tiny Blue Dot Foundation, the Templeton World Charity Foundation, the Distinguished Chair in Consciousness Science and the David P. White Chair in Sleep Medicine at the University of Wisconsin-Madison. L.M. was supported by Borse di studio e ricerca per la mobilità internazionale (Collegio Superiore, Bologna).

## AUTHOR CONTRIBUTIONS

Initial conceptualization, G.T., M.B., S.S.; Final conceptualization, G.T., M.B., K.D., G.F., L.A., C.C., W.M., and O.S.; Methodology, M.B., S.S., M.J., L.M., R.M.B., O.S., and G.T.; Formal analysis, M.B., W.M., G.T.; Validation, M.B., M.J., L.M., K.D., G.F., R.M.B., and G.T.; Writing – initial draft, M.B., S.S., and G.T.; Writing – final draft, G.T., M.B., M.J., and L.M.; Review & editing, all authors; Supervision, R.M.B., M.B., and G.T.; Funding acquisition, G.T.

## DECLARATION OF GENERATIVE AI AND AI-ASSISTED TECHNOLOGIES IN THE MANUSCRIPT PREPARATION PROCESS

During the last phases of the preparation of this work (since early 2026, but not in the previous several years), the authors used Claude (Anthropic) to write and run analysis code, draft figures, and generate tables; its use in the analyses is described in the Methods (Software and reproducibility). To aid with the organization of project material, Flux software was also used (fluxsci.org). The main text was written by the lead authors, and Claude was used only to insert updated values and minor edits suggested by co-authors. Parts of the Methods (primarily tables) were drafted and tested with Claude to ensure full independent reproducibility based on Methods alone. The authors take full responsibility for the content of the published article.

## METHODS

### Key resources table

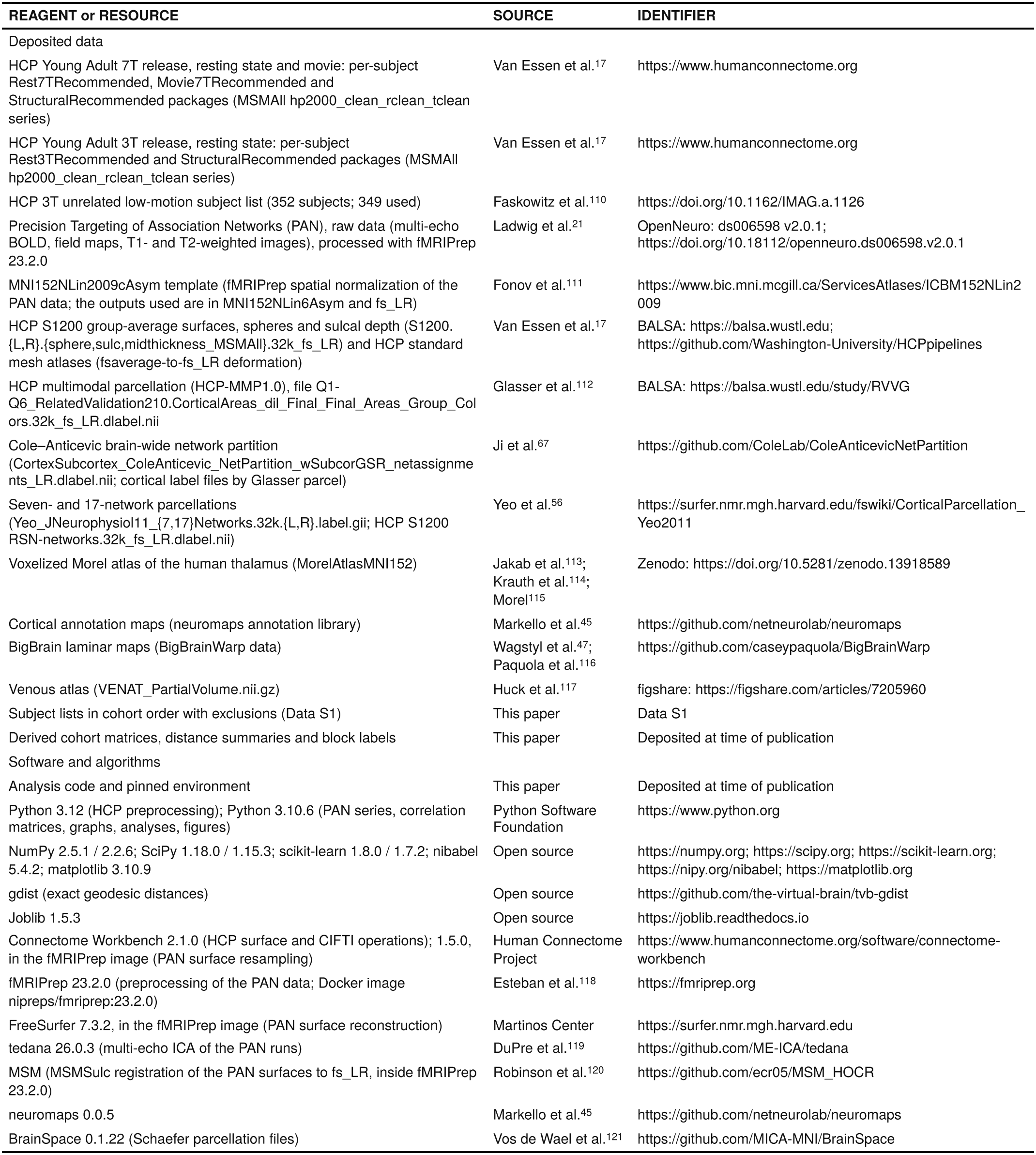

### Experimental model and study participant details

We used MRI data from three publicly available datasets: the Human Connectome Project (HCP) 7T and 3T releases, as minimally preprocessed by the HCP, and the Precision Targeting of Association Networks (PAN) dataset, whose raw data we processed ourselves (Data preprocessing). The HCP and PAN studies were approved by the institutional review boards of the institutions that acquired the data, and all participants gave written informed consent; the de-identified data were used here under the data use terms of each release. The influence of sex was examined for the main partition, by analyzing the women and the men of the HCP 7T cohort separately (Hyperparameter and robustness sweeps); sex was not otherwise examined. Sex was taken as recorded by the HCP; gender, ancestry, race and ethnicity were not analyzed.

#### HCP 7T

From 175 candidate subjects of the HCP Young Adult 7T release (the included subjects are listed in Data S1), we excluded participants based on the following criteria: (1) incomplete functional data acquisitions and (2) excessive head motion, defined as more than 50% of the time frames censored due to a framewise displacement (FD) > 0.25 mm. One further subject was excluded from the resting-state cohort for a run-level data issue listed by the HCP, and one subject (395756) could not be processed. Of the 24 subjects excluded from that cohort, 8 had incomplete acquisitions, 14 excessive motion, 1 the HCP-listed issue and 1 could not be processed (Data S1). Following exclusion, the 7T resting-state cohort analyzed here consisted of 151 subjects (89 females, 62 males, age range: 22–36+ years). The movie cohort consisted of 160 subjects (97 females and 63 males, age range 22–36+ years). In the movie cohort, 6 subjects were excluded due to incomplete acquisitions and 9 to excessive motion (the criteria were applied to each acquisition separately, so the two cohorts differ by 11 subjects). The functional data were acquired using a 7T gradient-echo EPI (GE-EPI) sequence (1.6 mm isotropic voxels, TR = 1.0 s). Both resting-state and movie-watching fMRI data comprised four runs acquired over two days, alternating phase-encoding directions each day (rest: PA and AP on each day; movie: AP and PA on day 1, PA and AP on day 2), totaling approximately 1 hour (3,600 TRs for rest, 3,655 for movie) of scanning. As is standard for the HCP pipeline, structural MRI data were derived from the subjects’ corresponding 3T 3D MPRAGE T1-weighted images with 0.7 mm isotropic voxels. Detailed acquisition parameters are provided in Table S1.

#### HCP 3T

We included 349 participants (182 females, 167 males, age range: 22–36+ years) from the preprocessed HCP Young Adult 3T release. We adopted the subject list and quality control criteria used by Faskowitz et al.^110^, which excluded subjects based on: (1) data quality issues (e.g., structural anomalies, surface segmenta-tion errors, head coil complications, or data requiring ICA-FIX manual reclassification), (2) excessive head motion (more than 50% of time frames censored at FD > 0.25 mm), and (3) familial relatedness. The list, of 352 subjects, was established by Ito et al.^122^; three of them (130114, 329844 and 908860) were not used, because the HCP lists all their 3T RL runs as processed with the wrong phase-encoding direction and recommends excluding them from analyses of the 3T functional data; the other 349 were all used (Data S1). Forty-three of the 349 are also in the 7T cohort. Structural MRI consisted of high-resolution 3D MPRAGE T1-weighted images (0.7 mm isotropic voxels). For functional data, we used only resting-state fMRI, acquired with a GE-EPI sequence (2.0 mm isotropic voxels, TR = 0.72 s). The resting-state data were collected in four runs over two days, two per day with opposite phase-encoding directions (day 1: RL, then LR; day 2: LR, then RL, the order used by the HCP from October 2012 onward), yielding approximately 57 minutes (4,800 TRs) of data per subject (detailed parameters in Table S1).

#### PAN

We included all 10 highly sampled participants (5 females, 5 males, age range: 18–33 years, mean age = 26.4 ± 4.8 years) from OpenNeuro published by Ladwig et al.^21^ (OpenNeuro ds006598, version 2.0.1, raw data). Structural MRI consisted of T1-weighted images acquired with 1.0 mm isotropic voxels and T2-weighted images, both used for surface reconstruction. Functional data were acquired on 3T Siemens Prisma scanners using a multi-echo multiband sequence (2.4 mm isotropic voxels, TR = 1.355 s) with an anterior-to-posterior (AP) phase-encoding direction. Detailed acquisition parameters are provided in Table S1.

### Method details

The processing chain from the per-subject correlations to the partition is summarized in Figure S2, and its parameters are listed in Table S9.

#### Data preprocessing

##### HCP 7T and 3T

For the HCP cohorts, we used the minimally pre-processed GIFTI and CIFTI data releases (the ICA-FIX-denoised, MSMAll-aligned series of the four runs concatenated, as released; file names in Table S8). The minimal preprocessing pipeline includes distortion and B1 field correction, motion correction, T1w/T2w alignment, nonlinear registration to MNI space, cortical surface segmentation and reconstruction, surface projection and resampling, and smoothing of the grayordinate series on the surface with a 2-mm FWHM Gaussian kernel (the pipeline setting for the 2-mm grayordinates used here)^123^. The released series (2025 release of the HCP Young Adult data, suffix _hp2000_clean_rclean_tclean) are high-pass filtered (hp2000), denoised by spatial ICA with FIX classification (clean) and its revised classification (rclean), and cleaned of global and semi-global structured noise by temporal ICA (tclean), which is designed to keep the global neural signal^124^; they are aligned across subjects by MSMAll, a surface registration driven by both anatomy and function. In the projection to the surface, voxels with an unusually high temporal coefficient of variation are excluded^123^. These are voxels whose coefficient exceeds the mean of their neighborhood (Gaussian, σ = 5 mm) by more than 0.5 standard deviations, mostly voxels that contain large vessels or lie at the edge of the parenchyma. In addition to the minimal preprocessing, we censored time frames with FD greater than 0.25 mm and linearly interpolated these frames from their neighboring frames for the nuisance regression; they were then removed, so that all correlations used retained frames only. FD is the sum of the absolute frame-to-frame translations (mm) and rotations (degrees, converted to arc length on a 50-mm-radius sphere), computed from the HCP movement regressors. No frames were discarded at the start of the runs, as the released series begin at steady state. We then regressed out an intercept and linear and quadratic trends over the concatenated runs and the mean white-matter and CSF signals, in one least-squares fit and without band-pass filtering (Table S8). The white-matter labels include the white matter within 5 mm of the cortex, so this regressor also carries some cortical signal. Finally, the cleaned BOLD signals were z-scored over the concatenated runs of each subject. No global signal was regressed; the control with global-signal regression is described under Hyperparameter and robustness sweeps.

##### PAN

The raw data (five echoes per frame, field maps, T1- and T2-weighted images) were processed with fMRIPrep 23.2.0^118^, the version the dataset’s authors used, with outputs in MNI152NLin6Asym (2 mm) and as CIFTI on the fs_LR 32k mesh (91k grayordinates), slice-timing correction off (Table S8). fMRIPrep reconstructed each subject’s cortical surfaces with FreeSurfer 7.3.2, registered them to fs_LR 32k by MSMSulc, and sampled the optimally combined BOLD series of each run on the subject’s own mid-thickness surface, without the exclusion of high-variation voxels applied to the HCP data. In each run, the initial frames that fMRIPrep flagged as not yet at steady state (one to three frames in 241 of the 250 runs) were dropped. Each run was denoised by multi-echo independent component analysis (ME-ICA) with tedana 26.0.3119,125, which separates the signal components whose amplitude scales with echo time, as the BOLD signal does, from those whose amplitude does not (motion, scanner and other non-BOLD artifacts). tedana was run with its default settings on the five echo-wise series of the run as preprocessed by fMRIPrep (native space, steady-state frames, fMRIPrep brain mask; Table S8). Frames with framewise displacement (fMRIPrep, the same definition as for HCP) above 0.25 mm were censored, replaced by linear interpolation from their neighboring frames for the nuisance fit and then removed, leaving 105 to 163 minutes of data per subject. An intercept, a linear and a quadratic trend, fMRIPrep’s mean white-matter and CSF signals and the time courses of all ME-ICA components entered one least-squares fit of each vertex’s series. The fitted contributions of the trend, the white-matter and CSF signals and the rejected components were removed; the accepted components, and any variance they share with the rejected ones, were left in the data (“non-aggressive” denoising strategy119); because the PAN runs are short (222 frames) and spread over several sessions, this fit was made run by run rather than over the concatenated runs as for HCP. As for the HCP series, no global signal was regressed. As controls, the analysis was repeated without ME-ICA (the same fit without the ICA components), without ME-ICA and with the six fMRIPrep motion parameters and their temporal derivatives added to the nuisance fit, and with ME-ICA and the global signal additionally regressed out, as for the HCP series (Table S6). The retained frames of all runs were concatenated in session and run order and z-scored once over the concatenation, so that single runs were not rescaled (4,634 to 7,223 frames per subject). Vertices with no signal in any run (430 to 621 per subject; all but 0 to 9 of them in the peri-allocortex, mostly the hippocampus, presubiculum and entorhinal cortex) were kept at zero, so that they correlated with no other vertex. fMRIPrep was run without a fixed random seed; the ICA of tedana used its default fixed seed.

#### Geodesic distances

Per-subject geodesic distances for each vertex on the fs_LR 32k surface were calculated for the HCP 3T and 7T subjects and for the PAN subjects. Using each subject’s native-space mid-thickness surface on the fs_LR 32k mesh (HCP-provided GIFTI files; Table S8), we extracted the triangulated mesh and computed the exact shortest path between two vertices using the *gdist* Python package (key resources table), which implements the Mitchell-Mount-Papadimitriou (MMP) exact geodesic algorithm. The medial wall (the vertices outside the HCP cortex mask, 2,796 L and 2,776 R of the 32,492 vertices) was excluded from the mesh to prevent any paths from routing through non-cortical areas; the peri-allocortex was kept in the mesh, so paths could cross it. The computation was run in parallel over vertices with the *Joblib* Python package (key resources table). For PAN, each subject’s fMRIPrep mid-thickness surface was first resampled to the fs_LR 32k mesh through its MSMSulc sphere (Connectome Workbench, barycentric), and distances up to 40 mm were then computed in the same way and on the same cortex mask.

Every analysis reported was performed within a hemisphere, on that hemisphere’s own triangulated mesh.

For each pair of vertices, the cohort geodesic distance is the median of the subjects’ individual geodesic distances (Table S8). This distance was used for the window, long-range cutoff, the distance profiles and the patch measures. The window is the short-distance range used to build the short-range graph, 3 ≤ *d* < 20 mm (Pairs, neighbors and partners; Edge-survival probability model). Each cohort used the median over its own subjects: the 151 subjects of the primary HCP 7T resting-state cohort, the 349 subjects of the HCP 3T cohort, the 160 subjects of the HCP 7T movie cohort, and the ten PAN subjects.

#### Connectivity, null floor and cohort graph

For each subject, the Pearson correlation *r* (not Fisher-transformed nor rectified) between the cleaned BOLD series of every pair of cortical vertices was computed after censoring of high-motion frames as described above.

##### Pairs, neighbors and partners

A candidate pair is a within-hemisphere vertex pair with neither vertex in the peri-allocortex (Cohort graph); *P* denotes the set of candidate pairs. The window candidates are the candidate pairs whose cohort geodesic distance *d*^*ij*^ lies in the window (Geodesic distances), *c* _win_ ={ ( *i, j* ) ∈ *P* : 3 ≤ *d*^*ij*^ < 20} . The neighbors of a vertex *i* are the vertices with which it forms a window candidate, *c* _win_ ( *i* ) = {*j* : ( *i, j* ) ∈ *c* _win_ }. An edge is a candidate pair that passes the null floor and the support rule (Cohort graph); *E* denotes the set of edges at all distances (the full-range graph), and the two vertices of an edge are partners. The window edges are the window candidates that are edges, *E* _win_ = {( *i, j* ) ∈ *E* : 3 ≤ *d* ^*ij*^ < 20 }; they form the short-range (window) graph. The window partners of a vertex *i* are its neighbors to which it is joined by a window edge, *N*^*i*^ = {*j* : ( *i, j* ) ∈ *E* _win_ } ⊆ *c* _win_ ( *i* ), and its window degree is their number, |*N*^*i*^|. Edges with one vertex in the peri-allocortex are defined by the same floor and support rule and enter only the long-range and connector analyses (Cohort graph).

##### Null floor

Each subject’s correlations were compared with that subject’s own null distribution: the correlations of 800,000 heterotopic vertex pairs, drawn once (with replacement) and the same for every subject, joining occipital cortex (V1–V4) to the contralateral frontal pole and to contralateral sensorimotor cortex, pairs with no plausible anatomical connection. The subject’s floor *τ*^*s*^ is the smallest correlation value at which the estimated false-discovery rate falls to *q* = 0.05. The estimate is the fraction of null pairs above the floor divided by the fraction of window candidates above the 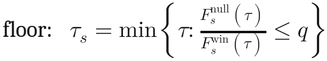, where 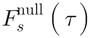 and 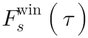 are the fractions of null pairs and window candidates whose correlation in subject *s* exceeds *τ*. The window candidates of the two hemispheres were pooled, so that each subject has a single floor. The graphs and the partition were then computed for each hemisphere separately. In each subject, the floor determines the support, to build the cohort graph (see Cohort graph).

##### Cohort graph

The support of a candidate pair (Pairs, neighbors and partners) is the number of subjects, out of the *N* subjects of the cohort (151 for the HCP 7T resting-state cohort), in which the pair’s correlation *r*^*s, ij*^ (vertices *i* and *j*, subject *s*) exceeds that subject’s null floor *τ*^*s*^, *S*^*ij*^ = | *s*: *r*^*s, ij*^ > *τ*^*s*^ |. A candidate pair is connected with an *edge* of the cohort graph when its support is a majority of the cohort (⌊*N* / 2⌋ + 1 subjects, e.g., 76 or more of 151), *E* = { (*i, j* ): *S*^*ij*^ ≥ ⌊*N* / 2⌋ + 1 }. The weight *w* of an edge is the mean correlation over all N subjects of the cohort, 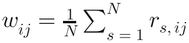, and only pairs with *w* > 0 are kept as edges (in the cohorts analyzed here, this condition removed no edge). The short-range (window) graph consists of the window edges. The full-range graph was created by applying the same rule to every within-hemisphere pair. The peri-allocortex is defined by whole HCP-MMP1.0 areas encompassing the hippocampal formation, entorhinal, presubicular and perirhinal cortex, piriform cortex, area 25, the parahippocampal areas, TF, s32, 33pr and the agranular anterior insula (Table S8; 3,536 vertices). It was excluded from floor calibration, null estimation, the window graph and all partitions. Support and mean *r* were nevertheless accumulated, with the same floors, for every pair of cortical vertices, including pairs with a peri-allocortex end and pairs across the hemispheres. These values give the full-range edges into the peri-allocortex, by the same floor and majority rule as for every other edge; they are used for the long-range and connector analyses (with them the full-range graph has 21,310,970 L and 22,375,674 R edges), and the homotopic edges. As a control, the subject floors and the majority support were replaced by a single cohort-wide threshold on mean *r* that keeps the same number of window edges (set for each hemisphere separately; *r* = 0.126 in both). The comparison is reported in Supplementary Results S2.

##### Area weighting

The surface area each fs_LR 32k vertex stands for varies (Supplementary Results S2). Therefore, we defined and applied the following area weights. For each subject s, the area of every triangle of the mid-thickness surface was divided equally among its three vertices, so the area of vertex *i* was one third of the summed area of the triangles that meet at it on the subject’s mid-thickness surface, 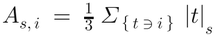, where |*t*|^*s*^ is the area of triangle *t* in subject *s*. The relative area *a*^*i*^ is the mean of *A*^*s, i*^ over the *N* subjects, divided by the mean of this quantity over the vertices of the same hemisphere outside the peri-allocortex, *V*^*h*^:

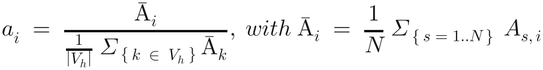

so that *a*^*i*^ averages 1 in each hemisphere (check values in Table S8).

For every cut and spectrum, each edge weight was multiplied by the areas of its two vertices, 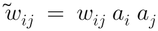, the lumped-mass discretization of the cortical sheet used for the mass matrix of geometric eigenmodes26. With this weighting, the volume of a block in the normalized cut measures cortical area rather than vertex count. The weighting enters the partitions, their spectra and restart profiles, and the edge probability *p* ( *d* ) . Quantities that concern the connection between two vertices (the floor, the support rule, thresholds, the ranking of partners and shortest paths) use the cohort weight *w* without it. Without the area weights the partition is nearly the same (Supplementary Results S2; Table S6).

##### Strip across the border

The vertex-level matrices of Figures 1B and 1D are drawn on a strip of 110 left-hemisphere vertices that runs from the interior of one block, across the border, into the interior of the other (Figure 2B). The strip was built from a seed and two anchors. The seed is a border vertex chosen by hand, in area 55b where it meets area 4 (border vertices as defined under *Distance to the border*; vertex indices in Table S8). One anchor was then placed in each block, 31–37 mm from the seed and well inside the block (27 and 34 mm from the border): one in area 3b and one in area 8Av, 71 mm apart. A vertex belongs to the strip when *d* ( *v, A* ) + *d* ( *v, B* ) ≤ *d* ( *A, B* ) + *width*, with *d* geodesic, *A* and *B* the anchors and *width* the strip width; the strip is thus a geodesic ellipse with the anchors as foci. Its vertices were ordered along the strip by *d* ( *v, A* ) − *d* ( *v, B* ), from the lattice anchor to the paralattice anchor, and thinned in that order, keeping a vertex only if it lay at least 5 mm from every vertex already kept. The width was raised from 2 mm in steps of 0.5 mm until at least 110 vertices survived the thinning (width = 13 mm), and the survivors were subsampled at evenly spaced ranks to 110. Counted from the lattice anchor, the vertices change block between the 49th and the 64th, where motor cortex meets premotor cortex and the frontal eye field (the strip is 13 mm wide, so vertices of the two blocks alternate along this stretch); the arrowheads in Figures 1B and 1D mark the middle of this stretch.

Figure 1B shows the representative subject of Figure 1A (572045): raw *r* for all pairs below the diagonal and, above it, only for the pairs above that subject’s floor (see Null floor). Figure 1D shows the cohort: the mean *r* of the edges of the full-range graph, *w*, below the diagonal, and the same restricted to 3 ≤ *d* < 20 mm above it (*τ*^*s*^ is defined under Null floor; the full-range graph is defined under Cohort graph, the two blocks under Spectral partitioning).

#### Distance-dependent regimes of connectivity

##### Edge-survival probability model

The boundary of the short-range connectivity regime, from which the window graph is built, was validated by fitting a two-component model to the distance decay of the edge-survival probability *p* ( *d* ), the probability that two cortical locations at distance *d* are joined by an edge, each vertex pair counted with the areas of its two vertices (Results, “Functional connectivity contains a distinct short-range component”; Figure 1C).

Distances are cohort geodesic distances from 0 to 300 mm, in 1-mm bins (⌊ *d*⌋ is the floor function applied to distance *d* ) . *P* is the set of candidate pairs and *E* the set of edges of the full-range cohort Graph (Pairs, neighbors and partners), and *P*^*d*^ = {(*i, j*) ∈ *P* : ⌊*d*^*ij*^ ⌋= ⌊*d*⌋} are the candidate pairs in the bin of *d. p* ( *d* ) is the fraction of these pairs that are edges:

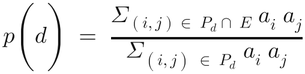

We compared a single exponential with the sum of two exponentials:

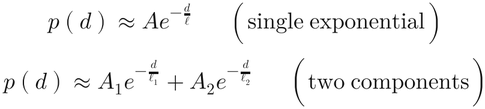

Each *l* is a length constant, the distance over which a component falls by a factor of *e*, and each *A* is the component’s amplitude, its value extrapolated to *d* = 0.

Both models were fit by least squares to log*p* ( *d* ), with each 1-mm bin weighted by the denominator of *p* ( *d* ), its area-weighted number of candidate pairs, divided by the mean over the fitted bins. Bins with *p* ( *d* ) = 0 were excluded. The single exponential was started from the best length constant on a grid from 2 to 400 mm in steps of 0.05 mm, with its amplitude fitted in closed form at each value. The two exponentials were started from every combination of *l*_1_ ∈ {5, 10, 15, 17, 20, 25, 35} mm, *l*_2_ ∈ {60, 100, 150, 220, 400, 800, 2,000, 10,000} mm and A_2_ ∈ {0.01, 0.03, 0.06, 0.1} (224 starts), with A_1_ started at the amplitude of the single-exponential fit. Each start was refined by the Nelder–Mead algorithm on the logarithms of the parameters, which keeps them positive, and the fit with the lowest weighted sum of squared residuals was kept. The two components were then labeled so that *l*_1_ < *l*_2_. The models were compared by the Bayesian information criterion (Statistical significance tests). In both hemispheres *l* _2_ grew without bound during fitting, so the long-range component is effectively constant. The fitted values and the distance at which the two components cross (the crossover) are given in the Results and in Table S8.

To find which connections produce the rise of *p* ( *d* ) beyond 200 mm (Results), each edge of the full-range cohort graph at d ≥ 200 mm was assigned to the pair of HCP-MMP1.0 areas that contain its two vertices, and the area pairs were ranked by their share of these edges, each edge counted with its weight *a*_*i*_ *a*_*j*_ as in *p* ( *d* ) . On the left, the twelve leading pairs each join PGi, PGs or STSvp to one of six dorsomedial and frontopolar prefrontal areas (9m, 10d, 9p, 9a, 8BL, 8Ad). TE1p, which adjoins STSvp and holds the next pair with these prefrontal areas (8Ad–TE1p, sixteenth), was added to the first group. On the right, nine of the fifteen leading pairs join the same two groups, and the others join inferior parietal and intraparietal areas (PGs, PGi, PFm, IP1) to p10p and a47r. *p* ( *d* ) was then recomputed without the vertex pairs that join the two groups, which were removed from the numerator and from the denominator (Table S8).

##### Short- and long-range bounds

The window 3 ≤ *d* < 20 mm (Geodesic distances) agreed with this fit: its outer bound lay at about one short-range length constant (1.0 *l* _1_ L and 0.9 *l* _1_ R), where the short-range component still exceeded the long-range one about tenfold, and its inner bound at 3 mm excluded the range in which the hemodynamic point-spread function and the volume-to-surface projection can produce correlations on their own (Supplementary Results S1). Long-range connections were defined as those of 60 mm or more, close to the crossover of the two components (65 mm L, 71 mm R). Beyond this distance, edge probability was dominated by the distance-independent component. Pairs at *d* ≥ 60 mm form the far band, used for the near/far ratio (Figure 3D2). The edges of the full-range cohort graph whose cohort geodesic distance is at least 60 mm are called *long-range edges, E*_*far*_ = {( *i, j* ) ∈ *E*_*ij*_ : *d* ≥ 60 mm} throughout; they define the long-range graphs and the connectors (Figure 4E–I). The fraction of the fitted short-range component that falls within 20 mm is 1 − exp(−20/*l*_1_).

##### Mean correlation profile

The mean correlation profile 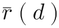 (Results) was computed over the candidate pairs closer than 20 mm as:

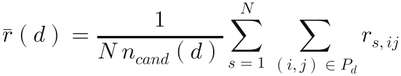

the mean correlation *r* of all within-hemisphere pairs outside the peri-allocortex in each 1-mm bin of cohort geodesic distance, first over the *n*_*cand*_ ( *d* ) pairs of the bin and then over the *N* = 151 subjects; *r*_*s, ij*_ is the unthresholded correlation of pair ( *i, j* ) in subject *s*, before any floor is applied. The value for random pairs is the mean *r* of 1,000,000 within-hemisphere pairs outside the peri-allocortex drawn with replacement (draw in Table S8), averaged over the subjects (mean *r* = 0.02).

##### Short-range density

The summed weight of a vertex’s window edges is 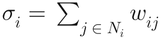 (*Pairs, neighbors and partners*) and its short-range density is this summed weight divided by its number of neighbors, 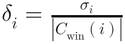 the weight per neighbor of the Results (Figure 1E); its unweighted density is the fraction of its neighbors that are window partners, 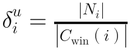. The two maps were compared by the Spearman correlation over the vertices of each hemisphere outside the peri-allocortex with at least 20 neighbors (*ρ* = 0.98 L and R). The block values in the Results (about three quarters and about half of the neighbors) are the medians of the unweighted density over the vertices of each block (Table S8). The regional values in the Results are medians over the vertices of the HCP-MMP1.0 areas listed in Table S8.

#### Spectral partitioning

Each hemisphere was partitioned on its own window graph by the normalized cut criterion of Shi and Malik^24^, applied to the giant connected component (the largest set of vertices all linked to one another by edges); vertices outside it were left unassigned. Let *W* be the weighted adjacency matrix of the area-weighted window graph (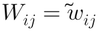 for pairs joined by an edge and zero otherwise; every cut and spectrum used weights multiplied by the vertex areas, see Area weighting) and *D* the diagonal matrix of its weighted degrees, *D*_*ii*_ = ∑_*j*_ *W*_*ij*_. For a split of the vertices into blocks *A* and *B*, the normalized cut is

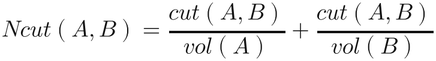

where *cut* ( *A, B* ) = ∑_*i*_ ∈ _*A, j*_ ∈ _*B*_ *W*_*ij*_ is the total weight of the edges between *A* and *B*, and *vol* ( *A* ) = ∑_*i*_ ∈ _*A*_ *D*_*ii*_ is the volume of *A*, the total weight of the edges at its vertices, each edge inside *A* counted once from each end; each term is thus the fraction of a block’s edge weight that crosses the border, and dividing by the volumes penalizes cuts that detach small sets. Since the exact minimization is NP-complete, the problem is relaxed to real values, where the minimizer is obtained from the second eigenvector of the symmetric normalized Laplacian *L* = *I* − *D*^− 1 / 2^*WD*^−1 / 2^. *L* was decomposed on the giant connected component (ARPACK through SciPy, shift-invert about −10^−6^, the eight smallest eigen-pairs; settings in Table S8), and the Fiedler vector was taken as *v*_2_ = *D*^−1 / 2^*u*, where 0 = *λ* ≤ *λ*_2_ ≤ … are the eigenvalues of *L* and *u*_2_ the eigenvector associated with *λ*_2_. The number of blocks was read from the eigenvalue ratios 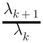 computed from the first eleven eigenvalues, obtained in a separate computation (Table S8; Figure 2A): a single large ratio at *k* = 2, after which the ratios fall toward one, fixed *K* = 2. Vertices were then ordered by their value on *v*_2_. Every split of the ordering into two groups, all vertices below a threshold vs. all above, was scored by the normalized cut. A margin of 200 vertices at either end was excluded (5% of the giant component for the PAN graphs, which are smaller), to avoid trivial splits that detach only a small fragment: the normalized cut penalizes small sides, but weakly, and in some graphs a weakly attached fragment otherwise gave the lowest cut. The split with the lowest normalized cut was taken as the initial partition and then refined (below).

The split was then refined by single-vertex descent. In each round, the vertices with more edge weight to the other block than to their own were ranked by this difference and moved in that order, in consecutive batches of about 100 (the ranked list divided into ⌊*m* / 100⌋ batches, *m* being the number of ranked vertices in the round). A batch was kept if it lowered the normalized cut. At the first batch that did not, its vertices were instead tried one at a time, each kept only if it lowered the normalized cut, and the round ended. Rounds were repeated until one moved no vertex, with a limit of 60 rounds. Finally, any piece of a block that was disconnected from the rest of that block in the window graph was assigned to the other block. As checks, the descent was repeated moving at each step only the single vertex that lowered the cut most, and the partition was recomputed after removing the temporal pole, anterior ventral temporal cortex and orbitofrontal cortex from the graph (Supplementary Results S2). Vertices outside the giant component remained unassigned (Results, “Spectral decomposition of cortical connectivity at 3–20 mm yields two blocks”; Figure 2B; Supplementary Results S2).

A split was reported as a partition only if its smaller side contained at least 5% of the assigned vertices (those of the giant component excluding the peri-allocortex). In the full-range graph (433 L and 425 R vertices) and the long-range graph at d ≥ 20 mm (right hemisphere), the lowest-cut split only detached a smaller fragment. We report these hemispheres as a single block, without an eigengap, and give the adjusted Rand index (ARI; defined below) of the lowest-cut split. Block 1 (lattice cortex) is the block that contains V1, block 2 (paralattice cortex) the other. In replications and controls the blocks were matched to the 7T blocks by overlap before the percentage of vertices in the same block was computed; the ARI does not depend on the labels.

To test whether the reported border is the only deep minimum of the normalized cut, we repeated the sweep and descent from 121 different starting splits, spaced evenly along the Fiedler ordering between the two margins. Each start was refined by descent until the cut no longer decreased, and we recorded the final cut and its agreement (ARI) with the reported 7T partition (orange in Figure 2C). Starts were counted as reaching the same border when the cut they reached was within 5% of the lowest one found; the depth of the cut is the ratio of the median cut reached over all starts to the lowest, 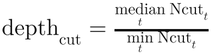 with *Ncut*_*t*_ the normalized cut reached from start *t*. The same descent from the same starts was run on the control graphs below (the geometric graph, which is the reference drawn in Figure 2C, the degree-preserving rewired graph and the graph with partner counts reassigned at random), and the same statistics are quoted for them. The cut profile is the normalized cut as a function of split position along the Fiedler ordering, computed for every split between the two margins before any descent. It was computed identically for the empirical graph and for every control graph. Its flatness is the fraction of these splits whose cut is at most 1.2 times the lowest cut of the profile, that is, splits nearly as good as the best one. For the left-hemisphere variant reached from some starts (Supplementary Results S2), the strip is the set of vertices that the two partitions assign differently; we report the share of its homotopic vertices in the posterior-central block of the right hemisphere, the homotopic ARI of each left partition with the right one, the block of the strip in each bootstrap replicate, and the share of the strip assigned to the anterior block in the other cohorts.

Partition agreement was measured by the adjusted Rand index (ARI; 1 for identical partitions, 0 for chance agreement) over the vertices assigned in both partitions, and by the percentage of vertices in the same block after matching the block labels between the two partitions (each block paired with the block it overlaps most). The agreement of the two hemispheres was measured at homotopic positions, taking as homotopic the vertices with the same index of the 32,492-vertex fs_LR 32k mesh in the two hemispheres (29,653 pairs). Percentages of a hemisphere are given relative to its cortical vertices (29,696 in the left hemisphere and 29,716 in the right). The counts differ because the HCP medial-wall masks of the two hemispheres differ.

#### Null models and controls for the partition

To establish whether the partition reflects the organization of short-range connectivity rather than a simpler property of the data, it was compared with a set of control graphs and alternative divisions of the cortex, each isolating one candidate explanation: the geometry of the cortical surface, the number of partners per vertex, connection density, the strength or length of the edges and the spatial smoothness of the graph’s own patterns. A final analysis assessed its stability across resampled cohorts (Supplementary Results S2 and S3; Table S6). Unless stated otherwise, each control was partitioned by the same procedure as the empirical graph and evaluated by the lowest normalized cut it reached, by the shape of its cut profile and by its agreement with the partition of record. Controls that define only which pairs are connected were given the empirical weights at random (see Nearest-partner graphs) and compared with the empirical graph, with the area factor applied as for every graph (Area weighting).

##### Nearest-partner graphs

Two control graphs keep the geometry of the cortical surface and the empirical number of partners of each vertex: (1) in the own-degree graph, each vertex is connected to as many of its nearest neighbors (by cohort geodesic distance, ties broken by pair index; a pair is an edge if either vertex selects the other) as it has partners in the empirical graph; (2) in the degree-scrambled graph, these partner counts are shuffled at random across the vertices of the hemisphere before the nearest neighbors are chosen. The shuffle was performed over all vertices of the hemisphere, the peri-allocortex included; because the peri-allocortex has no partners in the window graph, about 1,600 vertices outside it received a count of zero. Restricting the shuffle to the vertices outside the peri-allocortex gave the same result. Each edge of these graphs and of the rewired graphs below was given a weight drawn at random from the weights of the empirical window graph, so that the controls differ from the empirical graph only in which pairs are connected.

##### Geometric graph

Pairs were the window candidates with *d* ≤ *d* *, where *d* * (15.15 mm L, 15.25 mm R) makes the number of pairs equal to the number of edges of the functional window graph. Weights were 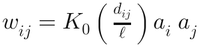, with *K*_0_ the modified Bessel function of the second kind of order zero (the steady-state Green’s function of diffusion with leakage on an infinite plane) and *l* the short-range length constant *l*_1_ of the edge-probability profile. The graph was divided by the procedure used for the partition and restarted from 121 splits along its Fiedler ordering (Figure 2C). In a wider search, the descent was started from 366 splits: the partition, 61 splits along each of the orderings by eigenvectors 2–6, and 60 regions grown by breadth-first search from random vertices to 10–50% of the hemisphere. Splits with a smaller side under 5% were excluded; a division was counted as unrelated to the border at ARI < 0.3. The same wider search was run on the empirical graph.

##### Degree-preserving rewiring

Each vertex was reconnected at random among its own neighbors, while keeping, on average, its empirical number of partners. We used the maximum-entropy model, the most random graph with these expected degrees: each window candidate ( *i, j* ) was included independently with probability 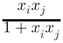, with the vertex parameters *x*_*i*_ fitted so that each vertex’s expected number of partners matched its empirical count (Table S8). Three random draws were made. In a distance-matched variant, each vertex also kept its distribution of partner distances: the random graph was drawn separately within each 1-mm band of cohort geodesic distance, so that each vertex’s expected number of partners in every band equaled its empirical number. One random graph was drawn per hemisphere. For each rewired graph we also computed the crossing ratio, as for the empirical graph: the number of edges crossing the reported border divided by the number expected if crossing pairs were connected at the within-block rate for pairs of the same length,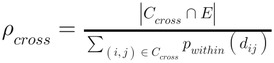(symbols as under *Crossing pairs*, below).

##### Dense foci

To test whether the partition simply separated the most densely connected regions from the rest, we took the top 5, 10, 20, 30, 40 and 50% of vertices by window degree (Pairs, neighbors and partners) as one block and the remaining vertices as the other, and computed the normalized cut of each split on the empirical window graph.

##### Weakest edges

The window graph was rebuilt from the weakest window candidates at the same edge count. Weights for them were initially defined as 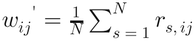 over all subjects (*N*=151) and then shifted to 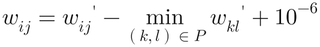, so that all were positive. Pairs were selected on the cohort mean r without area weights; the area factor was applied to the selected edges.

##### All pairs

Every window candidate was made an edge, without the floor and the support rule, with weight *w* = (max mean ( r ), 10^− 6^ ).

##### Strongest edges at any distance

To test whether the partition depended on restricting edges to short distances, we built a graph from the strongest edges of the full-range cohort graph, excluding the peri-allocortex, regardless of their length. We kept as many edges as the window graph contained, choosing those with the highest weight *w* = *mean* ( *r* ) (39% L and 38% R of the edges were 40 mm or longer).

##### Random edges

To test whether the partition depended on which pairs are strongly correlated, we built a graph with the same number of pairs as the empirical window graph drawn at random from all window candidates (Pairs, neighbors and partners), regardless of correlation strength. The empirical window weights were assigned to these edges at random (three random draws).

##### Smoothness-matched surrogates

To test whether the reported border was a better cut than any equally smooth division of the cortex, we generated 1,000 random maps with the same spatial smoothness as the graph’s own low-frequency patterns. Each map was a random combination of the 59 lowest-frequency eigenvectors *u*_*k*_ of the window graph’s symmetric normalized Laplacian, excluding the trivial first one (the first 60 eigenpairs were thus computed for this purpose): ∑_*k*_ *c*_*k*_ *u*_*k*_, with 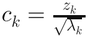(*λ*_*k*_ floored at 10^−3^*λ* ) and *z* drawn from a standard normal distribution, on the vertices of positive degree. This set leaves out the vertices with no window edge and can include small components outside the giant component. Weighting by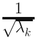 gives the smoother patterns larger amplitudes, as in a random field on the graph. Each surrogate map was multiplied by *D*^−1 / 2^ and swept exactly as the Fiedler vector *v*_2_ (Spectral partitioning), with the same 200-vertex margins and without descent. Along each sweep we recorded the lowest normalized cut, and we report the mean and the best surrogate value against the empirical one. The surrogates are built from the graph’s own eigenvectors, among which the Fiedler vector has the largest weight, so the comparison is descriptive, and no p value is reported.

##### Long-range and full-range graphs

To test whether long-range connections alone yielded the partition, the edges of the full-range cohort graph at *d* ≥ 20, 40 or 60 mm (at 60 mm, the long-range edges; see Short- and long-range bounds), or at all distances (same floors, support and weights; edges into the peri-allocortex kept, its vertices removed before partitioning), were partitioned using the same procedure.

##### Long-range graph against the canonical networks

For cutoffs from 20 to 70 mm in 5-mm steps and at 80 and 90 mm, the three leading non-trivial eigenvectors of the giant component 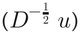 were correlated vertex by vertex (absolute Pearson *r*) with the first three functional gradients of Margulies et al.^55^ (neuromaps), and the graph was divided at pre-specified *K* (*K* = 2 by a median split of the Fiedler vector and *K* = 7 by k-means on the six leading non-trivial eigenvectors, each row scaled to unit length). Clusters were matched one to one to the seven networks of Yeo et al.^56^ by the Hungarian algorithm on the Dice matrix (the Dice coefficient of a cluster and a network is twice their overlap divided by the sum of their sizes, *Dice* 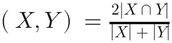: 1 for identical sets and 0 for disjoint ones), and scored by ARI against the Yeo parcellation and by the number of networks recovered at Dice ≥ 0.4. *η*^2^ is the share of the variance of the principal gradient (gradient 1 of Margulies et al., as above) accounted for by the main partition, 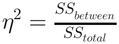 (the sum of squares of the gradient values explained by the block means over their total sum of squares), computed in each hemisphere over the lattice and paralattice vertices of the giant component of the *d* ≥ 20 mm graph (Supplementary Results S3; Figure S3 shows the left hemisphere).

##### Bootstrap over subjects

To test whether the original partition depended on the particular subjects in the cohort, the cohort graph was rebuilt from resampled cohorts. In each of 60 replicates, 151 subjects were drawn at random with replacement. For each of the 12,330,406 window candidates, the support was recomputed as the number of subjects above their floor, counting each subject as many times as it was drawn (each subject keeping its own floor *τ*_*s*_ ), and the weight as the mean correlation over the 151 draws. A pair was an edge when its support reached the majority of 76 and weight was defined as in *Cohort graph*. Each resulting graph was partitioned exactly as in the main analysis (spectral decomposition, sweep and descent), separately for each hemisphere, and compared with the main partition. Run with every subject drawn exactly once, the same code reproduced the main partition exactly (ARI 1.0).

#### Hyperparameter and robustness sweeps

To test the robustness of the results to methodological choices, every arbitrary parameter chosen for the main construction was varied one at a time, the rest of the analysis being re-run unchanged, and the resulting partition scored against the main partition (Table S6; Supplementary Results S2):

##### Window

The outer bound of the window was set at 12, 15, 18 and 20 mm, and the inner bound at 3, 5, 7, 8 and 10 mm. Each subject’s floor was kept at its value calibrated on the 3–20 mm window, so that only the range of distances changed.

##### Null floor

The per-subject floors were replaced by a single threshold on the cohort mean *r* chosen to keep the same number of edges (*r* = 0.126). Pairs were selected on the cohort mean r without the area weights, and the area factor was applied to the selected edges. In a second variant, each subject’s floor was replaced by a stricter per-subject threshold derived from the subject’s own distribution of *r* over all within-hemisphere pairs at *d* ≥ 3 mm outside the peri-allocortex: a Gaussian null was fitted to the central peak of this distribution (Efron’s central matching^126^), and the threshold was the lowest positive *r* at which the local false-discovery rate, 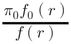, fell to 0.1 (median 0.149, range 0.109–0.249; Table S8).

##### Edge support

The majority rule (a vertex pair being accepted as an edge in the cohort graph only if *r* > *τ*_*s*_ in at least 76 of the 151 subjects) was replaced by a requirement for one third, two thirds or 90% of the cohort (51, 101 and 136 of the 151 subjects).

##### Edge weight

The definition of *w* = *mean* ( *r* ) over all subjects (see above) was replaced (1) by the mean of the margin *r* − *τ*_*s*_ over all subjects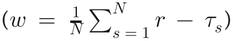, (2) by the mean of the margin over the supporting subjects only 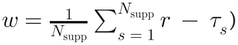, and (3) by the mean of the rectified margin 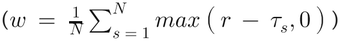.

##### Cut criterion

The sweep was scored by conductance,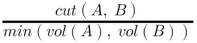, instead of the normalized cut.

##### Split halves

The 151 subjects were divided into disjoint halves of 76 and 75. Subjects were dealt in turn, in the order of the cohort list (Data S1), into six groups, and groups 1–3 formed one half and groups 4–6 the other. Each half was taken through the whole analysis pipeline independently (floors, majority support within the half, i.e., respectively 39 and 38 subjects, spectral partitioning and refinement), and the result was compared with the other half and with the full cohort.

##### Sex

The cohort was also divided by sex as recorded in the HCP data (89 women, 62 men), and each group was carried through the entire chain on its own, like a half, with its own majority rule (45 and 32 subjects).

##### Global-signal regression (GSR)

The mean time course of the cleaned, z-scored series over the 59,412 cortical vertices was regressed from every vertex of each subject’s concatenated time series (least squares, no intercept). The series were then restandardized (z-scored), and the analysis pipeline was repeated unchanged from the floors onward.

##### Peri-allocortex

The analysis was repeated without the peri-allo-cortex mask, with each subject’s floor recalibrated to include periallocortex pairs in the window (Table S6). To describe the third eigenmode of this unmasked graph (Discussion), each vertex was given the coordinates of the second and third eigenvectors of *L*_*sym*_ (the two leading non-trivial ones), scaled to unit length, and the vertices were divided into three clusters by k-means (50 initializations), as for the sectors (Sectors).

#### Controls for non-neural spread

##### Lower envelope

For each subject and each 1-mm bin of cohort-median geodesic distance, we took the 5^th^ percentile of the raw correlation *r* over all window candidates of both hemispheres ( 3 ≤ *d* < 20 mm, peri-allocortex excluded; linear interpolation) as the lower envelope of the correlation profile. A spatially uniform, non-neural process would add the same correlation to every pair at a given distance. The weakest correlations at that distance therefore set an upper limit on its contribution, and the lower envelope estimates this limit while being less sensitive to outliers than the minimum. For each subject, the lower envelope was fitted with a single exponential, 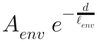, by nonlinear least squares on the untransformed 5^th^-percentile values, with each 1-mm bin represented by its center. Only bins below 20 mm with a positive envelope were used; the envelope reached zero by about 10 mm. We report *ℓ*_*env*_ as the mean ± SD over the 151 subjects. The upper bound on the non-neural share of the correlation at each distance is the ratio of the cohort-mean envelope to the cohort-mean median correlation at that distance, 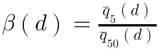, with 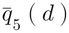 the cohort-mean lower envelope (5^th^ percentile) and 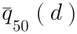 the cohort-mean median correlation in the bin of *d*. Where the envelope was measurable, the ratio used the measured values. Beyond that range, the envelope was replaced by the fitted curve, with *ℓ*_*env*_ fixed at its cohort mean and *A*_*env*_ refitted to the cohort-mean envelope (*A*_*env*_ = 0.864), and the median correlation was interpolated between bin centers (Table S8). Values quoted at 3, 5 and 7 mm are those of the 3–4, 5–6 and 7–8 mm bins (Supplementary Results S1).

##### Spatial smoothness

The HCP pipeline smooths the grayordinate series on the cortical surface with a Gaussian kernel of 2 mm FWHM (Data preprocessing). Our own steps (censoring, nuisance regression, z-scoring) act on each vertex over time and do not mix vertices. We measured two quantities (Supplementary Results S1; Table S8).

1. Effective smoothness. For each of the 151 subjects, the spatial smoothness of the series was estimated on the subject’s own mid-thickness surfaces (MSMAll, 32k fs_LR mesh) with wb_command - cifti-estimate-fwhm (Connectome Workbench 2.1.0), with each hemisphere as one component and one estimate for the whole series (option -whole-file). The estimate was made on the released series and on the series as analyzed, which were first written back into a CIFTI file on the same mesh. This estimate is the total smoothness of the data, including the applied kernel, the spatial correlation of the BOLD signal itself, and the mapping from volume to surface.
2. Correlation created by the kernel alone. White noise, independent across vertices (1,200 frames), was smoothed with the same kernel (wb_command -metric-smoothing, 2 mm FWHM) on the mid-thickness surfaces of three subjects (the first, middle and last of the cohort list), both hemispheres. The Pearson correlation between the smoothed series of two cortical vertices was then averaged over all pairs in 1-mm bins of cohort geodesic distance, and over the six runs.

##### Facing sulcal banks

For every edge of the window graph, the Euclidean distance between its two vertices on the template mid-thickness surface (HCP S1200, MSMAll) was compared with the template geodesic, the shortest path along the edges of that mesh within the cortex; the edges of the two hemispheres were pooled. A pair was classed as folded, joining the facing banks of a sulcus, when its geodesic distance was at least twice its Euclidean distance, and as flat, lying along the sheet, when the ratio was at most 1.15; pairs with intermediate ratios entered neither class.

Within each 2-mm bin of Euclidean distance (4–6, 6–8 and 8–10 mm), the mean weight of the folded pairs was compared with two controls taken from the flat pairs pool: (1) pairs at the same Euclidean distance (Euclidean-matched: the same straight-line distance, but a shorter path along the sheet), and (2) pairs at any Euclidean distance weighted to the folded pairs’ distribution of geodesic distance in 1-mm bins (geodesic-matched: the same path along the sheet, but a larger straight-line distance; for Cohen’s *d*, a sample of pool pairs was matched bin by bin). Each contrast was summarized by the difference of means and by Cohen’s *d* (Effect sizes). The comparison was repeated with the pairs restricted to those with both ends in the same HCP-MMP1.0 area, and by sweeping one distance from 6 to 20 mm while the other was held at 18–20 mm (geodesic) or 6–8 mm (Euclidean) (Supplementary Results S1).

##### Band decomposition of the weight map

The short-range density *δ*_*i*_ of each vertex (Short-range density, above) was recomputed from the edges and neighbors of one distance band at a time (3–5, 5–10, 10–15 and 15–20 mm), 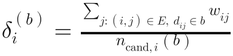, with *n*_*cand, i*_ ( *b* ) the number of neighbors of vertex *i* in that band. The ratio of the block means (lattice over paralattice, both hemispheres pooled) was reported for each band 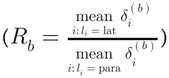 whether the contrast between the blocks depended on the shortest edges, where a uniform non-neural component could contribute, we compared the map built from all window edges (3–20 mm) with the map built from 10–20 mm edges only, beyond the distance at which the lower envelope reached zero (Supplementary Results S1).

##### Within-area profiles

For each subject, on the unthresholded correlation matrix, we computed the mean *r* of the window candidates with both vertices inside a single left-hemisphere area *A* (V1, V2 and 3b in lattice cortex; 46, 9a and 10d in paralattice cortex) in each 1-mm bin *b* of cohort geodesic distance, using only bins with at least 20 pairs, and averaged over subjects, 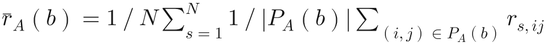, with *P*_*A*_(*b*) the window candidates inside *A* in bin *b, r*_*s,ij*_ the correlation of pair (*i, j*) in subject *s* and *N* the number of subjects. The values in the 5–6 mm and 19–20 mm bins were then averaged over the three areas of each block (Supplementary Results S1).

##### Venous exposure

Exposure to pial veins was estimated from the VENAT atlas^117^, the vessel partial volume from 7T quantitative susceptibility maps of 20 adults (MNI152, 0.5 mm). For 40 subjects of the 7T cohort (Table S8), the atlas was sampled by trilinear interpolation at each vertex of the subject’s 32k pial surface in MNI space and at 0.5-mm steps up to 3 mm outward along the white-to-pial direction; the exposure of a vertex is the maximum over these points, averaged over subjects. Exposure was compared between the blocks and related, within each block, to the short-range density and to the unweighted density (see Short-range density; Spearman’s ρ). The block ratio of short-range density was computed within quintiles of exposure, and the band ratios (Band decomposition of the weight map) were recomputed without the 10, 20 or 30% most exposed vertices of each hemisphere and their edges. The same vertices were removed from the window graph, and the descent (Spectral partitioning) was started from the reported border restricted to the remaining giant component. A vein would elongate the set of partners of a vertex along its course. Elongation was measured with the orientation tensor of the unit tangent vectors on the fs_LR sphere from a vertex to its window partners, as the axial anisotropy (*λ*_1_ − *λ*_2_)/(*λ*_1_ + *λ*_2_), with *λ*_1_ ≥ *λ*_2_ the two nonzero eigenvalues; it was compared with the same number of partners drawn in each 1-mm band from the vertex’s own neighbors (ten draws) and reported as the median excess over this null in the interior of each block (vertices with no neighbor in the other block; Supplementary Results S1).

##### Homotopic flatness

If homotopic correlations arose from signal spreading across the midline, they would be strongest near the midline and fall off steeply with distance from it. To test this, we regressed the homotopic weight of each vertex (Homotopic edges, below) on its distance from the midsagittal plane, measured as the absolute *x* coordinate on the template mid-thickness surface, *h*_*i*_ = *α* + *η*|*x*_*i*_| + *ε*_*i*_, with *h*_*i*_ the homotopic weight of vertex *i*. The regression used ordinary least squares over the left-hemisphere vertices with a supported homotopic edge (21,800 vertices). We compared the fitted slope and its 95% confidence interval with the decay that a spread with the lower envelope length constant ( *ℓ*_*env*_ ≈ 1.95 mm) would produce over the same range of distances (Supplementary Results S1).

#### The border between the blocks

##### Crossing pairs

The window candidates (3 ≤ *d*_*ij*_ < 20 mm) were classed by the block labels of record as within lattice cortex, within paralattice cortex or crossing the border. Denoting by *l*_*i*_ ∈ { *lat, para* } the block label of vertex *i* and by *d*_*ij*_ the cohort geodesic distance between vertices *i* and *j*, the three classes are (*C*_*lat*_ = { (*i, j*) : *l*_*i*_ = *l*_*j*_ = *lat*}, (*C*_*para*_ = { (*i, j*) : *l*_*i*_ = *l*_*j*_ = para } and (*C*_*cross*_ = { (*i, j*) : *l*_*i*_ ≠ *l*_*j*_ } For each class *c*, the edge probability (the fraction of its pairs that are edges of the cohort graph) was computed overall and in 1-mm bins of cohort geodesic distance, by count (without the area weights that enter *p* ( *d* ) under Edge-survival probability model): 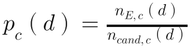, where *n*_*cand, c*_ ( *d* ) = | {( *i, j* ) ∈ *C*_C_ : ⌊*d*_*ij*_⌋ = ⌊*d*⌋ }|is the number of pairs of class *c* in the 1-mm bin containing *d* and *n*_*E*_, *C*_*c*_ ( *d* ) = | { ( *i, j* ) ∈ _C_ ∩ _*E*_: *⌊d*_*ij*_ *⌋* = *⌊d ⌋*}|the number of those that are edges of the cohort graph; the overall probability *p* is the same ratio with numerator and denominator summed over all bins. Crossing edge probability was compared with a distance-matched expectation: each crossing pair was assigned the within-block edge probability of its distance bin, with the two blocks pooled, 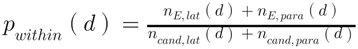, and these were averaged over the crossing pairs, 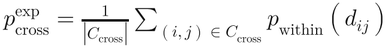 the fraction of crossing pairs expected to be edges if crossing pairs were connected as often as within-block pairs of the same length. We report the ratio of the observed fraction *p* _*cross*_ to this expected fraction, 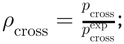 ratio below 1 means that pairs crossing the border are connected less often than distance alone predicts. The same comparison was made without any threshold, on the cohort-mean correlation of every window candidate (the mean of r over the N subjects, as for the weight w): each crossing pair was assigned the mean correlation of the within-block pairs of its 1-mm distance bin, the two blocks pooled, and the mean correlation of the crossing pairs was divided by the mean of these expected values (Results; Supplementary Results S3). The distance of a vertex from the border is its cohort geodesic distance from the nearest border vertex, *b*_*i*_ (defined under *Distance to the border*). Crossing pairs were also compared with the within-block pairs of each block separately, matched on two variables: geodesic distance (1-mm bins) and the distance of the farther endpoint from the border (0–2, 2–5, 5–10, 10–15 and 15–20 mm), the farther endpoint of the pair ( *i, j* ) lying at max (*b*_*i*_, *b*_*j*_) . Each combination of a distance bin and a border-distance bin defines a cell, *m*_*ij*_ = (*⌊d*_*ij*_ *⌋, γ* max (*b*_*i*_, *b*_*j*_ ))), with *γ* the index of the border-distance bin of the farther endpoint. For each block *l* ∈ { *lat, para* }, the expected rate for crossing pairs is the average of the within-block edge rates over cells, each cell weighted by the fraction of crossing pairs that fall in it: 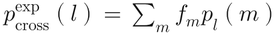, where *m* runs over the cells, 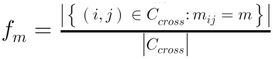 is the fraction of crossing pairs in cell *m* and 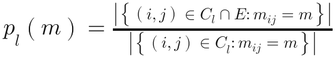 the edge probability of the within-block pairs of block *l* in that cell, so that the within-block pairs are given the same distribution of lengths and border distances as the crossing pairs. The support of each crossing pair (the number of subjects in which it exceeds the floor) and the mean weight of the crossing edges were taken from the same computation as for all other edges. For each vertex, the fraction of its window partners that lie in its own block was also computed 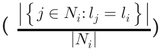.

##### Distance to the border

Border vertices, denoted ∂, are the vertices that lie within 2.2 mm (cohort geodesic, about one mesh spacing) of a vertex in the other block of the main partition, excluding the peri-allocortex and unassigned vertices. Each vertex *i* was assigned its cohort geodesic distance to the nearest border vertex, 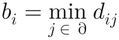, the minimum being taken over all border vertices, on either side of the border. For each vertex with at least 20 neighbors at 7–20 mm (starting at 7 mm, beyond the range in which almost every pair is an edge), we computed two quantities: the number of these neighbors, *n*_cand, *i*_ [ 7, 20 ), and the fraction of them that are edges,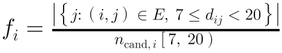 . Both were summarized by their medians in four groups of vertices: (1) border vertices, within 2.2 mm of the other block; (2) the interior of lattice cortex and (3) the interior of paralattice cortex, more than 20 mm from the border; and (4) all vertices within 20 mm of the border. The edge fraction was also summarized for each HCP-MMP1.0 area, as the median over the area’s vertices, together with the median distance of those vertices from the border (Supplementary Results S3, Location of the border). For the profile across the border (Supplementary Results S3, Connectivity density and pattern), the cohort-mean correlation and the edge fraction were averaged over all the window candidates of each vertex, and then over the vertices within 3 mm of the border and over those 10–20 mm from it on either side.

##### Temporal signal-to-noise

For each of the 151 subjects and each of the four released resting-state runs (Table S8), the temporal signal-to-noise ratio of every cortical vertex was computed as the temporal mean divided by the temporal standard deviation of the cleaned series, 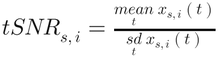, with *x*_*s, i*_ ( *t* ) the series of vertex *i* in subject *s*. The values were averaged over the runs. The median over subjects, 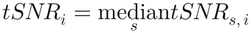, was summarized by the median over the same groups of vertices (Supplementary Results S3, Location of the border).

#### Within-block connectivity characterization

The two blocks were compared on the full-range cohort graph, which includes edges at all distances. Four aspects were assessed: the geometry of each vertex’s strongest partners, the falloff of connectivity with distance within large single areas, the strength of the homotopic edge to the opposite hemisphere, and the internal integration of each block (Figure 3). As in the window graph, pairs closer than 3 mm were left out of the partner-distance profiles, the near/far ratio and the within-area profiles, to limit the contribution of the hemodynamic point-spread function. Block contrasts of per-vertex measures are summarized by medians and Cohen’s *d* and tested by the spin test (Quantification and statistical analysis).

##### Partners and patches

The strongest partner set of a vertex, *Ω*_*i*_, is its 450 strongest same-hemisphere partners, ranked by the weight *w* of the edge joining them to it in the full-range cohort graph (all of its partners if it has fewer; ties in *w* broken arbitrarily), at all distances, pairs closer than 3 mm included. Its size, 450, was matched to the size of a vertex’s short-range neighborhood (on average 442 L and 441 R vertices lie within 3–20 mm), so that a vertex whose strongest connections were all short-range could fill its strongest partner set within that neighborhood. Patches are the groups obtained by single-linkage clustering of the partners at 10 mm on the cohort geodesic: two partners belong to the same patch if they are connected by a chain of partners, each less than 10 mm from the next. The linkage distance, 10 mm, is the distance at which the lower envelope of the correlation reached zero (Lower envelope), so that two groups of partners closer than this could be joined by non-neural spread alone. The number of partners (100 to 900; Table S8) and the linkage distance (5 mm) were varied on 8,000 vertices per hemisphere drawn at random from all vertices outside the peri-allocortex with a block label (Supplementary Results S4).

##### Illustrative seeds

Partners and patches were defined as for the cohort (Partners and patches, above), but from one single subject’s own correlations. We chose one subject (550439, whose floor, τ_*s*_ = 0.103, lay within the interquartile range of the cohort) and six illustrative seed vertices on the left hemisphere, three in lattice cortex (V1, V2, area 3b) and three in paralattice cortex (areas 46, 9a and 10d). Each seed was the vertex of its area with the smallest summed cohort geodesic distance to the other vertices of the same HCP-MMP1.0 area, except the V2 seed, a vertex of the dorsolateral occipital surface chosen manually because nearly all of its partners fall in one patch in this subject, as do those of the other two lattice seeds (vertex indices in Table S8). For each seed, the subject’s 450 strongest same-hemisphere pairs by *r* were taken and kept when *r* > τ_*s*_. Patches are colored, the seed is black (Figure 3A).

##### Partner geometry

For every cortical vertex outside the peri-allo-cortex, three measures were derived from its strongest partner set and the cohort geodesic: (1) Median partner distance: the median cohort geodesic distance from the vertex to its partners, 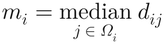 . Partners closer than 3 mm were included (see

Supplementary Results S4 for medians without them); (2) Number of patches: the number of patches of the strongest partner set (Partners and patches, above); (3) Largest-patch share: the fraction of partners in the largest patch, 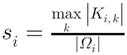, with *K*_*i, k*_ the *k*th patch of the vertex. Cohen’s *d* was computed on the base-10 logarithms of partner distance, which is right-skewed, and on raw values for the patch count and the largest-patch share.

##### Profiles of partner distance

For each vertex the fraction of its 450 partners in a 3–5 mm band and in each 5-mm band of cohort geodesic distance from 5 to 90 mm was computed, 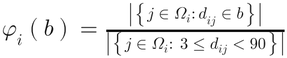, partners closer than 3 mm or farther than 90 mm being excluded from the profile and its denominator. The per-vertex fractions were then averaged within each block of the main partition and, for display, normalized to each block’s value in the 5–10 mm band (Figure 3D1). The ratio quoted in the Results is that of the paralattice block means in the 85–90 and 25–30 mm bands. The minimum quoted in the Results (20–30 mm) is the pair of adjacent bands with the lowest summed fraction, reported together as the 20–25 and 25–30 mm bands have nearly equal values.

##### Near/far ratio

For each vertex *i*, partners were counted in a near band, *b*_*near*_ = [ 3, 20 ) mm, and a far band, *b*_*far*_ = [ 60, ∞ ) mm (close to the crossover of the two components of the distance profile; see *Edge-survival probability model*). In each band the count was turned into a connection probability, the number of the vertex’s 450 strongest partners (Partners and patches) in the band over the number of cortical vertices of the hemisphere, periallocortex included, in the band, with a Haldane–Anscombe correction of 0.5 added to the count of partners so that the logarithm of the ratio stays finite for vertices with no partners in one of the two bands:

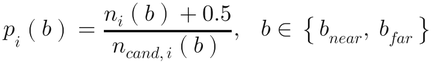

where *n*_*i*_ (*b)* = |{*j* ∈ *Ω* : *d* ∈ *b* }| is the number of the vertex’s partners in band *b* and *n*_*cand, i*_ ( *b* ) = | *j*: *d*_*ij*_ ∈ *b* | the number of cortical vertices in that band (peri-allocortex included). Figure 3D2 maps the base-10 logarithm of this ratio, centered at the Youden cut, 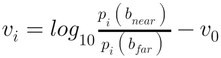 . The Youden cut is the threshold on the log ratio that best separates the two blocks: it maximizes the fraction of block 1 vertices above the threshold plus the fraction of block 2 vertices below it, searched over the 1^st^ to 99^th^ percentiles of the values from both blocks pooled (1.82 L and 1.81 R). The color scale saturates 1.5 log units on either side of the center. We also report, for each block, the fraction of vertices with no partner beyond 60 mm.

##### Within-area falloff

We selected two contiguous territories, V1–V2 in the posterior-central block and dorsolateral prefrontal cortex (areas 46, 9a, 10d) in the anterior block. For each territory, we plotted the weight *w* of every pair with both vertices inside it (pairs between its areas included) against its cohort geodesic distance, from 3 to 50 mm in 1-mm bins, using the full-range cohort graph. Pairs not supported by a majority of subjects were included with a weight of zero. The falloff over 3–50 mm was summarized by the curvature index κ, defined as twice the quadratic coefficient of a quadratic fit on the bin means, with both axes scaled to the unit interval, 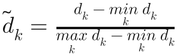 and 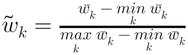, with *d*_*k*_ the center of bin_*k*_ (3.5 to 49.5 mm) and 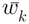 its mean weight, the quadratic being 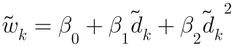, fitted by unweighted least squares over the bins, so that κ = 2β2 is the second derivative of the fitted curve in the scaled coordinates (κ = 0 for a straight-line decline, κ > 0 for a convex curve, one that falls steeply at short distances and then flattens). κ was computed in each hemisphere; Figure 3C shows the left.

##### Homotopic edges

For each vertex, the homotopic edge joins it to the vertex with the same mesh index in the other hemisphere (which the fs_LR 32k template aligns to the homotopic position), *i*’ (43 L and 63 R vertices have no homotopic counterpart because their counterpart lies in the medial wall). The weight *w* and support of each homotopic edge were taken from the same computation as for every other edge, and the edge counted as supported when it met the cohort’s majority rule. Unsupported homotopic edges were given a weight of zero, both here and in the block medians. The homotopic weight map was compared with each vertex’s short-range density (Short-range density; Figure 1E) by Spearman correlation, over all vertices outside the peri-allocortex with a homotopic counterpart. The fraction of supported homotopic edges was computed for each block and for the peri-allocortex (Figure 3F).

##### Internal integration

Two measures were used to assess integration of the blocks. (1) Coreness. The *k*-core of a graph is the largest subgraph in which every vertex has at least *k* partners within the subgraph. A vertex’s coreness is the highest *k* for which it belongs to the *k*-core, *c*_*i*_ = max { *k*: *i* ∈ *H*_*k*_ }, with *H*_*k*_ the *k*-core, so high coreness means a vertex lies in a densely interconnected core. Coreness was computed on the binary (unweighted) 3–20 mm window graph of each hemisphere. It was summarized for each block by its mean, its maximum and its survival curve (the fraction of vertices with coreness at least *k*, for every *k*). We also report the area under the ROC curve (AUC) for coreness as a measure separating the two blocks (Figure 3E). (2) Degree by distance band. For each vertex, we counted its edges in the full-range cohort graph in four bands of cohort geodesic distance: below 20 mm (including pairs closer than 3 mm), 20–50 mm, 50–100 mm and 100 mm or more. Each count was divided by the number of cortical vertices of the hemisphere in that band, and the fractions were averaged within each block (Figure S4B).

#### Within-lattice streams and hierarchy

##### Sectors

The 3–20 mm window graph was restricted to the edges with both ends in lattice cortex (2.66 L and 2.68 R million edges) and its symmetric normalized Laplacian decomposed as above. Its spectrum had no single gap (Supplementary Results S4), so the number of sectors within lattice cortex was set by reproducibility. We ran k-means clustering (50 initializations) for *K* = 2 to 7 on the *K* − 1 leading non-trivial eigenvectors of the lattice-cortex graph, with each vertex’s row scaled to unit length^25^. This was done on the full HCP 7T rest cohort and, separately, on each of the two halves of the cohort described above, each half with its own majority rule and mean *r*, always within the block identified as lattice cortex in the main partition. Each solution, refined by single-vertex descent as described below, was scored by three agreement values, all measured by ARI: (1) between the two hemispheres at homotopic positions, (2) between the two halves of the cohort, and (3) between each half and the full cohort. *K* = 4 was the only value at which all three agreements reached 0.85 or more (Table S8; Supplementary Results S4). As for the main partition, the k-means sectors were then refined by single-vertex descent of the *K*-way normalized cut, 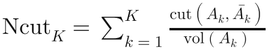, with *A* the vertices of sector *k* and Ā_*k*_ the rest of the lattice graph: in each round, every vertex with more edge weight to another sector than to its own was considered, in order of that excess, and moved to that sector if the move lowered Ncut_*K*_; rounds were repeated until no vertex moved. Sectors were named by the primary area they contain (visual, somatomotor, auditory-temporal) and, for the fourth, medial parietal, by its territory; each sector is the cluster that holds most of the vertices of V1, 3b, A1 or POS1, respectively, and at *K* = 4 each of these areas lay in a different cluster. To compare how cleanly the sectors and the main partition divide their graphs, we report the normalized cut divided by the number of blocks, <u>^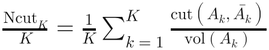^</u>, with *A* and Ā as above; this is the average fraction of a block’s edge weight that crosses to other blocks. Dividing by the number of blocks puts the four-sector division of the lattice and the two-way main partition of the hemisphere on the same scale, on which the sector cut is several times shallower (Supplementary Results S4).

##### Multimodal zone

The lattice partners of a lattice vertex are its partners in the full-range cohort graph that lie in lattice cortex, regardless of weight and distance. The multisector fraction is the fraction of them that lie in sectors (as partitioned at *K* = 4 with the procedure in *Sectors*) other than the vertex’s own, 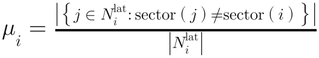, with 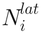 the set of lattice partners of vertex *i*. The multimodal zone was defined as the set of lattice vertices whose multisector fraction *μ*_*i*_ is at or above the 80^th^ percentile (Figure 4H).

##### Depth

Depth was computed separately for each sector and hemisphere, on the full-range cohort graph (all pairs at least 3 mm apart) restricted to edges with both ends in the sector, so that the order of the areas is not tied to the surface by the 20-mm bound of the window. For the visual, somatomotor and auditory-temporal sectors the vertex set was the largest connected component of the sector’s window graph. For the medial parietal sector it was medial parietal cortex proper (3,070 L and 3,120 R vertices). This is the cluster that holds POS1 when the lattice graph is divided by the procedure of Sectors at *K* = 6: k-means on the five leading non-trivial eigenvectors of the lattice graph, rows scaled to unit length, then the descent described under Sectors. This division separates it from the medial motor strip. The depth of a vertex is its distance through the graph from its sector’s base (listed below): the length *ζ*_*i*_ of the shortest path to it from any base vertex, computed with Dijkstra’s algorithm on the undirected weighted graph, with edge weights (lengths) 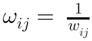 and 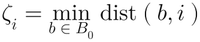, with *B*_0_ the base and dist ( *b, i* ) the shortest-path length from base vertex *b* to sector vertex *i*. Depths were then converted to percentiles within the sector’s vertex set, with tied values given their average rank. Low values therefore lie close to the base and 100 is the farthest vertex. Vertices that could not be reached from the base, if any, were given the highest rank. The base of the visual, somatomotor and auditory-temporal sectors was every vertex of the sector labeled V1, 3b or A1, respectively, in HCP-MMP1.0. For the medial parietal sector, which has no primary area, its border was chosen as base: the sector’s vertices sharing an edge of the fs_LR 32k mesh with a cortical vertex outside it, plus its vertices adjacent to the medial wall. Each side of the border taken alone (the side facing other cortex and the side facing the medial wall) and bases at the retrosplenial rim (RSC alone; RSC and ProS; RSC, ProS, POS1 and v23ab; RSC with the posterior cingulate areas) were compared with the border by the Spearman correlation, over the vertices of medial parietal cortex proper, of the resulting depth with per-vertex measures that vary with depth in the three sensory sectors: the number of patches, the near/far ratio, the number of long-range edges, the prefrontal share of long-range edges and the share of partners in other sectors (Supplementary Results S4). Depth from the border was also related to surface distance from it (the shortest path along the edges of the template mid-thickness mesh within the sector) and, with that distance partialled out, to the unweighted short-range density (Short-range density). Depth values of areas reported in the text are medians over the area’s vertices in the sector, for areas with at least 25 such vertices.

##### Comparison with known hierarchies

For each sensory stream, the areas were assigned levels from anatomical and physiological studies in primates (Table S8). For the areas with at least 25 vertices in the sector, the median depth was correlated with the level (Spearman ρ), and so was the median surface distance from the base, the geodesic distance along the mid-thickness surface from the nearest base vertex within the sector. Whether depth orders the areas beyond distance from the base was measured by the partial Spearman correlation of depth with level given surface distance. The partial correlation was compared with that of 100 rewired graphs, in which every edge kept its weight and one endpoint, chosen at random, and took as its other endpoint a vertex of the sector drawn at random in the same 1-mm bin of cohort geodesic distance from the kept endpoint (duplicate and self pairs dropped; Table S8); p is the fraction of the 101 values, the observed one included, that are at least as large as the observed one.

##### Convergence on medial parietal cortex

To describe the connections of the auditory-temporal sector with the rest of lattice cortex, we took the edges of the full-range cohort graph at *d* ≥ 20 mm that join a vertex of this sector (its largest connected component) to a vertex in another lattice sector. Each edge was classified by the sector of its far end: visual, somatomotor, medial parietal cortex proper, or the rest of the medial parietal sector. We report the share of edges in each class and, for the edges ending in medial parietal cortex proper, the HCP-MMP1.0 areas in which they end.

#### Lattice connectors

Two non-exclusive populations of lattice vertices, the *paralattice connectors* and the *peri-allocortex connectors*, were defined from the full-range cohort graph restricted to edges whose cohort geodesic distance was at least 60 mm (the long-range edges, see Short- and long-range bounds). Figure 4E shows, for each lattice vertex, the number of its long-range edges to any cortical vertex. Edges were classified by their other endpoint as lattice, paralattice or peri-allocortex. An edge to the peri-allocortex is a vertex pair with an end in the peri-allocortex whose correlation exceeds the subject’s floor (set on the window candidates) in a majority of subjects; similarly to other edges, its weight *w* is the cohort mean *r* . The same edges entered the long-range analyses (see *Cohort graph*, above); the full-range graph of Figure 3 excluded them.

##### Paralattice and peri-allocortex connectors

The two populations were identified by the same rule, applied to the long-range edges into paralattice cortex and into the peri-allocortex, respectively. Let *X* ∈ { paralattice, peri - allocortex } denote the target cortex. For each lattice vertex the number of its long-range edges into the target cortex was counted, 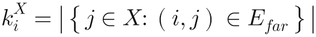, with *E* the long-range edges (cohort graph edges, see *Cohort graph*, with cohort geodesic distance *d* ≥ 60*mm*, see Short- and long-range bounds). The lattice vertices were ranked by the two 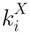 counts separately, and the connectors to a target were the top-ranked vertices that together send three quarters of all long-range edges from lattice cortex into that target, *T*_*X*_, the smallest set of top-ranked lattice vertices with 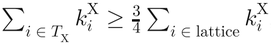, with any vertices tied at the lowest count included. Each connector population was compared with lattice cortex as a whole by its window degree (Pairs, neighbors and partners), summarized by the median over vertices. The length of its long-range edges into the target was summarized by their median cohort geodesic distance. Each connector was assigned to a region by the HCP-MMP1.0 area in which it lay: lateral temporal (TE1p, TE1m, TE1a, TE2p, TE2a, PHT, PH, STSvp, STSdp, STSva, STSda, TPOJ1, TPOJ2, TPOJ3, FST, MT, MST, STV, A5, TGd and TGv), inferior parietal and intraparietal (PF, PFt, PFop, PFm, PFcm, PGi, PGs, PGp, IP0, IP1, IP2, AIP, LIPv, LIPd, VIP, MIP, 7PL, 7PC and 7AL), medial parietal and posterior cingulate (7Am, 7Pm, 7m, 5mv, 5m, 31pv, 31pd, 31a, d23ab, v23ab, 23d, 23c, PCV, POS1, POS2, RSC, DVT and ProS), or other areas. The share of each region was counted for each population, and the connectors are shown by region in Figures 4F and 4G.

The long-range edges into non-lattice cortex of each population were classified by their endpoint outside lattice cortex as dorsolateral prefrontal (paralattice vertices of 46, 9-46d, a9-46v, p9-46v, 9a, 9p, 8Ad, 8Av, 8BL, 8C, IFSa, IFSp, i6-8 and s6-8), medial prefrontal (paralattice vertices of 10d, 10r, 10v, 9m, a24, p24, p32, d32, a32pr, 10pp and p24pr), ventrolateral and orbital prefrontal (paralattice vertices of 44, 45, 47l, 47m, 47s, IFJa, IFJp, a47r, p47r, 11l, 13l, OFC, pOFC, a10p, p10p, AVI, FOP4 and FOP5), other paralattice, parahippocampal (peri-allocortex vertices of PHA1, PHA2 and PHA3) and other peri-allocortex, and the share of each class was computed (Figure 4I). The input of the paralattice connectors to paralattice cortex was measured at the receiving end: for each paralattice vertex, the number of long-range edges it received from the paralattice connectors, 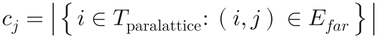, averaged over the paralattice vertices of each HCP-MMP1.0 area with at least 20 of them, 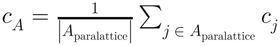, with *A*_paralattice_ the paralattice vertices of area A. Parieto-frontal channels: the paralattice connectors in PF, PFt, AIP and 7Am (anterior group) and in PGi, PGs, 7m and 31pv (posterior group) were compared by the share of their long-range edges into paralattice cortex that ends in 6r, 6a, 46 and IFSa, and in 8Ad, 8Av and 9m; the overlap of the two groups’ target distributions over paralattice areas is the sum over areas of the smaller of the two shares, 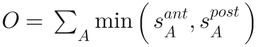 (0 for disjoint targets, 1 for identical ones; Table S8). The outline drawn in Figures 4F and 4G consists of the peri-allocortex connectors that share a mesh edge with a vertex outside the set, together with the vertices adjacent to them on the mesh inside the set; in Figure 4H it consists of the first of these rings only. Overlaps among the paralattice connectors, the peri-allocortex connectors and the multimodal zone were reported as percentages of each set, together with the share of all lattice-to-peri-allocortex long-range edges sent by each set and by their intersections (Supplementary Results S4, Connectors; Table S8).

##### Comparison with canonical networks

The paralattice and the peri-allocortex connectors were compared with the 7- and 17-network parcellations of Yeo et al.^56^ (fs_LR 32k label files, Table S8; the 17 labels were given the network names of Schaefer et al.^127^ in the order of the 17-network file and checked against their HCP-MMP1.0 composition). For each set, we computed the share of its vertices in each network, and its enrichment for that network: this share divided by the fraction of lattice cortex that the network occupies, 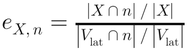 for set *X* and network *n* (1 = no enrichment). Both hemispheres were pooled (Supplementary Results S4, Connectors).

#### Thalamic connectivity

##### Preprocessing

For each of the 151 subjects of the 7T cohort we took the concatenated resting-state CIFTI series of the HCP release (*Data preprocessing*; Table S8). Every cortical vertex and every thalamic, striatal and cerebellar grayordinate was then cleaned in the same way: frames with FD > 0.25 mm were censored (interpolated for the fit and then removed, as in the cortical analysis), an intercept, a linear and a quadratic trend over the concatenated runs and the mean CSF signal were regressed out in one least-squares fit, and the series were z-scored over the retained frames. The mean white-matter signal, regressed in the cortical analysis, was not used here as its mask (*Data preprocessing*) includes the white matter next to the cortex and the internal capsule around the thalamus, and can therefore reflect cortical and thalamic signal. As a check, this model (trend, censoring and CSF) was compared with four others (trend only; trend and censoring; trend, white-matter and CSF signals; and the full cortical model, trend, censoring, white-matter and CSF signals), and was the one whose nucleus assignments agreed best with the established cortical targets of the specific nuclei (Table S3; Supplementary Results S5; Table S8). Thalamic grayordinates (2-mm grid, the same in every subject) were mapped, without further registration, to the nuclei of the MNI152 version of the 1-mm Morel atlas: each grayordinate took the nucleus most frequent among 27 points spaced 1 mm apart in its 2-mm cell (nearest atlas voxel at each point, background ignored), with the magnocellular and parvocellular divisions of the lateral geniculate nucleus merged and overlapping masks assigned to the smaller nucleus,^113–115^ and a nucleus was retained in a hemisphere when it held at least three grayordinates, giving 55 nucleus estimates over the two hemispheres (28 left, 27 right; Table S8). The lateral geniculate nucleus had no grayordinate in the CIFTI thalamus in any subject and was not analyzed.

##### Coupling measures

For each subject and nucleus estimate n, we computed partial correlations with the nucleus’s mean time course, in every case controlling for the mean signal of all other thalamic grayordinates of both hemispheres.

##### Cortical coupling

*r*_*n, v*_, with each vertex *v* of the ipsilateral cortex, averaged over the vertices of each block 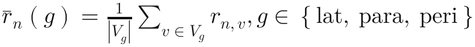 with vertex set *V*_*g*_ (*V*_lat_, *V*_para_ and *V*_peri_ as under Lattice connectors).

##### Striatal coupling

*r*_*n, str*_, with the mean time course of the ipsilateral striatum (all grayordinates of the caudate, putamen and accumbens).

##### Loop coupling

The mean of the striatal and cerebellar partial correlations, 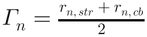.

##### Block preference

For each nucleus and each subject, we computed the difference between its mean partial correlation with two blocks, 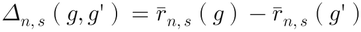, for three contrasts: paralattice minus lattice, peri-allocortex minus lattice and peri-allocortex minus paralattice (*g, g*’ ∈ { lat, para, peri } ). Each set of 151 paired differences was tested by the Wilcoxon signed-rank test and summarized by Cohen’s *d*_*z*_ (Quantification and statistical analysis). Nuclei were then assigned in two steps. (1) A nucleus was assigned to the peri-allocortex if both peri-allocortex contrasts were positive, each with *p* < 10^−3^ and *d*_*z*_ ≥ 0.3. (2) Otherwise, it was assigned to paralattice or lattice cortex if the paralattice-minus-lattice contrast had *p* < 10^−3^ and |*d*_*z*_| ≥ 0.3 in absolute value, according to the sign of *d*_*z*_. Nuclei meeting neither criterion were left unresolved (Figure 5B; Table S4).

##### Principal afferents

Each Morel nucleus was assigned a class by its principal afferents (first-order sensory relay, higher-order, basal-ganglia-recipient or intralaminar) and, where one exists, the subcortical output nucleus that projects to it (internal pallidum, substantia nigra pars reticulata or cerebellar nuclei), following the anatomical literature summarized in Table S3; the class and output nucleus of each nucleus are given in Table S4. Each class predicts the block its cortical targets fall in: nuclei receiving basal-ganglia or cerebellar output predict paralattice cortex, and first-order relays and the pulvinar predict lattice cortex. These predictions also served as the benchmark for the nuisance models (above; the nucleus estimates and their predicted blocks are listed in Table S8).

##### Loop closure

Across the nucleus estimates, we computed the Spearman correlation between the paralattice-minus-lattice preference (the cohort mean of its paired difference) and the loop coupling (defined above). Correlation analysis was repeated (1) with the striatal coupling alone and (2) with the cerebellar coupling alone. Finally, each one of the correlations with the striatal coupling and with the loop coupling was repeated without the six estimates of the anterior nuclei (anteroventral, anteromedial and lateral dorsal, both hemispheres; the anterodorsal nucleus has no estimate). Because the two hemisphere estimates of a nucleus are not independent, they were averaged before each correlation (28 nuclei; SG has a left estimate only), and significance was assessed by permuting the averaged values across the nuclei (10^5^ permutations; Figure 5C; Supplementary Results S5). The p values given beside ρ in Supplementary Results S5 are those of the two-sided Spearman test on the averaged estimates.

##### Vertex-level coupling in motor cortex

To see where, within motor and premotor cortex, each thalamic nucleus couples most strongly, the cortical partial correlations *r*_*n, v*_ were averaged, for each nucleus and subject, within groups of vertices defined by HCP-MMP1.0 area and by block:

Area 4 as a whole.

Caudal and rostral area 4. Each area-4 vertex was assigned to the caudal part if it lay nearer to area 3a, and to the rostral part if it lay nearer to areas 6d, 6a and 6v (Euclidean distance on the S1200 group mid-thickness surface).

Least and most effector-selective area 4. The lowest and highest terciles of effector selectivity over the area-4 vertices. A vertex’s selectivity is the largest of its contralateral foot, hand and tongue contrasts against the average of the five movements, in the HCP S1200 group motor task (Cohen’s *d* maps, MSMAll, 2-mm smoothing; Table S8).

The lattice and paralattice parts of areas 6d and 6mp. Both areas straddle the border.

Areas 3a and 3b.

The rest of lattice cortex: lattice cortex without areas 4, 6d, 6mp, 3a and 3b.

Paralattice premotor cortex: the paralattice vertices of 6d, 6a, 6r, 6v, 6ma, 6mp, SCEF, FEF, PEF, 55b, 24dd and 24dv.

The rest of paralattice cortex: paralattice vertices without paralattice premotor cortex vertices.

Two groups were compared by the paired difference of these averages across the 151 subjects, summarized by Cohen’s *d*_*z*_ and tested by the Wilcoxon signed-rank test, as for the block preference. The comparisons reported are area 4 against the rest of lattice cortex, rostral against caudal area 4, the least against the most effector-selective tercile, and the lattice against the paralattice part of 6d and of 6mp.

#### Generality and replication

The computation was repeated, with every parameter re-derived from the cohort considered (its own floor, majority of ⌊*N* / 2 ⌋+ 1 subjects, cohort geodesic; the null pairs, heterotopic pairs with no plausible direct or indirect connection, were those drawn for the 7T cohort). For each dataset, the partitions obtained were scored against the main 7T partition by ARI and by the percentage of vertices in the same block, alongside the agreement of the two hemispheres at homotopic positions. Each hemisphere was always partitioned on its own graph.

##### Split halves

The two halves of the 7T cohort are those described in Hyperparameter and robustness sweeps (Split halves).

##### Movie watching

The four HCP 7T movie runs of 160 subjects (the cohort geodesic was taken as the median over all 160; Data S1) were denoised as the resting-state runs. The movie data were analyzed in two forms. In the raw form, correlations were computed from the cleaned movie series, as for rest. In the residual form, we first removed the stimulus-driven signal shared by all subjects. For each vertex and each frame, we computed the mean of the z-scored series over the 160 subjects, using only the subjects who retained that frame after motion censoring (including the subject itself), 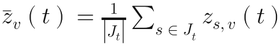, with *J*_*t*_ those subjects. Each subject’s cleaned series at that vertex was then regressed on this group-mean time course by least squares, separately for each subject and vertex over the subject’s retained frames: 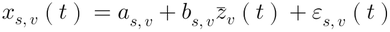, where the intercept *a*_*s, v*_ and the slope *b*_*s, v*_ are fitted for that subject and vertex. The intercept, although close to zero, was kept, because 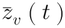 is averaged over a different set of subjects at each frame (*J*_*t*_) and so does not average exactly to zero over one subject’s retained frames. The residuals *ε*_*s, v*_ ( *t* ) were re-standardized (z-scored over the retained frames) before computing the correlations.

##### HCP 3T

The 349 subjects were analyzed by the same procedure (Figure 6C; Supplementary Results S6).

##### Precision fMRI (PAN)

The ten subjects of Ladwig et al.^21^, each with about two hours of resting-state data (Data preprocessing), were analyzed as a ten-subject cohort and each subject alone (Figure 6D,E). For a single subject an edge was a pair with *r* > *τ*_*s*_ in that subject, with weight *r* (computed on standardized concatenated series, *Data preprocessing*). Because the PAN graphs had fewer vertices in their giant components, the margin excluded from the Fiedler sweep was 5% of the giant component (at least 50 vertices) rather than a fixed 200 vertices (on the 7T cohort, the two margins gave an identical partition). A subject’s hemisphere counted as recovering the partition when its ARI against the 7T reference was at least 0.60. The cohort and the single subjects were also analyzed with global-signal regression (the mean over the cortical vertices, regressed after the nuisance fit as in the 7T control) (Supplementary Results S6).

For every cohort, two tables were made from the cohort’s own partition: the block that contains most of each HCP-MMP1.0 area’s vertices, by hemisphere (Table S2; the percentages are provided as source data), and the percentages of each of the twelve Cole– Anticevic networks’ vertices in lattice cortex, paralattice cortex and the peri-allocortex (Table S7). Network membership was taken from the network label files by Glasser parcel (Table S8).

#### Correspondence with cortical maps

We took 27 published cortical maps from the neuromaps annotation library (annotations listed in Table S8). Maps delivered on the fs_LR 32k mesh were used as they were. Maps on the fs_LR 4k and 164k meshes were resampled to it by nearest-vertex lookup on the fs_LR sphere. Maps on the fsaverage meshes were resampled through the fsaverage-to-fs_LR registration sphere with neuromaps (adaptive barycentric, area-weighted); fs_LR vertices that fell in the fsaverage medial wall (read from the neuromaps fsaverage medial-wall mask, desc-nomedialwall) were given no value. The two volumetric MNI152 PET maps were sampled by trilinear interpolation at the vertices of the fs_LR 32k template mid-thickness surface. Statistics were computed on the 59,412 vertices of the two hemispheres pooled, and vertices without a value in a map were left out of that map’s statistics. The evolutionary and developmental expansion maps of Hill et al.^128^ were delivered by that library for the right hemisphere only and were not used; the evolutionary expansion map of Table S5 was that of Xu et al.^129^, defined on both hemispheres. Each map was summarized by its mean within each block, by the Spearman correlation between its vertex values and the block ordinal (assigning value 1 to lattice cortex vertices, 2 to paralattice cortex, 3 to peri-allocortex; with three values and these block sizes, |ρ| cannot exceed 0.84), and by the Kruskal–Wallis effect size across the three blocks, 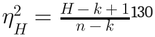 with *k* = 3. The correlation and the effect size were then tested against 1,000 rotations of the block map on the sphere (spin test, below), and the resulting *p* values were corrected across the 27 maps by the Benjamini–Hochberg procedure. No map survived this correction, so the associations were reported descriptively.

Laminar differentiation was estimated from the BigBrain six-layer segmentation. Its layer thickness maps on the fsaverage 164k mesh (BigBrainWarp^116^) were resampled to fs_LR 32k with neuromaps (adaptive barycentric, area-weighted), with the same fsaverage medial-wall mask (desc-nomedialwall). For each vertex, we computed the thickness of layer IV as a fraction of the total thickness of the six layers. This value was available for 58,261 vertices, peri-allocortex included (only the 1,151 vertices in the medial-wall mask or with zero BigBrain thickness have none). The quartiles of this fraction were computed over all 58,261 vertices with a value, peri-allocortex included, and the vertices were divided at these quartiles into four classes, labeled from the lowest to the highest fraction agranular, dysgranular, eulaminate and koniocortical^47^, after the cortical types whose granular-layer development they approximate. We report the block composition of each class and the class composition of each block. Because the segmentation was built for isocortex, the layer IV fraction was not used to characterize the peri-allocortex. To compare lattice and paralattice cortex, we computed the Spearman correlation between the layer IV fraction and block membership over all lattice and paralattice vertices (assigning 1 to lattice vertices, and 2 to paralattice ones), and reported the values descriptively.

### Quantification and statistical analysis

#### Agreement and association

Two partitions were compared by the adjusted Rand index (ARI; 1 for identical partitions, 0 for chance agreement) and by the percentage of vertices in the same block. For the percentage, block labels were first matched one to one, each block being paired with the block it overlaps most. Monotone association was measured by Spearman’s *ρ*. Clusters were matched to networks by the Dice coefficient with one-to-one Hungarian matching. The separation of the two blocks by a per-vertex measure was quantified by the area under the ROC curve and the Youden cut.

#### Effect sizes

Contrasts between two groups of vertices (Figure 3 block contrasts) and between two groups of vertex pairs (Facing sulcal banks) were summarized by Cohen’s *d*, computed with the root-mean-square of the two sample standard deviations, 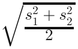 . Effect sizes were computed on the common logarithm for the right-skewed measures (partner distance, the near/far ratio) and on raw values for the others (patch count, the largest-patch share, and sulcal banks contrasts). Paired contrasts across subjects (thalamic analyses) were summarized by Cohen’s *d*_*z*_. The association of a published cortical map with the three blocks (lattice, paralattice, peri-allocortex) was summarized by the Kruskal–Wallis effect size 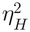, with k = 3 (Correspondence with cortical maps).

#### Statistical significance tests

The single- and two-exponential fits of the edge-survival probability were compared by the Bayesian information criterion (Edge-survival probability model). Statistics that compare a per-vertex map with the blocks were tested by 1,000 independent spatial permutations (spin test). For each one, a rotation was drawn uniformly at random (Table S8), applied to the left fs_LR 32k sphere and in mirror image to the right. Every vertex was reassigned the block label of the vertex nearest its rotated position. Vertices whose nearest rotated neighbor fell in the medial wall were dropped from that rotation, and so were vertices falling in the peri-allocortex for the block contrasts. The *p* value was the fraction of the 1,000 rotations reaching a value at least as large as the observed one, counting the observed value itself, so that 0.001 is the smallest attainable *p*. The spin test was applied to the block contrasts of the per-vertex measures of Figure 3 (Cohen’s *d*) and to the 27 published maps of Table S5 (Spearman’s ρ with the block ordinal and Kruskal–Wallis 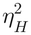). Cohen’s *d* and Spearman’s *ρ* were tested two-sided, on absolute values. Kruskal–Wallis 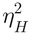was tested one-sided, being independent of the direction of the effect. For the 27 maps only, *p* values were corrected by the Benjamini–Hochberg procedure, separately for ρ and for 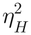.

Paired contrasts across subjects were tested by the Wilcoxon signed-rank test (Thalamic connectivity). The loop-closure correlations (Spearman’s ρ between the paralattice-minus-lattice preference of a nucleus and its striatal, cerebellar or loop coupling, averaged between left and right hemisphere) were tested by permutation. The coupling values of the 28 hemisphere-averaged nuclei were shuffled across the nuclei 10^5^ times (seed 20261104), and the correlation was recomputed on each shuffle. The p value was the fraction of shuffles whose |ρ| was at least as large as the observed one, counting the observed value itself.

The partial correlation of depth with hierarchical level was tested against 100 rewired graphs (Comparison with known hierarchies).

The slope of the homotopic weight on distance from the midline was reported with its 95% confidence interval from ordinary least squares (Homotopic flatness).

#### Conventions

For map and partition statistics, *n* denotes vertices; for the cohort construction, *N* denotes subjects. The variability of the partition across cohorts was estimated by a bootstrap over subjects, with 60 replicates and 2.5th–97.5th percentile intervals (Null models and controls for the partition). No data were excluded beyond the cohort-selection criteria stated above.

#### Software and reproducibility

Analyses were performed in Python. Preprocessing of the HCP series ran under Python 3.12 with NumPy 2.5.1, SciPy 1.18.0, scikit-learn 1.8.0 and nibabel 5.4.2; the PAN series, the correlation matrices, the cohort graphs, all later analyses and the figures ran under Python 3.10.6 with NumPy 2.2.6, SciPy 1.15.3, scikit-learn 1.7.2, nibabel 5.4.2 and matplotlib 3.10.9 (all package versions in Table S8). Spectral decomposition was performed through the ARPACK interface (scipy.sparse.linalg.eigsh); k-means and the ARI were taken from scikit-learn; CIFTI and GIFTI input and output were handled by nibabel. Geodesic distances were computed as exact surface geodesics on each subject’s own mid-thickness surface (gdist), and surface and CIFTI operations used the Connectome Work-bench; the PAN data were processed with fMRIPrep 23.2.0 (Docker image nipreps/fmriprep:23.2.0, with its FreeSurfer 7.3.2 and Connectome Workbench 1.5.0) and tedana 26.0.3. k-means was run with several random initializations (Table S8). The k-means initializations, the null pairs, the random-edge, degree-scrambled and rewired graphs, the shuffled weights, the smoothness-matched surrogates, the spin rotations and the bootstrap came from a single set of random draws. The analysis code was written and run with the assistance of Claude large language models (Anthropic) working under the direction of the authors. Every analysis is specified in these Methods and in Table S8, and the code will be deposited with the derived data (Data and code availability).

## SUPPLEMENTARY INFORMATION

Supplementary Results S1–S6, Figures S1–S4, and Tables S1–S9 follow the references.

Data S1. Subject lists of the four cohorts, in the order used by the analysis, with the excluded subjects and the reason for each exclusion (Excel file).

## SUPPLEMENTARY INFORMATION

### Supplementary Results S1. Short-range correlations and non-neural spread

To further validate the use of short-range functional correlations (3–20 mm), a range rarely examined owing to potential vascular, motion, or volume-conduction artifacts, we evaluated the spatial reach of non-neural spread against the actual behavior of short-range cortical signals.

Assuming a spatially uniform, non-neural process operating everywhere, its contribution at each distance cannot exceed the lowest correlation observed anywhere across the cortical sheet at that distance. This lower envelope (5^th^ percentile over all pairs), computed in each of the 151 subjects on the cohort-median geodesic, decays as A·exp(−d/ *l*env) with *l*env = 1.95 ± 0.34 mm and reaches zero by about 10 mm, whereas the short-range component of the edge probability decays with a length constant of 19.8 mm L and 22.1 mm R (Figure 1C), about ten times longer. Relative to the typical (median) correlation at the same distance, the envelope is 36% in the 3–4 mm bin, 20% at 5–6 mm, 11% at 7–8 mm and, by extrapolation of the fit, 3% at 10 mm, 0.3% at 15 mm and less than 0.1% at 20 mm.

The released series are smoothed on the surface by the HCP pipeline (Gaussian kernel, 2 mm FWHM). The effective smoothness of the series as analyzed is 4.31 ± 0.17 mm L and 4.28 ± 0.17 mm R FWHM (151 subjects; 4.40 ± 0.21 and 4.34 ± 0.23 mm as released), a value that includes the spatial correlation of the BOLD signal itself. The kernel alone, applied to white noise, creates a correlation of 0.02 at 3–4 mm and below 0.005 beyond 4 mm between vertices that share no signal (Methods; Table S8), so its contribution lies almost entirely below the 3-mm inner bound of the window. Short-range correlations also follow the intrinsic geometry of the cortical sheet rather than volume conduction. For the 48,809 retained edges joining facing banks of a sulcus, where geodesic distance is at least twice the Euclidean distance (median geodesic 17.6 mm against median Euclidean 7.3 mm), weights track surface distance. At Euclidean separations of 6–8 mm, the correlation weight lies close to geodesic-matched controls (0.202 against 0.191; Cohen’s d = 0.18) and well below Euclidean-matched controls (0.234; d = 0.47), corresponding to a gap to the Euclidean control that is about three times the gap to the geodesic control. The same holds when the pairs are restricted to a single cortical area (6–8 mm: 0.203 against 0.198 and 0.245). This excludes spread through the volume, such as volume conduction, partial voluming and the projection from volume to surface, but not a hemodynamic signal that spreads along the cortical sheet, such as one carried by pial veins (see below).

Correlation profiles within the 3–20 mm window also have properties that no spatial blurring kernel can produce. First, the short-range weight ratio between lattice and paralattice blocks increases across the window, rising from 1.03 at 3–5 mm to 1.61 at 10–15 mm and 2.44 at 15–20 mm (weight per neighbor within each band); rebuilding the weight map from 10–20 mm edges alone sharpens the contrast between the two blocks from 1.53 to 1.94. Second, within individual cortical areas, where tissue properties and local vasculature are likely uniform, correlations decay in an orderly way with distance. On the full correlation matrices, in all 151 subjects and on the cohort-median geodesic, lattice areas (V1, V2, 3b) and paralattice areas (46, 9a, 10d) have similar correlation magnitudes at 5 mm (mean r 0.33 against 0.38) and diverge with distance, the lattice areas being 1.5-fold higher by 20 mm (0.20 against 0.13). Third, homotopic vertices across the lateral convexity, which are separated by centimeters of subcortical space and belong to distinct vascular territories, track each vertex’s local short-range weight (Spearman 0.85 and 0.87; Figure 3F). These homotopic weights vary slowly with distance from the midline (slope −0.006 per 10 mm, 95% CI −0.006 to −0.005, against a mean of 0.233), whereas an artifactual spread with the 1.95-mm length constant of the envelope would decay by more than ten orders of magnitude across the same span. Finally, the partition does not depend on the shortest distances, where non-neural spread is largest: with the inner bound of the window raised to 5, 7 or 8 mm it stays in place (ARI 0.96–0.99; Supplementary Results S2, Table S6).

The partition does not align with known vascular topography. Venous and arterial density is highest in cingulate, cuneus, precuneus, parahippocampal, insular and temporal cortex, straddling both blocks and the peri-allocortex^1^; cerebral blood flow and blood volume are only weakly related to block membership (ρ = +0.09 and −0.23; Table S5); and quantitative measurements in macaque visual cortex find vascularization to be similar across areas, tracking metabolic demand rather than cell or synapse density^2^.

Large pial veins run along the cortical surface for millimeters to centimeters, so vertices along the same vein could share a signal that follows the sheet, reaches the 10–20 mm range and is not bounded by the lower envelope. We estimated the exposure of each vertex to pial veins, the vessel partial volume near the vertex, from a 7T venous atlas^3^, taking the maximum from the pial surface to 3 mm outward and averaging over 40 subjects (Methods). Lattice cortex was slightly more exposed than paralattice cortex (median vessel partial volume 0.024 against 0.013 L and 0.023 against 0.011 R). Within each block, however, exposure was weakly and negatively related to weight per neighbor (Spearman ρ = −0.09 to −0.21); the lattice excess was present at every level of exposure (lattice/paralattice ratio 1.36–2.06 L and 1.17–2.48 R across quintiles of exposure); and removing the 10–30% most exposed vertices left the lattice/paralattice ratio at 15–20 mm nearly unchanged (2.36–2.39 against 2.44) and the border in place (ARI ≥ 0.98). A vein would also elongate the set of partners of a vertex along its course. The partners of lattice vertices were barely more elongated than partners drawn at random at matched distances (excess axial anisotropy 0.016 L and 0.006 R), and not in relation to exposure; those of paralattice vertices were more elongated (0.041 L and 0.037 R), more so near veins (ρ = 0.37 L and 0.22 R). A venous contribution may thus be present in paralattice cortex, but there it would add correlation and reduce the contrast between the blocks. Finally, the HCP projection to the surface excludes voxels with locally high temporal variability, mostly voxels that contain large vessels (Methods), and the partition was recovered in the PAN data, which were projected without this exclusion.

### Supplementary Results S2. Refinement, uniqueness and robustness of the partition

#### Refinement

Spectral decomposition optimizes a cut that is normalized by total edge weight, which can misplace individual vertices near the border. We therefore refined the partition by single-vertex descent of the normalized cut: vertices on the border were moved to the block to which they are more strongly connected, in small batches or singly, accepting each move only if it lowered the normalized cut, until no move did. This moved 99 L (0.35%) and 154 R (0.55%) vertices and left the partition otherwise unchanged. An exact refinement, which at each step moves the single vertex that lowers the cut most, changes 2 more vertices in each hemisphere. The final blocks comprise 18,952 L and 18,966 R vertices (posterior-central) and 9,003 L and 8,955 R (anterior), with a normalized cut of 0.0108 L and 0.0124 R. The temporal pole, anterior ventral temporal cortex and orbitofrontal cortex have very few surviving edges (median 7 to 14 partners in the window, 2% to 5% of their neighbors there, against 73% L and 74% R in lattice and 48% L and 49% R in paralattice cortex), so their assignment is less clear-cut, but the partition is not altered by their removal (ARI 1.00 in both hemispheres; 100% of the remaining vertices in the same block).

##### Graph conductance

Scoring the splits along the Fiedler ordering by graph conductance, cut(A,B)/min[vol(A), vol(B)], which divides by the smaller block only, selects the same split as the normalized cut (ARI 1.00 in both hemispheres).

##### Mesh vs. surface area spacing

The vertices of the fs_LR 32k mesh are not evenly spaced on the cortex. On the individual mid-thickness surfaces, the area a vertex stands for ranges from 1.7 to 4.6 mm^2^ (5th–95th percentile), and a paralattice vertex stands for more cortex than a lattice vertex (3.3 against 2.9 mm^2^ on average) and therefore has fewer neighbors (394 against 464). Weighting each edge by the areas of its two vertices (Methods) makes every part of the cortical sheet count in proportion to its area, so that the normalized cut compares amounts of cortex rather than numbers of vertices. Without the area weights the partition is nearly the same (ARI 0.97 in both hemispheres; 221 L and 229 R vertices change block; Table S6).

#### Uniqueness

We tested other divisions of the ordering by restarting the descent from 121 splits spaced evenly along the Fiedler eigenvector. The starts that end within 5% of the lowest cut (15 L and 9 R, all beginning near the border) end on it or, on the left, on the variant described below (ARI 0.81–1.00 L and 0.96–1.00 R; every start that ends within 10% of the lowest cut agrees with the partition at ARI ≥ 0.81 L and 0.96 R); every other start stops at a local minimum that is shallower; the median start ends at 2.8 L and 2.8 R times the lowest cut (Figure 2C). Over 60 bootstrap replicates of the cohort, the partition agrees with the full-cohort partition at mean ARI 0.977 L (median 0.988) and 0.989 R (2.5th–97.5th percentiles 0.83– 0.995 and 0.978–0.995), with a normalized cut of 0.0106 L and 0.0124 R (0.0090–0.0117 and 0.0106–0.0139); 4 of the 60 left-hemisphere replicates give the variant described below.

A wider search started the descent from 366 splits: the main partition, 61 splits along each of the orderings by eigenvectors 2–6, and 60 regions grown from random vertices to 10–50% of the hemisphere (Methods). Divisions unrelated to the partition (ARI < 0.3) have a normalized cut at least 2.1 L and 2.3 R times that of the partition, and divisions at ARI < 0.6 at least 1.8 L and 1.7 R times.

In the left hemisphere, two partitions are close in cost (normalized cut 0.0105–0.0108). The first one, reported in the main text, keeps a ventral strip of the central region in the posterior-central block. A variant reached from some restarts moves this strip of about 1,050 vertices (ventral area 4, 3b, OP4, 3a, 1, OP2-3, PoI2, Ig, 43 and 6v) to the anterior block (ARI 0.85 with the reported partition). The main text reports the first partition because it is almost identical to the single partition identified in the right hemisphere (93% of the homologs of the strip vertices lie in the posterior-central block, homotopic ARI 0.97 against 0.84 for the second partition). Moreover, the strip lies in the posterior-central block in 56 of the 60 bootstrap replicates, and the other cohorts (movie, 3T, PAN) assign only 0.4–8% of it to the anterior block.

#### Robustness

The partition does not depend on the precise choice of short-range window. Neighboring windows give the same solution: a 3–18 mm window agrees with 3–20 mm at ARI 1.00 L and 0.99 R, a 3–15 mm window at 1.00 and 0.99, and a 3–12 mm window at 0.99 and 0.98, with block sizes within 1% of the hemisphere’s vertices. Raising the lower cutoff, which removes the shortest distances most exposed to the point-spread function, leaves the partition in place as well: a 5–20 mm window agrees at ARI 0.96 L and 0.96 R, a 7–20 mm window at 0.98 and 0.98, and an 8–20 mm window at 0.98 and 0.99.

The partition does not depend on how the floor is set: a stricter per-subject floor (local false-discovery rate 0.1, median τs 0.149; ARI 0.97 L and R) gives the same partition, and so does the replacement of the subject-specific floors by a single cohort-wide threshold, r > 0.126, chosen to retain the same number of edges, which selects 98% of the same pairs (ARI 0.99 L and R; Table S6; the processing chain and its parameters are summarized in Figure S2 and Table S9).

The partition is also robust to the number of subjects required to support a connection (one third or two thirds of the cohort instead of a majority: ARI 0.96–0.99; 90% of subjects, which retains a quarter of the connections: 0.72 L and 0.93 R). The convention used to average a connection’s weight across subjects does not move the right border (ARI ≥ 0.99) and, on the left, selects between the two partitions described above (Table S6). Split halves of the cohort, carried through the entire chain independently (null floors, minimum number of subjects supporting a connection, spectral decomposition and refinement) agree with each other at ARI 0.98 L and 0.98 R (99.5% and 99.6% of vertices in the same block) and with the full cohort at 0.98–0.99 in both hemispheres, with block sizes within 1%. Analyzed separately, the women (89) and the men (62) of the cohort give the same partition: ARI with the full cohort 0.98 L and 0.99 R for the women and 0.99 L and R for the men (Table S6), and 0.97 L and 0.98 R between the two groups.

The partition also does not depend on particular preprocessing steps. Regressing the global signal from every subject’s time series and applying the identical analysis pipeline recovers the same two-block organization in both hemispheres, with the same single eigengap (λ_3_/λ_2_ 5.08 L and 5.20 R) and 99.8% L and 99.7% R of vertices in the same block (ARI 0.99 L and 0.99 R; Table S6).

With the peri-allocortex left in, and each subject’s floor recalibrated to include its pairs, the two blocks are unchanged (ARI 0.99 L and 1.00 R; the peri-allocortex vertices are then counted in the blocks; Table S6).

### Supplementary Results S3. The role of geometry, density, and range

#### Geometry of the cortical surface

We asked to what extent geometry alone might establish a fault line in the cortical surface. To do so, we coupled every pair of vertices within the short-range window by the Green’s function of geodesic distance (Methods). On this graph, the lowest normalized cut divides each hemisphere front to back near the border identified based on the short-range functional connectivity (ARI with the partition 0.79 L, 0.78 R) with a small gap after the second eigenvalue (λ_3_/λ_2_ = 2.5 L, 2.6 R). The geometric border runs along the central sulcus: most of area 4 (75% L, 85% R), area 3a and area 6d go to the anterior side, and in both hemispheres most of the posterior insular area PoI2 goes to the posterior side. The partition in the main text lies one gyrus further forward, between primary motor and caudal premotor cortex and the rostral premotor and prefrontal cortex, corresponding to where tracing studies in the macaque place a change in anatomical connectivity patterns^4–6^. The geometric bottleneck is shallow and undistinguished. Its lowest cut (0.035 L and R) is 3.2 L and 2.8 R times more expensive than that of the partition (0.0108 L, 0.0124 R). More than a third of the splits along its Fiedler ordering come within 20% of its minimum (38% L, 39% R, against 4% L and 3% R for the functional graph); restarting the descent from 121 splits along the ordering gives a median cut only 1.32 L and 1.27 R times the lowest; and divisions unrelated to the border (ARI < 0.3) come within 6% L and 7% R of the minimum. The second eigenmode of the cortical surface likewise runs along the long axis of the hemisphere^7,8^.

The functional connectivity bottleneck is about three times deeper and, apart from the left-hemisphere variant described in Supplementary Results S2, has no near-equal alternative. It is also not simply a consequence of the smoothness of the graph. Divisions along 1,000 random maps with the same spatial smoothness as the graph’s own low-frequency modes give normalized cuts 7.2 L and 6.5 R times higher on average (Methods). The best of them (0.0131 L, 0.0149 R) are higher than the sweep of the empirical eigenvector before refinement (0.0110 L, 0.0127 R).

#### Connectivity density and pattern

The location of the border depends on differences in the density of short-range connections regardless of the specific arrangement of short-range connections of each vertex. However, that arrangement matters at the border, where it deepens the bottleneck.

If each vertex keeps its number of partners and their distances but the partners themselves are drawn at random from its 3–20 mm neighbors, the border stays in place (ARI 0.95 L and 0.93 R) with a single bottleneck. The same holds if the partners are drawn from the 3–20 mm neighbors without regard to distance or are replaced by each vertex’s nearest neighbors (ARI 0.94 and 0.93 L, 0.93 and 0.92 R; Methods). However, the density of each vertex is not enough by itself: if dense and sparse vertices are scattered over the cortex, what remains is a shallow fault line resembling that of the geometric graph (best normalized cut 0.031 L and 0.030 R, 2.9 and 2.4 times that of the partition, ARI 0.76 L and 0.77 R). Nor does the partition simply pick out the most densely connected vertices. If they are grouped into a block, whether the densest 5% or the densest 50% of the hemisphere, the normalized cut that separates that block from the rest is at least 16 times higher than at the border (Methods).

Over the 3–20 mm window, 10.6% L and 11.9% R of the vertex pairs that straddle the border are connected, against 53% and 54% expected from the within-block rates at the same distances, the two blocks pooled (ratio 0.20 and 0.22). The crossing pairs that are edges (27,492 L and 34,867 R) are also weaker and less consistently supported than the edges within either block: mean weight 0.17 L and 0.16 R, against 0.24 within the posterior-central block and 0.21 within the anterior block; mean support 98 L and 97 R of 151 subjects, against 119 and 120 for the posterior-central block, and 109 in both hemispheres for the anterior block. The deficit is present at every distance above 4 mm and widens with distance, from three-to fourfold at 10 mm to about ninefold at the outer bound of the window. Part of this deficit follows from the difference in density, but the actual partners of each vertex deepen it: when the partners are drawn at random with their number and distances preserved, the ratio rises to 0.31 L and 0.38 R, and the cut is 1.8 L and 2.0 R times shallower (normalized cut 0.019 L and 0.025 R; 2.0 to 2.6 times shallower with nearest or distance-blind random partners, 0.022 to 0.032). Indeed, nearly all of each vertex’s short-range partners lie within its own block (all of them for more than nine in ten of all vertices, 93% L and 91% R), for sparsely connected vertices as much as for dense ones.

The cortex on either side of the border is itself sparsely connected: within-block pairs of the same lengths as the crossing pairs and at the same distances from the border are connected in 28% L and 37% R of cases in the posterior-central block and in 23% L and 32% R in the anterior block, half to two thirds of the rate expected at the same lengths from the two blocks as a whole; pairs that cross the border are again connected less than half as often. Thus, the difference in density defines the location of the border, and the partners each vertex keeps make the bottleneck deeper.

The bottleneck is also present without any threshold. Taking for every pair of the window its raw correlation (mean of the cohort), the pairs that cross the border correlate about half as strongly as within-block pairs of the same length (observed over distance-matched expected mean r, 0.50 L and 0.51 R, against 0.20 L and 0.22 R for the fraction of pairs that are edges). Averaged over each vertex’s neighbors, the raw correlation dips at the border, 17% L and 22% R below its value on either side, and the fraction of its neighbors that are window partners falls from 0.37 L and 0.46 R on the posterior-central side and 0.39 L and 0.48 R on the anterior side to 0.25 L and 0.30 R at the border (means over the vertices within 3 mm of the border and over those 10–20 mm from it on either side; all neighbors).

#### Connectivity range and strength

Recomputing the cohort graph without the 3 and 20 mm cutoffs, over the full distance range, with every other parameter fixed, yields a single giant cluster: the lowest normalized cut only splits off a peripheral fragment of 433 L and 425 R vertices (ARI 0.03 L and 0.03 R). Furthermore, a graph built from long-range connections alone shows hardly any trace of the two blocks at any cutoff (at 20, 40 and 60 mm, ARI is −0.06, −0.05 and −0.04 L and −0.02, 0.05 and −0.01 R). The strongest pairs at any distance, taken in the same number as the edges of the experimental short-range graph, also produce no partition (ARI −0.06 L and −0.04 R).

Recomputing the short-range graph without thresholding for strength yields a shallow, undistinguished partition similar to the geometric one (Table S6). A graph with the same number of edges as the empirical window graph, drawn at random from all 3–20 mm pairs and given the empirical weights at random (each vertex keeps on average about 56% of its neighbors), still divides front to back near the border (ARI 0.83–0.86 L and 0.88–0.89 R over three draws), but its lowest cut is 0.051 L and R, 4.7 L and 4.1 R times that of the partition, and its spectrum has a smaller gap (λ_3_/λ_2_ = 2.5 L, 2.7 R). The weakest pairs of the short-range connectivity, taken in the same number as the edges of the experimental short-range graph, also yield no partition (ARI −0.01 L and −0.02 R).

Long-range connectivity recovers the established large-scale organization. We analyzed the long-range graph containing only the edges of the cohort graph longer than a minimum length (varied from 20 to 90 mm) up to the longest edges of the hemisphere. With a minimum length of 45 mm or more, the leading eigenvector of the graph reproduces the principal functional gradient^9^ (|r| = 0.83 L and 0.93 R, vertex by vertex). Divided into two (K = 2), the graph of edges longer than 45 mm separates mainly unimodal areas (somatomotor, visual and posterior parietal cortex) from mainly transmodal areas, and cuts across the main partition rather than reproducing it (Figure S3; ARI 0.11 L and 0.13 R). Conversely, the main partition accounts for only 13% L and 14% R of the variance of the principal gradient.

Divided into seven clusters (K = 7, by k-means), the same graph recovers six of the seven networks of Yeo et al.^10^ (Figure S3, left hemisphere; ARI 0.50 L and 0.51 R). Matching clusters to networks one to one, the Dice coefficient (Methods) is 0.83 visual, 0.77 default, 0.64 ventral attention, 0.57 dorsal attention, 0.55 frontoparietal and 0.46 somatomotor, with only the limbic network, which lies in the orbitofrontal and temporopolar signal-dropout zone, lacking a counterpart (right hemisphere: 0.83 visual, 0.79 default, 0.72 frontoparietal, 0.67 ventral attention, 0.56 dorsal attention and 0.49 somatomotor). The convergence is worth noting, since the two derivations differ in data and in method: here, spectral clustering (k-means on the leading eigenvectors of the normalized Laplacian) of long-range correlations at 7T, in 1.6-mm data smoothed on the surface with a 2-mm kernel; there, mixture-model clustering of binarized connectivity profiles at 3T, in 3-mm data smoothed on the surface with a 6-mm kernel.

#### Location of the border

The partition places its border in a belt of premotor, cingulate motor and opercular-insular cortex. We examined whether this primarily reflects a deficit of short-range edges or instead the amount of cortex available for sampling, the quality of the signal, or the folding of the sheet.

The deficit of short-range edges is not confined to the border but extends about 20 mm on either side of it. Measuring for each vertex the fraction of its 7–20 mm candidate pairs that are edges (including peri-allocortex), the value is 0.76 L and 0.76 R in the interior of lattice cortex and 0.52 L and 0.49 R in the interior of paralattice cortex, and falls to 0.05 L and 0.16 R within 2.2 mm of the border (medians over vertices; the interiors are the vertices more than 20 mm from the border). This cannot be explained by a deficit in the candidate pairs of a vertex at 7–20 mm, which is higher near the border than in either interior (489 L and 502 R within 20 mm of the border, against 427 L and 413 R in lattice cortex and 321 and 316 in paralattice cortex). Temporal signal-to-noise differs by less than 10% among these groups (97 L and 92 R at the border, 94 and 89 in lattice cortex, 95 and 97 in paralattice cortex; all 151 subjects, mean over the four runs) while the edge fraction varies more than tenfold.

The border does not follow the central sulcus but runs anterior to it, in front of area 4. Measured as the fraction of 7–20 mm candidate pairs that are edges, the banks and fundus of the sulcus are densely connected within lattice cortex (3b 0.75 L and 0.79 R; 3a 0.60 and 0.65), as is area 4 (0.52 and 0.58), whose median vertex lies 12 mm L and 11 mm R from the border. The connection deficit lies anterior to area 4, in the supplementary motor area (6mp; 0.12 L and 0.25 R), the cingulate motor areas 24dd and 24dv (0.00 in both hemispheres), dorsal and ventral premotor cortex (6d, 6a and 6v; 0.26–0.50), the frontal eye field (0.18 and 0.40) and, on the lateral surface, the frontal and parietal operculum and the posterior insula (FOP2, PoI1, PoI2 and PI, 0.00–0.03).

The insula is divided between the two blocks. The posterior insula and the parietal operculum belong to lattice cortex (Ig, 52, OP1, OP2-3 and OP4 at 98–100%, PoI1 and PI at 100% on the left; on the right PoI1 at 95% and PI at 100%), even though their vertices have few short-range edges: the median vertex of Ig has 51 L and 38 R partners in the window and of PI 18 and 16, against 300 L and 291 R for the median vertex of lattice cortex and 190 L and 198 R for that of paralattice cortex. The anterior insula and the frontal operculum lie in paralattice cortex (AVI, FOP1, FOP3, FOP4 and FOP5 at 0% lattice and MI at 1% or less; the agranular anterior insula is masked as peri-allocortex). Between the two lie PoI2 (37% L and 38% R lattice), area 43 (27% and 19%) and FOP2 (12% and 17%), the areas whose assignment moves most across cohorts, together with the premotor and cingulate motor strip.

The location of the border agrees with anatomical evidence about the intrinsic horizontal connectivity. In the macaque, primary visual, extrastriate, somatosensory and primary motor cortex share a patchy lattice of intrinsic projections, whereas in prefrontal areas 9 and 46 these projections form stripes 200–400 µm wide and 2–4 mm long, in patterns extending over 7–8 mm, reciprocally linked with distant stripes of the same dimensions^11–13^, and in motor cortex the horizontal collaterals form discontinuous patches confined to a body-part representation^14^. Area 4 therefore belongs with the sensory areas, as it does here, and the transition to the striped regime falls within premotor cortex; marmoset tracing places it inside area 6, between a caudal part receiving motor and somatosensory input and a rostral part receiving prefrontal input^15^. The T1w/T2w myelin map, which is high in area 4 and drops at its anterior edge^16,17^, the sensorimotor–association axis^18^ and the evolutionary and developmental expansion of cortex^19^ and its allometric scaling^20^ group motor with sensory cortex in the same way. Higher myelin content in lattice cortex is consistent with its shorter connections, since in sensory and motor cortex the T1w/T2w signal reflects intracortical myelin, much of it in the horizontal fibers of the deep layers^16^.

### Supplementary Results S4. Organization of lattice cortex

#### Partner patches and coreness

The following measures extend three contrasts of Figure 3 in the main text: whether the strongest partners of a vertex form one patch or several (Figure 3B), how deeply the vertex sits in the short-range graph (Figure 3E), and how it is coupled to its homotopic vertex (Figure 3F).

In the posterior-central block the partners of a vertex lie mostly in a single patch around it; in the anterior block they are spread over several. The largest patch contains a median of 84% of a vertex’s partners in the posterior-central block vs. 50% in the anterior block (86% vs. 52% R; d = 1.23 L and R, p = 0.001 and 0.002), and more than half of the anterior-block vertices have partners in five or more patches (60% L and 54% R), vs. 19% and 16% of the posterior-central block. The partners are correspondingly nearer in the posterior-central block: leaving out those closer than 3 mm, the median partner distance is 18.5 vs. 82.6 mm L and 18.7 vs. 93.2 mm R (d = 1.03 L and 1.12 R). The block difference in the number of patches holds when patches are defined differently: it is stable from 100 to 450 partners (Cohen’s d = 1.01–1.10 L, 0.98–1.02 R), declines beyond (0.85 L and 0.87 R at 900), and is similar with single linkage at 5 mm instead of 10 mm (0.98 L and 0.93 R).

The posterior-central block is also the densely interconnected core of the short-range graph. The k-core decomposition, which is computed across the entire hemisphere without reference to the partition, gives a mean coreness of 153 in the posterior-central block vs. 98 in the anterior block L and 153 vs. 101 R (Figure S4). Every vertex with coreness above 200 lies in the posterior-central block, and coreness alone separates the two blocks with an area under the ROC curve of 0.78 L and 0.73 R. The higher density extends beyond the window: counting each vertex’s partners in bands of geodesic distance, as a fraction of the vertices available in each band, the posterior-central block is denser within 20 mm (63% vs. 47–48%) and at 20–50 mm (17–18% vs. 8%), and the relation reverses only beyond 100 mm.

Homotopic coupling follows short-range coupling. The main text gives the correlation of homotopic weight with short-range weight outside the peri-allocortex (Figure 3F); including the peri-allocortex, whose vertices have no window edges, it is 0.85 L and 0.88 R. The homotopic edge is supported by a majority of subjects for 79% of the vertices of the posterior-central block and 75% of those of the anterior block, in both hemispheres, and for 10% L and 9% R of the peri-allocortex vertices.

#### Sectors

The spectrum of the lattice-cortex graph has no single gap (eigen-value ratios 1.28, 1.88, 1.51, 1.10 and 1.10 L and 1.10, 1.92, 1.30, 1.18 and 1.10 R for K = 2 to 6), so the number of sectors was set by reproducibility (Methods). For each K from 2 to 7, the k-means solution, refined by single-vertex descent, was compared between the two hemispheres at homotopic vertices, between two independent halves of the cohort, and between each half and the full cohort. K = 4 is the only value at which all three agreements reach 0.85: the hemispheres agree at ARI 0.92, the half-cohorts at 0.98 L and 0.94 R, and each half reproduces the full cohort at 0.96 to 0.99 (Table S8). At K = 3 the left half-cohorts agree at only 0.74, and at K = 5 and 6 the hemispheres at 0.59 and 0.63. However, the sectors are delineated by far shallower divisions than the main partition: by the same definition, the normalized cut per block is 0.0054 L and 0.0062 R for the main partition and 0.036 L and 0.045 R for the four sectors.

#### Hierarchies

Taking the primary areas as anchors for the respective sectors, the strong connections recover classical sensory streams and hierarchies^21^. We defined the depth of a vertex as its distance from the sector’s primary area (V1, area 3b or A1, the base of the hierarchy) along strong connections: the shortest path through the sector’s full-range cohort graph (all pairs at least 3 mm apart), with each edge counted as the inverse of its weight, so that a path through strong edges is short. Depth is expressed as a percentile within the sector (Figure 4A–D, dark to light; Methods). The visual stream runs from V1 (Figure 4A; left hemisphere, median depth 10) through V2 (32), V3 (38), V3A (46) and V4 (55) to V3B (66), LO1 (71) and PIT (76), then to MST (86), FST (90) and MT (91), and ends in FFC (97) on the ventral surface and in TPOJ2 and TPOJ3 (95) at the temporo-parieto-occipital junction. The somatomotor stream (Figure 4B) runs from 3b (4) through 3a (12), 1 (16) and 2 (33)^21,22^ to 7PC (45), PFt (48), AIP (58) and PF (62), and ends in PFm (85) and the angular gyrus (PGi 86, PGs 93). The auditory-temporal stream (Figure 4C) runs from A1 (1) through the belt (LBelt 4, MBelt 7) and parabelt (PBelt 9) to A4 (13), and then divides into the anterior and posterior branches described in the primate^23,24^. The anterior branch continues through A5 (22) and STSda (36) to STSva (60), TE1a (75) and the temporal pole (TGv 84, TGd 89). The posterior branch runs through STSdp (34) and TPOJ1 (39) to STSvp (57), and the right hemisphere is similar (STSvp 63). The posterior branch is shallower than the anterior one, and the areas beside it, STV (23) and PSL (21), lie at the level of A5 (22). This is consistent with the finding that in the human the posterior superior temporal gyrus responds as early as primary auditory cortex^25^.

Altogether, depth along strong connections agrees broadly with the hierarchies established by anatomy and physiology in primates^21–24^. Assigning the areas of each stream to levels from these studies (Methods), the median depth of the areas follows the levels at Spearman ρ = 0.87 L and 0.85 R in the visual sector, 0.81 L and 0.74 R in the somatomotor sector, 0.99 L and R along the anterior auditory branch and 0.89 L and 0.99 R along the posterior branch. Depth grows with distance from the primary area, and distance along the surface from the same base already gives much of this order (ρ = 0.67 L and 0.70 R visual, 0.81 L and R somatomotor, 0.89–0.99 auditory). The strong connections add to it in the visual, somatomotor and anterior auditory streams (partial ρ given surface distance 0.74 L and 0.68 R, 0.41 and 0.24, and 0.68 and 0.53), more than in each of 100 rewired graphs that keep every edge’s length and weight but replace one of its ends by another vertex of the sector at the same distance (p = 0.01, the smallest possible), but not along the posterior auditory branch (0.14 L, −0.51 R; p = 1.0).

The medial parietal sector has no primary area. Depth was computed in the same way, from the sector’s border (Figure 4D; Methods): the shallowest areas are DVT (4), MIP (14), 7PC (15), RSC (17), ProS (18) and 7AL (20), where the sector meets the visual and somatomotor sectors and the retrosplenial cortex. The deepest vertices lie in the dorsal posterior cingulate cortex and the central precuneus (PCV 74, 23c 76, 31pv 83, 7m 84, d23ab 84, 31a 86, 23d 91, 31pd 92), which include the posterior cingulate areas (23, 31) that in the macaque lie on the parieto-medial temporal pathway^26^. Because the sector has no primary area, the border was compared as base with four bases at the retrosplenial rim (RSC alone; RSC and ProS; RSC, ProS, POS1 and v23ab; RSC with the posterior cingulate areas; Methods), by the correlation of depth with five per-vertex measures that vary with depth in the three sensory sectors. Only from the border did depth follow the sensory sectors on all five, in both hemispheres. The side of the border that faces other cortex, taken alone, gave nearly the same depth (ρ = 0.91 L and 0.90 R with depth from the whole border), whereas the side that faces the medial wall behaved like the bases at the rim. From every base at the rim, depth reversed its relation to the number of long-range edges and to the prefrontal share of long-range edges (for RSC, ProS, POS1 and v23ab, Spearman ρ = −0.33 L and −0.15 R, vs. +0.27 and +0.17 from the border, and −0.81 and −0.84, vs. +0.37 and +0.37), so the border was kept as base. Depth largely follows surface distance from the border (ρ = 0.83 L and 0.81 R) except that vertices with a lower unweighted short-range density are higher in the hierarchy (partial ρ = −0.51 L and −0.52 R). The exception lies mostly in the anterior part of the sector, next to medial sensorimotor cortex (areas 23d, 23c and 5mv), where the unweighted short-range density is three to four times lower than in the middle of the precuneus (0.20–0.27 vs. 0.80–0.89, left hemisphere) and depth exceeds what distance from the border predicts.

The auditory-temporal sector connects with the top of the medial parietal hierarchy. Of its partners in other lattice sectors at d ≥ 20 mm, 25% L and 24% R lie in medial parietal cortex proper (the sector without its medial motor strip), mostly in 7m, 31pd and 31pv, as expected from the projections of the caudal superior temporal region to posterior cingulate and medial parietal cortex in the macaque^27,28^.

#### Connectors

The lattice vertices that together send three quarters of the lattice edges at d ≥ 60 mm into paralattice cortex make up a fifth of lattice cortex (19% L and R), while still having a higher number of short-range edges than the lattice as a whole (median 429 L and 428 R vs. 300 L and 291 R). Their long-range edges have a median length of 158 mm L and 154 mm R and go mainly to prefrontal cortex, most densely to its caudal and mid-lateral part, and rarely to orbitofrontal cortex, polar area 10pp or the cingulate motor areas (Figure 4I). They run in two largely separate channels: connectors in the anterior inferior parietal lobule, the intraparietal sulcus and the anterior precuneus (PF, PFt, AIP, 7Am) project chiefly to premotor and caudal lateral prefrontal cortex (6r, 6a, 46, IFSa), and those in the angular gyrus and posterior medial parietal cortex (PGi, PGs, 7m, 31pv) chiefly to 8Ad, 8Av and 9m, with hardly any overlap between the two. The paralattice connectors comprise 3,662 L and 3,540 R vertices, each with at least 772 L and 758 R long-range edges into paralattice cortex; the peri-allocortex connectors comprise 864 L and 630 R vertices (4.6% L and 3.3% R of lattice cortex), each with at least 39 L and 35 R long-range edges into the peri-allocortex. Of the multimodal zone (the fifth of lattice vertices with the largest share of their partners in other sectors), 40% L and 29% R are paralattice connectors. The vertices that are both paralattice connectors and multimodal send about a third of all lattice edges into the peri-allocortex. The lattice connectors do not correspond to any one canonical resting-state network. The paralattice connectors are spread over four of the seven networks of Yeo et al.^10^ (default, frontoparietal, dorsal attention and ventral attention), which hold 41%, 24%, 21% and 14% of them (enrichment over lattice cortex 2.4, 3.5, 1.5 and 1.6; under 1% in each of the other three). The peri-allocortex connectors are most enriched in the medial temporal subsystem of the default network (one of the 17 networks of Yeo et al.), which holds 20% of them against 1.9% of lattice cortex (enrichment 10.4); among the seven networks they lie in the default (45%), dorsal attention (28%), visual (14%) and frontoparietal (13%) networks.

### Supplementary Results S5. Thalamic nuclei and the two blocks

This section provides additional details concerning thalamic nuclei described in the main text (Figure 5, Table S4). Specifically, the mediodorsal nucleus’ parvocellular division prefers paralattice cortex in both hemispheres; its magnocellular division couples more strongly with the peri-allocortex than with either block in both hemispheres. The weak and non-selective prefrontal coupling of the mediodorsal nucleus at rest is in line with earlier work showing that its resting-state connectivity is distributed across several cortical networks^29,30^.

Of the anterior nuclei, all three couple more with paralattice cortex and with the peri-allocortex than with lattice cortex (Table S4); the anteromedial and lateral dorsal nuclei couple about equally with paralattice cortex and the peri-allocortex, whereas the anteroventral nucleus prefers paralattice cortex. Intralaminar thalamic nuclei (centromedian, central lateral, central medial, parafascicular) provide predominantly diffuse, modulatory inputs to superficial cortical layers^31^. The centromedian and parafascicular nuclei receive basal ganglia output^32^ and the central lateral nucleus cerebellar output^33^. Among the intralaminar nuclei, the central lateral and central medial nuclei prefer paralattice cortex, the centromedian nucleus prefers lattice cortex, and the parafascicular nucleus prefers paralattice cortex. The ventral medial nucleus, which receives basal ganglia output and projects diffusely to layer 1, has no consistent preference. The midline medioventral nucleus (reuniens), like the magnocellular mediodorsal division, couples most with the peri-allocortex.

The relation between loop coupling and cortical preference (main text) was tested with the two hemisphere estimates of each nucleus averaged, since they are not independent (Spearman ρ = +0.83 against the mean of the two partial correlations, n = 28 nuclei, p = 4 × 10^−8^; +0.82 with the anterior nuclei excluded, p = 6 × 10^−7^; +0.80 against the striatal and +0.80 against the cerebellar coupling alone; permuting the averaged values across the nuclei, p < 10^−5^; 2 × 10^−5^ with the anterior nuclei excluded). Anatomically, primary motor cortex is an exception: it lies in lattice cortex, although its main thalamic input comes from the ventral lateral nucleus, which relays cerebellar and pallidal output^34^. The ventral lateral nuclei couple with area 4 less than with the rest of lattice cortex (VLa d = −0.69 L and −0.85 R), and less still than with paralattice cortex, while the ventral posterior and centromedian nuclei couple with area 4 more than with the rest of lattice cortex (VPM d = +1.92 and +1.88; CM d = +1.77 and +1.56), as they do with somatosensory cortex. This result is based on the partial correlation of fMRI signals, which need not follow the strength of an anatomical projection. Within area 4, the vertices that couple more with the ventral lateral and centromedian nuclei are the least selective for movements of a single body part^35^.

Of the five nuisance models compared (Methods), the one used here (trend, censoring and CSF signal) agreed best with the established cortical targets of the specific nuclei: 36 of the 43 assignments that Table S3 predicts and 19 of 20 estimates for nuclei with well-established targets, against 32 or 33 and 17 or 18 for the other models. The models differed mainly by a uniform shift: in every model the effect sizes kept the order they had without nuisance regression (Spearman ρ ≥ 0.99), and adding the white-matter signal shifted all of them toward paralattice cortex (by about 0.12). Censoring alone reproduced the assignments obtained without any nuisance regression.

### Supplementary Results S6. Generality: replications and correspondence with cortical maps

#### The main partition is preserved during stimulus-driven activity

For the HCP 7T movie-watching cohort, the same inclusion criteria applied to the resting-state cohort were applied to the movie scans: 160 of the 175 candidates passed them (six incomplete acquisitions, nine motion failures). We constructed two datasets. In the raw dataset, per-subject correlation matrices were computed directly from the preprocessed movie time series. In the residual dataset, the stimulus-locked component was first removed by regressing, at each vertex, the inter-subject mean time course, leaving the correlations of what remains after subtraction of the shared evoked response.

Both movie datasets return the two blocks (Figure 6B, and Figure S1 for the residual dataset; the dashed line in Figures 6 and S1 is the resting-state border). Against the resting-state partition, the residual dataset agrees at ARI 0.98 L and 0.97 R (99.5% L and 99.3% R of vertices keeping their block), the raw dataset at 0.98 L and 0.97 R (99.4% L and 99.2% R). Block sizes match those at rest: 18,950 / 9,005 L and 19,077 / 8,844 R vertices in the residual dataset and 18,842 / 9,113 L and 18,951 / 8,970 R in the raw dataset, against 18,952 / 9,003 L and 18,966 / 8,955 R at rest. The two hemispheres, partitioned independently, agree at 99.3% and 99.3% of homotopic positions (ARI 0.97 and 0.97), as at rest (99.1%), and the spectrum keeps its single gap (λ_3_/λ_2_ = 5.03 L and 4.12 R residualized, 5.84 and 5.09 raw). Removing the stimulus-locked component makes almost no difference, showing that the main partition does not depend on the stimulus being present or absent from the signal.

The disagreement with resting-state is small (0.5–0.8% of vertices). Lattice cortex keeps 99.3–99.8% of its vertices and paralattice cortex 98.3–99.7%. The vertices that change block lie at the border, in the premotor, motor and cingulate motor belt (6d, 6mp, 4, 24dd, FEF) and the posterior insula and parietal operculum (PoI2, 43). A full per-parcel listing is provided in Table S2.

#### The main partition replicates at 3T in a larger cohort

The 3T replication used the HCP 3T cohort of unrelated, low-motion subjects (N = 349, 43 of them also in the 7T cohort; fs_LR 32k surface), with the same preprocessing chain except for field strength and the resulting per-subject noise. Every step was rederived from the 3T cohort itself: a null floor for each subject (median 0.088, quartiles 0.058 and 0.127), geodesic distances from the cohort’s own surfaces, and an edge required a majority of the 349 subjects (175), retaining 1,590,325 L and 1,553,266 R edges. Spectral decomposition and descent were then applied exactly as at 7T.

As at 7T, the 3T graph subdivides into two blocks (Figure 6C; the dashed line in Figure 6 is the 7T border), with block sizes 19,443 / 8,512 L vertices and 19,732 / 8,189 R (lattice, paralattice), against 18,952 / 9,003 L and 18,966 / 8,955 R at 7T. The spectrum has its largest gap after the second eigenvalue (λ_3_/λ_2_ = 5.1 L and 5.5 R). The 3T partition matches the 7T partition vertex for vertex at ARI 0.91 L (97.7% of vertices keeping their block) and 0.89 R (97.1%). In both hemispheres the border lies slightly more anteriorly at 3T: 6.4% L and 8.7% R of the 7T paralattice are assigned to lattice cortex, chiefly the frontal eye field, 55b, PEF, ventral premotor area 6v, area 43, the posterior insula (PoI2) and the frontal operculum (FOP2), while the 7T lattice is kept almost entirely (99.6% L and 99.9% R). The two 3T hemispheres, each partitioned alone, agree with each other at ARI 0.95 (98.9% of homotopic positions), close to the 7T hemispheres (0.97). Area by area, the assignments are otherwise those of the 7T partition: the auditory strip lies within lattice cortex, areas 52 and TA2 included, as do primary somatosensory cortex (3b, 100% in both hemispheres), the early visual areas and MT (100%), and the medial parietal areas; dorsolateral and medial prefrontal areas are paralattice (10d and p32, 0% lattice). A full per-parcel listing is provided in Table S2.

#### The main partition replicates at the single-subject level with precision fMRI

The Precision targeting of Association Networks dataset (PAN^36^; OpenNeuro ds006598) includes 10 deeply phenotyped individuals, each with about two hours of resting-state fMRI acquired on a 3T Siemens Prisma using a multi-echo multiband sequence (2.4 mm isotropic, TR = 1.355 s, multiband factor 6) with real-time motion monitoring. We processed the raw data with fMRIPrep 23.2.0, the pipeline the dataset’s authors used, on each subject’s own anatomy, and denoised each run by multi-echo ICA (Methods).

Each PAN subject was treated as an independent dataset. The series were cleaned as the HCP series, with the non-BOLD components identified by multi-echo ICA removed in the same fit and without global signal regression, and geodesic distances were measured on each subject’s own surface (Methods). The PAN preprocessing has no counterpart of the temporal-ICA cleanup that removes global structured noise from the HCP series^37^, and the null floor is more than twice as high as at 7T (median τ_s_ 0.28, against 0.11). In the cohort graph an edge had to pass in a majority of the ten subjects (six); in a single-subject graph an edge is a pair above that subject’s floor. Spectral decomposition and descent were applied as at 7T, separately for each hemisphere (Methods).

The ten-subject cohort graph matched the 7T reference at ARI 0.82 L and 0.86 R (95.5% L and 96.3% R of vertices), with the two hemispheres agreeing at 97.8% of homotopic positions (ARI 0.91) (Figure 6D). As in the HCP 3T cohort, the border lies more anteriorly than at 7T: 14.3% L and 9.9% R of the 7T paralattice are assigned to lattice cortex, chiefly the frontal eye field, 55b, PEF, ventral premotor area 6v, area 43 and the frontal operculum (FOP2) in both hemispheres and, on the left, 6r, FOP1, the posterior insula (PoI2) and parts of 6mp, 24dd and 6a, while the 7T lattice is kept almost entirely (more than 99.9% L and 99.2% R). The PAN partition agrees with the HCP 3T partition at ARI 0.89 L and 0.93 R. Scoring each single-subject hemisphere (Figure 6E) against the 7T reference, the lattice–paralattice division is recovered (ARI ≥ 0.60) in all 20 hemispheres (ARI 0.62 to 0.91; median 0.81). A full perparcel listing is provided in Table S2 (sub-PAN04).

With global-signal regression (the mean over the cortical vertices, regressed after the nuisance fit as in the 7T control), the cohort partition matches the 7T reference at ARI 0.92 L and 0.88 R, and 19 of the 20 single hemispheres reach ARI 0.60. Without multi-echo ICA, the cohort partition matches the 7T reference at ARI 0.92 L and 0.86 R, and 18 of the 20 single hemispheres reach ARI 0.60; without it and with the six motion parameters and their temporal derivatives added to the nuisance fit, at ARI 0.82 L and 0.85 R, with the left border in the anterior position found with multi-echo ICA (ARI 0.96 L and 0.94 R between the two partitions) (Tables S6 and S8).

#### Correspondence with published cortical maps and laminar differentiation

Each of the 27 maps (Table S5) was compared with the block ordinal index by Spearman ρ(block) (main text). Because this signed correlation assumes an ordering of the three blocks, we also report the Kruskal–Wallis effect size η^2^H. Each map was tested against 1,000 rotations of the block map on the sphere. Eight of the 54 tests fall below 0.05 and none survives correction across the 27 maps (Table S5): no map follows the partition more closely than a rotation of it does, so the correspondences below must be considered as descriptive.

The pattern of the MEG maps (main text) is consistent with TMS-EEG measurements of cortical natural frequency, whereby occipital cortex responds predominantly in the alpha band, superior parietal cortex in beta, and frontal and premotor cortex in the beta–gamma range^38^. The next strongest are the first principal component of cortical gene expression (−0.62: lattice high, paralattice low), theta power (+0.57), μ-opioid receptor density (+0.50), T1w/T2w myelin (−0.48), the sensorimotor–association axis (+0.47), the principal functional gradient (+0.37), 5-HT1A receptor density (+0.33) and cortical thickness (+0.33).

With each vertex assigned a laminar-differentiation type from BigBrain histology (main text), layer 4 occupies a larger fraction of cortical thickness in lattice than in paralattice cortex (medians 0.087 vs. 0.068; Spearman ρ = −0.32 with block membership). The proportion of lattice cortex increases from 44% in agranular cortex to 57% in dysgranular, 74% in eulaminate and 84% in koniocortical cortex, whereas paralattice decreases from 50% to 14%. Lattice cortex is more often granular (61% eulaminate or koniocortical) than paralattice cortex, which is more often agranular (71% agranular or dysgranular). Thus, although the partition follows a clear gradient of laminar differentiation, the overlap is incomplete and the lattice–paralattice boundary does not simply recapitulate cytoarchitectonic boundaries.

---

**Figure S1.**
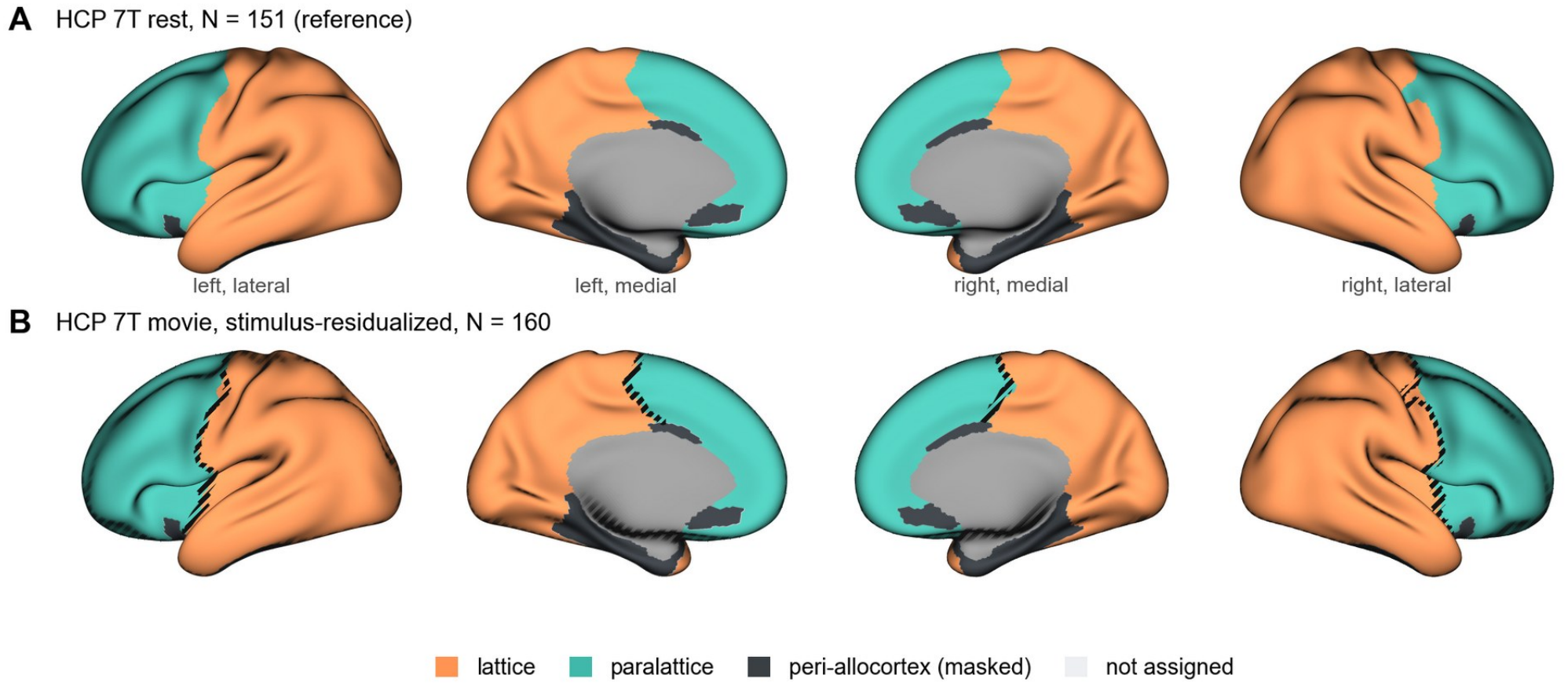
The main partition during movie watching after removal of the stimulus-locked component (related to Figure 6). Both hemispheres, lateral and medial views. **(A)** The HCP 7T resting-state reference partition (N = 151), as in Figure 6A. **(B)** HCP 7T movie watching (N = 160) after regression, at each vertex, of the inter-subject mean time course (ARI 0.98 L and 0.97 R against rest). Orange, lattice; teal, paralattice; gray, the peri-allocortex; near-white, vertices with no edge in the graph, not assigned. In B the dashed line marks the border of the resting-state partition shown in A.

**Figure S2.**
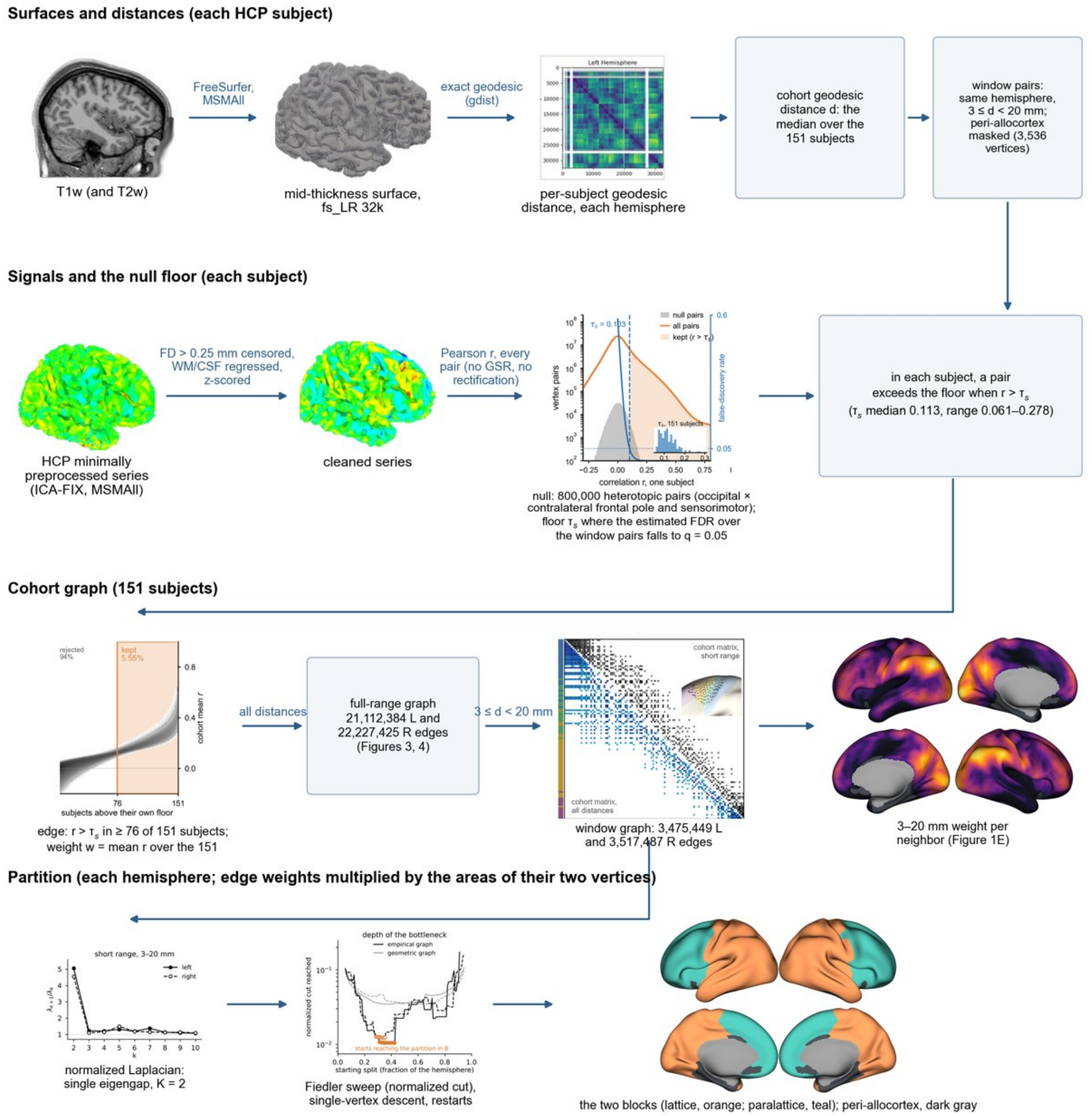
Diagram of the pipeline that generates the objects used in the analysis (related to Methods). The steps that turn the HCP data into the cohort graphs and the partition, with the quantities of record. Top row: each subject’s mid-thickness surface (fs_LR 32k) gives exact geodesic distances, whose median over the 151 subjects is the cohort distance *d*; the window is the within-hemisphere pairs with 3 ≤ *d* < 20 mm, the peri-allocortex (3,536 vertices) excluded. Second row: the minimally preprocessed series are censored (FD > 0.25 mm), cleaned of the white-matter and CSF signals and z-scored, and Pearson *r* is computed for every pair; each subject’s floor τ_*s*_ is set where the false-discovery rate estimated from 800,000 heterotopic null pairs falls to 0.05 over the window candidates. Third row: a pair is an edge when *r* > τ_*s*_ in at least 76 of the 151 subjects, with weight the mean *r* over all 151; the full-range and window graphs and the 3–20 mm weight per neighbor of each vertex. Bottom row, on the window graph with every edge weight multiplied by the areas of its two vertices: the eigengap of the normalized Laplacian, the Fiedler sweep with single-vertex descent, and the resulting blocks.

**Figure S3.**
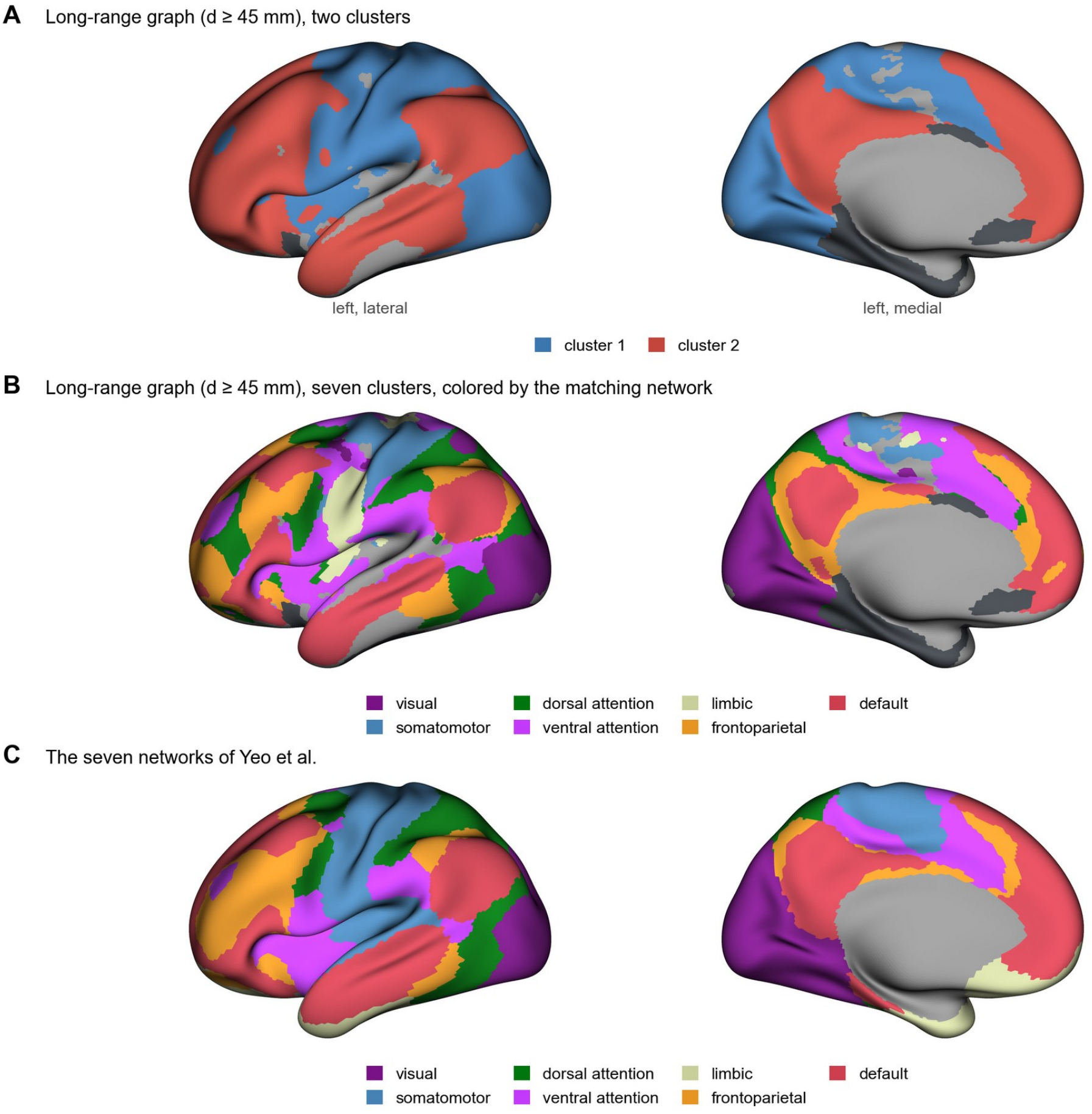
Long-range connectivity recovers the principal gradient and the canonical networks (related to Figure 2). Graph of the cohort edges at d ≥ 45 mm (HCP 7T, N = 151), left hemisphere, lateral and medial views. **(A)** Division into two clusters (median split of the leading non-trivial eigenvector). **(B)** Division into seven clusters (k-means on the six leading non-trivial eigenvectors), each cluster in the color of the network of Yeo et al.^10^ that it matches; the cluster over the central sulcus matches no network and takes the color of the limbic network, which has no counterpart. **(C)** The seven networks of Yeo et al. Gray, the peri-allocortex and the vertices outside the giant component of the graph.

**Figure S4.**
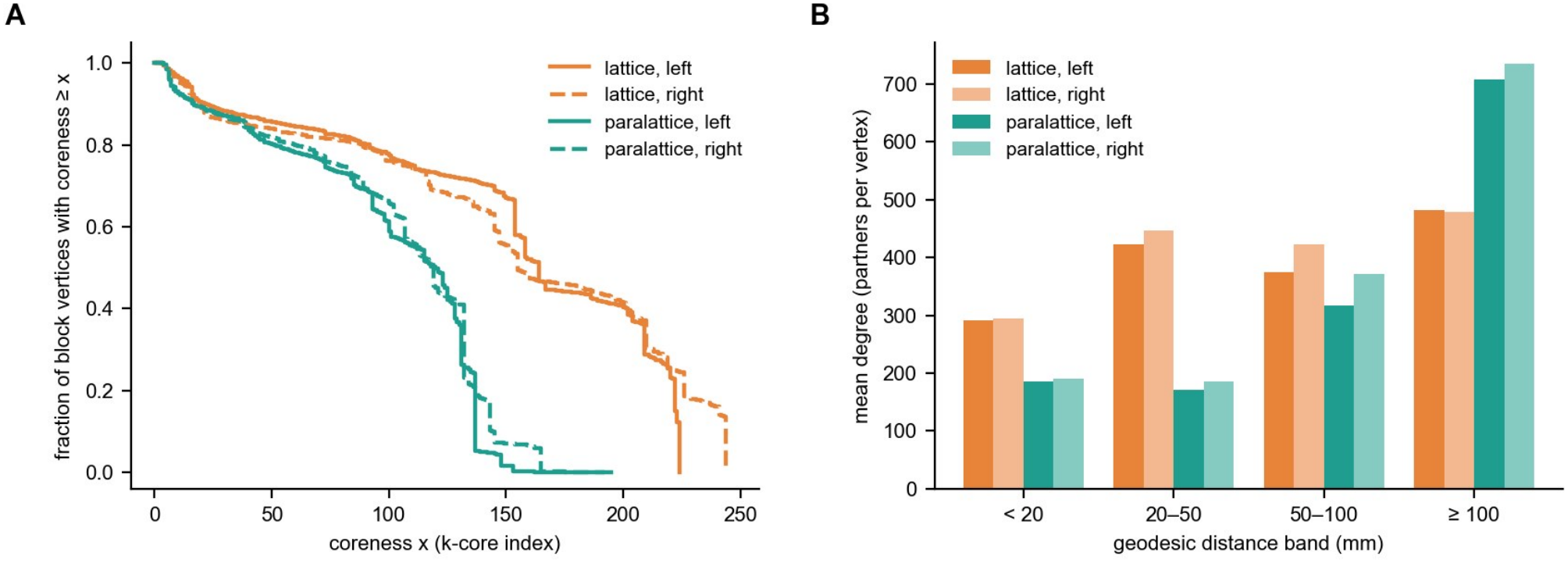
Coreness distribution and degree by distance band (related to Figure 3E). HCP 7T cohort (N = 151). **(A)** Survival curves of vertex coreness for lattice (orange) and paralattice (teal) in each hemisphere’s 3 ≤ d < 20 mm graph: the fraction of each block’s vertices with coreness at least x (mean 153 against 98 L and 153 against 101 R; maxima 224 and 244 against 195 and 192). **(B)** Mean degree per vertex by geodesic distance band in the full-range cohort graph, one bar per block and hemisphere.

**Table S1.** Acquisition parameters for HCP 7T, HCP 3T and PAN (related to Methods).

| Parameter | HCP 7T | HCP 3T | PAN |
|---|---|---|---|
| Scanner | Siemens Magnetom 7T | Siemens Skyra 3T | Siemens Prisma 3T |
| Head coil | Nova 32-channel Siemens head coil | Standard 32-Channel Siemens receive head coil | 64-channel head coil |
| Sequence | GE-EPI | GE-EPI | Multi-echo multiband |
| fMRI voxel size | 1.6 mm isotropic | 2.0 mm isotropic | 2.4 mm isotropic |
| TR | 1.0 sec | 0.72 sec | 1.355 sec |
| TE | 22.2 msec | 33.1 msec | Multi-echo<br>(five TEs, 12–91.2 ms) |
| Flip Angle | 45 | 52 | 59 |
| Multiband | 5 | 8 | 6 |
| In-plane accel. | 2 | Off | 2 |
| Slices | 85 | 72 | 54 |
| FOV | 208 x 208 mm | 208 x 180 mm | 216 mm |
| Phase Encoding | Rest – Day 1: PA/AP, Day 2: PA/AP; Movie – Day 1: AP/PA, Day 2: PA/AP | Day 1: RL/LR, Day 2: LR/RL | AP |
| Runs | rfMRI – 4 runs (900 frames/runs, ~ 1hrs)<br>Movie – 4 runs (921, 918, 915, 901 frames) | rfMRI – 4runs (1200 frames/runs, ~ 57 mins) | rfMRI – 22 to 33 runs per subject (222 frames/run; two runs of 194 and 201) |
| Structural MRI | 3T 3D MPRAGE T1-weighted<br>(0.7 mm isotropic) | 3T 3D MPRAGE T1-weighted<br>(0.7 mm isotropic) | T1-weighted and T2-weighted<br>(1.0 mm isotropic) |
| Participants | *N* = 151 (rest), 160 (movie)<br>(rest: 89 females, 62 males; movie: 97 females, 63 males,<br>age range: 22–36+ years) | *N* = 349<br>(182 females, 167 males,<br>age range: 22–36+ years) | *N* = 10<br>(5 females, 5 males,<br>age range: 18–33 years) |

**Table S2.**
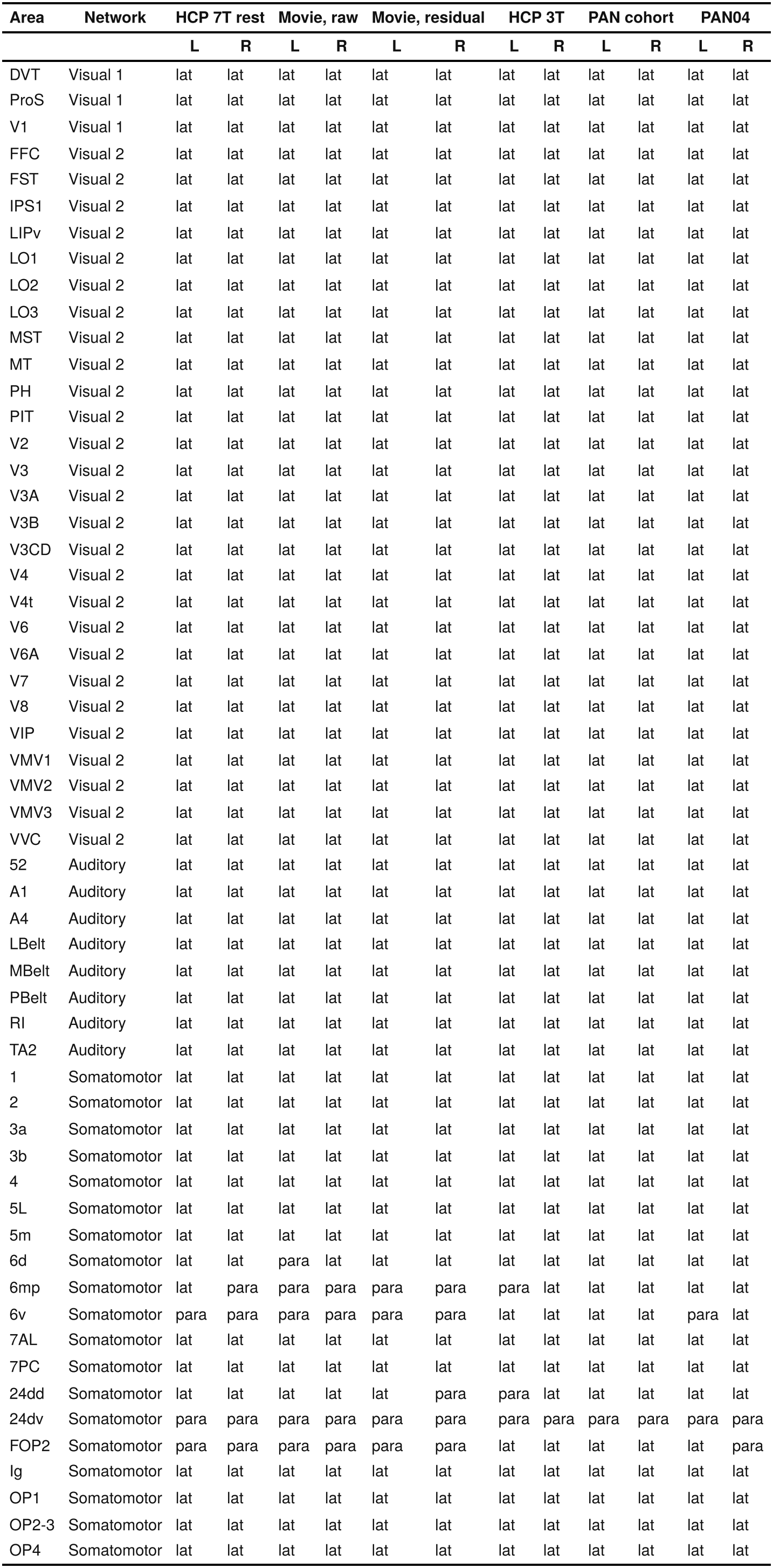
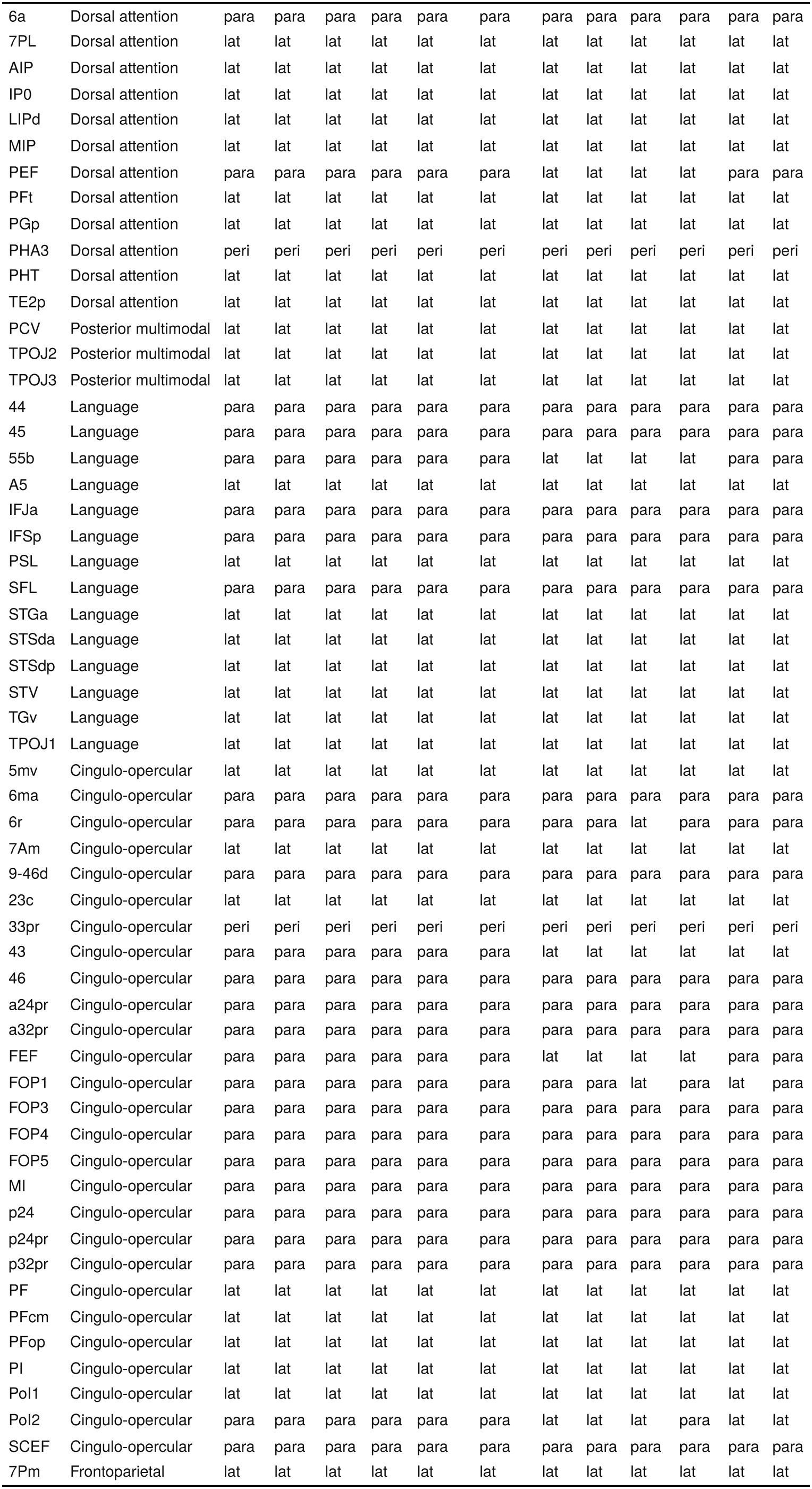
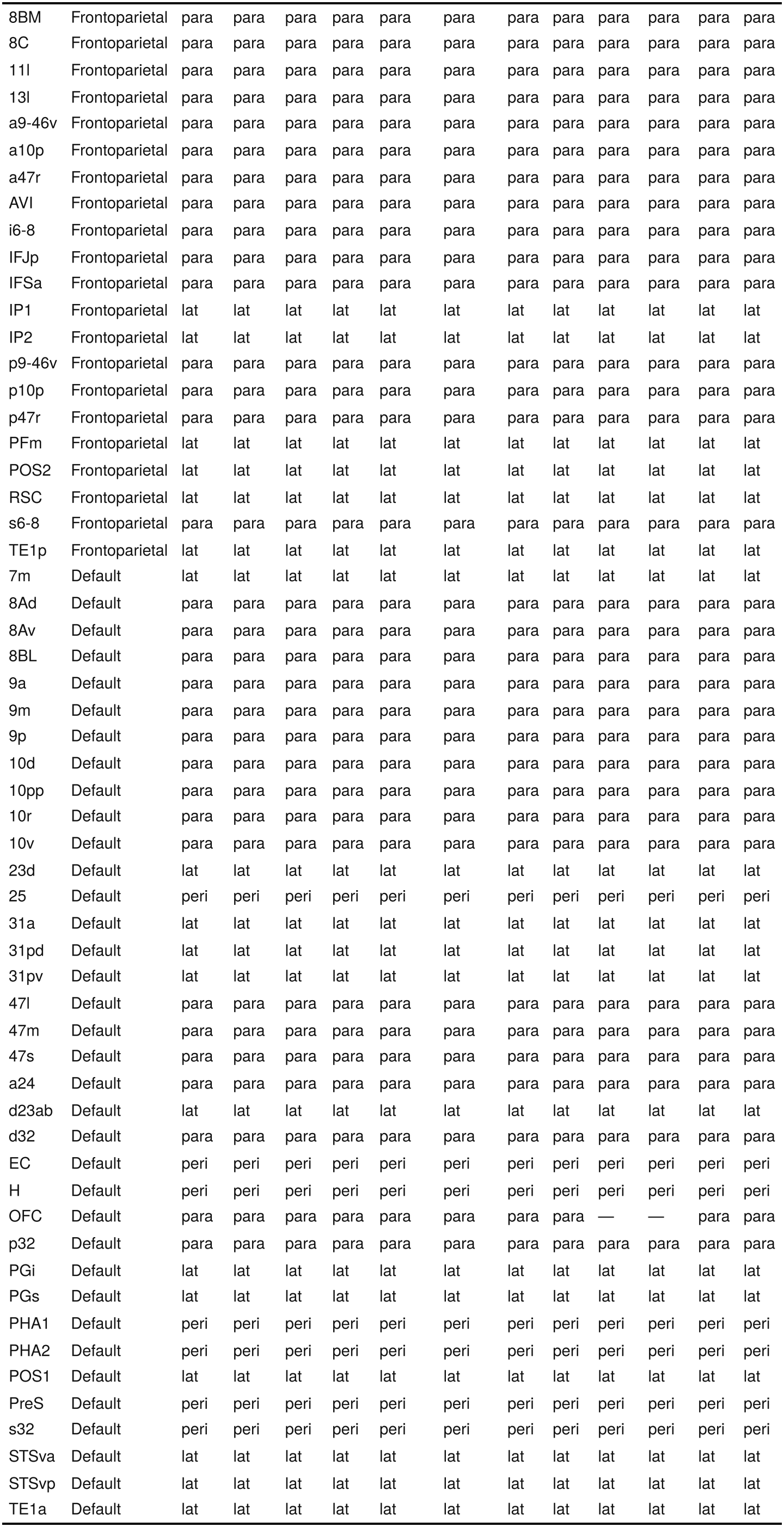
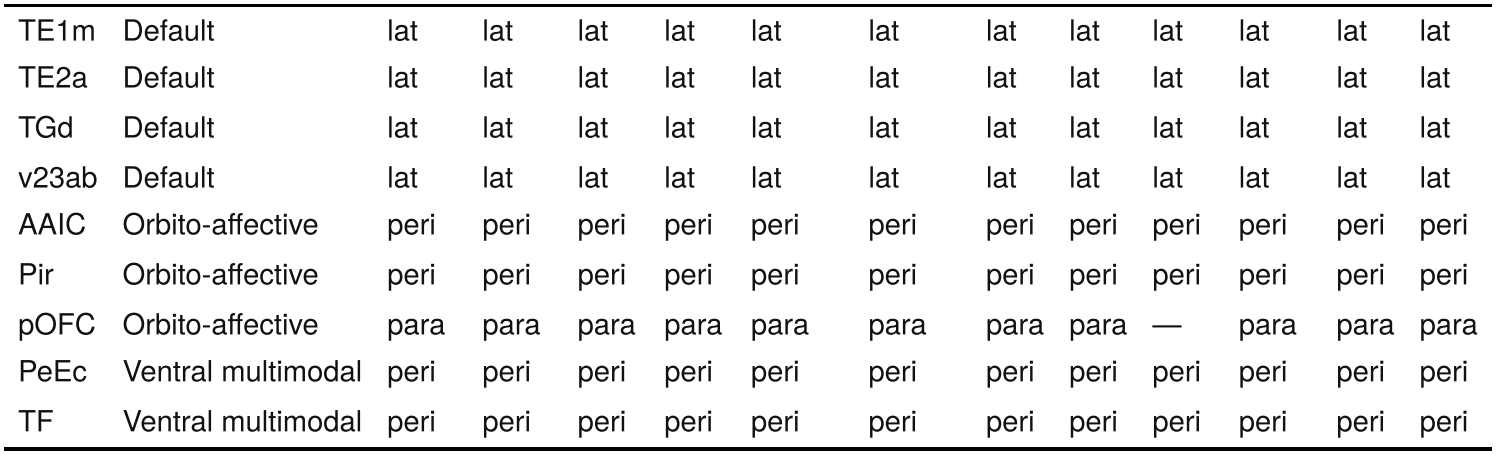
Block assignment of every cortical area in every cohort. For each of the 180 HCP-MMP1.0 areas, left (L) and right (R) hemisphere, the table gives the modal block (lat, lattice; para, paralattice; peri, the peri-allocortex) for the 7T resting-state partition of record, the two movie partitions, the 3T partition, the PAN cohort partition and one PAN subject (sub-PAN04); —, a majority of the area’s vertices are not assigned; the percentage of each area’s vertices assigned to each block is provided as source data.

**Table S3.** Thalamic afferent and loop recipiency for the two blocks and the peri-allocortex. For primate cortical areas, the table lists the principal specific thalamic nucleus providing afferent input (point-to-point, granular-layer, driving connections) and whether that nucleus is a target of basal-ganglia output (pallidal GPi / nigral SNr) or of cerebellar output (references in the first column). Class follows the driver criterion: first-order nuclei are driven by subcortical afferents, higher-order nuclei by cortical layer 5; the basal-ganglia-recipient nuclei (VA, VLo/VLa) have no excitatory driver of either kind and receive large pallidal or nigral inhibitory terminals instead^53^. Block = majority assignment from Figure 2 (HCP 7T, 3–20 mm, each hemisphere alone). “Specific loop” denotes a closed, topographically segregated cortico–basal-ganglia–thalamo–cortical loop through a dedicated relay, as distinct from the diffuse intralaminar system. Note that inferotemporal area TE also has a specific SNr loop, relayed by the magnocellular ventral anterior nucleus VAmc^54^, and its human homolog (TE1a–TE2p, with the temporal pole) lies in lattice cortex, an exception like AIP, M1 and caudal dorsal premotor area 6d.

| **Principal thalamic afferent** | **Class** | **Basal ganglia input** | **Cortical area(s)** | **Partition block** |
|---|---|---|---|---|
| ventral anterior (VApc), ventral lateral oral (VLo)<sup>39,40</sup> | specific, basal-ganglia-recipient | Yes (GPi) | premotor, SMA, pre-SMA | paralattice (6d lattice; 6mp split) |
| VA + paralaminar mediodorsal (MD)<sup>41</sup> | specific; VA basal-ganglia-recipient, MD higher-order | Yes | frontal eye field (area 8) | paralattice |
| MD parvocellular, VAmc<sup>42,43</sup> | specific; MD higher-order, VAmc basal-ganglia-recipient | Yes (SNr) | dlPFC (9, 46), vlPFC (44, 45, 47) | paralattice |
| MD magnocellular, VAmc<sup>43,44</sup> | specific; MD higher-order, VAmc basal-ganglia-recipient | Yes | orbitofrontal (11, 13) | paralattice |
| VA, MD (MDpc, MDdc), midline and intralaminar (Cdc, Pf, CL, Li)<sup>45</sup> | specific; VA basal-ganglia-recipient, MD higher-order; midline and intralaminar non-specific | Yes (VA, MD) | anterior cingulate (24, 32) | paralattice |
| lateral geniculate (LGN)<sup>46</sup> | specific, first-order | No | V1 | lattice |
| inferior + lateral pulvinar<sup>47</sup> | specific, higher-order | No | V2–V4, V3A/B, MT+, extrastriate | lattice |
| ventral posterior (VPL/VPM)<sup>33,46</sup> | specific, first-order | No | S1 (3a, 3b, 1, 2), S2 / operculum | lattice |
| medial geniculate (MGN)<sup>23</sup> | specific, first-order | No | A1, belt / parabelt | lattice |
| lateral posterior (LP), anterior pulvinar<sup>48</sup> | specific, higher-order | No | Superior parietal / IPS (LIP, VIP, MIP, area 7) | lattice |
| medial pulvinar, LP, lateral dorsal (LD)<sup>28,48,49</sup> | specific, higher-order | No | IPL connectors (PF, PFm, PGi/s, IP1/2), precuneus, lateral-temporal STS | lattice |
| pulvinar (medial, lateral, oral) and LP; SNr and dentate output via a thalamic relay not identified (candidates MDmf, VLc, CL)<sup>50</sup> | specific, higher-order | Yes | AIP (anterior intraparietal) | lattice (exception) |
| anterior nuclei (AD/AV/AM), LD; medial pulvinar, midline nuclei<sup>51,52</sup> | — | No | hippocampal formation, subiculum, entorhinal, perirhinal and parahippocampal cortex | peri-allocortex |
| VLo/VLa (pallidal, motor) + VPLo / area X (cerebellar)<sup>34</sup> | specific; VLo/VLa basal-ganglia-recipient, VPLo/X first-order | Motor side Yes, sensory side No | M1 (area 4) | lattice (exception) |
| intralaminar CM, CL, Pf<sup>31,32</sup> | intralaminar (non-specific) | project diffusely to layer 1 and to striatum | (frontal, layer 1) | diffuse |

**Table S4.** Coupling of individual thalamic nuclei with the two cortical blocks, the peri-allocortex and the subcortical loops (HCP 7T, N = 151). Thalamic grayordinates were labeled with the Morel atlas^55–57^; nuclei with fewer than three grayordinates in a hemisphere are omitted (the lateral geniculate nucleus has none). For each nucleus and subject, the mean time course was partially correlated with every cortical vertex of the same hemisphere, controlling for the mean signal of the rest of the thalamus, and averaged within each block and within the peri-allocortex; the columns give Cohen’s d of the paired difference across subjects (paralattice minus lattice, peri-allocortex minus lattice), left and right hemisphere. Striatal r is the partial correlation with the ipsilateral striatum (caudate, putamen, accumbens) and cerebellar r that with the contralateral cerebellar cortex, both controlling for the rest of the thalamus (cohort means). Rows are grouped by the block each nucleus couples with, lattice first, and within each group by class (following Table S3); the nucleus resolved in only one hemisphere (VPLa) is placed by that hemisphere. Loop input names the output nucleus (internal pallidum, substantia nigra pars reticulata, or cerebellar nuclei) that projects to the nucleus. Couples with gives the block preferred at Wilcoxon p < 10^−3^ and |d| ≥ 0.3 (the peri-allocortex requires both of its contrasts); unresolved otherwise. Across the 28 nuclei (the two hemisphere estimates of each averaged), loop coupling scales with paralattice preference (Spearman ρ = +0.83; see text). The series were censored for motion and cleaned of the CSF signal (Methods).

| Nucleus | Couples with |  | Loop input | Paralattice – lattice (d) |  | Peri-allocortex – lattice (d) |  | Striatal (r) |  | Cerebellar (r) |  |
|---|---|---|---|---|---|---|---|---|---|---|---|
|  | L | R |  | L | R | L | R | L | R | L | R |
| **Lattice** |  |  |  |  |  |  |  |  |  |  |  |
| *first-order, sensory* |  |  |  |  |  |  |  |  |  |  |  |
| Ventral posterior lateral, anterior (VPLa) | unresolved | lattice | none | −0.12 | −0.55 | −0.88 | −0.95 | +0.027 | +0.015 | +0.029 | +0.012 |
| Ventral posterior lateral, posterior (VPLp) | lattice | lattice | none | −0.37 | −1.01 | −1.25 | −1.54 | +0.051 | +0.018 | +0.053 | +0.021 |
| Ventral posterior medial (VPM) | lattice | lattice | none | −1.33 | −1.53 | −1.89 | −1.80 | +0.008 | −0.008 | +0.009 | −0.013 |
| Ventral posterior inferior (VPI) | lattice | lattice | none | −1.05 | −1.00 | −1.10 | −1.00 | −0.009 | −0.013 | +0.001 | −0.010 |
| Medial geniculate (MGN) | lattice | lattice | none | −0.88 | −0.51 | −0.86 | −0.34 | −0.011 | −0.003 | +0.000 | +0.009 |
| *higher-order* |  |  |  |  |  |  |  |  |  |  |  |
| Pulvinar, anterior (PuA) | lattice | lattice | none | −1.68 | −1.52 | −1.45 | −1.47 | −0.018 | −0.034 | −0.012 | −0.027 |
| Pulvinar, medial (PuM) | lattice | lattice | none | −0.95 | −0.86 | +0.15 | +0.06 | −0.048 | −0.049 | +0.047 | +0.035 |
| Pulvinar, lateral (PuL) | lattice | lattice | none | −1.00 | −0.79 | −0.99 | −0.97 | +0.010 | +0.007 | +0.013 | +0.024 |
| Lateral posterior (LP) | lattice | lattice | none | −0.33 | −0.75 | −0.95 | −1.31 | +0.031 | +0.007 | +0.048 | +0.070 |
| Posterior (Po) | lattice | lattice | – | −1.29 | −1.15 | −1.08 | −0.87 | −0.020 | −0.018 | −0.015 | −0.017 |
| Limitans (Li) | lattice | lattice | none | −0.63 | −0.76 | −0.75 | −0.81 | −0.012 | −0.025 | +0.016 | −0.002 |
| Suprageniculate (SG) | lattice | — | none | −1.12 | — | −0.91 | — | −0.019 | — | −0.006 | — |
| *intralaminar, caudal* |  |  |  |  |  |  |  |  |  |  |  |
| Centromedian (CM) | lattice | lattice | GPi | −1.09 | −1.20 | −1.73 | −1.94 | +0.016 | +0.001 | +0.009 | +0.001 |
| **Paralattice** |  |  |  |  |  |  |  |  |  |  |  |
| *basal-ganglia-recipient, motor* |  |  |  |  |  |  |  |  |  |  |  |
| Ventral anterior, parvocellular (VApc) | paralattice | paralattice | GPi | +1.68 | +1.86 | +0.85 | +0.49 | +0.105 | +0.108 | +0.109 | +0.143 |
| Ventral anterior, magnocellular (VAmc) | paralattice | paralattice | SNr | +1.27 | +1.45 | +0.94 | +0.55 | +0.036 | +0.036 | +0.051 | +0.053 |
| Ventral lateral anterior (VLa) | paralattice | paralattice | GPi | +1.27 | +1.15 | −0.38 | −0.63 | +0.089 | +0.094 | +0.081 | +0.110 |
| *first-order, motor* |  |  |  |  |  |  |  |  |  |  |  |
| Ventral lateral posterior, dorsal (VLpd) | paralattice | paralattice | cerebellar | +1.61 | +1.33 | +0.76 | +0.16 | +0.085 | +0.090 | +0.101 | +0.122 |
| Ventral lateral posterior, ventral (VLpv) | paralattice | paralattice | cerebellar | +1.06 | +0.44 | −0.79 | −1.20 | +0.093 | +0.076 | +0.111 | +0.105 |
| *first-order, limbic* |  |  |  |  |  |  |  |  |  |  |  |
| Anteroventral (AV) | paralattice | paralattice | none | +1.73 | +1.49 | +1.38 | +0.96 | +0.052 | +0.080 | +0.077 | +0.112 |
| Anteromedial (AM) | paralattice | paralattice | none | +0.97 | +1.07 | +1.30 | +1.39 | +0.034 | +0.050 | +0.036 | +0.048 |
| Lateral dorsal (LD) | paralattice | paralattice | none | +1.16 | +1.13 | +1.11 | +1.17 | +0.014 | +0.017 | +0.057 | +0.061 |
| *higher-order* |  |  |  |  |  |  |  |  |  |  |  |
| Mediodorsal, parvocellular (MDpc) | paralattice | paralattice | SNr | +0.71 | +0.32 | +0.22 | −0.76 | +0.019 | +0.014 | +0.076 | +0.082 |
| *intralaminar, rostral* |  |  |  |  |  |  |  |  |  |  |  |
| Central lateral (CL) | paralattice | paralattice | cerebellar | +0.97 | +0.85 | +0.29 | −0.26 | +0.023 | +0.011 | +0.095 | +0.099 |
| Central medial (CeM) | paralattice | paralattice | none | +0.72 | +1.16 | +1.02 | +1.01 | +0.026 | +0.032 | +0.024 | +0.035 |
| *intralaminar, caudal* |  |  |  |  |  |  |  |  |  |  |  |
| Parafascicular (Pf) | paralattice | paralattice | GPi/SNr | +0.41 | +0.61 | −0.77 | −1.18 | +0.042 | +0.041 | +0.056 | +0.072 |
| **Peri-allocortex** |  |  |  |  |  |  |  |  |  |  |  |
| *higher-order* |  |  |  |  |  |  |  |  |  |  |  |
| Mediodorsal, magnocellular (MDmc) | peri-allocortex | peri-allocortex | SNr | +0.23 | −0.18 | +1.86 | +1.47 | −0.011 | −0.029 | −0.002 | −0.030 |
| *midline* |  |  |  |  |  |  |  |  |  |  |  |
| Medioventral (reuniens) (MV) | peri-allocortex | peri-allocortex | – | +0.41 | +0.66 | +1.11 | +1.18 | +0.009 | +0.005 | −0.006 | +0.003 |
| **Unresolved** |  |  |  |  |  |  |  |  |  |  |  |
| *basal-ganglia-recipient, diffuse (layer 1)* |  |  |  |  |  |  |  |  |  |  |  |
| Ventral medial (VM) | unresolved | unresolved | SNr | +0.16 | +0.04 | −1.03 | −1.37 | +0.039 | +0.033 | +0.041 | +0.049 |

**Table S5.**
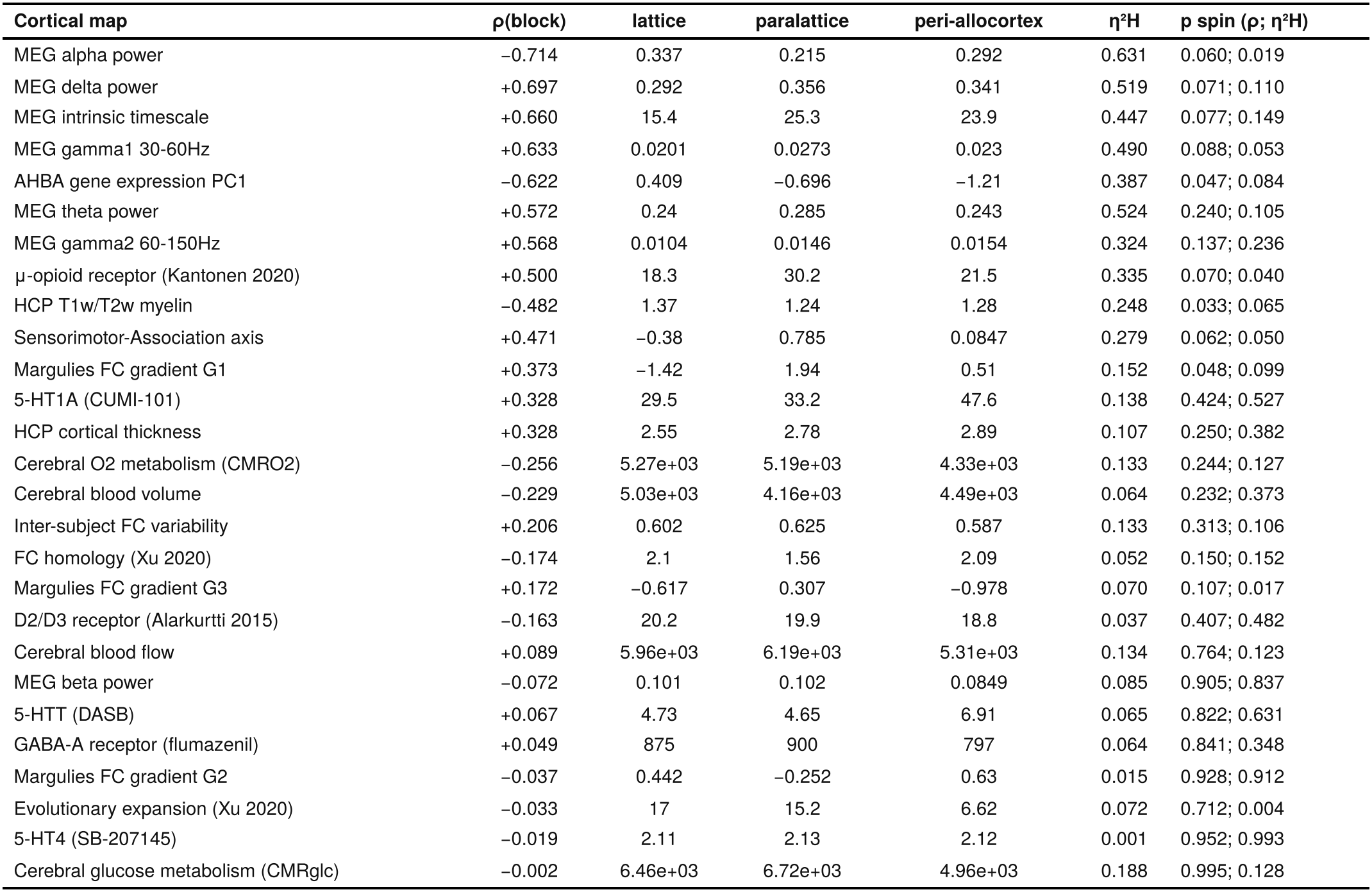
Descriptive correspondence of the blocks with published cortical maps (HCP 7T, N = 151). For each map, ρ(block) is the Spearman correlation between its per-vertex value and the block ordinal index (lattice = 1, paralattice = 2, peri-allocortex = 3); a positive value denotes a property that increases from lattice through paralattice to the peri-allocortex, a negative value one that decreases. The three block columns give each map’s mean value within that block, in the map’s own units, and η^2^H is the Kruskal–Wallis effect size, (H − k + 1)/(n − k) with k = 3 blocks^58^. The two are reported beside ρ because ρ is a monotonic summary and cannot see a map that peaks on the middle block: cerebral glucose metabolism, for instance, reads ρ = 0.00 on block means of 6.46 / 6.72 / 4.96 × 10^3^ and η^2^H = 0.188, and the same holds for functional connectivity gradient G3 and functional connectivity homology. A small ρ beside a large η^2^H therefore means the blocks differ but not in rank order. Maps are from the neuromaps library and are ordered by |ρ|. The sensorimotor–association axis is each vertex’s mean rank across ten maps^18^, among them T1w/T2w myelin, the principal functional gradient, gene expression PC1 and cortical thickness, and is not independent of them. The two Xu 2020 maps are from Xu et al.^59^; the expansion maps of Hill et al.^19^ are delivered for the right hemisphere only and were not used.^60^ The maps are spatially smooth and strongly inter-correlated and are not independent. The last column gives, for ρ and for η^2^H, the fraction of 1,000 rotations of the block map on the sphere reaching a value at least as large (the spin test, Methods; 0.001 is the smallest value attainable with 1,000 rotations). Eight of the 54 tests fall below 0.05 and none survives correction across the 27 maps (smallest Benjamini–Hochberg q = 0.26 for ρ and 0.11 for η^2^H; with 1,000 rotations these values vary between sets of rotations, from 0.22 to 0.26 for ρ and from 0.11 to 0.19 for η^2^H over 12 sets, but none falls below 0.05). The table should thus be considered as descriptive of the direction in which each property varies across the blocks.

| Cortical map | ρ(block) | lattice | paralattice | peri-allocortex | η<sup>2</sup>H | p spin (ρ; η<sup>2</sup>H) |
|---|---|---|---|---|---|---|
| MEG alpha power | −0.714 | 0.337 | 0.215 | 0.292 | 0.631 | 0.060; 0.019 |
| MEG delta power | +0.697 | 0.292 | 0.356 | 0.341 | 0.519 | 0.071; 0.110 |
| MEG intrinsic timescale | +0.660 | 15.4 | 25.3 | 23.9 | 0.447 | 0.077; 0.149 |
| MEG gamma1 30-60Hz | +0.633 | 0.0201 | 0.0273 | 0.023 | 0.490 | 0.088; 0.053 |
| AHBA gene expression PC1 | −0.622 | 0.409 | −0.696 | −1.21 | 0.387 | 0.047; 0.084 |
| MEG theta power | +0.572 | 0.24 | 0.285 | 0.243 | 0.524 | 0.240; 0.105 |
| MEG gamma2 60-150Hz | +0.568 | 0.0104 | 0.0146 | 0.0154 | 0.324 | 0.137; 0.236 |
| μ-opioid receptor (Kantonen 2020) | +0.500 | 18.3 | 30.2 | 21.5 | 0.335 | 0.070; 0.040 |
| HCP T1w/T2w myelin | −0.482 | 1.37 | 1.24 | 1.28 | 0.248 | 0.033; 0.065 |
| Sensorimotor-Association axis | +0.471 | −0.38 | 0.785 | 0.0847 | 0.279 | 0.062; 0.050 |
| Margulies FC gradient G1 | +0.373 | −1.42 | 1.94 | 0.51 | 0.152 | 0.048; 0.099 |
| 5-HT1A (CUMI-101) | +0.328 | 29.5 | 33.2 | 47.6 | 0.138 | 0.424; 0.527 |
| HCP cortical thickness | +0.328 | 2.55 | 2.78 | 2.89 | 0.107 | 0.250; 0.382 |
| Cerebral O2 metabolism (CMRO2) | −0.256 | 5.27e+03 | 5.19e+03 | 4.33e+03 | 0.133 | 0.244; 0.127 |
| Cerebral blood volume | −0.229 | 5.03e+03 | 4.16e+03 | 4.49e+03 | 0.064 | 0.232; 0.373 |
| Inter-subject FC variability | +0.206 | 0.602 | 0.625 | 0.587 | 0.133 | 0.313; 0.106 |
| FC homology (Xu 2020) | −0.174 | 2.1 | 1.56 | 2.09 | 0.052 | 0.150; 0.152 |
| Margulies FC gradient G3 | +0.172 | −0.617 | 0.307 | −0.978 | 0.070 | 0.107; 0.017 |
| D2/D3 receptor (Alarkurtti 2015) | −0.163 | 20.2 | 19.9 | 18.8 | 0.037 | 0.407; 0.482 |
| Cerebral blood flow | +0.089 | 5.96e+03 | 6.19e+03 | 5.31e+03 | 0.134 | 0.764; 0.123 |
| MEG beta power | −0.072 | 0.101 | 0.102 | 0.0849 | 0.085 | 0.905; 0.837 |
| 5-HTT (DASB) | +0.067 | 4.73 | 4.65 | 6.91 | 0.065 | 0.822; 0.631 |
| GABA-A receptor (flumazenil) | +0.049 | 875 | 900 | 797 | 0.064 | 0.841; 0.348 |
| Margulies FC gradient G2 | −0.037 | 0.442 | −0.252 | 0.63 | 0.015 | 0.928; 0.912 |
| Evolutionary expansion (Xu 2020) | −0.033 | 17 | 15.2 | 6.62 | 0.072 | 0.712; 0.004 |
| 5-HT4 (SB-207145) | −0.019 | 2.11 | 2.13 | 2.12 | 0.001 | 0.952; 0.993 |
| Cerebral glucose metabolism (CMRglc) | −0.002 | 6.46e+03 | 6.72e+03 | 4.96e+03 | 0.188 | 0.995; 0.128 |

**Table S6.** Robustness of the main partition to analysis choices (HCP 7T, N = 151). Each row re-derives the partition of each hemisphere with one element of the construction changed and scores it against the partition of record (peri-allocortex set aside; a null floor for each subject at q = 0.05; an edge in a majority of subjects with weight the cohort mean of r; 3 ≤ d < 20 mm; normalized-cut sweep and descent) by the adjusted Rand index (ARI; chance = 0, identity = 1); block sizes are lattice / paralattice. Rows: the window (3–18, 3–15, 3–12, 5–20, 7–20, 8–20 and 10–20 mm); the support rule (one third, two thirds, 90% of subjects); the sweep criterion (conductance); two halves of the cohort carried through the whole chain independently; the weight convention; the single cohort-wide threshold that retains the same number of edges; a stricter per-subject floor (local false-discovery rate 0.1); all pairs of the window, without floor or support rule; the partition without the area weights; the women and the men of the cohort, each group carried through the whole chain on its own; the 7T cohort with global-signal regression; the 7T cohort with the peri-allocortex left in (its vertices are then counted in the blocks); and the replications (movie, 3T and the PAN cohort, the latter also without multi-echo ICA, without it and with 12 motion regressors, and with global-signal regression), each scored against the partition of record.

| Construction | ARI, left | ARI, right | Blocks, left | Blocks, right |
|---|---|---|---|---|
| Partition of record (3–20 mm, majority support, null floor at q = 0.05, no GSR) | 1.000 | 1.000 | 18,952 / 9,003 | 18,966 / 8,955 |
| Support: one third of subjects | 0.957 | 0.968 | 18,771 / 9,184 | 19,076 / 8,845 |
| Support: two thirds of subjects | 0.971 | 0.987 | 18,743 / 9,136 | 18,847 / 8,997 |
| Support: 90% of subjects | 0.716 | 0.934 | 16,302 / 10,456 | 17,713 / 8,731 |
| Window 5–20 mm | 0.961 | 0.962 | 17,848 / 8,611 | 17,772 / 8,538 |
| Window 7–20 mm | 0.978 | 0.984 | 17,013 / 8,117 | 16,854 / 7,991 |
| Window 8–20 mm | 0.981 | 0.985 | 16,820 / 8,027 | 16,614 / 7,912 |
| Window 10–20 mm | 0.980 | 0.985 | 16,559 / 7,780 | 16,395 / 7,730 |
| Window 3–15 mm | 0.996 | 0.985 | 18,948 / 9,007 | 19,039 / 8,882 |
| Window 3–18 mm | 0.999 | 0.987 | 18,949 / 9,006 | 19,047 / 8,874 |
| Window 3–12 mm | 0.994 | 0.980 | 18,940 / 9,015 | 19,005 / 8,916 |
| Sweep criterion: conductance | 1.000 | 1.000 | 18,952 / 9,003 | 18,966 / 8,955 |
| Half A of the cohort (76 subjects) | 0.982 | 0.990 | 18,883 / 9,067 | 19,003 / 8,918 |
| Half B of the cohort (75 subjects) | 0.994 | 0.993 | 18,922 / 9,033 | 18,946 / 8,975 |
| Weight: mean of (r − τ<sub>s</sub>) over supporting subjects | 0.835 | 0.999 | 17,756 / 10,199 | 18,967 / 8,954 |
| Weight: mean of max(r − τ<sub>s</sub>, 0) over all subjects | 0.834 | 0.995 | 17,753 / 10,202 | 18,940 / 8,981 |
| Weight: mean of (r − τs) over all subjects | 0.983 | 0.993 | 18,864 / 9,091 | 18,969 / 8,952 |
| Single cohort-wide threshold on mean r retaining the same number of edges (r = 0.126) | 0.995 | 0.995 | 18,983 / 8,972 | 18,980 / 8,941 |
| Stricter per-subject floor: local false-discovery rate 0.1 (median τs 0.149) | 0.971 | 0.974 | 18,805 / 9,105 | 18,798 / 9,075 |
| All pairs of the window (no floor, no support rule; weight max(mean r, 10<sup>−6</sup>)) | 0.955 | 0.949 | 19,213 / 8,742 | 19,253 / 8,668 |
| Without the area weights | 0.968 | 0.967 | 19,065 / 8,890 | 18,951 / 8,970 |
| Women only (N = 89) | 0.979 | 0.991 | 18,833 / 9,122 | 18,927 / 8,994 |
| Men only (N = 62) | 0.987 | 0.989 | 18,985 / 8,955 | 19,010 / 8,897 |
| With global-signal regression | 0.993 | 0.988 | 18,973 / 8,982 | 18,945 / 8,976 |
| Peri-allocortex not masked | 0.987 | 0.998 | 20,282 / 9,414 | 20,408 / 9,308 |
| HCP 7T movie, raw (N = 160) | 0.975 | 0.967 | 18,842 / 9,113 | 18,951 / 8,970 |
| HCP 7T movie, stimulus-residualized (N = 160) | 0.980 | 0.972 | 18,950 / 9,005 | 19,077 / 8,844 |
| HCP 3T (N = 349) | 0.906 | 0.886 | 19,443 / 8,512 | 19,732 / 8,189 |
| PAN cohort (N = 10) | 0.822 | 0.856 | 19,998 / 7,363 | 19,457 / 8,001 |
| PAN cohort (N = 10), without multi-echo ICA | 0.915 | 0.864 | 19,098 / 8,713 | 19,763 / 8,006 |
| PAN cohort (N = 10), without multi-echo ICA, with 12 motion regressors | 0.820 | 0.849 | 20,163 / 7,661 | 19,903 / 7,883 |
| PAN cohort (N = 10), with global-signal regression | 0.921 | 0.880 | 19,503 / 8,452 | 19,351 / 8,570 |

**Table S7.** Block composition of the Cole–Anticevic networks in every cohort. For each network, the percentage of its vertices assigned to lattice/paralattice/peri-allocortex, in the same six constructions as Table S2; vertices that are not assigned are not counted, so the PAN percentages can sum to less than 100. Vertex counts per network (HCP 7T rest): Visual 1 2,148, Visual 2 6,787, Auditory 1,633, Somatomotor 10,107, Dorsal attention 3,807, Posterior multimodal 954, Language 3,231, Cingulo-opercular 9,102, Frontoparietal 8,222, Default 11,537, Orbito-affective 718, Ventral multimodal 1,166.

| **Network** | **HCP 7T rest** | **Movie, raw** | **Movie, residual** | **HCP 3T** | **PAN cohort** | **PAN04** |
|---|---|---|---|---|---|---|
| Visual 1 | 100.0/0.0/0.0 | 100.0/0.0/0.0 | 100.0/0.0/0.0 | 100.0/0.0/0.0 | 99.5/0.0/0.0 | 99.3/0.0/0.0 |
| Visual 2 | 100.0/0.0/0.0 | 100.0/0.0/0.0 | 100.0/0.0/0.0 | 100.0/0.0/0.0 | 99.4/0.0/0.0 | 99.7/0.0/0.0 |
| Auditory | 100.0/0.0/0.0 | 100.0/0.0/0.0 | 100.0/0.0/0.0 | 100.0/0.0/0.0 | 99.1/0.0/0.0 | 98.7/0.0/0.0 |
| Somatomotor | 88.1/11.9/0.0 | 86.9/13.1/0.0 | 88.6/11.4/0.0 | 91.6/8.4/0.0 | 94.0/5.8/0.0 | 92.7/7.3/0.0 |
| Dorsal attention | 74.5/20.1/5.4 | 74.4/20.3/5.4 | 74.4/20.2/5.4 | 76.4/18.2/5.4 | 80.1/14.4/5.4 | 77.3/17.4/5.4 |
| Posterior multimodal | 100.0/0.0/0.0 | 100.0/0.0/0.0 | 100.0/0.0/0.0 | 100.0/0.0/0.0 | 100.0/0.0/0.0 | 100.0/0.0/0.0 |
| Language | 60.4/39.6/0.0 | 60.3/39.7/0.0 | 60.5/39.5/0.0 | 65.0/35.0/0.0 | 63.8/35.3/0.0 | 60.1/39.2/0.0 |
| Cingulo-opercular | 40.7/58.7/0.6 | 40.7/58.7/0.6 | 41.3/58.1/0.6 | 48.1/51.3/0.6 | 51.0/48.2/0.6 | 44.6/54.6/0.6 |
| Frontoparietal | 43.4/55.7/0.9 | 43.4/55.7/0.9 | 43.4/55.7/0.9 | 43.4/55.7/0.9 | 43.5/52.8/0.9 | 43.3/54.2/0.9 |
| Default | 47.1/39.8/13.1 | 47.1/39.8/13.1 | 47.1/39.8/13.1 | 47.1/39.8/13.1 | 44.8/36.8/13.1 | 46.0/38.6/13.1 |
| Orbito-affective | 0.0/27.3/72.7 | 0.0/27.3/72.7 | 0.0/27.3/72.7 | 0.0/27.3/72.7 | 0.0/15.6/72.7 | 0.0/25.8/72.7 |
| Ventral multimodal | 0.0/0.0/100.0 | 0.0/0.0/100.0 | 0.0/0.0/100.0 | 0.0/0.0/100.0 | 0.0/0.0/100.0 | 0.0/0.0/100.0 |

**Table S8.**
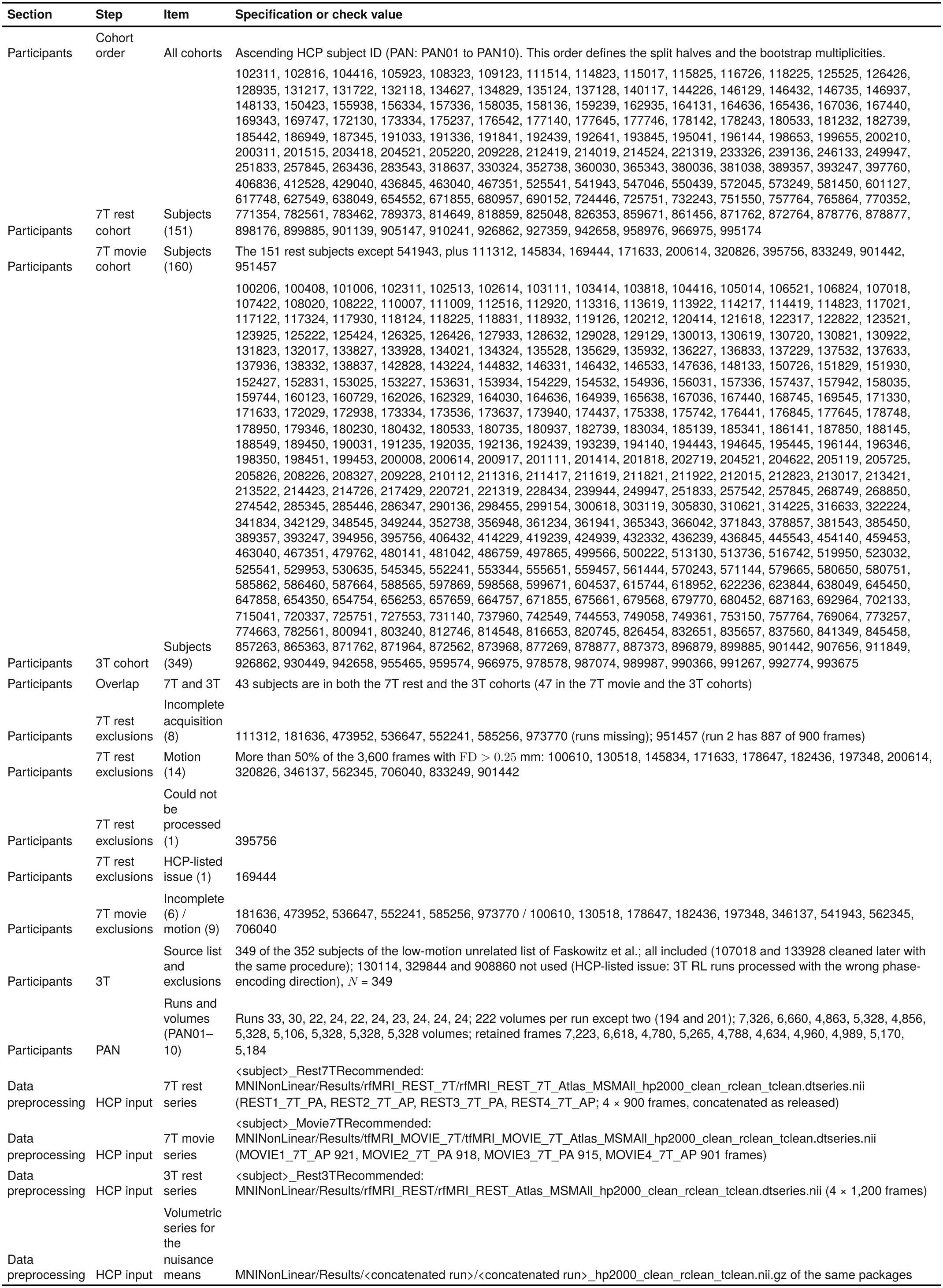
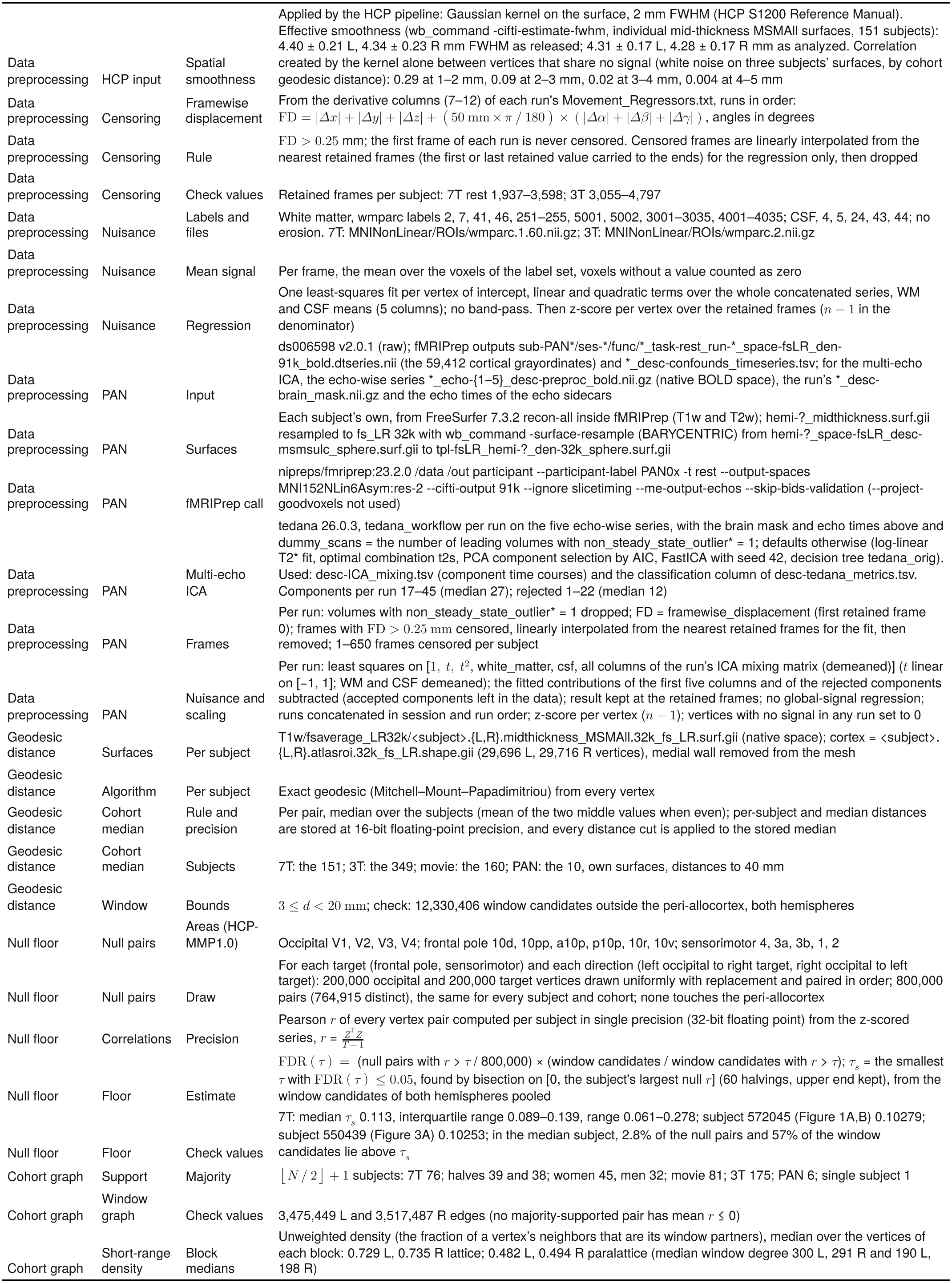
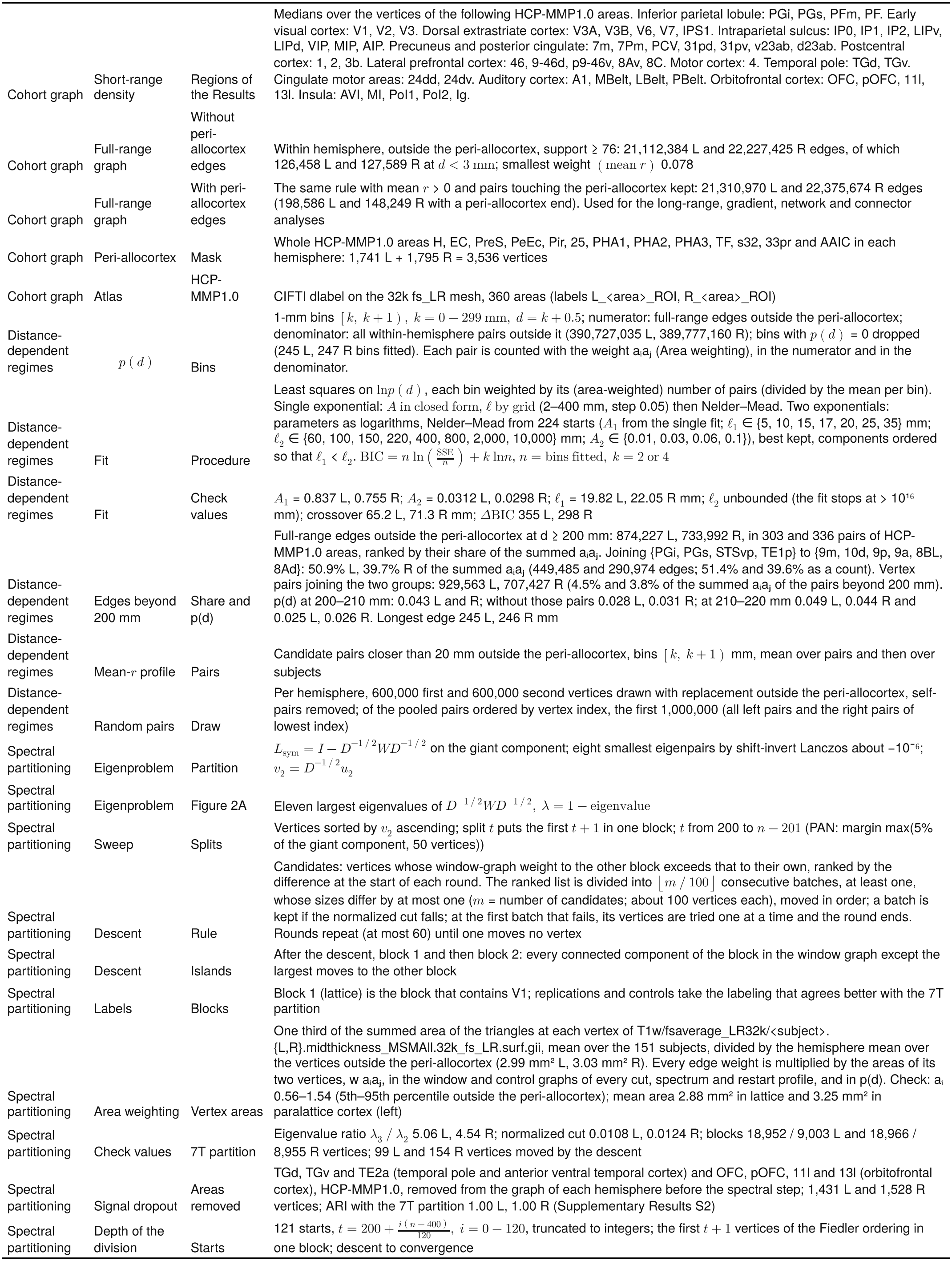
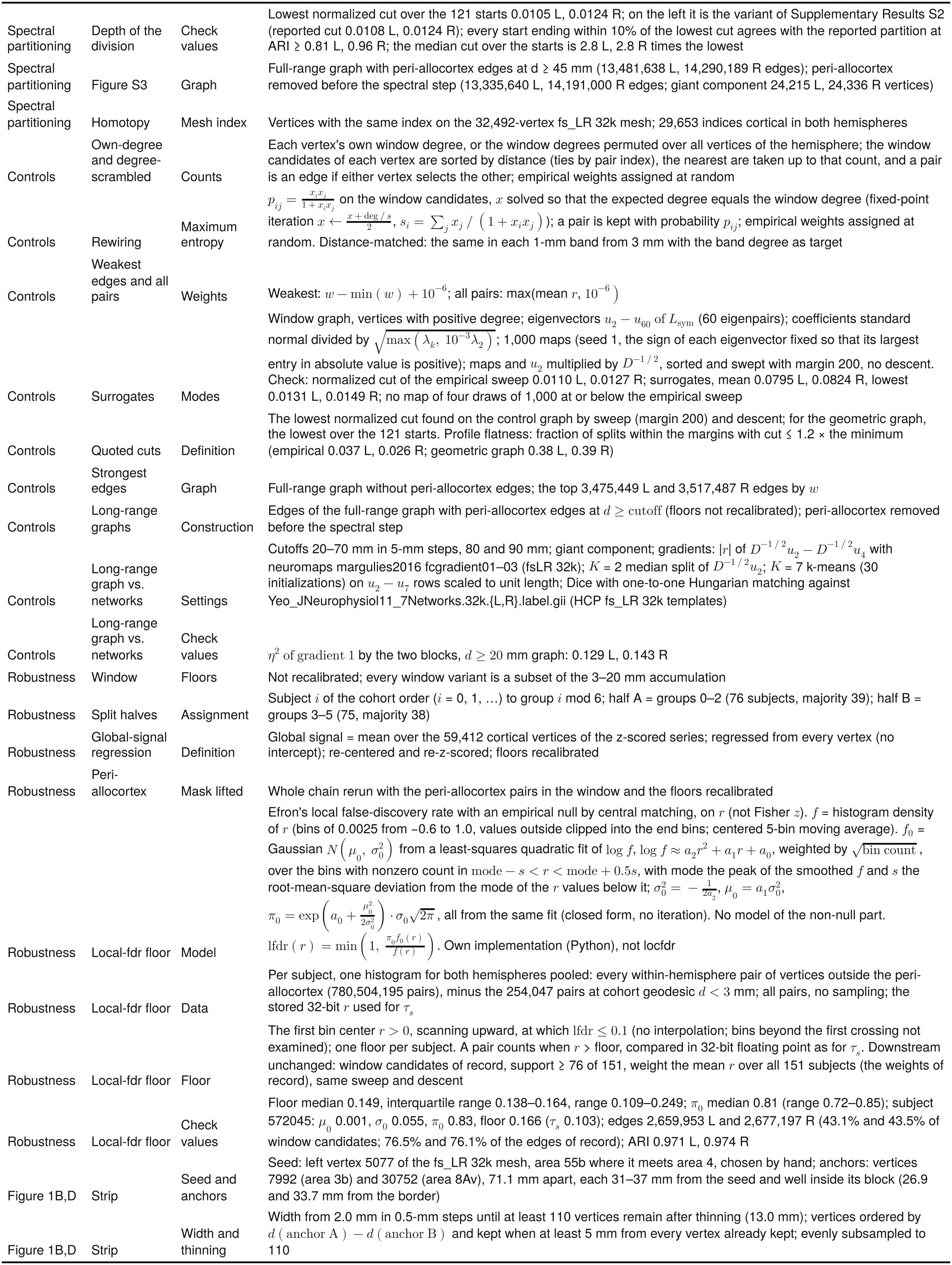
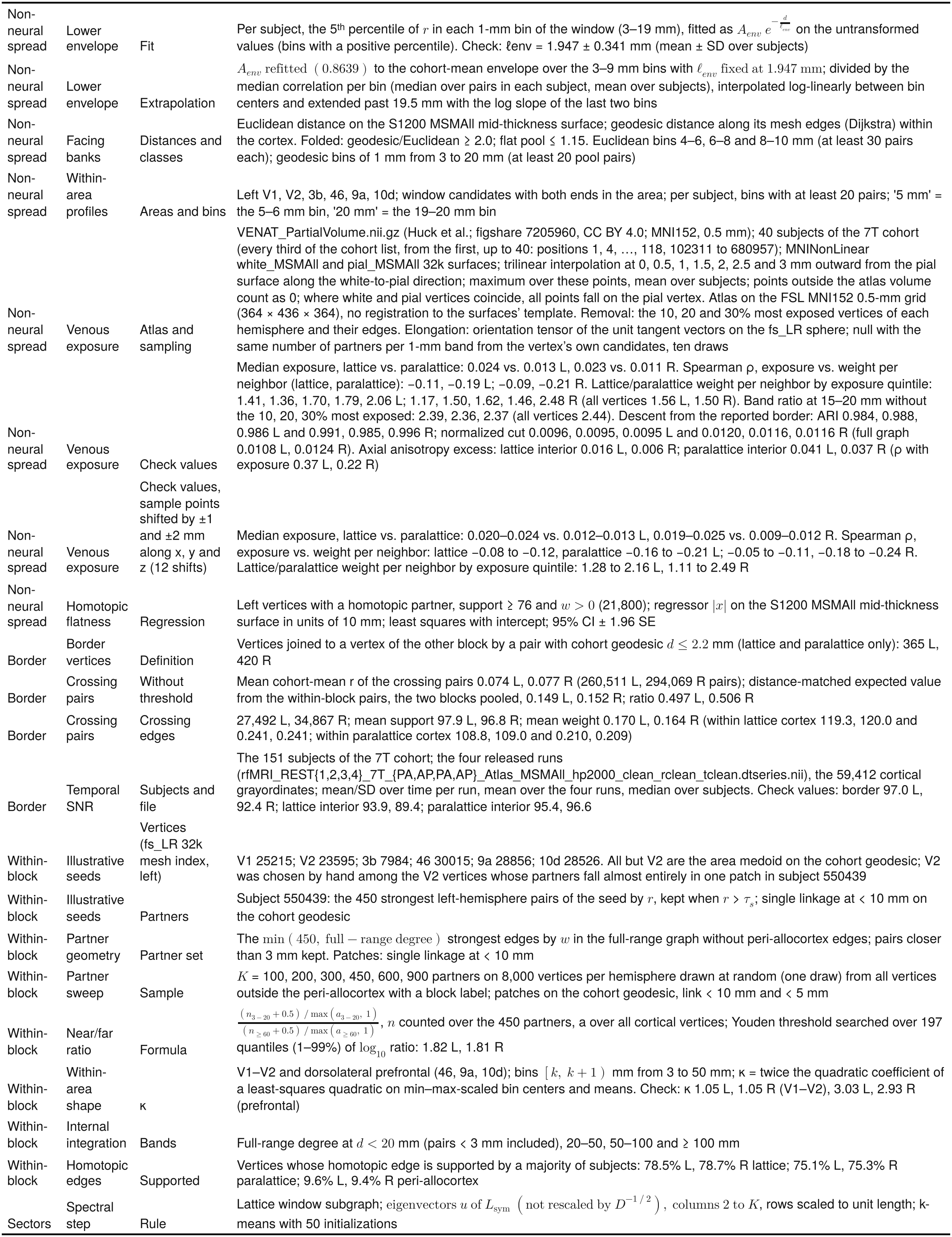
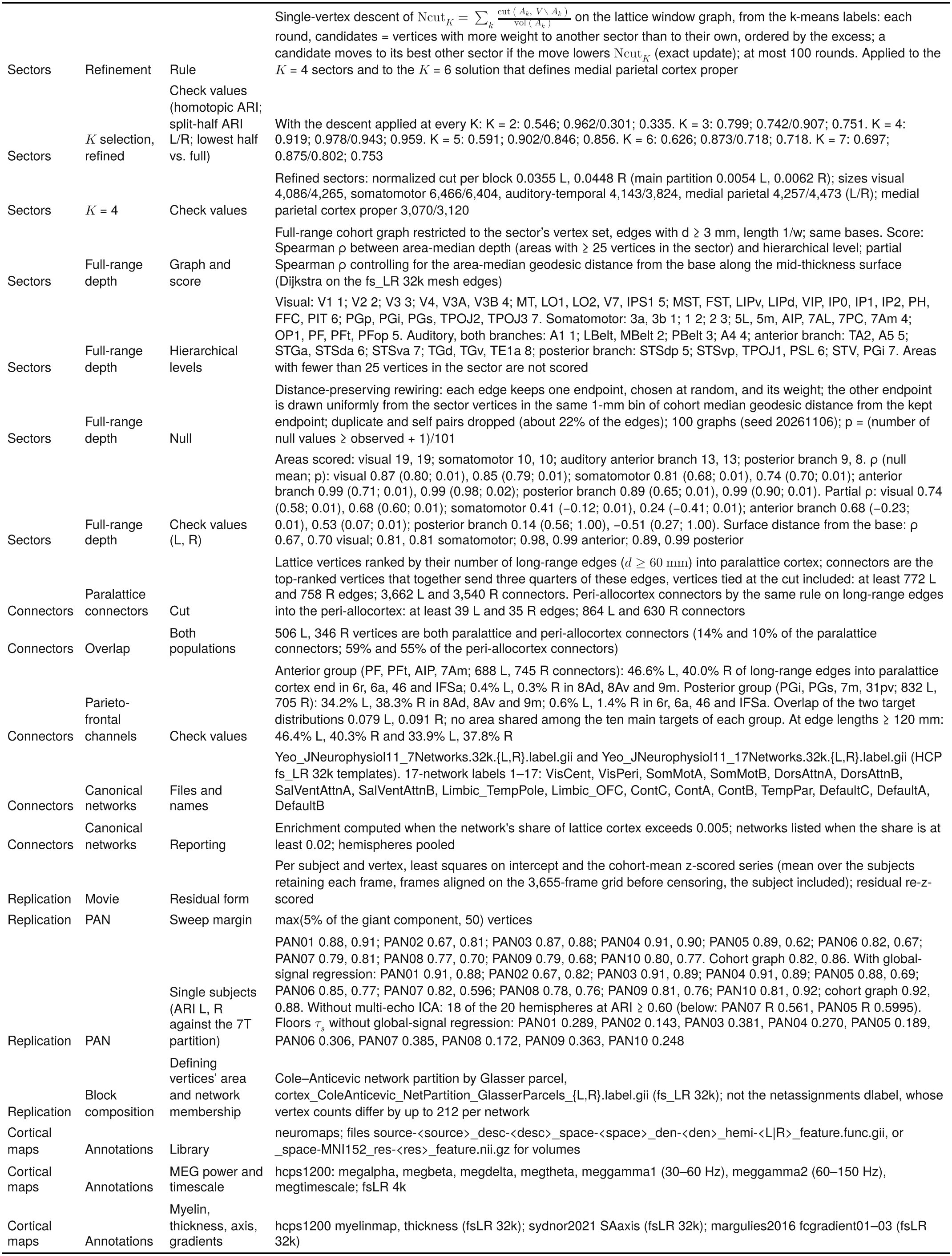
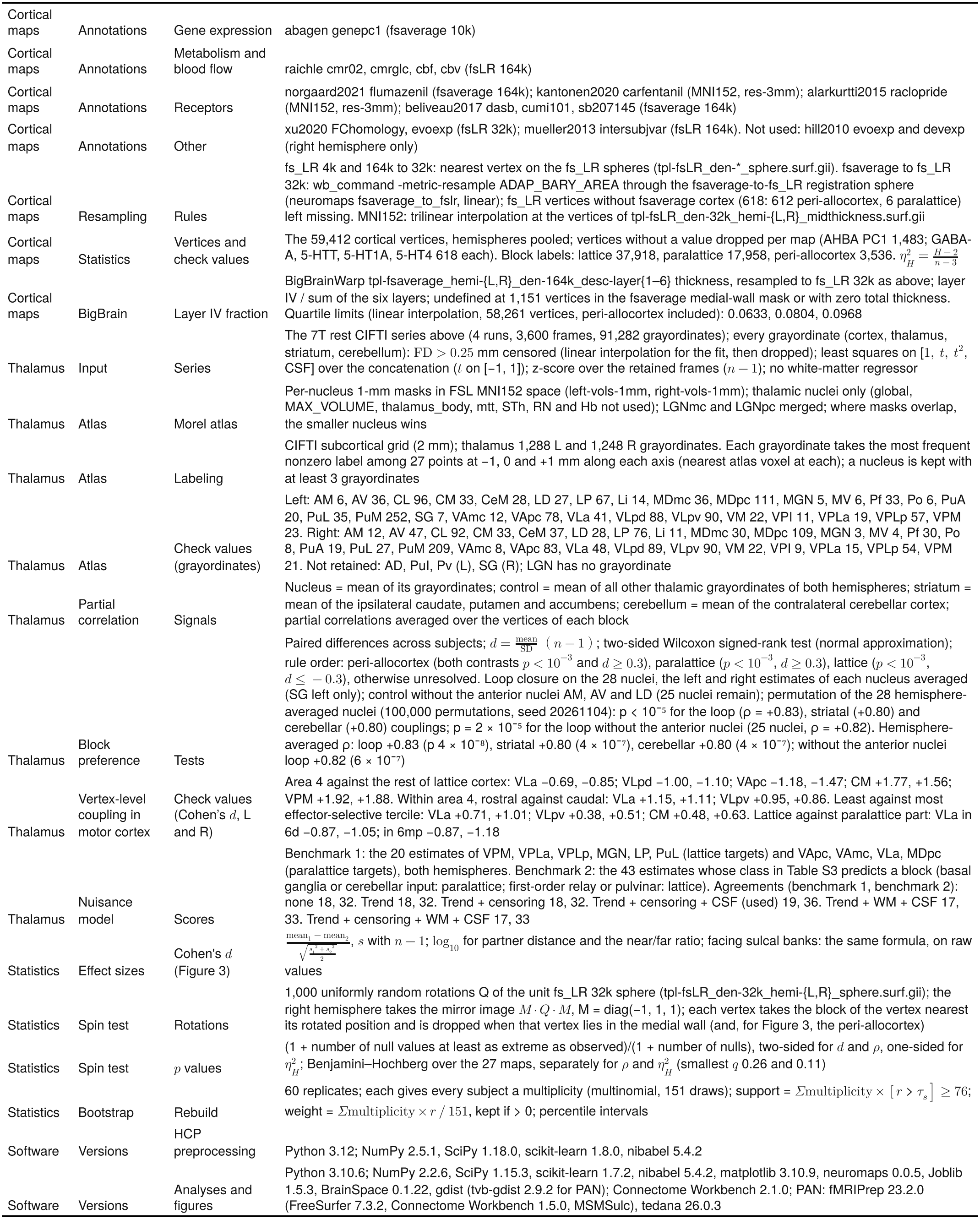
Reproduction details. The inputs from the public releases, the definitions and conventions that the Methods give in outline, and check values against which an independent implementation can be compared, in the order of the Methods. Subject lists are given in this table and, with the reason for each exclusion, in Data S1.

| Section | Step | Item | Specification or check value |
|---|---|---|---|
| Participants | Cohort order | All cohorts | Ascending HCP subject ID (PAN: PAN01 to PAN10). This order defines the split halves and the bootstrap multiplicities. |
| Participants | 7T rest cohort | Subjects (151) | 102311, 102816, 104416, 105923, 108323, 109123, 111514, 114823, 115017, 115825, 116726, 118225, 125525, 126426, 128935, 131217, 131722, 132118, 134627, 134829, 135124, 137128, 140117, 144226, 146129, 146432, 146735, 146937, 148133, 150423, 155938, 156334, 157336, 158035, 158136, 159239, 162935, 164131, 164636, 165436, 167036, 167440, 169343, 169747, 172130, 173334, 175237, 176542, 177140, 177645, 177746, 178142, 178243, 180533, 181232, 182739, 185442, 186949, 187345, 191033, 191336, 191841, 192439, 192641, 193845, 195041, 196144, 198653, 199655, 200210, 200311, 201515, 203418, 204521, 205220, 209228, 212419, 214019, 214524, 221319, 233326, 239136, 246133, 249947, 251833, 257845, 263436, 283543, 318637, 330324, 352738, 360030, 365343, 380036, 381038, 389357, 393247, 397760, 406836, 412528, 429040, 436845, 463040, 467351, 525541, 541943, 547046, 550439, 572045, 573249, 581450, 601127, 617748, 627549, 638049, 654552, 671855, 680957, 690152, 724446, 725751, 732243, 751550, 757764, 765864, 770352, 771354, 782561, 783462, 789373, 814649, 818859, 825048, 826353, 859671, 861456, 871762, 872764, 878776, 878877, 898176, 899885, 901139, 905147, 910241, 926862, 927359, 942658, 958976, 966975, 995174 |
| Participants | 7T movie cohort | Subjects (160) | The 151 rest subjects except 541943, plus 111312, 145834, 169444, 171633, 200614, 320826, 395756, 833249, 901442, 951457 |
| Participants | 3T cohort | Subjects (349) | 100206, 100408, 101006, 102311, 102513, 102614, 103111, 103414, 103818, 104416, 105014, 106521, 106824, 107018, 107422, 108020, 108222, 110007, 111009, 112516, 112920, 113316, 113619, 113922, 114217, 114419, 114823, 117021, 117122, 117324, 117930, 118124, 118225, 118831, 118932, 119126, 120212, 120414, 121618, 122317, 122822, 123521, 123925, 125222, 125424, 126325, 126426, 127933, 128632, 129028, 129129, 130013, 130619, 130720, 130821, 130922, 131823, 132017, 133827, 133928, 134021, 134324, 135528, 135629, 135932, 136227, 136833, 137229, 137532, 137633, 137936, 138332, 138837, 142828, 143224, 144832, 146331, 146432, 146533, 147636, 148133, 150726, 151829, 151930, 152427, 152831, 153025, 153227, 153631, 153934, 154229, 154532, 154936, 156031, 157336, 157437, 157942, 158035, 159744, 160123, 160729, 162026, 162329, 164030, 164636, 164939, 165638, 167036, 167440, 168745, 169545, 171330, 171633, 172029, 172938, 173334, 173536, 173637, 173940, 174437, 175338, 175742, 176441, 176845, 177645, 178748, 178950, 179346, 180230, 180432, 180533, 180735, 180937, 182739, 183034, 185139, 185341, 186141, 187850, 188145, 188549, 189450, 190031, 191235, 192035, 192136, 192439, 193239, 194140, 194443, 194645, 195445, 196144, 196346, 198350, 198451, 199453, 200008, 200614, 200917, 201111, 201414, 201818, 202719, 204521, 204622, 205119, 205725, 205826, 208226, 208327, 209228, 210112, 211316, 211417, 211619, 211821, 211922, 212015, 212823, 213017, 213421, 213522, 214423, 214726, 217429, 220721, 221319, 228434, 239944, 249947, 251833, 257542, 257845, 268749, 268850, 274542, 285345, 285446, 286347, 290136, 298455, 299154, 300618, 303119, 305830, 310621, 314225, 316633, 322224, 341834, 342129, 348545, 349244, 352738, 356948, 361234, 361941, 365343, 366042, 371843, 378857, 381543, 385450, 389357, 393247, 394956, 395756, 406432, 414229, 419239, 424939, 432332, 436239, 436845, 445543, 454140, 459453, 463040, 467351, 479762, 480141, 481042, 486759, 497865, 499566, 500222, 513130, 513736, 516742, 519950, 523032, 525541, 529953, 530635, 545345, 552241, 553344, 555651, 559457, 561444, 570243, 571144, 579665, 580650, 580751, 585862, 586460, 587664, 588565, 597869, 598568, 599671, 604537, 615744, 618952, 622236, 623844, 638049, 645450, 647858, 654350, 654754, 656253, 657659, 664757, 671855, 675661, 679568, 679770, 680452, 687163, 692964, 702133, 715041, 720337, 725751, 727553, 731140, 737960, 742549, 744553, 749058, 749361, 753150, 757764, 769064, 773257, 774663, 782561, 800941, 803240, 812746, 814548, 816653, 820745, 826454, 832651, 835657, 837560, 841349, 845458, 857263, 865363, 871762, 871964, 872562, 873968, 877269, 878877, 887373, 896879, 899885, 901442, 907656, 911849, 926862, 930449, 942658, 955465, 959574, 966975, 978578, 987074, 989987, 990366, 991267, 992774, 993675 |
| Participants | Overlap | 7T and 3T | 43 subjects are in both the 7T rest and the 3T cohorts (47 in the 7T movie and the 3T cohorts) |
| Participants | 7T rest exclusions | Incomplete acquisition (8) | 111312, 181636, 473952, 536647, 552241, 585256, 973770 (runs missing); 951457 (run 2 has 887 of 900 frames) |
| Participants | 7T rest exclusions | Motion (14) | More than 50% of the 3,600 frames with FD > 0.25 mm: 100610, 130518, 145834, 171633, 178647, 182436, 197348, 200614, 320826, 346137, 562345, 706040, 833249, 901442 |
| Participants | 7T rest exclusions | Could not be processed (1) | 395756 |
| Participants | 7T rest exclusions | HCP-listed issue (1) | 169444 |
| Participants | 7T movie exclusions | Incomplete (6) / motion (9) | 181636, 473952, 536647, 552241, 585256, 973770 / 100610, 130518, 178647, 182436, 197348, 346137, 541943, 562345, 706040 |
| Participants | 3T | Source list and exclusions | 349 of the 352 subjects of the low-motion unrelated list of Faskowitz et al.; all included (107018 and 133928 cleaned later with the same procedure); 130114, 329844 and 908860 not used (HCP-listed issue: 3T RL runs processed with the wrong phase-encoding direction), N = 349 |
| Participants | PAN | Runs and volumes (PAN01–10) | Runs 33, 30, 22, 24, 22, 24, 23, 24, 24, 24; 222 volumes per run except two (194 and 201); 7,326, 6,660, 4,863, 5,328, 4,856, 5,328, 5,106, 5,328, 5,328, 5,328 volumes; retained frames 7,223, 6,618, 4,780, 5,265, 4,788, 4,634, 4,960, 4,989, 5,170, 5,184 |
| Data preprocessing | HCP input | 7T rest series | <subject>_Rest7TRecommended: MNINonLinear/Results/rfMRI_REST_7T/rfMRI_REST_7T_Atlas_MSMAll_hp2000_clean_rclean_tclean.dtseries.nii (REST1_7T_PA, REST2_7T_AP, REST3_7T_PA, REST4_7T_AP; 4 × 900 frames, concatenated as released) |
| Data preprocessing | HCP input | 7T movie series | <subject>_Movie7TRecommended: MNINonLinear/Results/tfMRI_MOVIE_7T/tfMRI_MOVIE_7T_Atlas_MSMAll_hp2000_clean_rclean_tclean.dtseries.nii (MOVIE1_7T_AP 921, MOVIE2_7T_PA 918, MOVIE3_7T_PA 915, MOVIE4_7T_AP 901 frames) |
| Data preprocessing | HCP input | 3T rest series | <subject>_Rest3TRecommended: MNINonLinear/Results/rfMRI_REST/rfMRI_REST_Atlas_MSMAll_hp2000_clean_rclean_tclean.dtseries.nii (4 × 1,200 frames) |
| Data preprocessing | HCP input | Volumetric series for the nuisance means | MNINonLinear/Results/<concatenated run>/<concatenated run>_hp2000_clean_rclean_tclean.nii.gz of the same packages |
| Data preprocessing | HCP input | Spatial smoothness | Applied by the HCP pipeline: Gaussian kernel on the surface, 2 mm FWHM (HCP S1200 Reference Manual). Effective smoothness (wb_command -cifti-estimate-fwhm, individual mid-thickness MSMAll surfaces, 151 subjects): 4.40 ± 0.21 L, 4.34 ± 0.23 R mm FWHM as released; 4.31 ± 0.17 L, 4.28 ± 0.17 R mm as analyzed. Correlation created by the kernel alone between vertices that share no signal (white noise on three subjects' surfaces, by cohort geodesic distance): 0.29 at 1–2 mm, 0.09 at 2–3 mm, 0.02 at 3–4 mm, 0.004 at 4–5 mm |
| Data preprocessing | Censoring | Framewise displacement | From the derivative columns (7–12) of each run's Movement_Regressors.txt, runs in order: FD = \|Δx\| + \|Δy\| + \|Δz\| + ( 50 mm × π / 180 ) × ( \|Δα\| + \|Δβ\| + \|Δγ\| ) , angles in degrees |
| Data preprocessing | Censoring | Rule | FD > 0.25 mm; the first frame of each run is never censored. Censored frames are linearly interpolated from the nearest retained frames (the first or last retained value carried to the ends) for the regression only, then dropped |
| Data preprocessing | Censoring | Check values | Retained frames per subject: 7T rest 1,937–3,598; 3T 3,055–4,797 |
| Data preprocessing | Nuisance | Labels and files | White matter, wmparc labels 2, 7, 41, 46, 251–255, 5001, 5002, 3001–3035, 4001–4035; CSF, 4, 5, 24, 43, 44; no erosion. 7T: MNINonLinear/ROIs/wmparc.1.60.nii.gz; 3T: MNINonLinear/ROIs/wmparc.2.nii.gz |
| Data preprocessing | Nuisance | Mean signal | Per frame, the mean over the voxels of the label set, voxels without a value counted as zero |
| Data preprocessing | Nuisance | Regression | One least-squares fit per vertex of intercept, linear and quadratic terms over the whole concatenated series, WM and CSF means (5 columns); no band-pass. Then z-score per vertex over the retained frames (n − 1 in the denominator) |
| Data preprocessing | PAN | Input | ds006598 v2.0.1 (raw); fMRIPrep outputs sub-PAN*/ses-*/func/*_task-rest_run-*_space-fsLR_den-91k_bold.dtseries.nii (the 59,412 cortical grayordinates) and *_desc-confounds_timeseries.tsv; for the multi-echo ICA, the echo-wise series *_echo-{1–5}_desc-preproc_bold.nii.gz (native BOLD space), the run's *_desc-brain_mask.nii.gz and the echo times of the echo sidecars |
| Data preprocessing | PAN | Surfaces | Each subject's own, from FreeSurfer 7.3.2 recon-all inside fMRIPrep (T1w and T2w); hemi-?_midthickness.surf.gii resampled to fs_LR 32k with wb_command -surface-resample (BARYCENTRIC) from hemi-?_space-fsLR_desc-msmsulc_sphere.surf.gii to tpl-fsLR_hemi-?_den-32k_sphere.surf.gii |
| Data preprocessing | PAN | fMRIPrep call | nipreps/fmriprep:23.2.0 /data /out participant --participant-label PAN0x -t rest --output-spaces MNI152NLin6Asym:res-2 --cifti-output 91k --ignore slicetiming --me-output-echos --skip-bids-validation (--project-goodvoxels not used) |
| Data preprocessing | PAN | Multi-echo ICA | tedana 26.0.3, tedana_workflow per run on the five echo-wise series, with the brain mask and echo times above and dummy_scans = the number of leading volumes with non_steady_state_outlier* = 1; defaults otherwise (log-linear T2* fit, optimal combination t2s, PCA component selection by AIC, FastICA with seed 42, decision tree tedana_orig). Used: desc-ICA_mixing.tsv (component time courses) and the classification column of desc-tedana_metrics.tsv. Components per run 17–45 (median 27); rejected 1–22 (median 12) |
| Data preprocessing | PAN | Frames | Per run: volumes with non_steady_state_outlier* = 1 dropped; FD = framewise_displacement (first retained frame 0); frames with FD > 0.25 mm censored, linearly interpolated from the nearest retained frames for the fit, then removed; 1–650 frames censored per subject |
| Data preprocessing | PAN | Nuisance and scaling | Per run: least squares on [1, t, t<sup>2</sup>, white_matter, csf, all columns of the run's ICA mixing matrix (demeaned)] (t linear on [−1, 1]; WM and CSF demeaned); the fitted contributions of the first five columns and of the rejected components subtracted (accepted components left in the data); result kept at the retained frames; no global-signal regression; runs concatenated in session and run order; z-score per vertex (n − 1); vertices with no signal in any run set to 0 |
| Geodesic distance | Surfaces | Per subject | T1w/fsaverage_LR32k/<subject>.{L,R}.midthickness_MSMAll.32k_fs_LR.surf.gii (native space); cortex = <subject>.{L,R}.atlasroi.32k_fs_LR.shape.gii (29,696 L, 29,716 R vertices), medial wall removed from the mesh |
| Geodesic distance | Algorithm | Per subject | Exact geodesic (Mitchell–Mount–Papadimitriou) from every vertex |
| Geodesic distance | Cohort median | Rule and precision | Per pair, median over the subjects (mean of the two middle values when even); per-subject and median distances are stored at 16-bit floating-point precision, and every distance cut is applied to the stored median |
| Geodesic distance | Cohort median | Subjects | 7T: the 151; 3T: the 349; movie: the 160; PAN: the 10, own surfaces, distances to 40 mm |
| Geodesic distance | Window | Bounds | 3 ≤ d < 20 mm; check: 12,330,406 window candidates outside the peri-allocortex, both hemispheres |
| Null floor | Null pairs | Areas (HCP-MMP1.0) | Occipital V1, V2, V3, V4; frontal pole 10d, 10pp, a10p, p10p, 10r, 10v; sensorimotor 4, 3a, 3b, 1, 2 |
| Null floor | Null pairs | Draw | For each target (frontal pole, sensorimotor) and each direction (left occipital to right target, right occipital to left target): 200,000 occipital and 200,000 target vertices drawn uniformly with replacement and paired in order; 800,000 pairs (764,915 distinct), the same for every subject and cohort; none touches the peri-allocortex |
| Null floor | Correlations | Precision | Pearson r of every vertex pair computed per subject in single precision (32-bit floating point) from the z-scored series, r = $\frac{Z^{\mathrm{T}}Z}{T-1}$ |
| Null floor | Floor | Estimate | FDR ( τ ) = (null pairs with r > τ / 800,000) × (window candidates / window candidates with r > τ); τ<sub>s</sub> = the smallest τ with FDR ( τ ) ≤ 0.05, found by bisection on [0, the subject's largest null r] (60 halvings, upper end kept), from the window candidates of both hemispheres pooled |
| Null floor | Floor | Check values | 7T: median τ<sub>s</sub> 0.113, interquartile range 0.089–0.139, range 0.061–0.278; subject 572045 (Figure 1A,B) 0.10279; subject 550439 (Figure 3A) 0.10253; in the median subject, 2.8% of the null pairs and 57% of the window candidates lie above τ<sub>s</sub> |
| Cohort graph | Support | Majority | ⌊N / 2⌋ + 1 subjects: 7T 76; halves 39 and 38; women 45, men 32; movie 81; 3T 175; PAN 6; single subject 1 |
| Cohort graph | Window graph | Check values | 3,475,449 L and 3,517,487 R edges (no majority-supported pair has mean r ≤ 0) |
| Cohort graph | Short-range density | Block medians | Unweighted density (the fraction of a vertex's neighbors that are its window partners), median over the vertices of each block: 0.729 L, 0.735 R lattice; 0.482 L, 0.494 R paralattice (median window degree 300 L, 291 R and 190 L, 198 R) |
| Cohort graph | Short-range density | Regions of the Results | Medians over the vertices of the following HCP-MMP1.0 areas. Inferior parietal lobule: PGi, PGs, PFm, PF. Early visual cortex: V1, V2, V3. Dorsal extrastriate cortex: V3A, V3B, V6, V7, IPS1. Intraparietal sulcus: IP0, IP1, IP2, LIPv, LIPd, VIP, MIP, AIP. Precuneus and posterior cingulate: 7m, 7Pm, PCV, 31pd, 31pv, v23ab, d23ab. Postcentral cortex: 1, 2, 3b. Lateral prefrontal cortex: 46, 9-46d, p9-46v, 8Av, 8C. Motor cortex: 4. Temporal pole: TGd, TGv. Cingulate motor areas: 24dd, 24dv. Auditory cortex: A1, MBelt, LBelt, PBelt. Orbitofrontal cortex: OFC, pOFC, 11l, 13l. Insula: AVI, MI, PoI1, PoI2, Ig. |
| Cohort graph | Full-range graph | Without peri-allocortex edges | Within hemisphere, outside the peri-allocortex, support ≥ 76: 21,112,384 L and 22,227,425 R edges, of which 126,458 L and 127,589 R at d < 3 mm; smallest weight ( mean r ) 0.078 |
| Cohort graph | Full-range graph | With peri-allocortex edges | The same rule with mean r > 0 and pairs touching the peri-allocortex kept: 21,310,970 L and 22,375,674 R edges (198,586 L and 148,249 R with a peri-allocortex end). Used for the long-range, gradient, network and connector analyses |
| Cohort graph | Peri-allocortex | Mask | Whole HCP-MMP1.0 areas H, EC, PreS, PeEc, Pir, 25, PHA1, PHA2, PHA3, TF, s32, 33pr and AAIC in each hemisphere: 1,741 L + 1,795 R = 3,536 vertices |
| Cohort graph | Atlas | HCP-MMP1.0 | CIFTI dlabel on the 32k fs_LR mesh, 360 areas (labels L_<area>_ROI, R_<area>_ROI) |
| Distance-dependent regimes | p ( d ) | Bins | 1-mm bins [ k, k + 1 ) , k = 0 − 299 mm, d = k + 0.5; numerator: full-range edges outside the peri-allocortex; denominator: all within-hemisphere pairs outside it (390,727,035 L, 389,777,160 R); bins with p ( d ) = 0 dropped (245 L, 247 R bins fitted). Each pair is counted with the weight a<sub>i</sub>a<sub>j</sub> (Area weighting), in the numerator and in the denominator. |
| Distance-dependent regimes | Fit | Procedure | Least squares on ln p ( d ) , each bin weighted by its (area-weighted) number of pairs (divided by the mean per bin). Single exponential: A in closed form, ℓ by grid (2–400 mm, step 0.05) then Nelder–Mead. Two exponentials: parameters as logarithms, Nelder–Mead from 224 starts (A<sub>1</sub> from the single fit; ℓ<sub>1</sub> ∈ {5, 10, 15, 17, 20, 25, 35} mm; ℓ<sub>2</sub> ∈ {60, 100, 150, 220, 400, 800, 2,000, 10,000} mm; A<sub>2</sub> ∈ {0.01, 0.03, 0.06, 0.1}), best kept, components ordered so that ℓ<sub>1</sub> < ℓ<sub>2</sub>. BIC = n ln $\left(\frac{\mathrm{SSE}}{n}\right)$ + k ln n, n = bins fitted, k = 2 or 4 |
| Distance-dependent regimes | Fit | Check values | A<sub>1</sub> = 0.837 L, 0.755 R; A<sub>2</sub> = 0.0312 L, 0.0298 R; ℓ<sub>1</sub> = 19.82 L, 22.05 R mm; ℓ<sub>2</sub> unbounded (the fit stops at > 10<sup>16</sup> mm); crossover 65.2 L, 71.3 R mm; ΔBIC 355 L, 298 R |
| Distance-dependent regimes | Edges beyond 200 mm | Share and p(d) | Full-range edges outside the peri-allocortex at d ≥ 200 mm: 874,227 L, 733,992 R, in 303 and 336 pairs of HCP-MMP1.0 areas, ranked by their share of the summed a<sub>i</sub>a<sub>j</sub>. Joining {PGi, PGs, STSvp, TE1p} to {9m, 10d, 9p, 9a, 8BL, 8Ad}: 50.9% L, 39.7% R of the summed a<sub>i</sub>a<sub>j</sub> (449,485 and 290,974 edges; 51.4% and 39.6% as a count). Vertex pairs joining the two groups: 929,563 L, 707,427 R (4.5% and 3.8% of the summed a<sub>i</sub>a<sub>j</sub> of the pairs beyond 200 mm). p(d) at 200–210 mm: 0.043 L and R; without those pairs 0.028 L, 0.031 R; at 210–220 mm 0.049 L, 0.044 R and 0.025 L, 0.026 R. Longest edge 245 L, 246 R mm |
| Distance-dependent regimes | Mean-r profile | Pairs | Candidate pairs closer than 20 mm outside the peri-allocortex, bins [ k, k + 1 ) mm, mean over pairs and then over subjects |
| Distance-dependent regimes | Random pairs | Draw | Per hemisphere, 600,000 first and 600,000 second vertices drawn with replacement outside the peri-allocortex, self-pairs removed; of the pooled pairs ordered by vertex index, the first 1,000,000 (all left pairs and the right pairs of lowest index) |
| Spectral partitioning | Eigenproblem | Partition | L<sub>sym</sub> = I − D<sup>−1/2</sup>WD<sup>−1/2</sup> on the giant component; eight smallest eigenpairs by shift-invert Lanczos about −10<sup>−6</sup>; v<sub>2</sub> = D<sup>−1/2</sup>u<sub>2</sub> |
| Spectral partitioning | Eigenproblem | Figure 2A | Eleven largest eigenvalues of D<sup>−1/2</sup>WD<sup>−1/2</sup>, λ = 1 − eigenvalue |
| Spectral partitioning | Sweep | Splits | Vertices sorted by v<sub>2</sub> ascending; split t puts the first t + 1 in one block; t from 200 to n − 201 (PAN: margin max(5% of the giant component, 50 vertices)) |
| Spectral partitioning | Descent | Rule | Candidates: vertices whose window-graph weight to the other block exceeds that to their own, ranked by the difference at the start of each round. The ranked list is divided into ⌊m / 100⌋ consecutive batches, at least one, whose sizes differ by at most one (m = number of candidates; about 100 vertices each), moved in order; a batch is kept if the normalized cut falls; at the first batch that fails, its vertices are tried one at a time and the round ends. Rounds repeat (at most 60) until one moves no vertex |
| Spectral partitioning | Descent | Islands | After the descent, block 1 and then block 2: every connected component of the block in the window graph except the largest moves to the other block |
| Spectral partitioning | Labels | Blocks | Block 1 (lattice) is the block that contains V1; replications and controls take the labeling that agrees better with the 7T partition |
| Spectral partitioning | Area weighting | Vertex areas | One third of the summed area of the triangles at each vertex of T1w/fsaverage_LR32k/<subject>.{L,R}.midthickness_MSMAll.32k_fs_LR.surf.gii, mean over the 151 subjects, divided by the hemisphere mean over the vertices outside the peri-allocortex (2.99 mm² L, 3.03 mm² R). Every edge weight is multiplied by the areas of its two vertices, w a<sub>i</sub>a<sub>j</sub>, in the window and control graphs of every cut, spectrum and restart profile, and in p(d). Check: a<sub>i</sub> 0.56–1.54 (5th–95th percentile outside the peri-allocortex); mean area 2.88 mm² in lattice and 3.25 mm² in paralattice cortex (left) |
| Spectral partitioning | Check values | 7T partition | Eigenvalue ratio λ<sub>3</sub> / λ<sub>2</sub> 5.06 L, 4.54 R; normalized cut 0.0108 L, 0.0124 R; blocks 18,952 / 9,003 L and 18,966 / 8,955 R vertices; 99 L and 154 R vertices moved by the descent |
| Spectral partitioning | Signal dropout | Areas removed | TGd, TGv and TE2a (temporal pole and anterior ventral temporal cortex) and OFC, pOFC, 11l and 13l (orbitofrontal cortex), HCP-MMP1.0, removed from the graph of each hemisphere before the spectral step; 1,431 L and 1,528 R vertices; ARI with the 7T partition 1.00 L, 1.00 R (Supplementary Results S2) |
| Spectral partitioning | Depth of the division | Starts | 121 starts, t = 200 + $\frac{i(n-400)}{120}$, i = 0 − 120, truncated to integers; the first t + 1 vertices of the Fiedler ordering in one block; descent to convergence |
| Spectral partitioning | Depth of the division | Check values | Lowest normalized cut over the 121 starts 0.0105 L, 0.0124 R; on the left it is the variant of Supplementary Results S2 (reported cut 0.0108 L, 0.0124 R); every start ending within 10% of the lowest cut agrees with the reported partition at ARI ≥ 0.81 L, 0.96 R; the median cut over the starts is 2.8 L, 2.8 R times the lowest |
| Spectral partitioning | Figure S3 | Graph | Full-range graph with peri-allocortex edges at d ≥ 45 mm (13,481,638 L, 14,290,189 R edges); peri-allocortex removed before the spectral step (13,335,640 L, 14,191,000 R edges; giant component 24,215 L, 24,336 R vertices) |
| Spectral partitioning | Homotopy | Mesh index | Vertices with the same index on the 32,492-vertex fs_LR 32k mesh; 29,653 indices cortical in both hemispheres |
| Controls | Own-degree and degree-scrambled | Counts | Each vertex's own window degree, or the window degrees permuted over all vertices of the hemisphere; the window candidates of each vertex are sorted by distance (ties by pair index), the nearest are taken up to that count, and a pair is an edge if either vertex selects the other; empirical weights assigned at random |
| Controls | Rewiring | Maximum entropy | p<sub>ij</sub> = $\frac{x_i x_j}{1+x_i x_j}$ on the window candidates, x solved so that the expected degree equals the window degree (fixed-point iteration x ← $\frac{x+\deg/s}{2}$, s<sub>i</sub> = Σ<sub>j</sub> x<sub>j</sub> / ( 1 + x<sub>i</sub>x<sub>j</sub>)); a pair is kept with probability p<sub>ij</sub>; empirical weights assigned at random. Distance-matched: the same in each 1-mm band from 3 mm with the band degree as target |
| Controls | Weakest edges and all pairs | Weights | Weakest: w − min ( w ) + 10<sup>−6</sup>; all pairs: max(mean r, 10<sup>−6</sup>) |
| Controls | Surrogates | Modes | Window graph, vertices with positive degree; eigenvectors u<sub>2</sub> − u<sub>60</sub> of L<sub>sym</sub> (60 eigenpairs); coefficients standard normal divided by $\sqrt{\max\left(\lambda_k, 10^{-3}\lambda_2\right)}$; 1,000 maps (seed 1, the sign of each eigenvector fixed so that its largest entry in absolute value is positive); maps and u<sub>2</sub> multiplied by D<sup>−1/2</sup>, sorted and swept with margin 200, no descent. Check: normalized cut of the empirical sweep 0.0110 L, 0.0127 R; surrogates, mean 0.0795 L, 0.0824 R, lowest 0.0131 L, 0.0149 R; no map of four draws of 1,000 at or below the empirical sweep |
| Controls | Quoted cuts | Definition | The lowest normalized cut found on the control graph by sweep (margin 200) and descent; for the geometric graph, the lowest over the 121 starts. Profile flatness: fraction of splits within the margins with cut ≤ 1.2 × the minimum (empirical 0.037 L, 0.026 R; geometric graph 0.38 L, 0.39 R) |
| Controls | Strongest edges | Graph | Full-range graph without peri-allocortex edges; the top 3,475,449 L and 3,517,487 R edges by w |
| Controls | Long-range graphs | Construction | Edges of the full-range graph with peri-allocortex edges at d ≥ cutoff (floors not recalibrated); peri-allocortex removed before the spectral step |
| Controls | Long-range graph vs. networks | Settings | Cutoffs 20–70 mm in 5-mm steps, 80 and 90 mm; giant component; gradients: \|r\| of D<sup>−1/2</sup>u<sub>2</sub> − D<sup>−1/2</sup>u<sub>4</sub> with neuromaps margulies2016 fcgradient01–03 (fsLR 32k); K = 2 median split of D<sup>−1/2</sup>u<sub>2</sub>; K = 7 k-means (30 initializations) on u<sub>2</sub> − u<sub>7</sub> rows scaled to unit length; Dice with one-to-one Hungarian matching against Yeo_JNeurophysiol11_7Networks.32k.{L,R}.label.gii (HCP fs_LR 32k templates) |
| Controls | Long-range graph vs. networks | Check values | η<sup>2</sup> of gradient 1 by the two blocks, d ≥ 20 mm graph: 0.129 L, 0.143 R |
| Robustness | Window | Floors | Not recalibrated; every window variant is a subset of the 3–20 mm accumulation |
| Robustness | Split halves | Assignment | Subject i of the cohort order (i = 0, 1, …) to group i mod 6; half A = groups 0–2 (76 subjects, majority 39); half B = groups 3–5 (75, majority 38) |
| Robustness | Global-signal regression | Definition | Global signal = mean over the 59,412 cortical vertices of the z-scored series; regressed from every vertex (no intercept); re-centered and re-z-scored; floors recalibrated |
| Robustness | Peri-allocortex | Mask lifted | Whole chain rerun with the peri-allocortex pairs in the window and the floors recalibrated |
| Robustness | Local-fdr floor | Model | Efron's local false-discovery rate with an empirical null by central matching, on r (not Fisher z). f = histogram density of r (bins of 0.0025 from −0.6 to 1.0, values outside clipped into the end bins; centered 5-bin moving average). f<sub>0</sub> = Gaussian N$\left(\mu_0, \sigma_0^2\right)$ from a least-squares quadratic fit of log f, log f ≈ a<sub>2</sub>r<sup>2</sup> + a<sub>1</sub>r + a<sub>0</sub>, weighted by $\sqrt{\mathrm{bin\ count}}$, over the bins with nonzero count in mode − s < r < mode + 0.5s, with mode the peak of the smoothed f and s the root-mean-square deviation from the mode of the r values below it; σ<sub>0</sub><sup>2</sup> = −$\frac{1}{2a_2}$, μ<sub>0</sub> = a<sub>1</sub>σ<sub>0</sub><sup>2</sup>, π<sub>0</sub> = exp$\left(a_0 + \frac{\mu_0^2}{2\sigma_0^2}\right)$ · σ<sub>0</sub>√2π , all from the same fit (closed form, no iteration). No model of the non-null part. lfdr ( r ) = min$\left(1, \frac{\pi_0 f_0(r)}{f(r)}\right)$. Own implementation (Python), not locfdr |
| Robustness | Local-fdr floor | Data | Per subject, one histogram for both hemispheres pooled: every within-hemisphere pair of vertices outside the peri-allocortex (780,504,195 pairs), minus the 254,047 pairs at cohort geodesic d < 3 mm; all pairs, no sampling; the stored 32-bit r used for τ<sub>s</sub> |
| Robustness | Local-fdr floor | Floor | The first bin center r > 0, scanning upward, at which lfdr ≤ 0.1 (no interpolation; bins beyond the first crossing not examined); one floor per subject. A pair counts when r > floor, compared in 32-bit floating point as for τ<sub>s</sub>. Downstream unchanged: window candidates of record, support ≥ 76 of 151, weight the mean r over all 151 subjects (the weights of record), same sweep and descent |
| Robustness | Local-fdr floor | Check values | Floor median 0.149, interquartile range 0.138–0.164, range 0.109–0.249; π<sub>0</sub> median 0.81 (range 0.72–0.85); subject 572045: μ<sub>0</sub> 0.001, σ<sub>0</sub> 0.055, π<sub>0</sub> 0.83, floor 0.166 (τ<sub>s</sub> 0.103); edges 2,659,953 L and 2,677,197 R (43.1% and 43.5% of window candidates; 76.5% and 76.1% of the edges of record); ARI 0.971 L, 0.974 R |
| Figure 1B,D | Strip | Seed and anchors | Seed: left vertex 5077 of the fs_LR 32k mesh, area 55b where it meets area 4, chosen by hand; anchors: vertices 7992 (area 3b) and 30752 (area 8Av), 71.1 mm apart, each 31–37 mm from the seed and well inside its block (26.9 and 33.7 mm from the border) |
| Figure 1B,D | Strip | Width and thinning | Width from 2.0 mm in 0.5-mm steps until at least 110 vertices remain after thinning (13.0 mm); vertices ordered by d ( anchor A ) − d ( anchor B ) and kept when at least 5 mm from every vertex already kept; evenly subsampled to 110 |
| Non-neural spread | Lower envelope | Fit | Per subject, the 5<sup>th</sup> percentile of r in each 1-mm bin of the window (3–19 mm), fitted as A<sub>env</sub> $e^{-\frac{d}{\ell_{env}}}$ on the untransformed values (bins with a positive percentile). Check: ℓenv = 1.947 ± 0.341 mm (mean ± SD over subjects) |
| Non-neural spread | Lower envelope | Extrapolation | A<sub>env</sub> refitted ( 0.8639 ) to the cohort-mean envelope over the 3–9 mm bins with ℓ<sub>env</sub> fixed at 1.947 mm; divided by the median correlation per bin (median over pairs in each subject, mean over subjects), interpolated log-linearly between bin centers and extended past 19.5 mm with the log slope of the last two bins |
| Non-neural spread | Facing banks | Distances and classes | Euclidean distance on the S1200 MSMAll mid-thickness surface; geodesic distance along its mesh edges (Dijkstra) within the cortex. Folded: geodesic/Euclidean ≥ 2.0; flat pool ≤ 1.15. Euclidean bins 4–6, 6–8 and 8–10 mm (at least 30 pairs each); geodesic bins of 1 mm from 3 to 20 mm (at least 20 pool pairs) |
| Non-neural spread | Within-area profiles | Areas and bins | Left V1, V2, 3b, 46, 9a, 10d; window candidates with both ends in the area; per subject, bins with at least 20 pairs; '5 mm' = the 5–6 mm bin, '20 mm' = the 19–20 mm bin |
| Non-neural spread | Venous exposure | Atlas and sampling | VENAT_PartialVolume.nii.gz (Huck et al.; figshare 7205960, CC BY 4.0; MNI152, 0.5 mm); 40 subjects of the 7T cohort (every third of the cohort list, from the first, up to 40: positions 1, 4, …, 118, 102311 to 680957); MNINonLinear white_MSMAll and pial_MSMAll 32k surfaces; trilinear interpolation at 0, 0.5, 1, 1.5, 2, 2.5 and 3 mm outward from the pial surface along the white-to-pial direction; maximum over these points, mean over subjects; points outside the atlas volume count as 0; where white and pial vertices coincide, all points fall on the pial vertex. Atlas on the FSL MNI152 0.5-mm grid (364 × 436 × 364), no registration to the surfaces' template. Removal: the 10, 20 and 30% most exposed vertices of each hemisphere and their edges. Elongation: orientation tensor of the unit tangent vectors on the fs_LR sphere; null with the same number of partners per 1-mm band from the vertex's own candidates, ten draws |
| Non-neural spread | Venous exposure | Check values | Median exposure, lattice vs. paralattice: 0.024 vs. 0.013 L, 0.023 vs. 0.011 R. Spearman ρ, exposure vs. weight per neighbor (lattice, paralattice): −0.11, −0.19 L; −0.09, −0.21 R. Lattice/paralattice weight per neighbor by exposure quintile: 1.41, 1.36, 1.70, 1.79, 2.06 L; 1.17, 1.50, 1.62, 1.46, 2.48 R (all vertices 1.56 L, 1.50 R). Band ratio at 15–20 mm without the 10, 20, 30% most exposed: 2.39, 2.36, 2.37 (all vertices 2.44). Descent from the reported border: ARI 0.984, 0.988, 0.986 L and 0.991, 0.985, 0.996 R; normalized cut 0.0096, 0.0095, 0.0095 L and 0.0120, 0.0116, 0.0116 R (full graph 0.0108 L, 0.0124 R). Axial anisotropy excess: lattice interior 0.016 L, 0.006 R; paralattice interior 0.041 L, 0.037 R (ρ with exposure 0.37 L, 0.22 R) |
| Non-neural spread | Venous exposure | Check values, sample points shifted by ±1 and ±2 mm along x, y and z (12 shifts) | Median exposure, lattice vs. paralattice: 0.020–0.024 vs. 0.012–0.013 L, 0.019–0.025 vs. 0.009–0.012 R. Spearman ρ, exposure vs. weight per neighbor: lattice −0.08 to −0.12, paralattice −0.16 to −0.21 L; −0.05 to −0.11, −0.18 to −0.24 R. Lattice/paralattice weight per neighbor by exposure quintile: 1.28 to 2.16 L, 1.11 to 2.49 R |
| Non-neural spread | Homotopic flatness | Regression | Left vertices with a homotopic partner, support ≥ 76 and w > 0 (21,800); regressor \|x\| on the S1200 MSMAll mid-thickness surface in units of 10 mm; least squares with intercept; 95% CI ± 1.96 SE |
| Border | Border vertices | Definition | Vertices joined to a vertex of the other block by a pair with cohort geodesic d ≤ 2.2 mm (lattice and paralattice only): 365 L, 420 R |
| Border | Crossing pairs | Without threshold | Mean cohort-mean r of the crossing pairs 0.074 L, 0.077 R (260,511 L, 294,069 R pairs); distance-matched expected value from the within-block pairs, the two blocks pooled, 0.149 L, 0.152 R; ratio 0.497 L, 0.506 R |
| Border | Crossing pairs | Crossing edges | 27,492 L, 34,867 R; mean support 97.9 L, 96.8 R; mean weight 0.170 L, 0.164 R (within lattice cortex 119.3, 120.0 and 0.241, 0.241; within paralattice cortex 108.8, 109.0 and 0.210, 0.209) |
| Border | Temporal SNR | Subjects and file | The 151 subjects of the 7T cohort; the four released runs (rfMRI_REST{1,2,3,4}_7T_{PA,AP,PA,AP}_Atlas_MSMAll_hp2000_clean_rclean_tclean.dtseries.nii), the 59,412 cortical grayordinates; mean/SD over time per run, mean over the four runs, median over subjects. Check values: border 97.0 L, 92.4 R; lattice interior 93.9, 89.4; paralattice interior 95.4, 96.6 |
| Within-block | Illustrative seeds | Vertices (fs_LR 32k mesh index, left) | V1 25215; V2 23595; 3b 7984; 46 30015; 9a 28856; 10d 28526. All but V2 are the area medoid on the cohort geodesic; V2 was chosen by hand among the V2 vertices whose partners fall almost entirely in one patch in subject 550439 |
| Within-block | Illustrative seeds | Partners | Subject 550439: the 450 strongest left-hemisphere pairs of the seed by r, kept when r > τ<sub>s</sub>; single linkage at < 10 mm on the cohort geodesic |
| Within-block | Partner geometry | Partner set | The min ( 450, full − range degree ) strongest edges by w in the full-range graph without peri-allocortex edges; pairs closer than 3 mm kept. Patches: single linkage at < 10 mm |
| Within-block | Partner sweep | Sample | K = 100, 200, 300, 450, 600, 900 partners on 8,000 vertices per hemisphere drawn at random (one draw) from all vertices outside the peri-allocortex with a block label; patches on the cohort geodesic, link < 10 mm and < 5 mm |
| Within-block | Near/far ratio | Formula | $\frac{(n_{3-20}+0.5)/\max(a_{3-20}, 1)}{(n_{\geq 60}+0.5)/\max(a_{\geq 60}, 1)}$, n counted over the 450 partners, a over all cortical vertices; Youden threshold searched over 197 quantiles (1–99%) of log<sub>10</sub> ratio: 1.82 L, 1.81 R |
| Within-block | Within-area shape | κ | V1–V2 and dorsolateral prefrontal (46, 9a, 10d); bins [ k, k + 1 ) mm from 3 to 50 mm; κ = twice the quadratic coefficient of a least-squares quadratic on min–max-scaled bin centers and means. Check: κ 1.05 L, 1.05 R (V1–V2), 3.03 L, 2.93 R (prefrontal) |
| Within-block | Internal integration | Bands | Full-range degree at d < 20 mm (pairs < 3 mm included), 20–50, 50–100 and ≥ 100 mm |
| Within-block | Homotopic edges | Supported | Vertices whose homotopic edge is supported by a majority of subjects: 78.5% L, 78.7% R lattice; 75.1% L, 75.3% R paralattice; 9.6% L, 9.4% R peri-allocortex |
| Sectors | Spectral step | Rule | Lattice window subgraph; eigenvectors u of L<sub>sym</sub> ( not rescaled by D<sup>−1/2</sup>) , columns 2 to K, rows scaled to unit length; k-means with 50 initializations |
| Sectors | Refinement | Rule | Single-vertex descent of Ncut<sub>K</sub> = $\sum_k \frac{\mathrm{cut}(A_k, V \setminus A_k)}{\mathrm{vol}(A_k)}$ on the lattice window graph, from the k-means labels: each round, candidates = vertices with more weight to another sector than to their own, ordered by the excess; a candidate moves to its best other sector if the move lowers Ncut<sub>K</sub> (exact update); at most 100 rounds. Applied to the K = 4 sectors and to the K = 6 solution that defines medial parietal cortex proper |
| Sectors | K selection, refined | Check values (homotopic ARI; split-half ARI L/R; lowest half vs. full) | With the descent applied at every K: K = 2: 0.546; 0.962/0.301; 0.335. K = 3: 0.799; 0.742/0.907; 0.751. K = 4: 0.919; 0.978/0.943; 0.959. K = 5: 0.591; 0.902/0.846; 0.856. K = 6: 0.626; 0.873/0.718; 0.718. K = 7: 0.697; 0.875/0.802; 0.753 |
| Sectors | K = 4 | Check values | Refined sectors: normalized cut per block 0.0355 L, 0.0448 R (main partition 0.0054 L, 0.0062 R); sizes visual 4,086/4,265, somatomotor 6,466/6,404, auditory-temporal 4,143/3,824, medial parietal 4,257/4,473 (L/R); medial parietal cortex proper 3,070/3,120 |
| Sectors | Full-range depth | Graph and score | Full-range cohort graph restricted to the sector's vertex set, edges with d ≥ 3 mm, length 1/w; same bases. Score: Spearman ρ between area-median depth (areas with ≥ 25 vertices in the sector) and hierarchical level; partial Spearman ρ controlling for the area-median geodesic distance from the base along the mid-thickness surface (Dijkstra on the fs_LR 32k mesh edges) |
| Sectors | Full-range depth | Hierarchical levels | Visual: V1 1; V2 2; V3 3; V4, V3A, V3B 4; MT, LO1, LO2, V7, IPS1 5; MST, FST, LIPv, LIPd, VIP, IP0, IP1, IP2, PH, FFC, PIT 6; PGp, PGi, PGs, TPOJ2, TPOJ3 7. Somatomotor: 3a, 3b 1; 1 2; 2 3; 5L, 5m, AIP, 7AL, 7PC, 7Am 4; OP1, PF, PFt, PFop 5. Auditory, both branches: A1 1; LBelt, MBelt 2; PBelt 3; A4 4; anterior branch: TA2, A5 5; STGa, STSda 6; STSva 7; TGd, TGv, TE1a 8; posterior branch: STSdp 5; STSvp, TPOJ1, PSL 6; STV, PGi 7. Areas with fewer than 25 vertices in the sector are not scored |
| Sectors | Full-range depth | Null | Distance-preserving rewiring: each edge keeps one endpoint, chosen at random, and its weight; the other endpoint is drawn uniformly from the sector vertices in the same 1-mm bin of cohort median geodesic distance from the kept endpoint; duplicate and self pairs dropped (about 22% of the edges); 100 graphs (seed 20261106); p = (number of null values ≥ observed + 1)/101 |
| Sectors | Full-range depth | Check values (L, R) | Areas scored: visual 19, 19; somatomotor 10, 10; auditory anterior branch 13, 13; posterior branch 9, 8. ρ (null mean; p): visual 0.87 (0.80; 0.01), 0.85 (0.79; 0.01); somatomotor 0.81 (0.68; 0.01), 0.74 (0.70; 0.01); anterior branch 0.99 (0.71; 0.01), 0.99 (0.98; 0.02); posterior branch 0.89 (0.65; 0.01), 0.99 (0.90; 0.01). Partial ρ: visual 0.74 (0.58; 0.01), 0.68 (0.60; 0.01); somatomotor 0.41 (−0.12; 0.01), 0.24 (−0.41; 0.01); anterior branch 0.68 (−0.23; 0.01), 0.53 (0.07; 0.01); posterior branch 0.14 (0.56; 1.00), −0.51 (0.27; 1.00). Surface distance from the base: ρ 0.67, 0.70 visual; 0.81, 0.81 somatomotor; 0.98, 0.99 anterior; 0.89, 0.99 posterior |
| Connectors | Paralattice connectors | Cut | Lattice vertices ranked by their number of long-range edges (d ≥ 60 mm) into paralattice cortex; connectors are the top-ranked vertices that together send three quarters of these edges, vertices tied at the cut included: at least 772 L and 758 R edges; 3,662 L and 3,540 R connectors. Peri-allocortex connectors by the same rule on long-range edges into the peri-allocortex: at least 39 L and 35 R edges; 864 L and 630 R connectors |
| Connectors | Overlap | Both populations | 506 L, 346 R vertices are both paralattice and peri-allocortex connectors (14% and 10% of the paralattice connectors; 59% and 55% of the peri-allocortex connectors) |
| Connectors | Parieto-frontal channels | Check values | Anterior group (PF, PFt, AIP, 7Am; 688 L, 745 R connectors): 46.6% L, 40.0% R of long-range edges into paralattice cortex end in 6r, 6a, 46 and IFSa; 0.4% L, 0.3% R in 8Ad, 8Av and 9m. Posterior group (PGi, PGs, 7m, 31pv; 832 L, 705 R): 34.2% L, 38.3% R in 8Ad, 8Av and 9m; 0.6% L, 1.4% R in 6r, 6a, 46 and IFSa. Overlap of the two target distributions 0.079 L, 0.091 R; no area shared among the ten main targets of each group. At edge lengths ≥ 120 mm: 46.4% L, 40.3% R and 33.9% L, 37.8% R |
| Connectors | Canonical networks | Files and names | Yeo_JNeurophysiol11_7Networks.32k.{L,R}.label.gii and Yeo_JNeurophysiol11_17Networks.32k.{L,R}.label.gii (HCP fs_LR 32k templates). 17-network labels 1–17: VisCent, VisPeri, SomMotA, SomMotB, DorsAttnA, DorsAttnB, SalVentAttnA, SalVentAttnB, Limbic_TempPole, Limbic_OFC, ContC, ContA, ContB, TempPar, DefaultC, DefaultA, DefaultB |
| Connectors | Canonical networks | Reporting | Enrichment computed when the network's share of lattice cortex exceeds 0.005; networks listed when the share is at least 0.02; hemispheres pooled |
| Replication | Movie | Residual form | Per subject and vertex, least squares on intercept and the cohort-mean z-scored series (mean over the subjects retaining each frame, frames aligned on the 3,655-frame grid before censoring, the subject included); residual re-z-scored |
| Replication | PAN | Sweep margin | max(5% of the giant component, 50) vertices |
| Replication | PAN | Single subjects (ARI L, R against the 7T partition) | PAN01 0.88, 0.91; PAN02 0.67, 0.81; PAN03 0.87, 0.88; PAN04 0.91, 0.90; PAN05 0.89, 0.62; PAN06 0.82, 0.67; PAN07 0.79, 0.81; PAN08 0.77, 0.70; PAN09 0.79, 0.68; PAN10 0.80, 0.77. Cohort graph 0.82, 0.86. With global-signal regression: PAN01 0.91, 0.88; PAN02 0.67, 0.82; PAN03 0.91, 0.89; PAN04 0.91, 0.89; PAN05 0.88, 0.69; PAN06 0.85, 0.77; PAN07 0.82, 0.596; PAN08 0.78, 0.76; PAN09 0.81, 0.76; PAN10 0.81, 0.92; cohort graph 0.92, 0.88. Without multi-echo ICA: 18 of the 20 hemispheres at ARI ≥ 0.60 (below: PAN07 R 0.561, PAN05 R 0.5995). Floors τ<sub>s</sub> without global-signal regression: PAN01 0.289, PAN02 0.143, PAN03 0.381, PAN04 0.270, PAN05 0.189, PAN06 0.306, PAN07 0.385, PAN08 0.172, PAN09 0.363, PAN10 0.248 |
| Replication | Block composition | Defining vertices' area and network membership | Cole–Anticevic network partition by Glasser parcel, cortex_ColeAnticevic_NetPartition_GlasserParcels_{L,R}.label.gii (fs_LR 32k); not the netassignments dlabel, whose vertex counts differ by up to 212 per network |
| Cortical maps | Annotations | Library | neuromaps; files source-<source>_desc-<desc>_space-<space>_den-<den>_hemi-<L\|R>_feature.func.gii, or _space-MNI152_res-<res>_feature.nii.gz for volumes |
| Cortical maps | Annotations | MEG power and timescale | hcps1200: megalpha, megbeta, megdelta, megtheta, meggamma1 (30–60 Hz), meggamma2 (60–150 Hz), megtimescale; fsLR 4k |
| Cortical maps | Annotations | Myelin, thickness, axis, gradients | hcps1200 myelinmap, thickness (fsLR 32k); sydnor2021 SAaxis (fsLR 32k); margulies2016 fcgradient01–03 (fsLR 32k) |
| Cortical maps | Annotations | Gene expression | abagen genepc1 (fsaverage 10k) |
| Cortical maps | Annotations | Metabolism and blood flow | raichle cmr02, cmrglc, cbf, cbv (fsLR 164k) |
| Cortical maps | Annotations | Receptors | norgaard2021 flumazenil (fsaverage 164k); kantonen2020 carfentanil (MNI152, res-3mm); alarkurtti2015 raclopride (MNI152, res-3mm); beliveau2017 dasb, cumi101, sb207145 (fsaverage 164k) |
| Cortical maps | Annotations | Other | xu2020 FChomology, evoexp (fsLR 32k); mueller2013 intersubjvar (fsLR 164k). Not used: hill2010 evoexp and devexp (right hemisphere only) |
| Cortical maps | Resampling | Rules | fs_LR 4k and 164k to 32k: nearest vertex on the fs_LR spheres (tpl-fsLR_den-*_sphere.surf.gii). fsaverage to fs_LR 32k: wb_command -metric-resample ADAP_BARY_AREA through the fsaverage-to-fs_LR registration sphere (neuromaps fsaverage_to_fslr, linear); fs_LR vertices without fsaverage cortex (618: 612 peri-allocortex, 6 paralattice) left missing. MNI152: trilinear interpolation at the vertices of tpl-fsLR_den-32k_hemi-{L,R}_midthickness.surf.gii |
| Cortical maps | Statistics | Vertices and check values | The 59,412 cortical vertices, hemispheres pooled; vertices without a value dropped per map (AHBA PC1 1,483; GABA-A, 5-HTT, 5-HT1A, 5-HT4 618 each). Block labels: lattice 37,918, paralattice 17,958, peri-allocortex 3,536. η<sub>H</sub><sup>2</sup> = $\frac{H-2}{n-3}$ |
| Cortical maps | BigBrain | Layer IV fraction | BigBrainWarp tpl-fsaverage_hemi-{L,R}_den-164k_desc-layer{1–6} thickness, resampled to fs_LR 32k as above; layer IV / sum of the six layers; undefined at 1,151 vertices in the fsaverage medial-wall mask or with zero total thickness. Quartile limits (linear interpolation, 58,261 vertices, peri-allocortex included): 0.0633, 0.0804, 0.0968 |
| Thalamus | Input | Series | The 7T rest CIFTI series above (4 runs, 3,600 frames, 91,282 grayordinates); every grayordinate (cortex, thalamus, striatum, cerebellum): FD > 0.25 mm censored (linear interpolation for the fit, then dropped); least squares on [1, t, t<sup>2</sup>, CSF] over the concatenation (t on [−1, 1]); z-score over the retained frames (n − 1); no white-matter regressor |
| Thalamus | Atlas | Morel atlas | Per-nucleus 1-mm masks in FSL MNI152 space (left-vols-1mm, right-vols-1mm); thalamic nuclei only (global, MAX_VOLUME, thalamus_body, mtt, STh, RN and Hb not used); LGNmc and LGNpc merged; where masks overlap, the smaller nucleus wins |
| Thalamus | Atlas | Labeling | CIFTI subcortical grid (2 mm); thalamus 1,288 L and 1,248 R grayordinates. Each grayordinate takes the most frequent nonzero label among 27 points at −1, 0 and +1 mm along each axis (nearest atlas voxel at each); a nucleus is kept with at least 3 grayordinates |
| Thalamus | Atlas | Check values (grayordinates) | Left: AM 6, AV 36, CL 96, CM 33, CeM 28, LD 27, LP 67, Li 14, MDmc 36, MDpc 111, MGN 5, MV 6, Pf 33, Po 6, PuA 20, PuL 35, PuM 252, SG 7, VAmc 12, VApc 78, VLa 41, VLpd 88, VLpv 90, VM 22, VPI 11, VPLa 19, VPLp 57, VPM 23. Right: AM 12, AV 47, CL 92, CM 33, CeM 37, LD 28, LP 76, Li 11, MDmc 30, MDpc 109, MGN 3, MV 4, Pf 30, Po 8, PuA 19, PuL 27, PuM 209, VAmc 8, VApc 83, VLa 48, VLpd 89, VLpv 90, VM 22, VPI 9, VPLa 15, VPLp 54, VPM 21. Not retained: AD, PuI, Pv (L), SG (R); LGN has no grayordinate |
| Thalamus | Partial correlation | Signals | Nucleus = mean of its grayordinates; control = mean of all other thalamic grayordinates of both hemispheres; striatum = mean of the ipsilateral caudate, putamen and accumbens; cerebellum = mean of the contralateral cerebellar cortex; partial correlations averaged over the vertices of each block |
| Thalamus | Block preference | Tests | Paired differences across subjects; d = $\frac{\mathrm{mean}}{\mathrm{SD}}$ ( n − 1 ) ; two-sided Wilcoxon signed-rank test (normal approximation); rule order: peri-allocortex (both contrasts p < 10<sup>−3</sup> and d ≥ 0.3), paralattice (p < 10<sup>−3</sup>, d ≥ 0.3), lattice (p < 10<sup>−3</sup>, d ≤ −0.3), otherwise unresolved. Loop closure on the 28 nuclei, the left and right estimates of each nucleus averaged (SG left only); control without the anterior nuclei AM, AV and LD (25 nuclei remain); permutation of the 28 hemisphere-averaged nuclei (100,000 permutations, seed 20261104): p < 10<sup>−5</sup> for the loop (ρ = +0.83), striatal (+0.80) and cerebellar (+0.80) couplings; p = 2 × 10<sup>−5</sup> for the loop without the anterior nuclei (25 nuclei, ρ = +0.82). Hemisphere-averaged ρ: loop +0.83 (p 4 × 10<sup>−8</sup>), striatal +0.80 (4 × 10<sup>−7</sup>), cerebellar +0.80 (4 × 10<sup>−7</sup>); without the anterior nuclei loop +0.82 (6 × 10<sup>−7</sup>) |
| Thalamus | Vertex-level coupling in motor cortex | Check values (Cohen's d, L and R) | Area 4 against the rest of lattice cortex: VLa −0.69, −0.85; VLpd −1.00, −1.10; VApc −1.18, −1.47; CM +1.77, +1.56; VPM +1.92, +1.88. Within area 4, rostral against caudal: VLa +1.15, +1.11; VLpv +0.95, +0.86. Least against most effector-selective tercile: VLa +0.71, +1.01; VLpv +0.38, +0.51; CM +0.48, +0.63. Lattice against paralattice part: VLa in 6d −0.87, −1.05; in 6mp −0.87, −1.18 |
| Thalamus | Nuisance model | Scores | Benchmark 1: the 20 estimates of VPM, VPLa, VPLp, MGN, LP, PuL (lattice targets) and VApc, VAmc, VLa, MDpc (paralattice targets), both hemispheres. Benchmark 2: the 43 estimates whose class in Table S3 predicts a block (basal ganglia or cerebellar input: paralattice; first-order relay or pulvinar: lattice). Agreements (benchmark 1, benchmark 2): none 18, 32. Trend 18, 32. Trend + censoring 18, 32. Trend + censoring + CSF (used) 19, 36. Trend + WM + CSF 17, 33. Trend + censoring + WM + CSF 17, 33 |
| Statistics | Effect sizes | Cohen's d (Figure 3) | $\frac{\mathrm{mean}_1 - \mathrm{mean}_2}{\sqrt{\frac{s_1^2+s_2^2}{2}}}$, s with n − 1; log<sub>10</sub> for partner distance and the near/far ratio; facing sulcal banks: the same formula, on raw values |
| Statistics | Spin test | Rotations | 1,000 uniformly random rotations Q of the unit fs_LR 32k sphere (tpl-fsLR_den-32k_hemi-{L,R}_sphere.surf.gii); the right hemisphere takes the mirror image M · Q · M, M = diag(−1, 1, 1); each vertex takes the block of the vertex nearest its rotated position and is dropped when that vertex lies in the medial wall (and, for Figure 3, the peri-allocortex) |
| Statistics | Spin test | p values | (1 + number of null values at least as extreme as observed)/(1 + number of nulls), two-sided for d and ρ, one-sided for η<sub>H</sub><sup>2</sup>; Benjamini–Hochberg over the 27 maps, separately for ρ and η<sub>H</sub><sup>2</sup> (smallest q 0.26 and 0.11) |
| Statistics | Bootstrap | Rebuild | 60 replicates; each gives every subject a multiplicity (multinomial, 151 draws); support = Σmultiplicity × [ r > τ<sub>s</sub>] ≥ 76; weight = Σmultiplicity × r / 151, kept if > 0; percentile intervals |
| Software | Versions | HCP preprocessing | Python 3.12; NumPy 2.5.1, SciPy 1.18.0, scikit-learn 1.8.0, nibabel 5.4.2 |
| Software | Versions | Analyses and figures | Python 3.10.6; NumPy 2.2.6, SciPy 1.15.3, scikit-learn 1.7.2, nibabel 5.4.2, matplotlib 3.10.9, neuromaps 0.0.5, Joblib 1.5.3, BrainSpace 0.1.22, gdist (tvb-gdist 2.9.2 for PAN); Connectome Workbench 2.1.0; PAN: fMRIPrep 23.2.0 (FreeSurfer 7.3.2, Connectome Workbench 1.5.0, MSMSulc), tedana 26.0.3 |

**Table S9.**
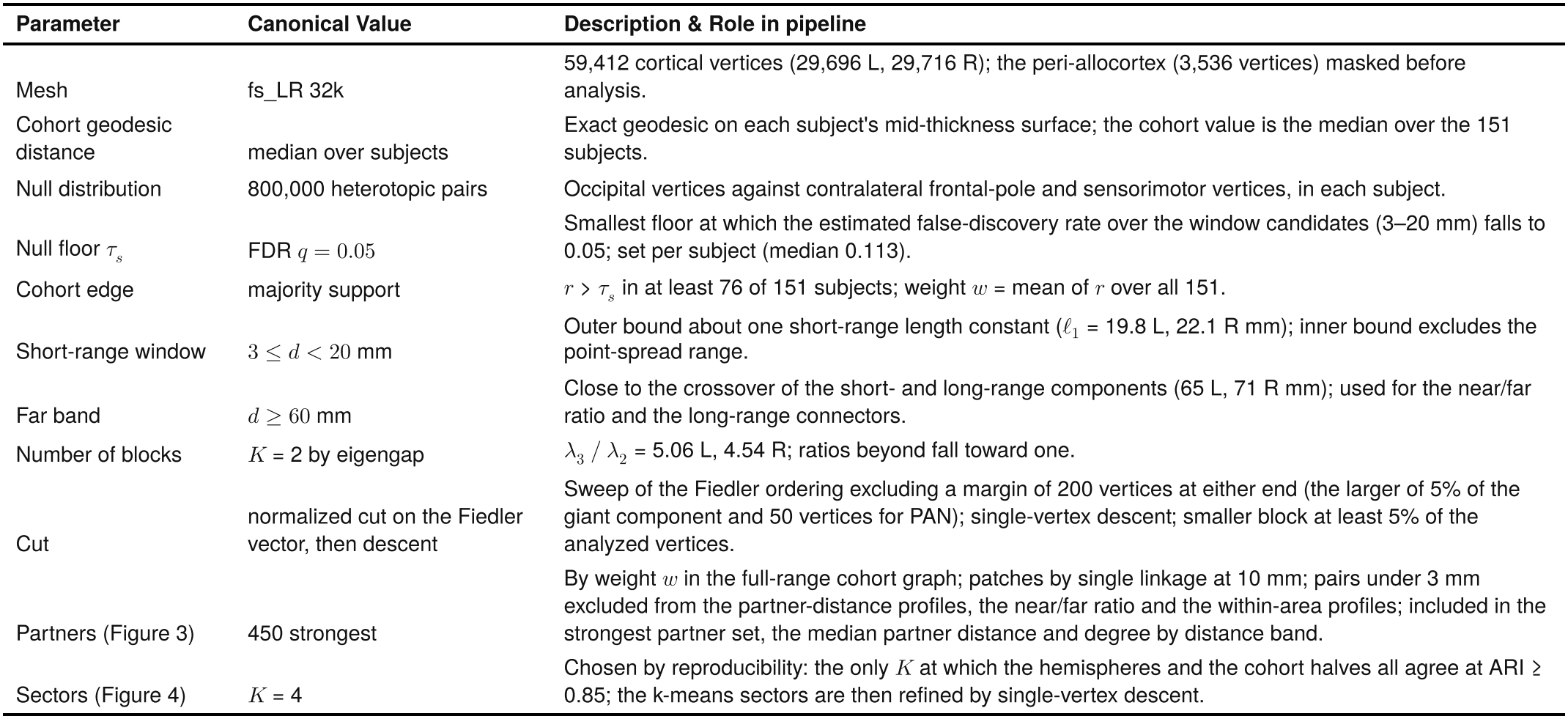
Hyperparameters for short-range graph extraction and partitioning (related to Methods).

| Parameter | Canonical Value | Description & Role in pipeline |
|---|---|---|
| Mesh | fs_LR 32k | 59,412 cortical vertices (29,696 L, 29,716 R); the peri-allocortex (3,536 vertices) masked before analysis. |
| Cohort geodesic distance | median over subjects | Exact geodesic on each subject's mid-thickness surface; the cohort value is the median over the 151 subjects. |
| Null distribution | 800,000 heterotopic pairs | Occipital vertices against contralateral frontal-pole and sensorimotor vertices, in each subject. |
| Null floor τ<sub>s</sub> | FDR *q* = 0.05 | Smallest floor at which the estimated false-discovery rate over the window candidates (3–20 mm) falls to 0.05; set per subject (median 0.113). |
| Cohort edge | majority support | *r* > τ<sub>s</sub> in at least 76 of 151 subjects; weight *w* = mean of *r* over all 151. |
| Short-range window | 3 ≤ *d* < 20 mm | Outer bound about one short-range length constant (ℓ<sub>1</sub> = 19.8 L, 22.1 R mm); inner bound excludes the point-spread range. |
| Far band | *d* ≥ 60 mm | Close to the crossover of the short- and long-range components (65 L, 71 R mm); used for the near/far ratio and the long-range connectors. |
| Number of blocks | *K* = 2 by eigengap | λ<sub>3</sub> / λ<sub>2</sub> = 5.06 L, 4.54 R; ratios beyond fall toward one. |
| Cut | normalized cut on the Fiedler vector, then descent | Sweep of the Fiedler ordering excluding a margin of 200 vertices at either end (the larger of 5% of the giant component and 50 vertices for PAN); single-vertex descent; smaller block at least 5% of the analyzed vertices. |
| Partners (Figure 3) | 450 strongest | By weight *w* in the full-range cohort graph; patches by single linkage at 10 mm; pairs under 3 mm excluded from the partner-distance profiles, the near/far ratio and the within-area profiles; included in the strongest partner set, the median partner distance and degree by distance band. |
| Sectors (Figure 4) | *K* = 4 | Chosen by reproducibility: the only *K* at which the hemispheres and the cohort halves all agree at ARI ≥ 0.85; the k-means sectors are then refined by single-vertex descent. |

## REFERENCES

1. Sporns, O., Tononi, G., and Kötter, R. (2005). The human connectome: A structural description of the human brain. PLoS Comput Biol 1, e42. 10.1371/journal.pcbi.0010042.

2. MICrONS Consortium (2025). Functional connectomics spanning multiple areas of mouse visual cortex. Nature 640, 435–447. 10.1038/s41586-025-08790-w.

3. Shapson-Coe, A., Januszewski, M., Berger, D.R., Pope, A., Wu, Y., Blakely, T., Schalek, R.L., Li, P.H., Wang, S., Maitin-Shepard, J., et al. (2024). A petavoxel fragment of human cerebral cortex reconstructed at nanoscale resolution. Science 384, eadk4858. 10.1126/science.adk4858.

4. Amir, Y., Harel, M., and Malach, R. (1993). Cortical hierarchy reflected in the organization of intrinsic connections in macaque monkey visual cortex. Journal of Comparative Neurology 334, 19–46. 10.1002/cne.903340103.

5. Majka, P., Bai, S., Bakola, S., Bednarek, S., Chan, J.M., Jermakow, N., Passarelli, L., Reser, D.H., Theodoni, P., Worthy, K.H., et al. (2020). Open access resource for cellular-resolution analyses of corticocortical connectivity in the marmoset monkey. Nat. Commun. 11, 1133. 10.1038/s41467-020-14858-0.

6. Markov, N.T., Ercsey-Ravasz, M.M., Ribeiro Gomes, A.R., Lamy, C., Magrou, L., Vezoli, J., Misery, P., Falchier, A., Quilodran, R., Gariel, M.A., et al. (2014). A weighted and directed interareal connectivity matrix for macaque cerebral cortex. Cereb Cortex 24, 17–36. 10.1093/cercor/bhs270.

7. Sepulcre, J., Liu, H., Talukdar, T., Martincorena, I., Yeo, B.T., and Buckner, R.L. (2010). The organization of local and distant functional connectivity in the human brain. PLoS Comput Biol 6, e1000808. 10.1371/journal.pcbi.1000808.

8. Tomasi, D., and Volkow, N.D. (2010). Functional connectivity density mapping. Proceedings of the National Academy of Sciences 107, 9885–9890. 10.1073/pnas.1001414107.

9. Zang, Y., Jiang, T., Lu, Y., He, Y., and Tian, L. (2004). Regional homogeneity approach to fMRI data analysis. Neuroimage 22, 394–400. 10.1016/j.neuroimage.2003.12.030.

10. Xu, A.G., Qian, M., Tian, F., Xu, B., Friedman, R.M., Wang, J., Song, X., Sun, Y., Chernov, M.M., Cayce, J.M., et al. (2019). Focal infrared neural stimulation with high-field functional MRI: A rapid way to map mesoscale brain connectomes. Science Advances 5. 10.1126/sciadv.aau7046.

11. Mountcastle, V.B. (1997). The columnar organization of the neocortex. Brain 120, 701–722. 10.1093/brain/120.4.701.

12. Tononi, G., Sporns, O., and Edelman, G.M. (1992). Reentry and the problem of integrating multiple cortical areas: simulation of dynamic integration in the visual system. Cereb Cortex 2, 310–335. 10.1093/cercor/2.4.310.

13. Tononi, G., Sporns, O., and Edelman, G.M. (1994). A measure for brain complexity: relating functional segregation and integration in the nervous system. Proc Natl Acad Sci U S A 91, 5033–5037. 10.1073/pnas.91.11.5033.

14. Barbas, H., Zikopoulos, B., and John, Y.J. (2022). The inevitable inequality of cortical columns. Front Syst Neurosci 16, 921468. 10.3389/fnsys.2022.921468.

15. Tononi, G. (2008). Consciousness as integrated information: a provisional manifesto. Biol Bull 215, 216–242. 10.2307/25470707.

16. Tononi, G., Boly, M., Massimini, M., and Koch, C. (2016). Integrated information theory: from consciousness to its physical substrate. Nat. Rev. Neurosci. 17, 450–461. 10.1038/nrn.2016.44.

17. Van Essen, D.C., Smith, S.M., Barch, D.M., Behrens, T.E.J., Yacoub, E., Ugurbil, K., and WU-Minn Hcp Consortium (2013). The WU-Minn Human Connectome Project: an overview. Neuroimage 80, 62–79. 10.1016/j.neuroimage.2013.05.041.

18. Markov, N.T., Misery, P., Falchier, A., Lamy, C., Vezoli, J., Quilodran, R., Gariel, M.A., Giroud, P., Ercsey-Ravasz, M., Pilaz, L.J., et al. (2011). Weight consistency specifies regularities of macaque cortical networks. Cereb Cortex 21, 1254–1272. 10.1093/cercor/bhq201.

19. Ercsey-Ravasz, M., Markov, N.T., Lamy, C., Van Essen, D.C., Knoblauch, K., Toroczkai, Z., and Kennedy, H. (2013). A Predictive Network Model of Cerebral Cortical Connectivity Based on a Distance Rule. Neuron 80, 184–197. 10.1016/j.neuron.2013.07.036.

20. Grill-Spector, K., and Weiner, K.S. (2014). The functional architecture of the ventral temporal cortex and its role in categorization. Nat Rev Neurosci 15, 536–548. 10.1038/nrn3747.

21. Ladwig, Z., Kermani, K.Z., Park, Y., Housteau, E., Dworetsky, A., Labora, N., Hernandez, J.J., Dorn, M., Smith, D.M., Nee, D.E., et al. (2026). Precision fMRI reveals densely interdigitated network patches with conserved motifs in the lateral prefrontal cortex. Neuron 114, 3499–3516.e8. 10.1016/j.neuron.2026.04.011.

22. Tsao, D.Y., Schweers, N., Moeller, S., and Freiwald, W.A. (2008). Patches of face-selective cortex in the macaque frontal lobe. Nature neuroscience 11, 877–879. 10.1038/NN.2158.

23. von Economo, C., and Koskinas, G.N. (2008). Atlas of Cytoarchitectonics of the Adult Human Cerebral Cortex (Karger).

24. Shi, J., and Malik, J. (2000). Normalized cuts and image segmentation. IEEE Trans. Pattern Anal. Mach. Intell. 22, 888–905. 10.1109/34.868688.

25. Ng, A., Jordan, M., and Weiss, Y. (2001). On spectral clustering: analysis and an algorithm. In Advances in Neural Information Processing Systems, T. Dietterich, S. Becker, and Z. Ghahramani, eds. (MIT Press).

26. Pang, J.C., Aquino, K.M., Oldehinkel, M., Robinson, P.A., Fulcher, B.D., Breakspear, M., and Fornito, A. (2023). Geometric constraints on human brain function. Nature 618, 566–574. 10.1038/s41586-023-06098-1.

27. Normand, F., Gajwani, M., Cao, T., Cruddas, J., Sangchooli, A., Oldham, S., Holmes, A., Robinson, P.A., Pang, J.C., and Fornito, A. (2026). Geometric constraints on the architecture of mammalian cortical connectomes. Cell 189,5283–5303.e17. 10.1016/j.cell.2026.05.048.

28. Alexander-Bloch, A.F., Shou, H., Liu, S., Satterthwaite, T.D., Glahn, D.C., Shinohara, R.T., Vandekar, S.N., and Raznahan, A. (2018). On testing for spatial correspondence between maps of human brain structure and function. NeuroImage 178, 540–551. 10.1016/j.neuroimage.2018.05.070.

29. Gilbert, C.D., and Wiesel, T.N. (1989). Columnar specificity of intrinsic horizontal and corticocortical connections in cat visual cortex. The Journal of Neuroscience 9, 2432–2442. 10.1523/jneurosci.09-07-02432.1989.

30. Lund, J.S., Angelucci, A., and Bressloff, P.C. (2003). Anatomical substrates for functional columns in macaque monkey primary visual cortex. Cereb. Cortex 13, 15–24. 10.1093/cercor/13.1.15.

31. Rockland, K.S., and Lund, J.S. (1983). Intrinsic laminar lattice connections in primate visual cortex. J. Comp. Neurol. 216, 303–318. 10.1002/cne.902160307.

32. Watakabe, A., Skibbe, H., Nakae, K., Abe, H., Ichinohe, N., Rachmadi, M.F., Wang, J., Takaji, M., Mizukami, H., Woodward, A., et al. (2023). Local and long-distance organization of prefrontal cortex circuits in the marmoset brain. Neuron 111, 2258–2273 e10. 10.1016/j.neuron.2023.04.028.

33. Seidman, S.B. (1983). Network structure and minimum degree. Soc. Networks5, 269–287. 10.1016/0378-8733(83)90028-x.

34. Bosking, W.H., Kretz, R., Pucak, M.L., and Fitzpatrick, D. (2000). Functional specificity of callosal connections in tree shrew striate cortex. J Neurosci 20, 2346–2359. 10.1523/JNEUROSCI.20-06-02346.2000.

35. Innocenti, G.M., Schmidt, K., Milleret, C., Fabri, M., Knyazeva, M.G., Battaglia-Mayer, A., Aboitiz, F., Ptito, M., Caleo, M., Marzi, C.A., et al. (2022). The functional characterization of callosal connections. Prog Neurobiol 208, 102186. 10.1016/j.pneurobio.2021.102186.

36. Jones, E.G. (2001). The thalamic matrix and thalamocortical synchrony. Trends in Neurosciences 24, 595–601. 10.1016/s0166-2236(00)01922-6.

37. Sherman, S.M. (2016). Thalamus plays a central role in ongoing cortical functioning. Nat Neurosci 19, 533–541. 10.1038/nn.4269.

38. Lanciego, J.L., Luquin, N., and Obeso, J.A. (2012). Functional Neuroanatomy of the Basal Ganglia. Cold Spring Harbor Perspectives in Medicine 2, a009621– a009621. 10.1101/cshperspect.a009621.

39. McFarland, N.R., and Haber, S.N. (2002). Thalamic Relay Nuclei of the Basal Ganglia Form Both Reciprocal and Nonreciprocal Cortical Connections, Linking Multiple Frontal Cortical Areas. The Journal of Neuroscience 22, 8117–8132. 10.1523/JNEUROSCI.22-18-08117.2002.

40. Mengual, E., de las Heras, S., Erro, E., Lanciego, J.L., and Giménez-Amaya, J.M. (1999). Thalamic interaction between the input and the output systems of the basal ganglia. Journal of Chemical Neuroanatomy 16, 187–200. 10.1016/s0891-0618(99)00010-1.

41. Alexander, G.E., DeLong, M.R., and Strick, P.L. (1986). Parallel organization of functionally segregated circuits linking basal ganglia and cortex. Annu Rev Neurosci 9, 357–381. 10.1146/annurev.ne.09.030186.002041.

42. Bostan, A.C., and Strick, P.L. (2018). The basal ganglia and the cerebellum: nodes in an integrated network. Nat Rev Neurosci 19, 338–350. 10.1038/s41583-018-0002-7.

43. Middleton, F.A., and Strick, P.L. (2002). Basal-ganglia “Projections” to the Prefrontal Cortex of the Primate. Cerebral Cortex 12, 926–935. 10.1093/cercor/12.9.926.

44. Hoover, J.E., and Strick, P.L. (1999). The Organization of Cerebellar and Basal Ganglia Outputs to Primary Motor Cortex as Revealed by Retrograde Transneuronal Transport of Herpes Simplex Virus Type 1. The Journal of Neuroscience 19, 1446–1463. 10.1523/JNEUROSCI.19-04-01446.1999.

45. Markello, R.D., Hansen, J.Y., Liu, Z.-Q., Bazinet, V., Shafiei, G., Suarez, L.E., Blostein, N., Seidlitz, J., Baillet, S., Satterthwaite, T.D., et al. (2022). neuromaps: structural and functional interpretation of brain maps. Nat. Methods 19, 1472–1479. 10.1038/s41592-022-01625-w.

46. Shafiei, G., Baillet, S., and Misic, B. (2022). Human electromagnetic and haemodynamic networks systematically converge in unimodal cortex and diverge in transmodal cortex. PLOS Biol. 20, e3001735. 10.1371/journal.pbio.3001735.

47. Wagstyl, K., Larocque, S., Cucurull, G., Lepage, C., Cohen, J.P., Bludau, S., Palomero-Gallagher, N., Lewis, L.B., Funck, T., Spitzer, H., et al. (2020). BigBrain 3D atlas of cortical layers: Cortical and laminar thickness gradients diverge in sensory and motor cortices. PLOS Biology 18, e3000678. 10.1371/journal.pbio.3000678.

48. Schmahmann, J.D., and Pandya, D.N. (2006). Fiber Pathways of the Brain (Oxford University Press).

49. Shastin, D., Genc, S., Parker, G.D., Koller, K., Tax, C.M., Evans, J., Hamandi, K., Gray, W.P., Jones, D.K., and Chamberland, M. (2022). Surface-based tracking for short association fibre tractography. Neuroimage 260, 119423. 10.1016/j.neuroimage.2022.119423.

50. Russo, S., Pigorini, A., Mikulan, E., Sarasso, S., Rubino, A., Zauli, F.M., Parmigiani, S., d’Orio, P., Cattani, A., Francione, S., et al. (2021). Focal lesions induce large-scale percolation of sleep-like intracerebral activity in awake humans. NeuroImage 234, 117964. 10.1016/j.neuroimage.2021.117964.

51. Shmuel, A., Yacoub, E., Chaimow, D., Logothetis, N.K., and Ugurbil, K. (2007). Spatio-temporal point-spread function of fMRI signal in human gray matter at 7 tesla. Neuroimage 35, 539–552. 10.1016/j.neuroimage.2006.12.030.

52. Turner, R. (2002). How much cortex can a vein drain? Downstream dilution of activation-related cerebral blood oxygenation changes. Neuroimage 16, 1062–1067. 10.1006/nimg.2002.1082.

53. Ciantar, K.G., Farrugia, C., Galdi, P., Scerri, K., Xu, T., and Bajada, C.J. (2022). Geometric effects of volume-to-surface mapping of fMRI data. Brain Struct. Funct. 227, 2457–2464. 10.1007/s00429-022-02536-4.

54. Jeganathan, J., Koussis, N.C., Paton, B., Phogat, R., Pang, J., Mansour L S., Zalesky, A., and Breakspear, M. (2025). Spurious correlations in surface-based functional brain imaging. Imaging Neuroscience 3, imag_a_00478. 10.1162/imag_a_00478.

55. Margulies, D.S., Ghosh, S.S., Goulas, A., Falkiewicz, M., Huntenburg, J.M., Langs, G., Bezgin, G., Eickhoff, S.B., Castellanos, F.X., Petrides, M., et al. (2016). Situating the default-mode network along a principal gradient of macroscale cortical organization. Proc Natl Acad Sci U S A 113, 12574–12579. 10.1073/pnas.1608282113.

56. Yeo, B.T., Krienen, F.M., Sepulcre, J., Sabuncu, M.R., Lashkari, D., Hollinshead, M., Roffman, J.L., Smoller, J.W., Zollei, L., Polimeni, J.R., et al. (2011). The organization of the human cerebral cortex estimated by intrinsic functional connectivity. J Neurophysiol 106, 1125–1165. 10.1152/jn.00338.2011.

57. Engel, S.A., Glover, G.H., and Wandell, B.A. (1997). Retinotopic organization in human visual cortex and the spatial precision of functional MRI. Cerebral Cortex 7, 181–192. 10.1093/cercor/7.2.181.

58. Felleman, D.J., and Van Essen, D.C. (1991). Distributed hierarchical processing in the primate cerebral cortex. Cereb. Cortex 1, 1–47. 10.1093/cercor/1.1.1.

59. Kaas, J.H. (2000). Organizing principles of sensory representations. Novartis Found Symp 228, 188–198; discussion 198-205. 10.1002/0470846631.ch13.

60. Lund, J.S., Yoshioka, T., and Levitt, J.B. (1993). Comparison of intrinsic connectivity in different areas of macaque monkey cerebral cortex. Cereb. Cortex 3, 148–162. 10.1093/cercor/3.2.148.

61. Tanigawa, H., Wang, Q., and Fujita, I. (2005). Organization of Horizontal Axons in the Inferior Temporal Cortex and Primary Visual Cortex of the Macaque Monkey. Cerebral Cortex 15, 1887–1899. 10.1093/cercor/bhi067.

62. Ghazanfar, A.A., and Schroeder, C.E. (2006). Is neocortex essentially multisensory? Trends in Cognitive Sciences 10, 278–285. 10.1016/j.tics.2006.04.008.

63. van den Heuvel, M.P., and Sporns, O. (2011). Rich-club organization of the human connectome. The Journal of Neuroscience 31, 15775–15786. 10.1523/JNEUROSCI.3539-11.2011.

64. van den Heuvel, M.P., and Sporns, O. (2013). Network hubs in the human brain. Trends Cogn. Sci. (Regul. Ed.) 17, 683–696. 10.1016/j.tics.2013.09.012.

65. Bertolero, M.A., Yeo, B.T.T., and D’Esposito, M. (2015). The modular and integrative functional architecture of the human brain. Proceedings of the National Academy of Sciences 112, E6798–E6807. 10.1073/pnas.1510619112.

66. Power, J.D., Schlaggar, B.L., Lessov-Schlaggar, C.N., and Petersen, S.E. (2013). Evidence for Hubs in Human Functional Brain Networks. Neuron 79, 798–813. 10.1016/j.neuron.2013.07.035.

67. Ji, J.L., Spronk, M., Kulkarni, K., Repovš, G., Anticevic, A., and Cole, M.W. (2019). Mapping the human brain’s cortical-subcortical functional network organization. NeuroImage 185, 35–57. 10.1016/j.neuroimage.2018.10.006.

68. Atasoy, S., Donnelly, I., and Pearson, J. (2016). Human brain networks function in connectome-specific harmonic waves. Nature Communications 7, 10340. 10.1038/ncomms10340.

69. Xia, J., Liu, C., Li, J., Meng, Y., Yang, S., Chen, H., and Liao, W. (2024). Decomposing cortical activity through neuronal tracing connectome-eigenmodes in marmosets. Nat. Commun. 15, 2289. 10.1038/s41467-024-46651-8.

70. Wang, Y., Royer, J., Park, B.Y., Vos de Wael, R., Lariviere, S., Tavakol, S., Rodriguez-Cruces, R., Paquola, C., Hong, S.J., Margulies, D.S., et al. (2023). Long-range functional connections mirror and link microarchitectural and cognitive hierarchies in the human brain. Cereb Cortex 33, 1782–1798. 10.1093/cercor/bhac172.

71. Gordon, E.M., Rajesh, A., Chauvin, R.J., Labonte, A., Adeyemo, B., Dworetsky, A., Lynch, C.J., Krimmel, S.R., Cho, P., Wang, A., et al. (2026). Parallel processing chains span cytoarchitectures to organize association cortex. bioRxiv, 2026.04.21.717753. 10.64898/2026.04.21.717753.

72. Mesulam, M. (1998). From sensation to cognition. Brain 121, 1013–1052. 10.1093/brain/121.6.1013.

73. Barbas, H., and Rempel-Clower, N. (1997). Cortical structure predicts the pattern of corticocortical connections. Cerebral Cortex 7, 635–646. 10.1093/cercor/7.7.635.

74. Campagnola, L., Seeman, S.C., Chartrand, T., Kim, L., Hoggarth, A., Gamlin, C., Ito, S., Trinh, J., Davoudian, P., Radaelli, C., et al. (2022). Local connectivity and synaptic dynamics in mouse and human neocortex. Science 375, eabj5861. 10.1126/science.abj5861.

75. Stepanyants, A., Martinez, L.M., Ferecskó, A.S., and Kisvárday, Z.F. (2009). The fractions of short- and long-range connections in the visual cortex. Proceedings of the National Academy of Sciences 106, 3555–3560. 10.1073/pnas.0810390106.

76. Braitenberg, V., and Schüz, A. (1991). Anatomy of the cortex: statistics and geometry (Springer-Verlag).

77. Cossell, L., Iacaruso, M.F., Muir, D.R., Houlton, R., Sader, E.N., Ko, H., Hofer, S.B., and Mrsic-Flogel, T.D. (2015). Functional organization of excitatory synaptic strength in primary visual cortex. Nature 518, 399–403. 10.1038/nature14182.

78. Song, S., Sjöström, P.J., Reigl, M., Nelson, S., and Chklovskii, D.B. (2005). Highly Nonrandom Features of Synaptic Connectivity in Local Cortical Circuits. PLoS Biology 3, e68. 10.1371/journal.pbio.0030068.

79. Yoshimura, Y., Dantzker, J.L.M., and Callaway, E.M. (2005). Excitatory cortical neurons form fine-scale functional networks. Nature 433, 868–873. 10.1038/nature03252.

80. Larkum, M.E., Nevian, T., Sandler, M., Polsky, A., and Schiller, J. (2009). Synaptic integration in tuft dendrites of layer 5 pyramidal neurons: a new unifying principle. Science 325, 756–760. 10.1126/science.1171958.

81. Polsky, A., Mel, B.W., and Schiller, J. (2004). Computational subunits in thin dendrites of pyramidal cells. Nature Neuroscience 7, 621–627. 10.1038/nn1253.

82. Salin, P.A., and Bullier, J. (1995). Corticocortical connections in the visual system: structure and function. Physiol Rev 75, 107–154. 10.1152/physrev.1995.75.1.107.

83. Levitt, J.B., Lewis, D.A., Yoshioka, T., and Lund, J.S. (1993). Topography of pyramidal neuron intrinsic connections in macaque monkey prefrontal cortex (areas 9 and 46). J. Comp. Neurol. 338, 360–376. 10.1002/cne.903380304.

84. Pucak, M.L., Levitt, J.B., Lund, J.S., and Lewis, D.A. (1996). Patterns of intrinsic and associational circuitry in monkey prefrontal cortex. J. Comp. Neurol. 376, 614–630. 10.1002/(SICI)1096-9861(19961223)376:4%3C614::AID-CNE9%3E3.0.CO;2-4.

85. Melchitzky, D.S., Sesack, S.R., Pucak, M.L., and Lewis, D.A. (1998). Synaptic targets of pyramidal neurons providing intrinsic horizontal connections in monkey prefrontal cortex. The Journal of Comparative Neurology 390, 211–224. 10.1002/(SICI)1096-9861(19980112)390:2%3C211::AID-CNE4%3E3.0.CO;2-4.

86. Bajada, C.J., Schreiber, J., and Caspers, S. (2019). Fiber length profiling: A novel approach to structural brain organization. NeuroImage 186, 164–173.10.1016/j.neuroimage.2018.10.070.

87. DeNardo, L.A., Berns, D.S., DeLoach, K., and Luo, L. (2015). Connectivity of mouse somatosensory and prefrontal cortex examined with trans-synaptic tracing. Nat Neurosci 18, 1687–1697. 10.1038/nn.4131.

88. Magrou, L., Joyce, M.K.P., Froudist-Walsh, S., Datta, D., Wang, X.J., Martinez-Trujillo, J., and Arnsten, A.F.T. (2024). The meso-connectomes of mouse, marmoset, and macaque: network organization and the emergence of higher cognition. Cereb Cortex 34, bhae174. 10.1093/cercor/bhae174.

89. Elston, G.N. (2003). Cortex, cognition and the cell: new insights into the pyramidal neuron and prefrontal function. Cereb. Cortex 13, 1124–1138. 10.1093/cercor/bhg093.

90. Fusi, S., Miller, E.K., and Rigotti, M. (2016). Why neurons mix: high dimensionality for higher cognition. Current Opinion in Neurobiology 37, 66–74. 10.1016/j.conb.2016.01.010.

91. Rigotti, M., Barak, O., Warden, M.R., Wang, X.J., Daw, N.D., Miller, E.K., and Fusi, S. (2013). The importance of mixed selectivity in complex cognitive tasks. Nature 497, 585–590. 10.1038/nature12160.

92. Kanari, L., Shi, Y., Arnaudon, A., Barros-Zulaica, N., Benavides-Piccione, R., Coggan, J.S., DeFelipe, J., Hess, K., Mansvelder, H.D., Mertens, E.J., et al. (2025). Of mice and men: dendritic architecture differentiates human from mouse neuronal networks. iScience 28, 112928. 10.1016/j.isci.2025.112928.

93. Rikhye, R.V., Gilra, A., and Halassa, M.M. (2018). Thalamic regulation of switching between cortical representations enables cognitive flexibility. Nature Neuroscience 21, 1753–1763. 10.1038/s41593-018-0269-z.

94. Schmitt, L.I., Wimmer, R.D., Nakajima, M., Happ, M., Mofakham, S., and Halassa, M.M. (2017). Thalamic amplification of cortical connectivity sustains attentional control. Nature 545, 219–223. 10.1038/nature22073.

95. García-Cabezas, M.Á., Zikopoulos, B., and Barbas, H. (2019). The Structural Model: a theory linking connections, plasticity, pathology, development and evolution of the cerebral cortex. Brain Structure and Function 224, 985–1008. 10.1007/s00429-019-01841-9.

96. Luppino, G., Matelli, M., Camarda, R., and Rizzolatti, G. (1993). Corticocortical connections of area F3 (SMA-proper) and area F6 (pre-SMA) in the macaque monkey. Journal of Comparative Neurology 338, 114–140. 10.1002/cne.903380109.

97. Luppino, G., Rozzi, S., Calzavara, R., and Matelli, M. (2003). Prefrontal and agranular cingulate projections to the dorsal premotor areas F2 and F7 in the macaque monkey. European Journal of Neuroscience 17, 559–578. 10.1046/j.1460-9568.2003.02476.x.

98. Rizzolatti, G., and Luppino, G. (2001). The Cortical Motor System. Neuron 31, 889–901. 10.1016/S0896-6273(01)00423-8.

99. Matelli, M., and Luppino, G. (1996). Thalamic input to mesial and superior area 6 in the macaque monkey. J. Comp. Neurol. 372, 59–87. 10.1002/(SICI)1096-9861(19960812)372:1%3C59::AID-CNE6%3E3.0.CO;2-L.

100. Fuster, J.M. (2001). The prefrontal cortex—an update: time is of the essence. Neuron 30, 319–333. 10.1016/s0896-6273(01)00285-9.

101. Jones, E.G., and Powell, T.P.S. (1970). An anatomical study of converging sensory pathways within the cerebral cortex of the monkey. Brain 93, 793–820. 10.1093/brain/93.4.793.

102. Nee, D.E., and D’Esposito, M. (2016). The hierarchical organization of the lateral prefrontal cortex. eLife 5, e12112. 10.7554/eLife.12112.

103. Badre, D., and Nee, D.E. (2018). Frontal Cortex and the Hierarchical Control of Behavior. Trends in Cognitive Sciences 22, 170–188. 10.1016/j.tics.2017.11.005.

104. Petrides, M., and Pandya, D.N. (1984). Projections to the frontal cortex from the posterior parietal region in the rhesus monkey. Journal of Comparative Neurology 228, 105–116. 10.1002/cne.902280110.

105. Cavada, C., and Goldman-Rakic, P.S. (1989). Posterior parietal cortex in rhesus monkey: II. Evidence for segregated corticocortical networks linking sensory and limbic areas with the frontal lobe. Journal of Comparative Neurology 287, 422–445. 10.1002/cne.902870403.

106. Schmidt, M.E., Aganj, I., Stockmann, J., Bilgic, B., Chang, Y., Hoge, W.S., Kirilina, E., Weiskopf, N., and Nasr, S. (2025). Unraveling the mesoscale resting-state functional connectivity of ocular dominance columns in humans using high-resolution functional MRI. Communications Biology 8.10.1038/s42003-025-09108-2.

107. Murphy, K., Birn, R.M., Handwerker, D.A., Jones, T.B., and Bandettini, P.A. (2009). The impact of global signal regression on resting state correlations: Are anti-correlated networks introduced? NeuroImage 44, 893–905. 10.1016/j.neuroimage.2008.09.036.

108. Murphy, K., and Fox, M.D. (2017). Towards a consensus regarding global signal regression for resting state functional connectivity MRI. NeuroImage 154, 169–173. 10.1016/j.neuroimage.2016.11.052.

109. Saad, Z.S., Gotts, S.J., Murphy, K., Chen, G., Jo, H.J., Martin, A., and Cox, R.W. (2012). Trouble at Rest: How Correlation Patterns and Group Differences Become Distorted After Global Signal Regression. Brain Connectivity 2, 25–32. 10.1089/brain.2012.0080.

110. Faskowitz, J., Bandettini, P.A., and Gonzalez-Castillo, J. (2026). Mapping high-amplitude fMRI edge time series events across time and space. Imaging Neuroscience 4. 10.1162/IMAG.a.1126.

111. Fonov, V., Evans, A.C., Botteron, K., Almli, C.R., McKinstry, R.C., and Collins,D.L. (2011). Unbiased average age-appropriate atlases for pediatric studies. NeuroImage 54, 313–327. 10.1016/j.neuroimage.2010.07.033.

112. Glasser, M.F., Coalson, T.S., Robinson, E.C., Hacker, C.D., Harwell, J., Yacoub, E., Ugurbil, K., Andersson, J., Beckmann, C.F., Jenkinson, M., et al. (2016). A multimodal parcellation of human cerebral cortex. Nature 536, 171–178. 10.1038/nature18933.

113. Jakab, A., Blanc, R., Berényi, E.L., and Székely, G. (2012). Generation of individualized thalamus target maps by using statistical shape models and thalamocortical tractography. American Journal of Neuroradiology 33, 2110–2116. 10.3174/ajnr.A3140.

114. Krauth, A., Blanc, R., Poveda, A., Jeanmonod, D., Morel, A., and Székely, G. (2010). A mean three-dimensional atlas of the human thalamus: Generation from multiple histological data. NeuroImage 49, 2053–2062. 10.1016/j.neuroimage.2009.10.042.

115. Morel, A. (2007). Stereotactic atlas of the human thalamus and basal ganglia (Informa Healthcare) 10.3109/9781420016796.

116. Paquola, C., Royer, J., Lewis, L.B., Lepage, C., Glatard, T., Wagstyl, K., DeKraker, J., Toussaint, P.J., Valk, S.L., Collins, L., et al. (2021). The BigBrainWarp toolbox for integration of BigBrain 3D histology with multimodal neuroimaging. Elife 10. 10.7554/eLife.70119.

117. Huck, J., Wanner, Y., Fan, A.P., Jäger, A.-T., Grahl, S., Schneider, U., Villringer, A., Steele, C.J., Tardif, C.L., Bazin, P.-L., et al. (2019). High resolution atlas of the venous brain vasculature from 7 T quantitative susceptibility maps. Brain Structure and Function 224, 2467–2485. 10.1007/s00429-019-01919-4.

118. Esteban, O., Markiewicz, C.J., Blair, R.W., Moodie, C.A., Isik, A.I., Erramuzpe, A., Kent, J.D., Goncalves, M., DuPre, E., Snyder, M., et al. (2019). fMRIPrep: a robust preprocessing pipeline for functional MRI. Nature Methods 16, 111–116. 10.1038/s41592-018-0235-4.

119. DuPre, E., Salo, T., Ahmed, Z., Bandettini, P., Bottenhorn, K., Caballero-Gaudes, C., Dowdle, L., Gonzalez-Castillo, J., Heunis, S., Kundu, P., et al. (2021). TE-dependent analysis of multi-echo fMRI with tedana. Journal of Open Source Software 6, 3669. 10.21105/joss.03669.

120. Robinson, E.C., Garcia, K., Glasser, M.F., Chen, Z., Coalson, T.S., Makropoulos, A., Bozek, J., Wright, R., Schuh, A., Webster, M., et al. (2018). Multimodal surface matching with higher-order smoothness constraints. NeuroImage 167, 453–465. 10.1016/j.neuroimage.2017.10.037.

121. Vos de Wael, R., Benkarim, O., Paquola, C., Lariviere, S., Royer, J., Tavakol, S., Xu, T., Hong, S.-J., Langs, G., Valk, S., et al. (2020). BrainSpace: a toolbox for the analysis of macroscale gradients in neuroimaging and connectomics datasets. Communications Biology 3. 10.1038/s42003-020-0794-7.

122. Ito, T., Hearne, L.J., and Cole, M.W. (2020). A cortical hierarchy of localized and distributed processes revealed via dissociation of task activations, connectivity changes, and intrinsic timescales. NeuroImage 221, 117141. 10.1016/j.neuroimage.2020.117141.

123. Glasser, M.F., Sotiropoulos, S.N., Wilson, J.A., Coalson, T.S., Fischl, B., Andersson, J.L., Xu, J., Jbabdi, S., Webster, M., Polimeni, J.R., et al. (2013). The minimal preprocessing pipelines for the Human Connectome Project. NeuroImage 80, 105–124. 10.1016/j.neuroimage.2013.04.127.

124. Glasser, M.F., Coalson, T.S., Bijsterbosch, J.D., Harrison, S.J., Harms, M.P., Anticevic, A., Van Essen, D.C., and Smith, S.M. (2018). Using temporal ICA to selectively remove global noise while preserving global signal in functional MRI data. NeuroImage 181, 692–717. 10.1016/j.neuroimage.2018.04.076.

125. Kundu, P., Inati, S.J., Evans, J.W., Luh, W.-M., and Bandettini, P.A. (2012). Differentiating BOLD and non-BOLD signals in fMRI time series using multi-echo EPI. NeuroImage 60, 1759–1770. 10.1016/j.neuroimage.2011.12.028.

126. Efron, B. (2004). Large-Scale Simultaneous Hypothesis Testing. Journal of the American Statistical Association 99, 96–104. 10.1198/016214504000000089.

127. Schaefer, A., Kong, R., Gordon, E.M., Laumann, T.O., Zuo, X.N., Holmes, A.J., Eickhoff, S.B., and Yeo, B.T.T. (2018). Local-Global Parcellation of the Human Cerebral Cortex from Intrinsic Functional Connectivity MRI. Cereb Cortex 28, 3095–3114. 10.1093/cercor/bhx179.

128. Hill, J., Inder, T., Neil, J., Dierker, D., Harwell, J., and Van Essen, D. (2010). Similar patterns of cortical expansion during human development and evolution. Proceedings of the National Academy of Sciences 107, 13135–13140. 10.1073/pnas.1001229107.

129. Xu, T., Nenning, K.-H., Schwartz, E., Hong, S.-J., Vogelstein, J.T., Goulas, A., Fair, D.A., Schroeder, C.E., Margulies, D.S., Smallwood, J., et al. (2020). Cross-species functional alignment reveals evolutionary hierarchy within the connectome. Neuroimage 223, 117346. 10.1016/j.neuroimage.2020.117346.

130. Tomczak, M., and Tomczak, E. (2014). The need to report effect size estimates revisited. An overview of some recommended measures of effect size. Trends in Sport Sciences 21, 19–25.

## SUPPLEMENTAL REFERENCES

1. Bernier, M., Cunnane, S.C., and Whittingstall, K. (2018). The morphology of the human cerebrovascular system. Human Brain Mapping 39, 4962–4975. 10.1002/hbm.24337.

2. Weber, B., Keller, A.L., Reichold, J., and Logothetis, N.K. (2008). The Microvascular System of the Striate and Extrastriate Visual Cortex of the Macaque. Cerebral Cortex 18, 2318–2330. 10.1093/cercor/bhm259.

3. Huck, J., Wanner, Y., Fan, A.P., Jäger, A.-T., Grahl, S., Schneider, U., Villringer, A., Steele, C.J., Tardif, C.L., Bazin, P.-L., et al. (2019). High resolution atlas of the venous brain vasculature from 7 T quantitative susceptibility maps. Brain Structure and Function 224, 2467–2485. 10.1007/s00429-019-01919-4.

4. Luppino, G., Matelli, M., Camarda, R., and Rizzolatti, G. (1993). Corticocortical connections of area F3 (SMA-proper) and area F6 (pre-SMA) in the macaque monkey. Journal of Comparative Neurology 338, 114–140. 10.1002/cne.903380109.

5. Luppino, G., Rozzi, S., Calzavara, R., and Matelli, M. (2003). Prefrontal and agranular cingulate projections to the dorsal premotor areas F2 and F7 in the macaque monkey. European Journal of Neuroscience 17, 559–578. 10.1046/j.1460-9568.2003.02476.x.

6. Rizzolatti, G., and Luppino, G. (2001). The Cortical Motor System. Neuron 31, 889–901. 10.1016/S0896-6273(01)00423-8.

7. Pang, J.C., Aquino, K.M., Oldehinkel, M., Robinson, P.A., Fulcher, B.D., Breakspear, M., and Fornito, A. (2023). Geometric constraints on human brain function. Nature 618, 566–574. 10.1038/s41586-023-06098-1.

8. Normand, F., Gajwani, M., Cao, T., Cruddas, J., Sangchooli, A., Oldham, S., Holmes, A., Robinson, P.A., Pang, J.C., and Fornito, A. (2026). Geometric constraints on the architecture of mammalian cortical connectomes. Cell 189, 5283–5303.e17. 10.1016/j.cell.2026.05.048.

9. Margulies, D.S., Ghosh, S.S., Goulas, A., Falkiewicz, M., Huntenburg, J.M., Langs, G., Bezgin, G., Eickhoff, S.B., Castellanos, F.X., Petrides, M., et al. (2016). Situating the default-mode network along a principal gradient of macroscale cortical organization. Proc Natl Acad Sci U S A 113, 12574–12579. 10.1073/pnas.1608282113.

10. Yeo, B.T., Krienen, F.M., Sepulcre, J., Sabuncu, M.R., Lashkari, D., Hollinshead, M., Roffman, J.L., Smoller, J.W., Zollei, L., Polimeni, J.R., et al. (2011). The organization of the human cerebral cortex estimated by intrinsic functional connectivity. J Neurophysiol 106, 1125–1165. 10.1152/jn.00338.2011.

11. Lund, J.S., Yoshioka, T., and Levitt, J.B. (1993). Comparison of intrinsic connectivity in different areas of macaque monkey cerebral cortex. Cereb. Cortex 3, 148–162. 10.1093/cercor/3.2.148.

12. Levitt, J.B., Lewis, D.A., Yoshioka, T., and Lund, J.S. (1993). Topography of pyramidal neuron intrinsic connections in macaque monkey prefrontal cortex (areas 9 and 46). J. Comp. Neurol. 338, 360–376. 10.1002/cne.903380304.

13. Pucak, M.L., Levitt, J.B., Lund, J.S., and Lewis, D.A. (1996). Patterns of intrinsic and associational circuitry in monkey prefrontal cortex. J. Comp. Neurol. 376, 614–630. 10.1002/(SICI)1096-9861(19961223)376:4%3C614::AID-CNE9%3E3.0.CO;2-4.

14. Huntley, G.W., and Jones, E.G. (1991). Relationship of intrinsic connections to forelimb movement representations in monkey motor cortex: a correlative anatomic and physiological study. Journal of Neurophysiology 66, 390–413. 10.1152/jn.1991.66.2.390.

15. Burman, K.J., Bakola, S., Richardson, K.E., Reser, D.H., and Rosa, M.G.P. (2014). Patterns of afferent input to the caudal and rostral areas of the dorsal premotor cortex (6DC and 6DR) in the marmoset monkey. Journal of Comparative Neurology 522, 3683–3716. 10.1002/cne.23633.

16. Glasser, M.F., and Van Essen, D.C. (2011). Mapping human cortical areas in vivo based on myelin content as revealed by T1- and T2-weighted MRI. The Journal of Neuroscience 31, 11597–11616. 10.1523/jneurosci.2180-11.2011.

17. Burt, J.B., Demirtas, M., Eckner, W.J., Navejar, N.M., Ji, J.L., Martin, W.J., Bernacchia, A., Anticevic, A., and Murray, J.D. (2018). Hierarchy of transcriptomic specialization across human cortex captured by structural neuroimaging topography. Nat. Neurosci. 21, 1251–1259. 10.1038/s41593-018-0195-0.

18. Sydnor, V.J., Larsen, B., Bassett, D.S., Alexander-Bloch, A., Fair, D.A., Liston, C., Mackey, A.P., Milham, M.P., Pines, A., Roalf, D.R., et al. (2021). Neurodevelopment of the association cortices: Patterns, mechanisms, and implications for psychopathology. Neuron 109, 2820–2846. 10.1016/j.neuron.2021.06.016.

19. Hill, J., Inder, T., Neil, J., Dierker, D., Harwell, J., and Van Essen, D. (2010). Similar patterns of cortical expansion during human development and evolution. Proceedings of the National Academy of Sciences 107, 13135–13140. 10.1073/pnas.1001229107.

20. Reardon, P.K., Seidlitz, J., Vandekar, S., Liu, S., Patel, R., Park, M.T.M., Alexander-Bloch, A., Clasen, L.S., Blumenthal, J.D., Lalonde, F.M., et al. (2018). Normative brain size variation and brain shape diversity in humans. Science 360, 1222–1227. 10.1126/science.aar2578.

21. Felleman, D.J., and Van Essen, D.C. (1991). Distributed hierarchical processing in the primate cerebral cortex. Cereb. Cortex 1, 1–47. 10.1093/cercor/1.1.1.

22. Iwamura, Y. (1998). Hierarchical somatosensory processing. Current Opinion in Neurobiology 8, 522–528. 10.1016/s0959-4388(98)80041-x.

23. Kaas, J.H., and Hackett, T.A. (2000). Subdivisions of auditory cortex and processing streams in primates. Proc. Natl. Acad. Sci. 97, 11793–11799.10.1073/pnas.97.22.11793.

24. Rauschecker, J.P., and Scott, S.K. (2009). Maps and streams in the auditory cortex: nonhuman primates illuminate human speech processing. Nature Neuroscience 12, 718–724. 10.1038/nn.2331.

25. Hamilton, L.S., Oganian, Y., Hall, J., and Chang, E.F. (2021). Parallel and distributed encoding of speech across human auditory cortex. Cell 184, 4626–4639.e13. 10.1016/j.cell.2021.07.019.

26. Kravitz, D.J., Saleem, K.S., Baker, C.I., and Mishkin, M. (2011). A new neural framework for visuospatial processing. Nat Rev Neurosci 12, 217–230. 10.1038/nrn3008.

27. Seltzer, B., and Pandya, D.N. (2009). Posterior cingulate and retrosplenial cortex connections of the caudal superior temporal region in the rhesus monkey. Experimental Brain Research 195, 325–334. 10.1007/s00221-009-1795-4.

28. Parvizi, J., Van Hoesen, G.W., Buckwalter, J., and Damasio, A. (2006). Neural connections of the posteromedial cortex in the macaque. Proceedings of the National Academy of Sciences 103, 1563–1568. 10.1073/pnas.0507729103.

29. Hwang, K., Bertolero, M.A., Liu, W.B., and D’Esposito, M. (2017). The Human Thalamus Is an Integrative Hub for Functional Brain Networks. The Journal of Neuroscience 37, 5594–5607. 10.1523/JNEUROSCI.0067-17.2017.

30. Kumar, V.J., Van Oort, E., Scheffler, K., Beckmann, C.F., and Grodd, W. (2017). Functional anatomy of the human thalamus at rest. Neuroimage 147, 678–691. 10.1016/j.neuroimage.2016.12.071.

31. Jones, E.G. (2001). The thalamic matrix and thalamocortical synchrony. Trends in Neurosciences 24, 595–601. 10.1016/s0166-2236(00)01922-6.

32. Smith, Y., Raju, D.V., Pare, J.-F., and Sidibe, M. (2004). The thalamostriatal system: a highly specific network of the basal ganglia circuitry. Trends in Neurosciences 27, 520–527. 10.1016/j.tins.2004.07.004.

33. Kalil, K. (1981). Projections of the cerebellar and dorsal column nuclei upon the thalamus of the rhesus monkey. Journal of Comparative Neurology 195, 25–50. 10.1002/cne.901950105.

34. Hoover, J.E., and Strick, P.L. (1999). The Organization of Cerebellar and Basal Ganglia Outputs to Primary Motor Cortex as Revealed by Retrograde Transneuronal Transport of Herpes Simplex Virus Type 1. The Journal of Neuroscience 19, 1446–1463. 10.1523/JNEUROSCI.19-04-01446.1999.

35. Gordon, E.M., Chauvin, R.J., Van, A.N., Rajesh, A., Nielsen, A., Newbold, D.J., Lynch, C.J., Seider, N.A., Krimmel, S.R., Scheidter, K.M., et al. (2023). A somatocognitive action network alternates with effector regions in motor cortex. Nature 617, 351–359. 10.1038/s41586-023-05964-2.

36. Ladwig, Z., Kermani, K.Z., Park, Y., Housteau, E., Dworetsky, A., Labora, N., Hernandez, J.J., Dorn, M., Smith, D.M., Nee, D.E., et al. (2026). Precision fMRI reveals densely interdigitated network patches with conserved motifs in the lateral prefrontal cortex. Neuron 114, 3499–3516.e8. 10.1016/j.neuron.2026.04.011.

37. Glasser, M.F., Coalson, T.S., Bijsterbosch, J.D., Harrison, S.J., Harms, M.P., Anticevic, A., Van Essen, D.C., and Smith, S.M. (2018). Using temporal ICA to selectively remove global noise while preserving global signal in functional MRI data. NeuroImage 181, 692–717. 10.1016/j.neuroimage.2018.04.076.

38. Rosanova, M., Casali, A., Bellina, V., Resta, F., Mariotti, M., and Massimini, M. (2009). Natural frequencies of human corticothalamic circuits. J Neurosci 29, 7679–7685. 10.1523/JNEUROSCI.0445-09.2009.

39. Alexander, G.E., DeLong, M.R., and Strick, P.L. (1986). Parallel organization of functionally segregated circuits linking basal ganglia and cortex. Annu Rev Neurosci 9, 357–381. 10.1146/annurev.ne.09.030186.002041.

40. Matelli, M., and Luppino, G. (1996). Thalamic input to mesial and superior area 6 in the macaque monkey. J. Comp. Neurol. 372, 59–87. 10.1002/(SICI)1096-9861(19960812)372:1%3C59::AID-CNE6%3E3.0.CO;2-L.

41. Lynch, J.C., Hoover, J.E., and Strick, P.L. (1994). Input to the primate frontal eye field from the substantia nigra, superior colliculus, and dentate nucleus demonstrated by transneuronal transport. Experimental Brain Research 100, 181–186. 10.1007/BF00227293.

42. McFarland, N.R., and Haber, S.N. (2002). Thalamic Relay Nuclei of the Basal Ganglia Form Both Reciprocal and Nonreciprocal Cortical Connections,Linking Multiple Frontal Cortical Areas. The Journal of Neuroscience 22, 8117–8132. 10.1523/JNEUROSCI.22-18-08117.2002.

44. Ray, J.P., and Price, J.L. (1993). The organization of projections from the mediodorsal nucleus of the thalamus to orbital and medial prefrontal cortex in macaque monkeys. Journal of Comparative Neurology 337, 1–31. 10.1002/cne.903370102.

45. Vogt, B.A., Pandya, D.N., and Rosene, D.L. (1987). Cingulate cortex of the rhesus monkey: I. Cytoarchitecture and thalamic afferents. J. Comp. Neurol. 262, 256–270. 10.1002/cne.902620207.

46. Sherman, S.M. (2016). Thalamus plays a central role in ongoing cortical functioning. Nat Neurosci 19, 533–541. 10.1038/nn.4269.

47. Kaas, J.H., and Lyon, D.C. (2007). Pulvinar contributions to the dorsal and ventral streams of visual processing in primates. Brain Research Reviews 55, 285–296. 10.1016/j.brainresrev.2007.02.008.

48. Schmahmann, J.D., and Pandya, D.N. (1990). Anatomical investigation of projections from thalamus to posterior parietal cortex in the rhesus monkey: A WGA-HRP and fluorescent tracer study. Journal of Comparative Neurology 295, 299–326. 10.1002/cne.902950212.

49. Mufson, E.J., and Mesulam, M.M. (1984). Thalamic connections of the insula in the rhesus monkey and comments on the paralimbic connectivity of the medial pulvinar nucleus. Journal of Comparative Neurology 227, 109–120. 10.1002/cne.902270112.

50. Clower, D.M., Dum, R.P., and Strick, P.L. (2005). Basal ganglia and cerebellar inputs to ‘AIP.’ Cereb. Cortex 15, 913–920. 10.1093/cercor/bhh190.

51. Aggleton, J.P., O’Mara, S.M., Vann, S.D., Wright, N.F., Tsanov, M., and Erichsen, J.T. (2010). Hippocampal-anterior thalamic pathways for memory: uncovering a network of direct and indirect actions. Eur J Neurosci 31, 2292–2307. 10.1111/j.1460-9568.2010.07251.x.

52. Insausti, R., Amaral, D.G., and Cowan, W.M. (1987). The entorhinal cortex of the monkey: III. Subcortical afferents. Journal of Comparative Neurology 264, 396–408. 10.1002/cne.902640307.

53. Rovó, Z., Ulbert, I., and Acsády, L. (2012). Drivers of the primate thalamus. The Journal of Neuroscience 32, 17894–17908. 10.1523/JNEUROSCI.2815-12.2012.

54. Middleton, F.A., and Strick, P.L. (1996). The temporal lobe is a target of output from the basal ganglia. Proc. Natl. Acad. Sci. 93, 8683–8687. 10.1073/pnas.93.16.8683.

55. Krauth, A., Blanc, R., Poveda, A., Jeanmonod, D., Morel, A., and Székely, G. (2010). A mean three-dimensional atlas of the human thalamus: Generation from multiple histological data. NeuroImage 49, 2053–2062. 10.1016/j.neuroimage.2009.10.042.

56. Morel, A. (2007). Stereotactic atlas of the human thalamus and basal ganglia (Informa Healthcare) 10.3109/9781420016796.

57. Jakab, A., Blanc, R., Berényi, E.L., and Székely, G. (2012). Generation of individualized thalamus target maps by using statistical shape models and thalamocortical tractography. American Journal of Neuroradiology 33, 2110–2116. 10.3174/ajnr.A3140.

58. Tomczak, M., and Tomczak, E. (2014). The need to report effect size estimates revisited. An overview of some recommended measures of effect size. Trends in Sport Sciences 21, 19–25.

59. Xu, T., Nenning, K.-H., Schwartz, E., Hong, S.-J., Vogelstein, J.T., Goulas, A., Fair, D.A., Schroeder, C.E., Margulies, D.S., Smallwood, J., et al. (2020). Cross-species functional alignment reveals evolutionary hierarchy within the connectome. Neuroimage 223, 117346. 10.1016/j.neuroimage.2020.117346.

60. Markello, R.D., Hansen, J.Y., Liu, Z.-Q., Bazinet, V., Shafiei, G., Suarez, L.E., Blostein, N., Seidlitz, J., Baillet, S., Satterthwaite, T.D., et al. (2022). neuromaps: structural and functional interpretation of brain maps. Nat. Methods 19, 1472–1479. 10.1038/s41592-022-01625-w.

